## Supplementary Information for "Combining two Genetic Sexing Strains allows sorting of non-transgenic males for *Aedes* genetic control"

**List of Supplementary Tables:**

Supplementary Table 1

Supplementary Table 2

Supplementary Table 3

**List of Supplementary Data:**

Supplementary Data 1

**List of Supplementary Data found in a separate document:**

Supplementary Data 2

Supplementary Data 3

Supplementary Data 4

**Supplementary Table 1: Summary and significance of statistical tests performed.** More details can be found in **Supplementary Data 2**.

| **Sex ratio (%)** | | |
|---|---|---|
| **Pairwise comparison** | **Estimate (SE) *** | ***p*-value** |
| Aaeg-M *vs.* Bra (WT) | -0.06 (0.09) | 0.903 |
| Aaeg-m *vs.* Bra (WT) | -0.01 (0.08) | 0.995 |
| Aaeg-CS *vs.* Bra (WT) | 0.05 (0.09) | 0.935 |
| Aal-M *vs.* BiA (WT) | -0.09 (0.11) | 0.847 |
| Aal-m *vs.* BiA (WT) | -0.09 (0.12) | 0.858 |
| Aal-CS *vs.* BiA (WT) | -0.03 (0.11) | 0.990 |
| **Fecundity (%)** | | |
| **Pairwise comparison** | **Estimate (SE) **** | ***p*-value** |
| Aaeg-M *vs.* Bra (WT) | -0.14 (0.06) | 0.075 |
| Aaeg-m *vs.* Bra (WT) | 0.11 (0.06) | 0.188 |
| Aaeg-CS *vs.* Bra (WT) | 0.74 (0.05) | < 0.001 *** |
| Aal-M *vs.* BiA (WT) | 0.59 (0.04) | < 0.001 *** |
| Aal-m *vs.* BiA (WT) | 0.04 (0.05) | 0.813 |
| Aal-CS *vs.* BiA (WT) | -0.06 (0.05) | 0.632 |
| **Egg hatching rate (%)** | | |
| **Pairwise comparison** | **Estimate (SE) *** | ***p*-value** |
| Aaeg-M *vs.* Bra (WT) | -0.17 (0.10) | 0.334 |
| Aaeg-m *vs.* Bra (WT) | 0.99 (0.11) | < 0.001 *** |
| Aaeg-CS *vs.* Bra (WT) | 0.83 (0.10) | < 0.001 *** |
| Aal-M *vs.* BiA (WT) | 0.15 (0.04) | 0.002 ** |
| Aal-m *vs.* BiA (WT) | -0.03 (0.06) | 0.9646 |
| Aal-CS *vs.* BiA (WT) | 0.10 (0.04) | 0.046 * |
| **Larva to adult survival (%)** | | |
| **Pairwise comparison** | **Estimate (SE)** | ***p*-value** |
| Aaeg-M *vs.* Bra (WT) | -4.87 (3.83) | 0.592 |

| | | |
|---|---|---|
| Aaeg-m *vs*. Bra (WT) | 1.00 (4.22) | 0.996 |
| Aaeg-CS *vs*. Bra (WT) | 3.75 (4.22) | 0.831 |
| Aal-M *vs*. BiA (WT) | 18.13 (4.00) | 0.003 ** |
| Aal-m *vs*. BiA (WT) | 16.63 (4.00) | 0.006 ** |
| Aal-CS *vs*. BiA (WT) | 5.62 (4.00) | 0.520 |
| **Male competitiveness (%)** | | |
| **Difference from expected** | **Estimate (SE) *** | ***p*-value** |
| Aaeg-M *vs*. Bra (WT) | 0.04 (0.04) | 0.318 |
| Aaeg-M *vs*. Aaeg-CS | 0.08 (0.04) | 0.057 |
| Aal-M *vs*. BiA (WT) | -0.42 (0.07) | < 0.001 |
| Aal-CS *vs*. BiA (WT) | 0.05 (0.05) | 0.34 |
| **Flight test (%)** | | |
| **Pairwise comparison** | **Estimate (SE) *** | ***p*-value** |
| Aaeg-M *vs*. Bra (WT) | -0.22 (0.19) | 0.461 |
| Aaeg-CS *vs*. Bra (WT) | -0.05 (0.19) | 0.966 |
| Aal-M *vs*. BiA (WT) | -0.36 (0.18) | 0.103 |
| Aal-CS *vs*. BiA (WT) | 0.35 (0.19) | 0.148 |
| **Male survival** | | |
| **Pairwise comparison (7 days)** | **Estimate (SE)** | ***p*-value** |
| Aaeg-M *vs*. Bra (WT) | 0.00 (0.01) | 0.875 |
| Aaeg-CS *vs*. Bra (WT) | 0.01 (0.01) | 0.226 |
| Aal-M *vs*. BiA (WT) | 0.00 (0.01) | 1.000 |
| Aal-CS *vs*. BiA (WT) | -0.01 (0.01) | 0.802 |
| **Pairwise comparison (14 days)** | **Estimate (SE)** | ***p*-value** |
| Aaeg-M *vs*. Bra (WT) | 0.00 (0.01) | 0.997 |
| Aaeg-CS *vs*. Bra (WT) | 0.01 (0.01) | 0.831 |
| Aal-M *vs*. BiA (WT) | 0.00 (0.01) | 1.000 |

| | | |
|---|---|---|
| Aal-CS *vs.* BiA (WT) | 0.00 (0.01) | 0.996 |

* estimated difference in logit of individuals

**Supplementary Table 2: Sequences of markers marking out and within the 63Mbp with high male-female genetic differentiation in *Ae. Aegypti.***

| | |
|---|---|
| 137972_left_side_Xspe_163341238 | GGAAAAAGGGTTTAAAAAATAACTGCATCAACCAACTATGCAAAAGCCAACTAGTACACACAGCTGACGTACAAACACTGCTGGAGACACTCAGCTTATTTTCCGCTGCAGGGTTATGTCATTGCAGGCAACACAT |
| 220255_Xspe_166482560 | TCGACCTTGGCTCGGAGTTTCCGTCGCGTTTGACCCAAAACCGGCGTACATAGGAAAGCATCCGAACCAAGGTGGTCAGATTCGTTGTAACCGGCGATACGGTCCATTTCGGCAATCGCCGAAACCGGTGTTTTGG |
| 31870_right_side_Xspe_174626199 | CAGCCAATCCGAAAAGCAAACACAAAGCCTGCCCGAGTTCGTCTGCTTTCCCGATGGGGTGCAATAGAGCAAAAGTTTCTCTTTTGTCTATGCGGTGATATTTTTGCGAAACTCGCGATAGAAAGCCTTTCCTGTG |

**Supplementary Table 3: Cumulative number of fluorescent and negative males and females between generations 3 and 6 in 11 *Ae. albopictus* lines carrying an M-linked transgene.** Underlined numbers are observations that may correspond to recombination events or to the presence of additional, less sex-linked transgene insertions. Lines showing such events were thus discarded. The selected M-linked line (Aal-M) is the one named "4.4Y" in the table.

| Line | Positive males | Negative males | Positive females | Negative females | Total |
|---|---|---|---|---|---|
| **SM1** | 181 | 0 | <u>3</u> | 316 | 500 |
| **SM2** | 53 | 0 | <u>34</u> | 25 | 112 |
| **SM3** | 301 | 0 | 0 | 625 | 926 |
| **SM5** | 259 | <u>11</u> | <u>26</u> | 384 | 680 |
| **SM6** | 22 | 0 | <u>1</u> | 17 | 40 |
| **SM7** | 81 | 0 | <u>30</u> | 63 | 174 |
| **SM11** | 79 | 0 | <u>11</u> | 51 | 141 |
| **SM12** | 647 | 0 | 0 | 623 | 1270 |
| **1.2R** | 666 | 0 | 0 | 569 | 1235 |
| **4.4Y = Aal-M** | 905 | 0 | 0 | 813 | 1718 |
| **5.2GR** | 403 | 0 | 0 | 310 | 713 |

**Supplementary Data 1: 1239bp *Ae. albopictus* genomic sequence flanking the Aal-M transposon and 1647 bp *Ae. albopictus* genomic sequence flanking the Aal-m transposon.** Complete flanking sequences and detailed explanations can be found in Supplementary Data 4.

Aal-M:

TTAACCACGAATTCTTTTAGGGATTCCTCCAGAGTCCTACCAGAAATTCATTTAGGGATAT
CTCCAGGAAATCCTTGAGAAATTCGTCCAGGGATCCAGGAATTCCTCCAGAGTATCCTTC
AGGCATTCCGGTCAGAAATCTTTAAGGGGTTCATCCTCGGATTTGTTCAGCAATCCATCC
ATGAATTCATCAAGAATACGCTCTGCTGCTTCTGCAAATGAATTCTTTCAGAATTTCGTCC
TGTGATTCTTTCAGGGATATTTATTATTGAGATTATTTTTTCCGAGAAATCCTTCAAGAAG
TTATCCATTGATTTTTTTCAAGAACTCCTCCACAAGTTTTTTCAGGGATGTTTAGAGAGTG
TCCTTCAATGATTCTTCCAAAAAATAGTCAAGTCACTCGTTTAGAAATTCCCTCGGCAAT
ACCTCAAGGATTCCCTTCAGCGATTGTTTTAGGACTTATCCTAGGGGTTTGTTCAAATATT
ACATAACGCTAAGGGGGAGAGAGGGGGTCTAGCTCTGTGTTACGATTCATACAACATTC
TTAAAGTTTTCATATAAAATTTGTTACGTGGGGGAGGGAGGGGGTCTTAAATTGTGAAAT
TTTGCGTTACGTAATATTTGAAAGAACCTTAGGATTTCTTCAGGGATTCCACCAGGGACC
TCCTTTCGAGTTTTTTTTTTGTAAAGATTCTTTCAGAAATTTCTCCAAGAAATTGCCTAAA
AATTTTCCAGGAATCTCTTAAGCAGTTCCTTCAGGGATTGTATCAGGAATTCCTATACCTA
ATAAATCTTTCAGGGATTCTTCAAAGCAACACGCATGTTACAGAAGTGTCACGGCAGTG
CGCAAGGTTTTGTTTTCGAAAAAGTCACTGTGACCTACTTTATACAAATAAGTACCTACA
TGCTTGAATCGCTTACAAACANTTTTTCCAGGGAATCTATCAGGAATTCACCCAGAGAA
AACTTTTTTGGAATTTTCCCAGGGTTTGTCCGACAATTACCATAAGAAAAAGTATCAGGA
GTTCCCCTAGGAAWTCCTTCTCAAATTCCTTCATTTTGAATATATTCTAGCGATTCGTCTA
GGAATCCCTTCAGGTGAATTTCAGGATTTTTTCAAGAACTTCATTATTTGCGAGTTCTCC
GGAAATACCTCAAGAATTTCTCCAGGGATTTCTTACAGGAATACCTCGAGGGTTTCCTCT
AGTGGTAGTTACAAGGTTTAACGAGAAGCCTTCA

Aal-m:

TTAAGCTAGTATTTACATTCTGGTTTCCGATTTAGTTTAATAAGAATCAATCCGCAAACAG
AGATAAGATTTTGTATCCTGGCCCGAGGTAGTGGCTCACAATGGCTCAACGCAGACCCA
AGCAACACAGAAAACATCTTTGGACCATCAGTATTTCCAAAGATGTTTTTTAAGTGAATT
ATAAGCGATTCAGAAAACATCTTGTGTTAATATCCAGCTCTAATTATGCTGATCGAAAACA
AACAACATCAAACATGTTATTGTCTATATCACGTATGAAATATGTTTATATAACTATCAGTG
AATCAAAATTCAACCATCGCGCAACAATAATGACAATGTCGCAACCTGTATTTGTTGCGA
CAAAACAACCCATGCGACATATGTTTGTCCCAACTTGAAAATAATGGGATTAACATCGTA
CACCACGAGTTTAGAGATTTTAAGCCTTGCATTGCGATTTTCGAACGGTTACTTCGAGTC
GGTAAACACTGCAACTCTGGTGAAGCTCTATCATCAAACAGTTTCGACTTTGGATTAGTA
ATAATTGATTATTCACGAGTTGAAAAGTACGCAACAAATTCAGCTTCCGTTTTGTCTGCA
ACAAAAAATTTCGAGATCATGTCATTATCTGTCACCAGTAGGGATAAAGACTATCCATCA
CATCTTCTTTGTTGCTTGTTTTGTGCCTCTTTTGTCTGACAATTCTCTTCACTGATAACTAT
TATATAACAGAAACATCAAGTTGCTAATTCAATCATATGTAAAAGCAAACGCGACAGAAC
AACAGAAAGAAGATAAAGATAAAAATAAACCAAAATGCAAACATTGTACCGTCTACCCC
CGATGGTTTGAACGACACCTCATGCAAACCAACGGGGTTCTTTTTTTAATTTGAACTTTT
AGTAACCCTGTGGACATCAACAAACGTACTTTTGGTAACCTTATTTGTTGTTTTGTTTTGA
TTCTGCGTTCCGTTTCACCGCGTTCCATGGGCAGAATGACGTTTGAACCATTTTTAGTTT
GAACGATGTGCAGATTGGCGGGGGTCAAATTAAAAAGTGTTCAGATTAGATGCTGTCAA

ACCAATGGGGGTAGACGGTATGCATTGATTTTAACCATTCCTAGGAGGCTCTGGTCGATG
GGGGAGAGTGAGTGGTTCAAGGGACCACGTCCCCCACGTCCGACGAACGTCCCCGCGC
TAATTAGGATCACCATACACAAATTTGGCCATCATCCTCAAGGCCAAAATGTACAGGCTG
CTGATCTACCAACTGGTCTGTCGAATACTATCCAGAAACACCGCGCAACTTGTATTCCGG
CCTATGTGAAAGAATCAGCATCCAACAGTCGGTGCTATTCCAACTCCAGCGTGAATTCCG
GCACTCAAACTTCACCTCTCTGCCGTGTGCAGCATAGTTGTCCCATGTTCCAAAACAGC
AAACTGAGAAAAACGCGTTTTAAGTTTCAGCATTGTTTCCATCTCTACGGACAAGTGTA
ATAAAAACACAAAATGAATCCACCATCATAGCGTTAGAATTGGTATTTATCATTGAAGTTT
GAACAAAAAGTTTCATAAACATACAATTTACTTTGATTACACGGTCCGACAAAAAATCA
ACTTTTGTTCTGCCCAGCTCTTAAATT

**Supplementary Data 2: Detailed statistical outputs**

(separate document)

**Supplementary Data 3: Production costs and initial parameters**

(separate document)

**Supplementary Data 4: Plasmid and genomic integration nucleotide sequences**

(separate document)

### Supplementary Data 2: Detailed statistical outputs

### Initial settings

#### R version

platform x86_64-w64-mingw32
arch x86_64
os mingw32
system x86_64, mingw32 major 4
minor 1.0
year 2021
svn rev 80317
language R
version.string R version 4.1.0 (2021-05-18) nickname Camp Pontanezen

#### Packages

```
set.seed(2501)

libs <- c(
  'plyr', 'dplyr', 'tidyr', 'stringr',
  'ggeffects', 'lme4', 'multcomp',
  'performance', 'reshape2', 'icenReg',
  'survminer', 'survtools', 'survival',
  'ggplot2', 'ggpubr', 'viridis', 'cowplot'
)
invisible(lapply(libs, library, character.only = T))
```

Versions:

plyr__1.8.6, dplyr__1.0.7, tidyr__1.1.3, stringr__1.4.0

ggeffects__1.0.2, lme4__1.1-26, multcomp__1.4-16

performance__0.7.0, reshape2__1.4.4, icenReg__2.0.15

survminer__0.4.9, survtools__0.1, survival__3.2-11

ggplot2__3.3.5, ggpubr__0.4.0, viridis__0.5.1

#### Plot common settings

```
# Panel background color
species.bg = c('Ae. aegypti' = 'white', 'Ae. albopictus' = 'grey')

# Aeg line colors
line.color.aeg = c('Bra (WT)' = "#24ff24", 'Aaeg-M' = "#004949",
                   'Aaeg-m' = "#009292", 'Aaeg-CS' = "#006ddb")

# Alb line colors
line.color.alb = c('BiA (WT)' = "#24ff24", 'Aal-M' = "#004949",
                   'Aal-m' = "#009292", 'Aal-CS' = "#006ddb")

# Set of colors
pal <- c('1' = "#000000", '2' = "#004949", '3' = "#009292",
         '4' = "#ff6db6", '5' = "#ffb6db", '6' = "#490092",
         '7' = "#006ddb", '8' = "#b66dff", '9' = "#6db6ff",
         '10' = "#b6dbff", '11' = "#920000", '12' = "#924900",
         '13' = "#db6d00", '14' = "#24ff24", '15' = "#ffff6d")
```

### Figure 3

#### Sex ratio

##### Aegypti

###### Data .

```
# Data measurements
sr.aeg.rep = read.csv('data/Sex-ratio_aeg.csv', header = T, sep = ';')
sr.aeg.rep$sr = sr.aeg.rep$Males / (sr.aeg.rep$Males + sr.aeg.rep$Females) * 100
sr.aeg.rep$Line = factor(sr.aeg.rep$Line,
                         levels = c('Bra (WT)', 'Aaeg-M', 'Aaeg-m' , 'Aaeg-CS'))
head(sr.aeg.rep)
```

```
  Replicate     Line Males Females       sr
1         2 Bra (WT)    97     100 49.23858
2         3 Bra (WT)    97      87 52.71739
3         4 Bra (WT)   126      95 57.01357
4         1   Aaeg-M   562     512 52.32775
5         2   Aaeg-M   912     837 52.14408
6         3   Aaeg-M   592     580 50.51195
```

```
# Build binomial dataset
males.aeg =
  rbind(
    data.frame(replicate = sr.aeg.rep[1,1],
               treatment = sr.aeg.rep[1,2],
               result = rep(1, sr.aeg.rep[1,3])),
    data.frame(replicate = sr.aeg.rep[1,1],
               treatment = sr.aeg.rep[1,2],
               result = rep(0, sr.aeg.rep[1,4])))

for(i in 2:nrow(sr.aeg.rep)){
  males.aeg =
    rbind(
      males.aeg,
      data.frame(replicate = sr.aeg.rep[i,1],
                 treatment = sr.aeg.rep[i,2],
                 result = rep(1, sr.aeg.rep[i,3])),
      data.frame(replicate = sr.aeg.rep[i,1],
                 treatment = sr.aeg.rep[i,2],
                 result = rep(0, sr.aeg.rep[i,4]))
    )

}
males.aeg$treatment = factor(x = males.aeg$treatment,
                             levels = c('Bra (WT)', 'Aaeg-M', 'Aaeg-m' , 'Aaeg-CS'))
males.aeg$Species = "Ae. aegypti"

summary.sr.aeg = males.aeg %>% group_by(treatment) %>%
  dplyr::summarise(value = n(),
            N = max(replicate)) %>% data.frame() %>%
```

```
  mutate(pos = c(1, 2, 3,4))

head(males.aeg)
```

```
  replicate treatment result     Species
1         2  Bra (WT)      1 Ae. aegypti
2         2  Bra (WT)      1 Ae. aegypti
3         2  Bra (WT)      1 Ae. aegypti
4         2  Bra (WT)      1 Ae. aegypti
5         2  Bra (WT)      1 Ae. aegypti
6         2  Bra (WT)      1 Ae. aegypti
```

###### Model .

```
# Generalized linear model with Bernouilli distribution
sr.aeg = glm(formula = 'result ~ treatment',
             family = binomial,
             data = males.aeg)
summary(sr.aeg)
```

```
Call:
glm(formula = "result ~ treatment", family = binomial, data = males.aeg)

Deviance Residuals:
   Min      1Q  Median      3Q     Max
-1.253  -1.228   1.103   1.128   1.148

Coefficients:
                  Estimate Std. Error z value Pr(>|z|)
(Intercept)       0.126414   0.081677   1.548    0.122
treatmentAaeg-M  -0.057801   0.087599  -0.660    0.509
treatmentAaeg-m  -0.009136   0.085106  -0.107    0.915
treatmentAaeg-CS  0.050668   0.087750   0.577    0.564

(Dispersion parameter for binomial family taken to be 1)

    Null deviance: 21476  on 15531  degrees of freedom
Residual deviance: 21470  on 15528  degrees of freedom
AIC: 21478

Number of Fisher Scoring iterations: 3
```

```
# Pairwise comparison
pairewise.aeg = glht(sr.aeg, mcp(treatment="Tukey"))

summary(pairewise.aeg)
```

```
     Simultaneous Tests for General Linear Hypotheses

Multiple Comparisons of Means: Tukey Contrasts
```

```
Fit: glm(formula = "result ~ treatment", family = binomial, data = males.aeg)

Linear Hypotheses:
                         Estimate Std. Error z value Pr(>|z|)
Aaeg-M - Bra (WT) == 0  -0.057801   0.087599  -0.660   0.9063
Aaeg-m - Bra (WT) == 0  -0.009136   0.085106  -0.107   0.9995
Aaeg-CS - Bra (WT) == 0  0.050668   0.087750   0.577   0.9347
Aaeg-m - Aaeg-M == 0     0.048666   0.039679   1.226   0.5924
Aaeg-CS - Aaeg-M == 0    0.108469   0.045071   2.407   0.0689 .
Aaeg-CS - Aaeg-m == 0    0.059804   0.040012   1.495   0.4216
---
Signif. codes:  0 '***' 0.001 '**' 0.01 '*' 0.05 '.' 0.1 ' ' 1
(Adjusted p values reported -- single-step method)
```

Replication

```
      Line N    n
1 Bra (WT) 4  602
2   Aaeg-M 3 3995
3   Aaeg-m 3 7017
4  Aaeg-CS 3 3918
```

Model fit quality

```
check_model(sr.aeg)
```

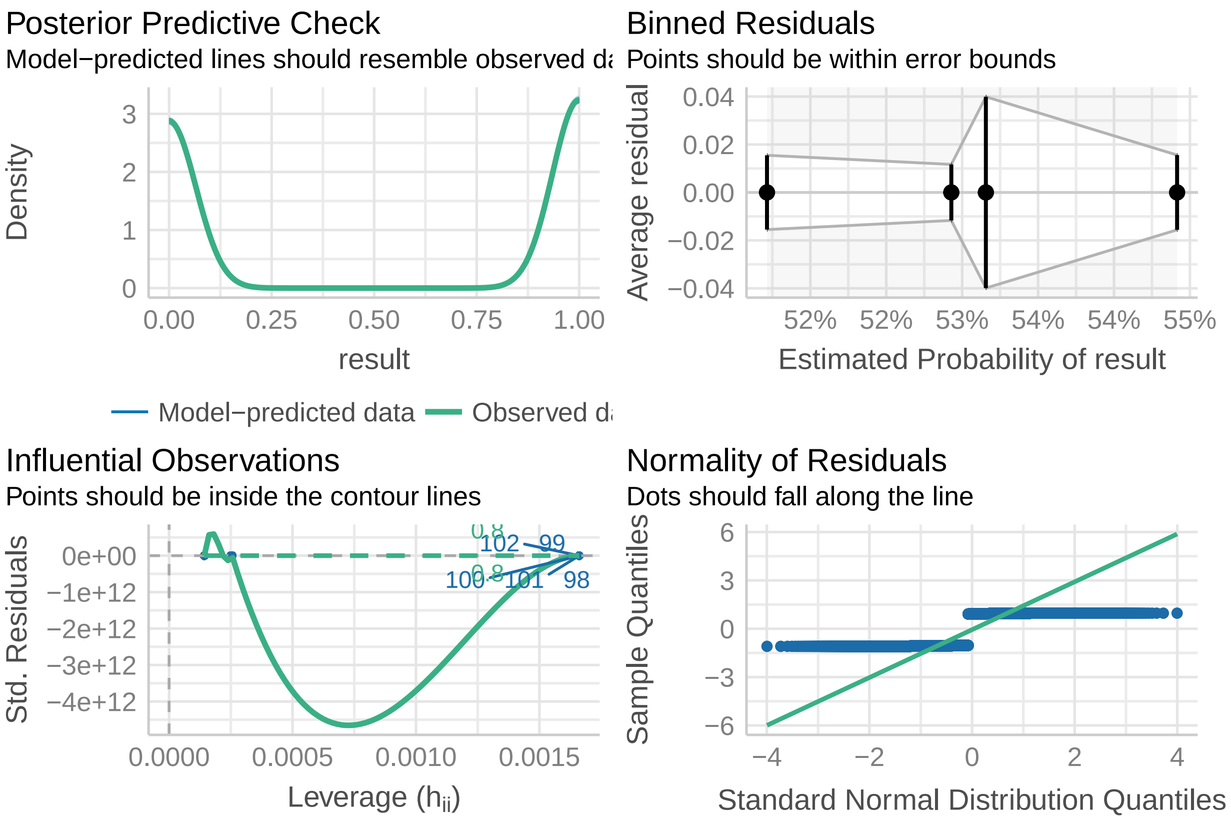

**Plot .**

```
# Extracion of the results
pred.sr.aeg = ggpredict(model = sr.aeg, terms = c("treatment")) %>% data.frame()
pred.sr.aeg$Species = "Ae. aegypti"

# plot
sr.plot.aeg =
  ggplot(data =  NULL) +
  geom_point(data = sr.aeg.rep,
             aes(x= jitter(as.numeric(factor(Line)), factor = .5),
                 y = sr , color = Line), alpha = .6, size = 2) +
  geom_point(data = pred.sr.aeg,
             aes(x = as.numeric(x),
                 y = predicted * 100), color = 'black',
             size = 1.5) +
  geom_errorbar(data = pred.sr.aeg,
                aes(x = as.numeric(x),
                    ymin = conf.low* 100,
                    ymax = conf.high* 100),
                color = 'black',
                width = 0.05) +

  geom_point(data = males.aeg,
             aes(x = jitter(as.numeric(factor(treatment)), factor = 2),
                 y = jitter((result * 95), factor = .25), color = treatment),
             alpha = .25,
             size = .01
  ) +
  annotate(geom = 'text', x= summary.sr.aeg$pos, y=-8,
           label = 'N = 3',
           size = 3) +
  annotate(geom = 'text', x= summary.sr.aeg$pos, y=-17,
           label = paste0('n = ', summary.sr.aeg$value),
           size = 3) +
  annotate(geom = 'line', x= 1:4, y=118) +
  annotate(geom = 'text', x= 2.5, y=123,
           label = 'n.s.',
           size = 3) +
  annotate(geom = 'line', x= 1:3, y=111) +
  annotate(geom = 'text', x= 2, y=116,
           label = 'n.s.',
           size = 3) +
  annotate(geom = 'line', x= 1:2, y=103) +
  annotate(geom = 'text', x= 1.5, y=109,
           label = 'n.s.',
           size = 3) +
  scale_x_continuous(breaks = 1:4,
                     labels = c("Bra (WT)",
                                "Aaeg-M",
                                "Aaeg-m",
                                "Aaeg-CS"),
                     limits = c(.5,4.5), guide = guide_axis(angle = 30)) +
```

```
  scale_y_continuous(sec.axis = sec_axis(~., breaks = c(0,95),
                                         labels = c('Females', 'Males'),
                                         name = 'Sex (binomial)'),
                     limits = c(-20,126),
                     breaks = c(0,25,50,75,100)) +
  scale_color_manual(values = line.color.aeg, labels = names(line.color.aeg)) +
  labs(x = 'Line',
       y = 'Percentage of males (%)') +
  theme_classic() +
  theme(panel.background =
          element_rect(fill = species.bg[names(species.bg)=="Ae. aegypti"]),
        legend.position = 'none',
        axis.title.y.right = element_text(color = 'black'),
        axis.text.y.right = element_text( angle = -45,
                                          hjust = 0, vjust = 0),
        axis.ticks.y.right = element_line(color = 'black'),
        axis.line.y.right = element_line(color = 'black'))

sr.plot.aeg
```

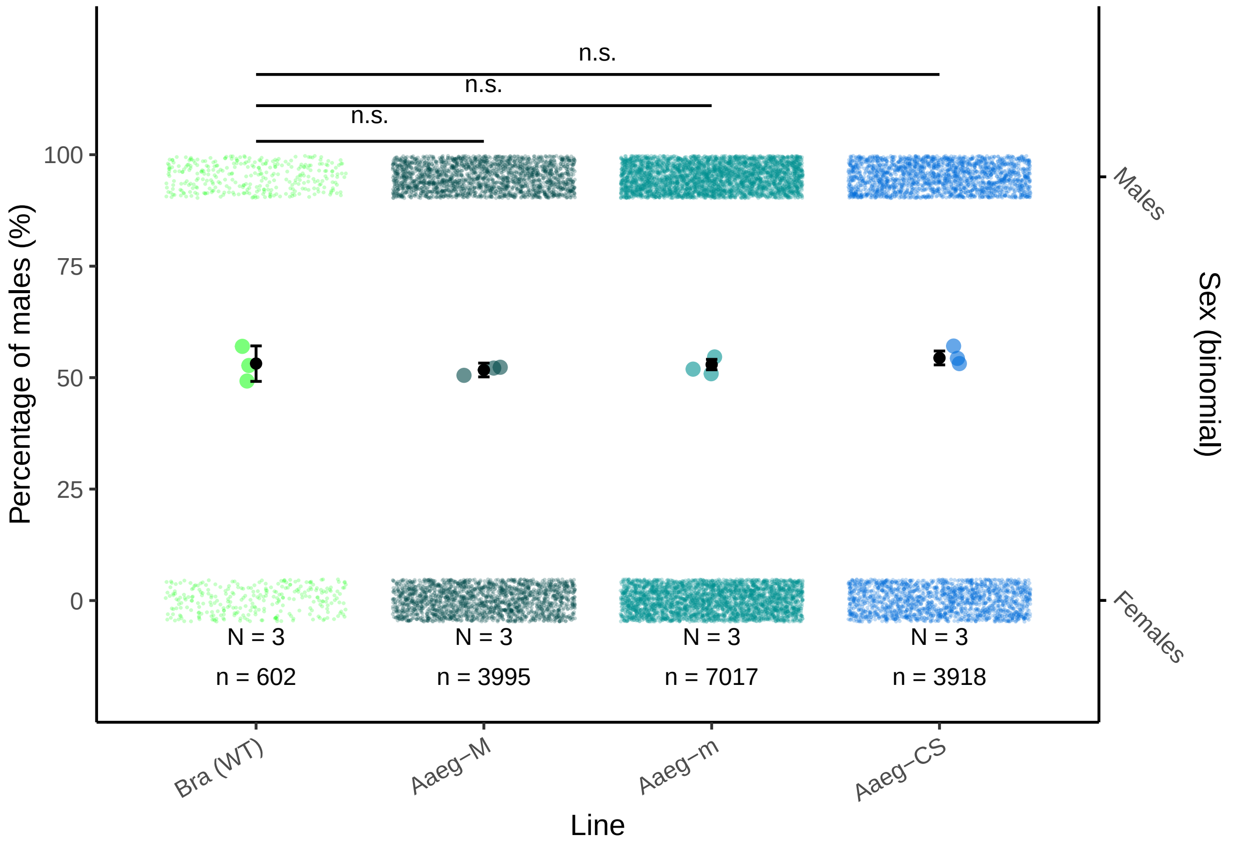

**Albopictus**

**Data** .

```r
# Data measurements
sr.alb.rep = read.table("data/Sex-ratio_albo.csv", header=T, sep=";")
sr.alb.rep$sr = sr.alb.rep$Males /
  (sr.alb.rep$Males + sr.alb.rep$Females) * 100
sr.alb.rep$Line = factor(sr.alb.rep$Line,
                         levels = c('BiA (WT)', 'Aal-M', 'Aal-m', 'Aal-CS'))
head(sr.alb.rep)
```

```
  Replicate   Line Males Females       sr
1         A  Aal-m   167     166 50.15015
2         B  Aal-m   245     246 49.89817
3         C  Aal-m   293     274 51.67549
4         D Aal-CS   136     119 53.33333
5         E Aal-CS   533     493 51.94932
6         F Aal-CS   510     471 51.98777
```

```r
# Build binomial dataset
males.albo =
  rbind(
    data.frame(replicate = sr.alb.rep[1,1],
               treatment = sr.alb.rep[1,2],
               result = rep(1, sr.alb.rep[1,3])),
    data.frame(replicate = sr.alb.rep[1,1],
               treatment = sr.alb.rep[1,2],
               result = rep(0, sr.alb.rep[1,4])))

for(i in 2:nrow(sr.alb.rep)){
  males.albo =
    rbind(
      males.albo,
      data.frame(replicate = sr.alb.rep[i,1],
                 treatment = sr.alb.rep[i,2],
                 result = rep(1, sr.alb.rep[i,3])),
      data.frame(replicate = sr.alb.rep[i,1],
                 treatment = sr.alb.rep[i,2],
                 result = rep(0, sr.alb.rep[i,4]))
    )

}
males.albo$treatment= factor(males.albo$treatment,
                             levels = c('BiA (WT)', 'Aal-M', 'Aal-m', 'Aal-CS'))
males.albo$Species = "Ae. albopictus"

summary.sr.alb = males.albo %>% group_by(treatment) %>%
  dplyr::summarise(value = n()) %>% data.frame() %>%
  mutate(pos = c(1, 2, 3,4))

head(males.albo)
```

```
  replicate treatment result        Species
1         A     Aal-m      1 Ae. albopictus
2         A     Aal-m      1 Ae. albopictus
3         A     Aal-m      1 Ae. albopictus
```

```
4          A     Aal-m      1 Ae. albopictus
5          A     Aal-m      1 Ae. albopictus
6          A     Aal-m      1 Ae. albopictus
```

**Model** .

```
# Generalized linear model with Bernouilli distribution
sr.albo = glm(formula = 'result ~ treatment',
              family = binomial,
              data = males.albo)
summary(sr.albo)
```

```

Call:
glm(formula = "result ~ treatment", family = binomial, data = males.albo)

Deviance Residuals:
   Min      1Q  Median      3Q     Max
-1.228  -1.189   1.128   1.166   1.166

Coefficients:
                Estimate Std. Error z value Pr(>|z|)
(Intercept)      0.11842    0.10388   1.140    0.254
treatmentAal-M  -0.09122    0.11350  -0.804    0.422
treatmentAal-m  -0.09110    0.11690  -0.779    0.436
treatmentAal-CS -0.03349    0.11208  -0.299    0.765

(Dispersion parameter for binomial family taken to be 1)

    Null deviance: 8226.0  on 5936  degrees of freedom
Residual deviance: 8224.4  on 5933  degrees of freedom
AIC: 8232.4

Number of Fisher Scoring iterations: 3
```

```
# Pairwise comparison
pairewise.albo = glht(sr.albo, mcp(treatment="Tukey"))

summary(pairewise.albo)
```

```

     Simultaneous Tests for General Linear Hypotheses

Multiple Comparisons of Means: Tukey Contrasts

Fit: glm(formula = "result ~ treatment", family = binomial, data = males.albo)

Linear Hypotheses:
                         Estimate Std. Error z value Pr(>|z|)
Aal-M - BiA (WT) == 0  -0.0912194  0.1135026  -0.804    0.847
Aal-m - BiA (WT) == 0  -0.0910976  0.1169041  -0.779    0.858
Aal-CS - BiA (WT) == 0 -0.0334861  0.1120801  -0.299    0.990
Aal-m - Aal-M == 0      0.0001218  0.0704883   0.002    1.000
```

```
Aal-CS - Aal-M == 0      0.0577333  0.0621609   0.929    0.781
Aal-CS - Aal-m == 0      0.0576115  0.0681741   0.845    0.826
(Adjusted p values reported -- single-step method)
```

Replication

```
# A tibble: 4 x 3
  Line          N     n
  <fct>     <int> <int>
1 BiA (WT)      3   372
2 Aal-M         3  1912
3 Aal-m         3  1391
4 Aal-CS        3  2262
```

Model fit quality

```
check_model(sr.albo)
```

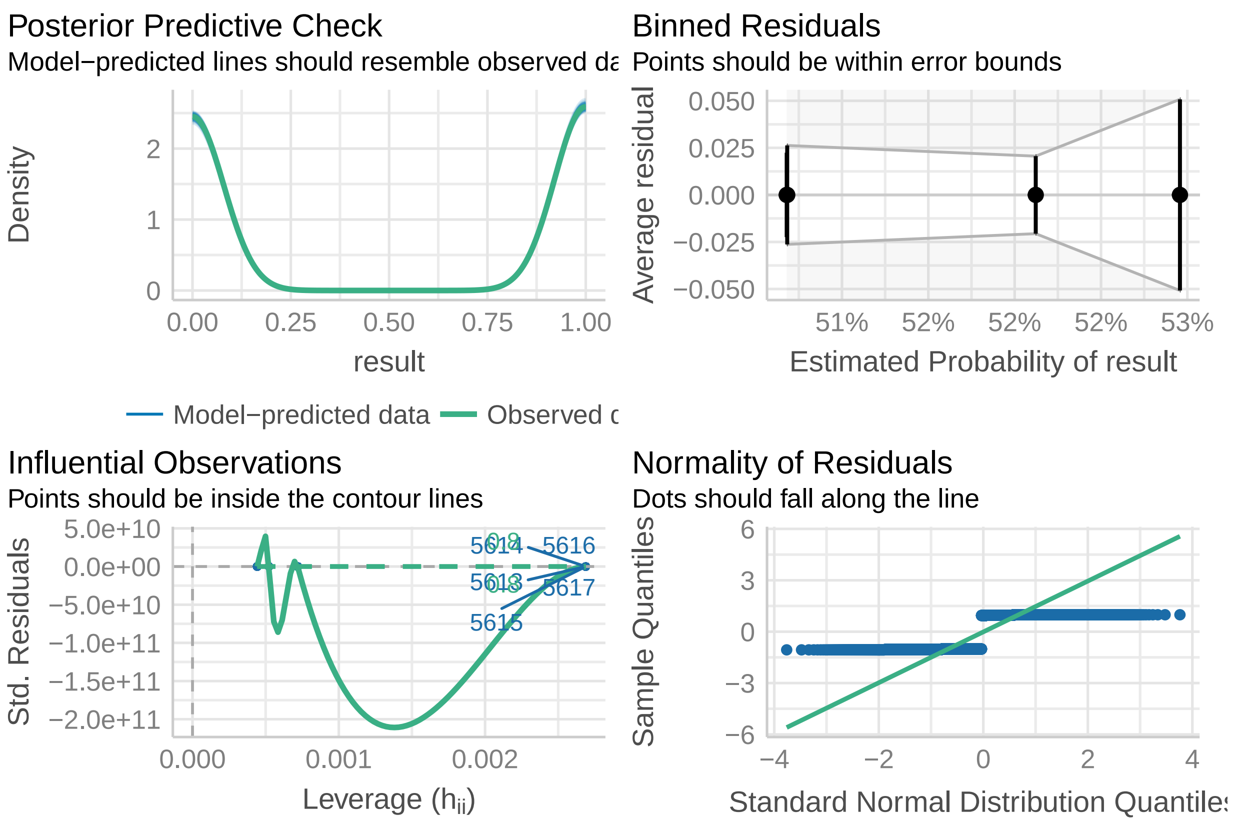

**Plot** .

```
# Extracion of the results
pred.sr.albo = ggpredict(model = sr.albo, terms = c("treatment")) %>%
  data.frame()
pred.sr.albo$Species = "Ae. albopictus"
```

```r
# plot
sr.plot.albo =
  ggplot(data =  NULL) +
  geom_point(data = sr.alb.rep,
             aes(x= jitter(as.numeric(factor(Line)), factor = .5),
                 y = sr ,
                 color = Line),
             alpha = .6,
             size = 2) +
  geom_point(data = pred.sr.albo,
             aes(x = as.numeric(x),
                 y = predicted * 100),
             size = 1.5, color = 'black') +
  geom_errorbar(data = pred.sr.albo,
                aes(x = as.numeric(x),
                    ymin = conf.low* 100,
                    ymax = conf.high* 100),
                width = 0.05,
                color = 'black') +
  geom_point(data = males.albo,
             aes(x = jitter(as.numeric(factor(treatment)), factor = 2),
                 y = jitter((result * 95), factor = .25),
                 color = treatment),
             alpha = .25,
             size = .01
  ) +
  annotate(geom = 'text', x= summary.sr.alb$pos, y=-8,
           label = 'N = 3',
           size = 3) +
  annotate(geom = 'text', x= summary.sr.alb$pos, y=-17,
           label = paste0('n = ', summary.sr.alb$value),
           size = 3) +
  annotate(geom = 'line', x= 1:4, y=118) +
  annotate(geom = 'text', x= 2.5, y=123,
           label = 'n.s.',
           size = 3) +
  annotate(geom = 'line', x= 1:3, y=111) +
  annotate(geom = 'text', x= 2, y=116,
           label = 'n.s.',
           size = 3) +
  annotate(geom = 'line', x= 1:2, y=103) +
  annotate(geom = 'text', x= 1.5, y=109,
           label = 'n.s.',
           size = 3) +
  scale_x_continuous(breaks = 1:4,
                     labels = c("BiA (WT)",
                                "Aal-M",
                                "Aal-m",
                                "Aal-CS"),
                     limits = c(.5,4.5),
                     guide = guide_axis(angle = 30)) +
  scale_y_continuous(sec.axis = sec_axis(~., breaks = c(0,95),
```

```
                                          labels = c('Females', 'Males'),
                                          name = 'Sex (binomial)'),
                      limits = c(-20,126),
                      breaks = c(0,25,50,75,100)) +
  scale_color_manual(values = line.color.alb, labels = names(line.color.alb)) +
  labs(x = 'Line',
       y = 'Percentage of males (%)') +
  theme_classic() +
  theme(panel.background =
          element_rect(fill = species.bg[names(species.bg)=="Ae. albopictus"]),
        legend.position = 'none',
        axis.title.y.right = element_text(color = 'black'),
        axis.text.y.right = element_text(angle = -45,
                                          hjust = 0, vjust = 0),
        axis.ticks.y.right = element_line(color = 'black'),
        axis.line.y.right = element_line(color = 'black'))

sr.plot.albo
```

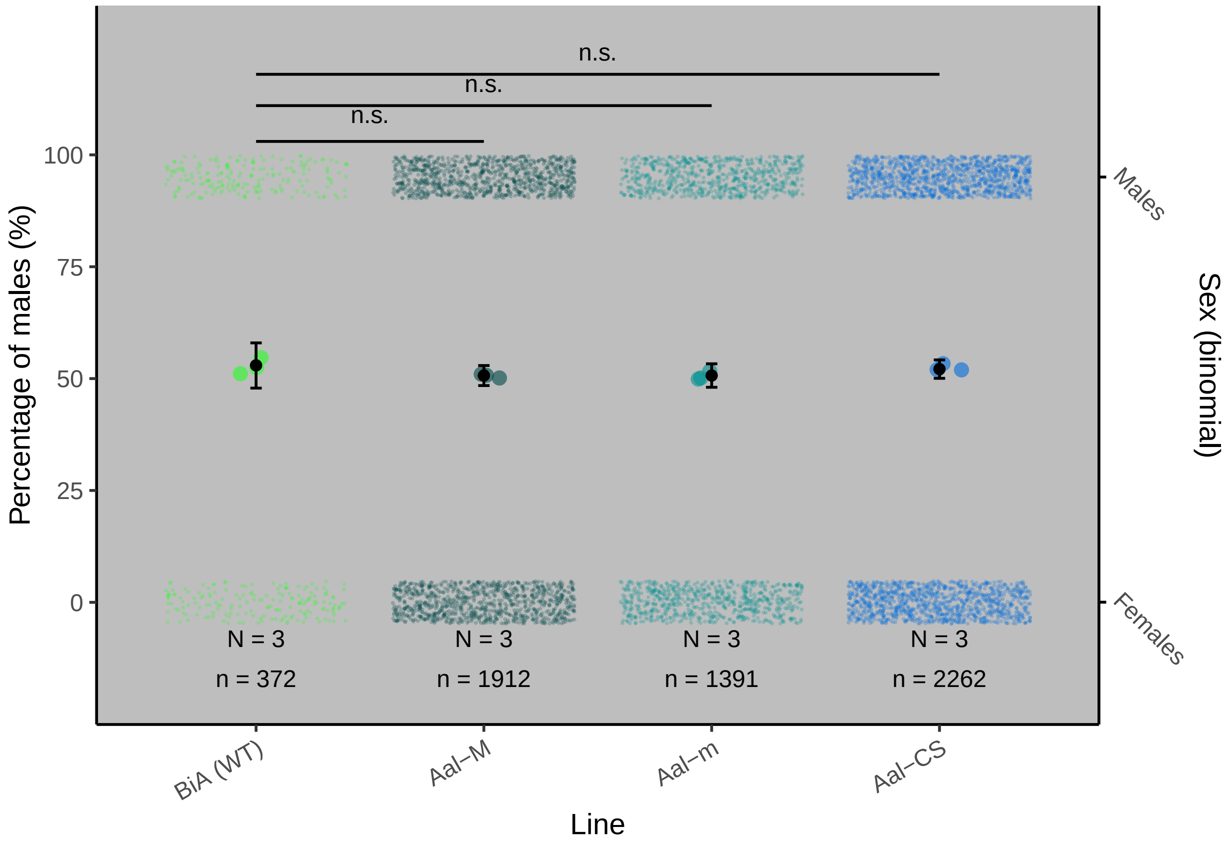

#### Hatching rate

##### Aegypti

###### Data .

```r
# Data measurements
hr.aeg = read.table("data/H-rate_aeg.csv", header = T, sep = ";")
hr.aeg$hr = hr.aeg$Larvae/hr.aeg$Eggs * 100
hr.aeg$Line = factor(hr.aeg$Line, levels = c("Bra (WT)",
                                             "Aaeg-M",
                                             "Aaeg-m",
                                             "Aaeg-CS"))
head(hr.aeg)
```

```
  Replicate     Line Eggs Larvae Unhatched       hr
1         1 Bra (WT)  220    140        80 63.63636
2         2 Bra (WT)  235    141        94 60.00000
3         3 Bra (WT)  272    166       106 61.02941
4         1   Aaeg-M  435    259       176 59.54023
5         2   Aaeg-M  297    157       140 52.86195
6         3   Aaeg-M  291    172       119 59.10653
```

```r
# Build binomial dataset
hatch.aegypti =
  rbind(
    data.frame(replicate = hr.aeg[1,1],
               treatment = hr.aeg[1,2],
               result = rep(0, hr.aeg[1,5])),
    data.frame(replicate = hr.aeg[1,1],
               treatment = hr.aeg[1,2],
               result = rep(1, hr.aeg[1,4])))

for(i in 2:nrow(hr.aeg)){
  hatch.aegypti =
    rbind(
      hatch.aegypti,
      data.frame(replicate = hr.aeg[i,1],
                 treatment = hr.aeg[i,2],
                 result = rep(0, hr.aeg[i,5])),
      data.frame(replicate = hr.aeg[i,1],
                 treatment = hr.aeg[i,2],
                 result = rep(1, hr.aeg[i,4]))
    )

}
hatch.aegypti$treatment=factor(hatch.aegypti$treatment,
                               levels = c("Bra (WT)", "Aaeg-M", "Aaeg-m", "Aaeg-CS"))
hatch.aegypti$Species = 'Ae. aegypti'

summary.hatch.aeg = hatch.aegypti %>% group_by(treatment) %>%
  dplyr::summarise(value = n()) %>% data.frame() %>%
  mutate(pos = c(1, 2, 3, 4))

head(hatch.aegypti)
```

```
  replicate treatment result     Species
1         1  Bra (WT)      0 Ae. aegypti
2         1  Bra (WT)      0 Ae. aegypti
```

```
3         1  Bra (WT)      0 Ae. aegypti
4         1  Bra (WT)      0 Ae. aegypti
5         1  Bra (WT)      0 Ae. aegypti
6         1  Bra (WT)      0 Ae. aegypti
```

##### Model .

```r
# Generalized linear model with Bernouilli distribution
hatch.aegypti_stat = glm(formula = 'result ~ treatment',
                         family = binomial,
                         data = hatch.aegypti)
summary(hatch.aegypti_stat)
```

```

Call:
glm(formula = "result ~ treatment", family = binomial, data = hatch.aegypti)

Deviance Residuals:
    Min       1Q   Median       3Q      Max
-1.8245  -1.3078   0.6479   0.6934   1.0524

Coefficients:
                 Estimate Std. Error z value Pr(>|z|)
(Intercept)       0.46777    0.07621   6.138 8.38e-10 ***
treatmentAaeg-M  -0.16639    0.09904  -1.680   0.0929 .
treatmentAaeg-m   0.98677    0.10649   9.266  < 2e-16 ***
treatmentAaeg-CS  0.83507    0.10166   8.214  < 2e-16 ***
---
Signif. codes:  0 '***' 0.001 '**' 0.01 '*' 0.05 '.' 0.1 ' ' 1

(Dispersion parameter for binomial family taken to be 1)

    Null deviance: 5089.2  on 4242  degrees of freedom
Residual deviance: 4871.9  on 4239  degrees of freedom
AIC: 4879.9

Number of Fisher Scoring iterations: 4
```

```r
# Pairwise comparison
pairewise.h.aeg = glht(hatch.aegypti_stat,
                       mcp(treatment="Tukey"))

h.aeg<-summary(pairewise.h.aeg)
h.aeg
```

```

     Simultaneous Tests for General Linear Hypotheses

Multiple Comparisons of Means: Tukey Contrasts

Fit: glm(formula = "result ~ treatment", family = binomial, data = hatch.aegypti)

Linear Hypotheses:
```

```
                        Estimate Std. Error z value Pr(>|z|)
Aaeg-M - Bra (WT) == 0  -0.16639    0.09904  -1.680    0.334
Aaeg-m - Bra (WT) == 0   0.98677    0.10649   9.266   <1e-04 ***
Aaeg-CS - Bra (WT) == 0  0.83507    0.10166   8.214   <1e-04 ***
Aaeg-m - Aaeg-M == 0     1.15316    0.09763  11.812   <1e-04 ***
Aaeg-CS - Aaeg-M == 0    1.00145    0.09233  10.846   <1e-04 ***
Aaeg-CS - Aaeg-m == 0   -0.15170    0.10029  -1.513    0.429
---
Signif. codes:  0 '***' 0.001 '**' 0.01 '*' 0.05 '.' 0.1 ' ' 1
(Adjusted p values reported -- single-step method)
```

Replication

```
# A tibble: 4 x 3
  Treatment      N     n
  <fct>      <int> <int>
1 Bra (WT)       3   727
2 Aaeg-M         3  1023
3 Aaeg-m         3  1178
4 Aaeg-CS        3  1315
```

Model fit quality

```
check_model(hatch.aegypti_stat)
```

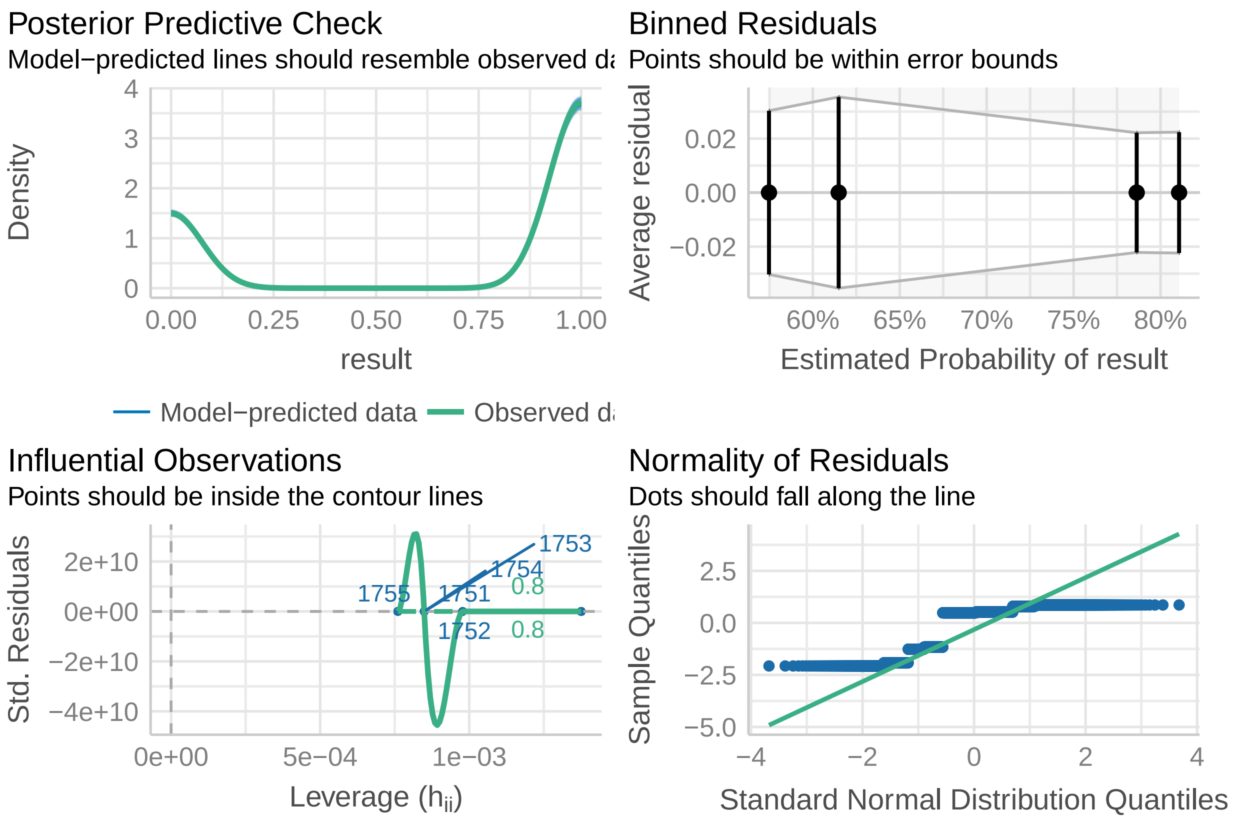

**Plot .**

```
# Extraction of the results
pred.aeg = data.frame(ggpredict(model = hatch.aegypti_stat,
                                terms = c("treatment")))
pred.aeg$x = factor(pred.aeg$x, levels = c("Bra (WT)",
                                           "Aaeg-M",
                                           "Aaeg-m",
                                           "Aaeg-CS"))
pred.aeg$Species = 'Ae. aegypti'

# plot
hr.plot.aeg =
  ggplot(data =  NULL) +
  geom_point(data = hr.aeg,
             aes(x = jitter(as.numeric(Line), factor = 1),
                 y = hr, color = Line),
             size = 2, alpha = .6) +
  geom_point(data = pred.aeg,
             aes(x = as.numeric(x),
                 y = predicted * 100),
             size = 1.5 , color = 'black') +
  geom_errorbar(data = pred.aeg,
                aes(x = as.numeric(x),
                    ymin = conf.low* 100,
                    ymax = conf.high* 100),
                width = 0.05,
                color = 'black') +
  geom_point(data = hatch.aegypti,
             aes(x = jitter(as.numeric(factor(treatment)), factor = 2),
                 y = jitter((result * 95), factor = .25),
                 color = treatment),
             size = .05
  ) +
  annotate(geom = 'text', x= summary.hatch.aeg$pos, y=-8,
           label = 'N = 3',
           size = 3) +
  annotate(geom = 'text', x= summary.hatch.aeg$pos, y=-17,
           label = paste0('n = ', summary.hatch.aeg$value),
           size = 3) +
  annotate(geom = 'line', x= 1:4, y=120.5) +
  annotate(geom = 'text', x= 2.5, y=123,
           label = '***',
           size = 3) +
  annotate(geom = 'line', x= 1:3, y=112) +
  annotate(geom = 'text', x= 2, y=116,
           label = '***',
           size = 3) +
  annotate(geom = 'line', x= 1:2, y=103) +
  annotate(geom = 'text', x= 1.5, y=109,
           label = 'n.s.',
           size = 3) +
  scale_x_continuous(breaks = 1:4,
```

```
                     labels = c("Bra (WT)",
                                "Aaeg-M",
                                "Aaeg-m",
                                "Aaeg-CS"),
                     limits = c(.5,4.5), guide = guide_axis(angle = 30)) +
  scale_y_continuous(sec.axis = sec_axis(~., breaks = c(0,95),
                                         labels = c('Unhatched', 'Hatched'),
                                         name = 'Hatching status (binomial)'),
                     limits = c(-20,126),
                     breaks = c(0,25,50,75,100)) +
  scale_color_manual(values = line.color.aeg, labels = names(line.color.aeg)) +
  labs(x = 'Line',
       y = 'Hatched eggs (%)') +
  theme_classic() +
  theme(legend.position = 'none',
        axis.title.y.right = element_text(color = 'black'),
        axis.text.y.right = element_text(angle = -45,
                                         hjust = 0, vjust = 0),
        axis.ticks.y.right = element_line(color = 'black'),
        axis.line.y.right = element_line(color = 'black'))

hr.plot.aeg
```

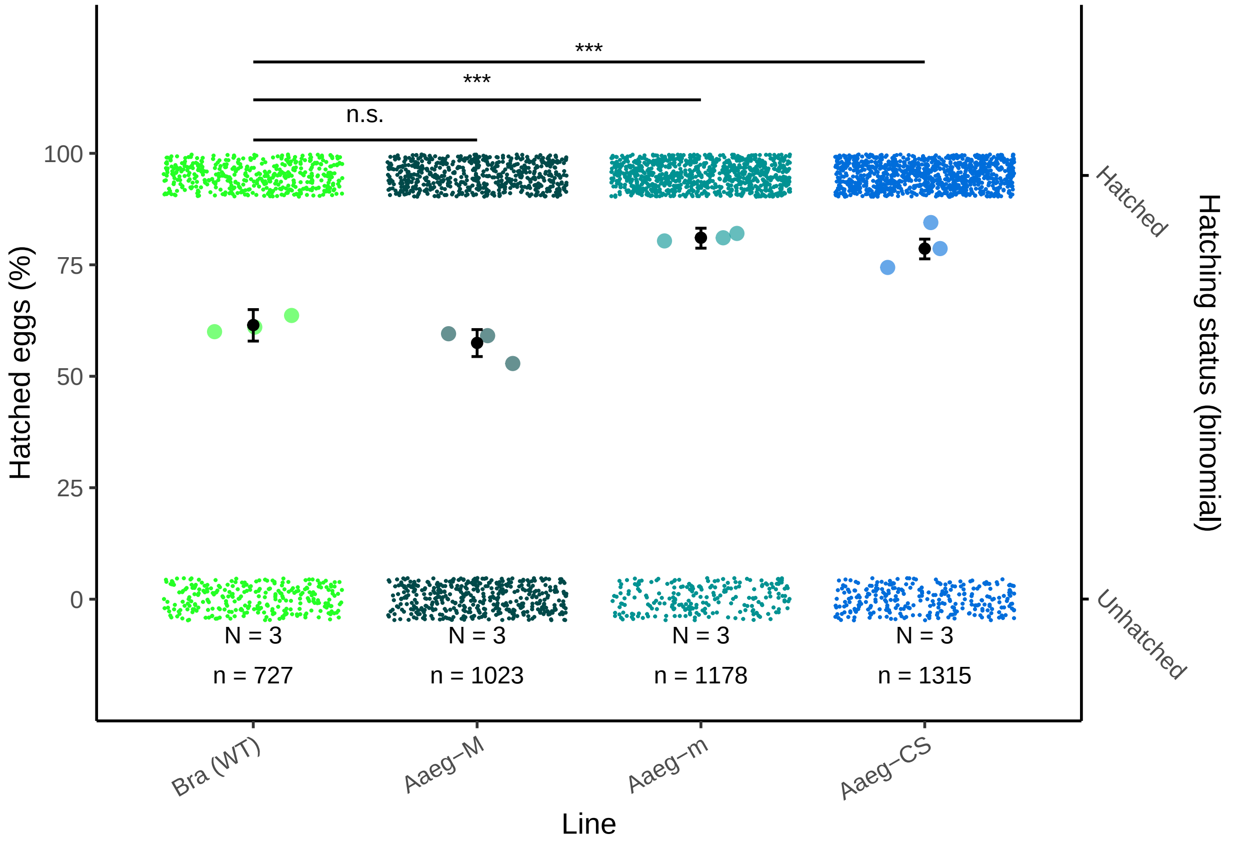

##### Albopictus

###### Data .

```
# Data measurements
hr.alb = read.table("data/H-rate_albo.csv", header = T, sep = ";")
hr.alb$hr = hr.alb$Larvae/hr.alb$Eggs * 100
hr.alb$Line = factor(hr.alb$Line, levels = c("BiA (WT)",
                                             "Aal-M",
                                             "Aal-m",
                                             "Aal-CS"))
head(hr.alb)
```

```
  Replicate     Line Eggs Larvae Unhatched       hr
1         1 BiA (WT)  965    555       410 57.51295
2         2 BiA (WT)  750    461       289 61.46667
3         3 BiA (WT) 1025    539       486 52.58537
4         4 BiA (WT)  730    326       404 44.65753
5         5 BiA (WT)  756    347       409 45.89947
6         1    Aal-M  927    632       295 68.17691
```

```
# Build binomial dataset
hatch.albo =
  rbind(
    data.frame(replicate = hr.alb[1,1],
               treatment = hr.alb[1,2],
               result = rep(0, hr.alb[1,5])),
    data.frame(replicate = hr.alb[1,1],
               treatment = hr.alb[1,2],
               result = rep(1, hr.alb[1,4])))

for(i in 2:nrow(hr.alb)){
  hatch.albo =
    rbind(
      hatch.albo,
      data.frame(replicate = hr.alb[i,1],
                 treatment = hr.alb[i,2],
                 result = rep(0, hr.alb[i,5])),
      data.frame(replicate = hr.alb[i,1],
                 treatment = hr.alb[i,2],
                 result = rep(1, hr.alb[i,4]))
    )

}
hatch.albo$treatment=factor(hatch.albo$treatment,
                            levels = c("BiA (WT)", "Aal-M", "Aal-m", "Aal-CS"))
hatch.albo$Species = 'Ae. albopictus'

summary.hatch.albo = hatch.albo %>%
  group_by(treatment) %>%
  dplyr::summarise(value = n()) %>%
  data.frame() %>%
  mutate(pos = c(1, 2, 3, 4))
```

```r
head(hatch.albo)
```

```
  replicate treatment result        Species
1         1  BiA (WT)      0 Ae. albopictus
2         1  BiA (WT)      0 Ae. albopictus
3         1  BiA (WT)      0 Ae. albopictus
4         1  BiA (WT)      0 Ae. albopictus
5         1  BiA (WT)      0 Ae. albopictus
6         1  BiA (WT)      0 Ae. albopictus
```

**Model** .

```r
# Generalized linear model with Bernouilli distribution
hatch.albo_stat = glm(formula = 'result ~ treatment',
                      family = binomial,
                      data = hatch.albo)
summary(hatch.albo_stat)
```

```
Call:
glm(formula = "result ~ treatment", family = binomial, data = hatch.albo)

Deviance Residuals:
   Min      1Q  Median      3Q     Max
-1.289  -1.267   1.070   1.091   1.144

Coefficients:
                Estimate Std. Error z value Pr(>|z|)
(Intercept)      0.10896    0.03081   3.536 0.000406 ***
treatmentAal-M   0.14952    0.04263   3.507 0.000453 ***
treatmentAal-m  -0.02901    0.05747  -0.505 0.613746
treatmentAal-CS  0.09815    0.03801   2.582 0.009816 **
---
Signif. codes:  0 '***' 0.001 '**' 0.01 '*' 0.05 '.' 0.1 ' ' 1

(Dispersion parameter for binomial family taken to be 1)

    Null deviance: 25862  on 18771  degrees of freedom
Residual deviance: 25844  on 18768  degrees of freedom
AIC: 25852

Number of Fisher Scoring iterations: 3
```

```r
# Pairwise comparison
pairewise.h.alb = glht(hatch.albo_stat,
                       mcp(treatment="Tukey"))

h.albo <- summary(pairewise.h.alb)
h.albo
```

```
     Simultaneous Tests for General Linear Hypotheses
```

```
Multiple Comparisons of Means: Tukey Contrasts

Fit: glm(formula = "result ~ treatment", family = binomial, data = hatch.albo)

Linear Hypotheses:
                       Estimate Std. Error z value Pr(>|z|)
Aal-M - BiA (WT) == 0   0.14952    0.04263   3.507  0.00240 **
Aal-m - BiA (WT) == 0  -0.02901    0.05747  -0.505  0.95667
Aal-CS - BiA (WT) == 0  0.09815    0.03801   2.582  0.04601 *
Aal-m - Aal-M == 0     -0.17853    0.05676  -3.145  0.00856 **
Aal-CS - Aal-M == 0    -0.05137    0.03693  -1.391  0.49668
Aal-CS - Aal-m == 0     0.12716    0.05338   2.382  0.07723 .
---
Signif. codes:  0 '***' 0.001 '**' 0.01 '*' 0.05 '.' 0.1 ' ' 1
(Adjusted p values reported -- single-step method)
```

Replication

```
# A tibble: 4 x 3
  Treatment     N     n
  <fct>     <int> <int>
1 BiA (WT)      5  4226
2 Aal-M         5  4684
3 Aal-m         5  1702
4 Aal-CS        5  8160
```

Model fit quality

```
check_model(hatch.albo_stat)
```

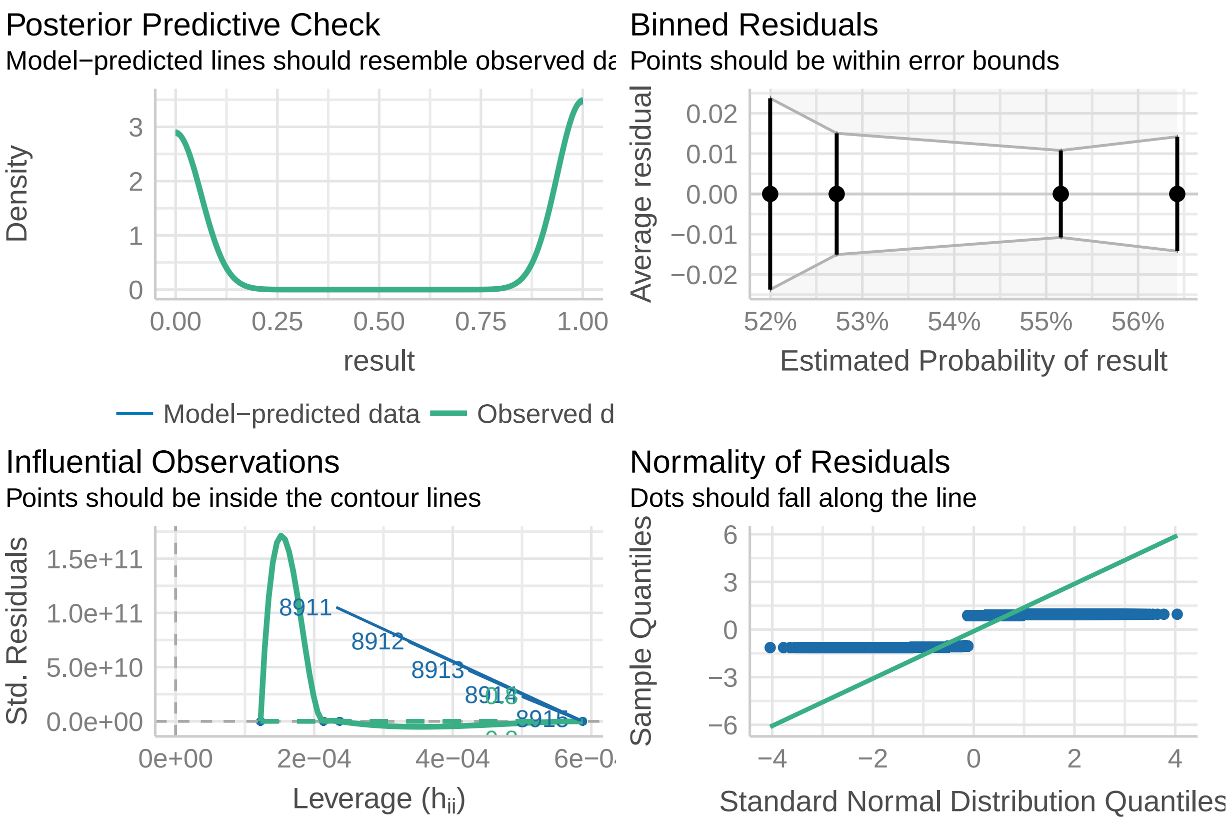

**Plot** .

```
# Extracion of the results
pred.alb = data.frame(ggpredict(model = hatch.albo_stat,
                                terms = c("treatment")))
pred.alb$x = as.character(pred.alb$x)
pred.alb$x = factor(pred.alb$x, levels = c("BiA (WT)",
                                           "Aal-M",
                                           "Aal-m",
                                           "Aal-CS"))
pred.alb$Species = 'Ae. albopictus'
pred.alb$group = as.factor(2)

# plot
hr.plot.alb =
  ggplot(data =  NULL) +
  geom_point(data = hr.alb,
             aes(x = jitter(as.numeric(Line), factor = 1),
                 y = hr, color = Line),
             size = 2, alpha = .6) +
  geom_point(data = pred.alb,
             aes(x = as.numeric(x),
                 y = predicted * 100),
             size = 1.5 , color = 'black') +
  geom_errorbar(data = pred.alb,
```

```
              aes(x = as.numeric(x),
                  ymin = conf.low* 100,
                  ymax = conf.high* 100),
              width = 0.05,
              color = 'black') +
geom_point(data = hatch.albo,
           aes(x = jitter(as.numeric(factor(treatment)), factor = 2),
               y = jitter((result * 95), factor = .25),
               color = treatment),
           size = .05
) +
annotate(geom = 'text', x= summary.hatch.albo$pos, y=-8,
         label = 'N = 5',
         size = 3) +
annotate(geom = 'text', x= summary.hatch.albo$pos, y=-17,
         label = paste0('n = ', summary.hatch.albo$value),
         size = 3) +
annotate(geom = 'line', x= 1:4, y=120.5) +
annotate(geom = 'text', x= 2.5, y=123,
         label = '*',
         size = 3) +
annotate(geom = 'line', x= 1:3, y=113) +
annotate(geom = 'text', x= 2, y=116,
         label = 'n.s.',
         size = 3) +
annotate(geom = 'line', x= 1:2, y=103) +
annotate(geom = 'text', x= 1.5, y=109,
         label = '**',
         size = 3) +
scale_x_continuous(breaks = 1:4,
                   labels = c("BiA (WT)",
                              "Aal-M",
                              "Aal-m",
                              "Aal-CS"),
                   limits = c(.5,4.5), guide = guide_axis(angle = 30)) +
scale_y_continuous(sec.axis = sec_axis(~., breaks = c(0,95),
                                       labels = c('Unhatched', 'Hatched'),
                                       name = 'Hatching status (binomial)'),
                   limits = c(-20,126),
                   breaks = c(0,25,50,75,100)) +
scale_color_manual(values = line.color.alb,
                   labels = names(line.color.alb)) +
labs(x = 'Line',
     y = 'Hatched eggs (%)') +
theme_classic() +
theme(panel.background =
        element_rect(fill =species.bg[names(species.bg)=="Ae. albopictus"]),
      legend.position = 'none',
      axis.title.y.right = element_text(color = 'black'),
      axis.text.y.right = element_text(angle = -45,
                                       hjust = 0, vjust = 0),
      axis.ticks.y.right = element_line(color = 'black'),
      axis.line.y.right = element_line(color = 'black'))
```

```
hr.plot.alb
```

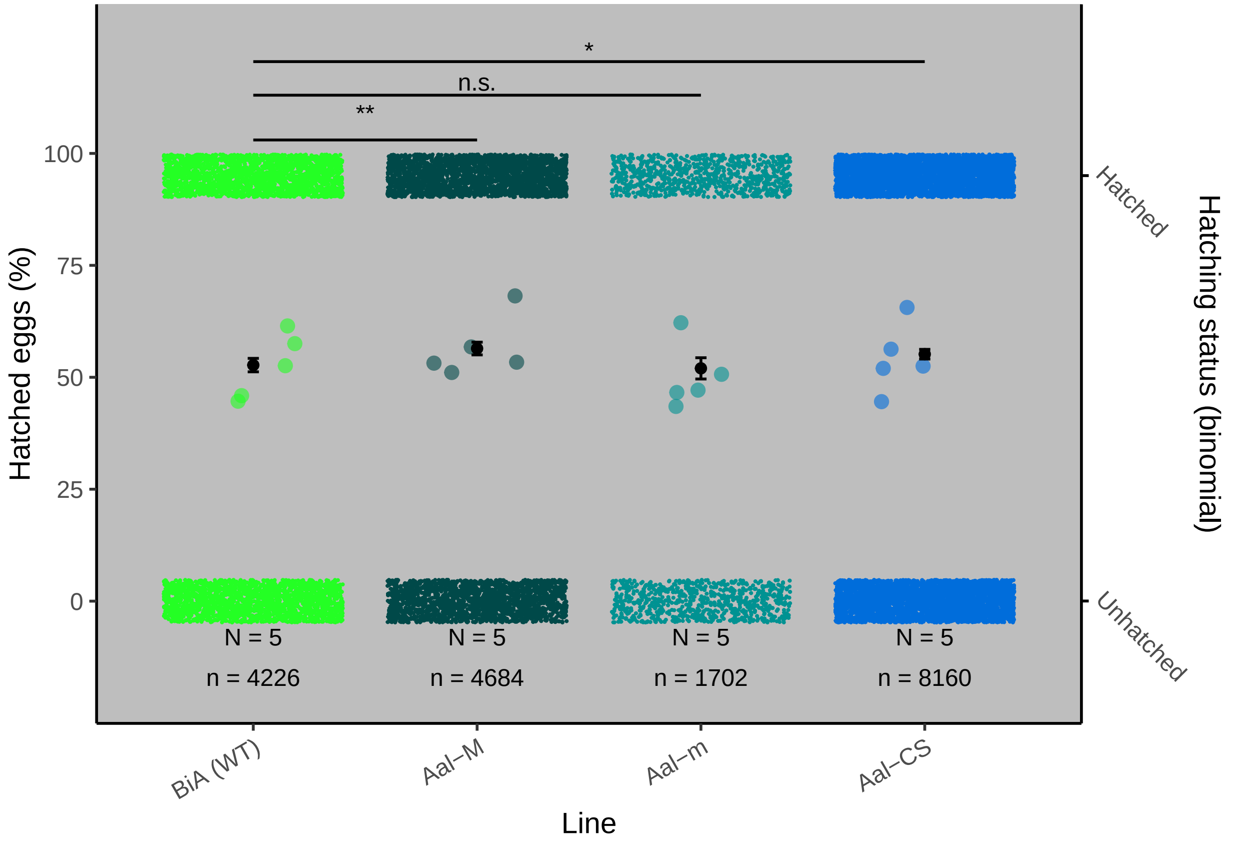

#### Fecundity

##### Aegypti

###### Data .

```
fecund_aeg = read.table("data/Fecundity_aeg.csv", header=TRUE, sep=";")

fecund_aeg = fecund_aeg %>% filter(Eggs != "NA")

fecund_aeg$Line = factor(fecund_aeg$Line,
                         levels = c('Bra (WT)', 'Aaeg-M', 'Aaeg-m', 'Aaeg-CS'))

fecund_aeg$Species = "Ae. aegypti"

head(fecund_aeg)
```

```
      Line Well Eggs     Species
1 Bra (WT)   A1   33 Ae. aegypti
2 Bra (WT)   A2    0 Ae. aegypti
3 Bra (WT)   A3    0 Ae. aegypti
```

```
4 Bra (WT)   A4   53 Ae. aegypti
5 Bra (WT)   A5   96 Ae. aegypti
6 Bra (WT)   A6    7 Ae. aegypti
```

```
fecund.summary.aeg = fecund_aeg %>%
  group_by(Line) %>%
  dplyr::summarise(value = n()) %>%
  data.frame() %>%
  mutate(pos = c(1, 2, 3, 4))
```

##### Model .

```
fecund_aeg.glm = glm(formula = 'Eggs ~ Line',
                     data = fecund_aeg, family = 'poisson')
mod = MASS::glm.nb(formula = 'Eggs ~ Line',
                     data = fecund_aeg)

mod1=pscl::hurdle(Eggs ~ Line,
             data = fecund_aeg,
             dist = 'negbin')
mod2 = pscl::zeroinfl(Eggs ~ Line,
             data = fecund_aeg,
             dist = 'poisson')
summary(mod2)
```

```
Call:
pscl::zeroinfl(formula = Eggs ~ Line, data = fecund_aeg, dist = "poisson")

Pearson residuals:
   Min     1Q Median     3Q    Max
-2.181 -1.491 -0.460  1.429  4.219

Count model coefficients (poisson with log link):
            Estimate Std. Error z value Pr(>|z|)
(Intercept)  3.62041    0.03968  91.232  < 2e-16 ***
LineAaeg-M  -0.19724    0.05819  -3.389 0.000701 ***
LineAaeg-m  -0.06843    0.05711  -1.198 0.230803
LineAaeg-CS  0.59582    0.04893  12.176  < 2e-16 ***

Zero-inflation model coefficients (binomial with logit link):
            Estimate Std. Error z value Pr(>|z|)
(Intercept)  -0.8873     0.4491  -1.976   0.0482 *
LineAaeg-M   -0.2113     0.6511  -0.325   0.7455
LineAaeg-m   -0.8473     0.7706  -1.100   0.2715
LineAaeg-CS  -0.6168     0.7122  -0.866   0.3865
---
Signif. codes:  0 '***' 0.001 '**' 0.01 '*' 0.05 '.' 0.1 ' ' 1

Number of iterations in BFGS optimization: 1
Log-likelihood: -1056 on 8 Df
```

```
summary(mod)
```

```

Call:
MASS::glm.nb(formula = "Eggs ~ Line", data = fecund_aeg, init.theta = 0.4279000025,
    link = log)

Deviance Residuals:
    Min       1Q   Median       3Q      Max
-2.0419  -1.3305  -0.1797   0.3794   1.0646

Coefficients:
            Estimate Std. Error z value Pr(>|z|)
(Intercept)   3.2756     0.3146  10.413   <2e-16 ***
LineAaeg-M   -0.1401     0.4451  -0.315    0.753
LineAaeg-m    0.1139     0.4664   0.244    0.807
LineAaeg-CS   0.7400     0.4539   1.630    0.103
---
Signif. codes:  0 '***' 0.001 '**' 0.01 '*' 0.05 '.' 0.1 ' ' 1

(Dispersion parameter for Negative Binomial(0.4279) family taken to be 1)

    Null deviance: 110.57  on 89  degrees of freedom
Residual deviance: 105.84  on 86  degrees of freedom
AIC: 776.53

Number of Fisher Scoring iterations: 1

              Theta:  0.4279
          Std. Err.:  0.0651

 2 x log-likelihood:  -766.5330
```

```
check_distribution(mod)
```

```

# Distribution of Model Family

Predicted Distribution of Residuals

 Distribution Probability
       normal         34%
      tweedie         25%
         beta         19%

Predicted Distribution of Response

               Distribution Probability
 neg. binomial (zero-infl.)        100%
```

```
check_homogeneity(mod)
```

```

OK: There is not clear evidence for different variances across groups (Bartlett Test, p = 0.433).
```

```r
check_zeroinflation(mod)
```

```
# Check for zero-inflation

   Observed zeros: 20
  Predicted zeros: 14
            Ratio: 0.70
```

```r
check_singularity(mod)
```

```
[1] FALSE
```

```r
performance::compare_performance(mod1, fecund_aeg.glm, mod2, mod)
```

```
# Comparison of Model Performance Indices

Name           |    Model |      AIC | AIC weights |      BIC | BIC weights |   RMSE |  Sigma | Score_l
------------------------------------------------------------------------------------------------------
mod1           |   hurdle |  768.797 |       0.980 |  791.295 |       0.244 | 31.040 | 32.719 |    -5.1
fecund_aeg.glm |      glm | 3463.728 |     < 0.001 | 3473.727 |     < 0.001 | 31.040 |  6.007 |   -19.1
mod2           | zeroinfl | 2127.533 |     < 0.001 | 2147.531 |     < 0.001 | 31.040 | 32.519 |   -11.7
mod            |   negbin |  776.533 |       0.020 |  789.032 |       0.756 | 31.040 |  1.109 |    -4.7
```

```r
check_model(mod)
```

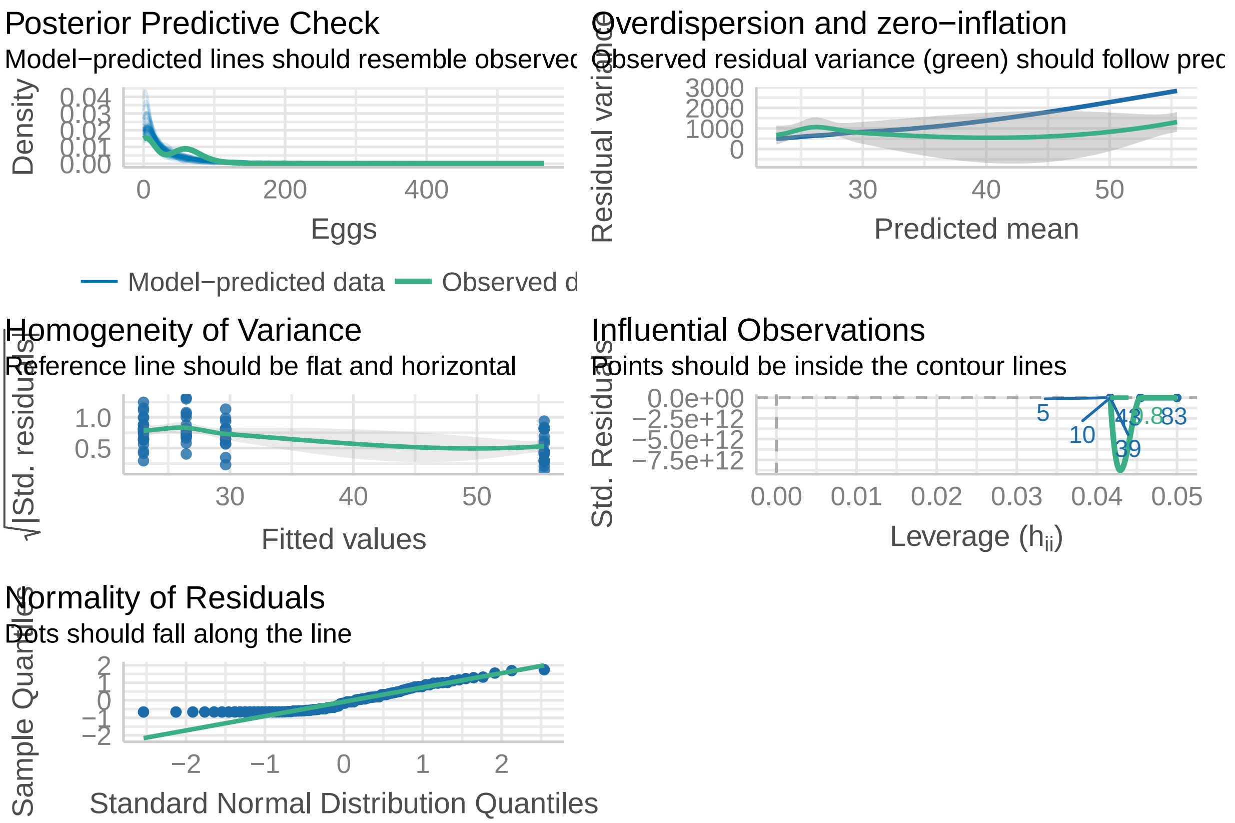

```r
pscl::hurdle.control(mod1)
```

```
$method

Call:
pscl::hurdle(formula = Eggs ~ Line, data = fecund_aeg, dist = "negbin")

Count model coefficients (truncated negbin with log link):
(Intercept)   LineAaeg-M   LineAaeg-m  LineAaeg-CS
    3.59467     -0.20305     -0.07031      0.60761
Theta = 1.0192

Zero hurdle model coefficients (binomial with logit link):
(Intercept)   LineAaeg-M   LineAaeg-m  LineAaeg-CS
     0.8873       0.2113       0.8473       0.6168

$maxit
[1] 10000

$trace
[1] FALSE

$separate
[1] TRUE

$start
NULL

$fnscale
[1] -1

$hessian
[1] TRUE

$reltol
[1] 1.646361e-10
```

```r
summary(mod1)
```

```

Call:
pscl::hurdle(formula = Eggs ~ Line, data = fecund_aeg, dist = "negbin")

Pearson residuals:
    Min      1Q  Median      3Q     Max
-0.8767 -0.7530 -0.2053  0.5867  1.9791

Count model coefficients (truncated negbin with log link):
            Estimate Std. Error z value Pr(>|z|)
(Intercept)  3.59467    0.24406  14.728   <2e-16 ***
LineAaeg-M  -0.20305    0.34025  -0.597   0.5507
LineAaeg-m  -0.07031    0.34473  -0.204   0.8384
```

```
LineAaeg-CS  0.60761    0.33873   1.794   0.0728 .
Log(theta)   0.01902    0.20161   0.094   0.9248
Zero hurdle model coefficients (binomial with logit link):
            Estimate Std. Error z value Pr(>|z|)
(Intercept)   0.8873     0.4491   1.976   0.0482 *
LineAaeg-M    0.2113     0.6511   0.325   0.7455
LineAaeg-m    0.8473     0.7706   1.100   0.2715
LineAaeg-CS   0.6168     0.7122   0.866   0.3865
---
Signif. codes:  0 '***' 0.001 '**' 0.01 '*' 0.05 '.' 0.1 ' ' 1

Theta: count = 1.0192
Number of iterations in BFGS optimization: 11
Log-likelihood: -375.4 on 9 Df
```

```r
fec_aeg_sum = summary(fecund_aeg.glm)
fec_aeg_sum
```

```
Call:
glm(formula = "Eggs ~ Line", family = "poisson", data = fecund_aeg)

Deviance Residuals:
    Min       1Q   Median       3Q      Max
-10.531   -6.659   -1.429    3.857   10.410

Coefficients:
            Estimate Std. Error z value Pr(>|z|)
(Intercept)  3.27557    0.03968  82.542   <2e-16 ***
LineAaeg-M  -0.14008    0.05819  -2.407   0.0161 *
LineAaeg-m   0.11389    0.05711   1.994   0.0461 *
LineAaeg-CS  0.73999    0.04893  15.122   <2e-16 ***
---
Signif. codes:  0 '***' 0.001 '**' 0.01 '*' 0.05 '.' 0.1 ' ' 1

(Dispersion parameter for poisson family taken to be 1)

    Null deviance: 3503.5  on 89  degrees of freedom
Residual deviance: 3103.4  on 86  degrees of freedom
AIC: 3463.7

Number of Fisher Scoring iterations: 6
```

```r
emmeans::emmeans(mod1, specs = 'Line', mode = 'response')
```

```
 Line     emmean   SE df lower.CL upper.CL
 Bra (WT)   26.5 3.49 81     19.5     33.4
 Aaeg-M     23.0 2.73 81     17.6     28.4
 Aaeg-m     29.6 2.82 81     24.0     35.3
 Aaeg-CS    55.5 5.62 81     44.3     66.6

Confidence level used: 0.95
```

```r
# Pairwise comparison
pairewise.f.aeg = glht(fecund_aeg.glm,
                       mcp(Line = "Tukey"))

f.aeg <- summary(pairewise.f.aeg)
f.aeg
```

```
     Simultaneous Tests for General Linear Hypotheses

Multiple Comparisons of Means: Tukey Contrasts

Fit: glm(formula = "Eggs ~ Line", family = "poisson", data = fecund_aeg)

Linear Hypotheses:
                        Estimate Std. Error z value Pr(>|z|)
Aaeg-M - Bra (WT) == 0  -0.14008    0.05819  -2.407   0.0747 .
Aaeg-m - Bra (WT) == 0   0.11389    0.05711   1.994   0.1882
Aaeg-CS - Bra (WT) == 0  0.73999    0.04893  15.122   <0.001 ***
Aaeg-m - Aaeg-M == 0     0.25397    0.05914   4.294   <0.001 ***
Aaeg-CS - Aaeg-M == 0    0.88007    0.05130  17.157   <0.001 ***
Aaeg-CS - Aaeg-m == 0    0.62610    0.05006  12.507   <0.001 ***
---
Signif. codes:  0 '***' 0.001 '**' 0.01 '*' 0.05 '.' 0.1 ' ' 1
(Adjusted p values reported -- single-step method)
```

```r
pred.fecund.aeg =
  emmeans::emmeans(mod1, specs = 'Line', mode = 'response') %>%
  data.frame() %>%
  dplyr::select(
    Line, mean.pro = emmean,
    sd.pro = SE,
    conf.inf = lower.CL,
    conf.sup = upper.CL
  )

head(pred.fecund.aeg)
```

```
      Line mean.pro   sd.pro conf.inf conf.sup
1 Bra (WT) 26.45834 3.486389 19.52151 33.39516
2   Aaeg-M 23.00000 2.731288 17.56559 28.43440
3   Aaeg-m 29.64999 2.818363 24.04234 35.25765
4  Aaeg-CS 55.45453 5.623214 44.26610 66.64297
```

Replication

```
      Line  N
1 Bra (WT) 24
2   Aaeg-M 24
3   Aaeg-m 20
4  Aaeg-CS 22
```

Model fit quality

```
check_model(fecund_aeg.glm)
```

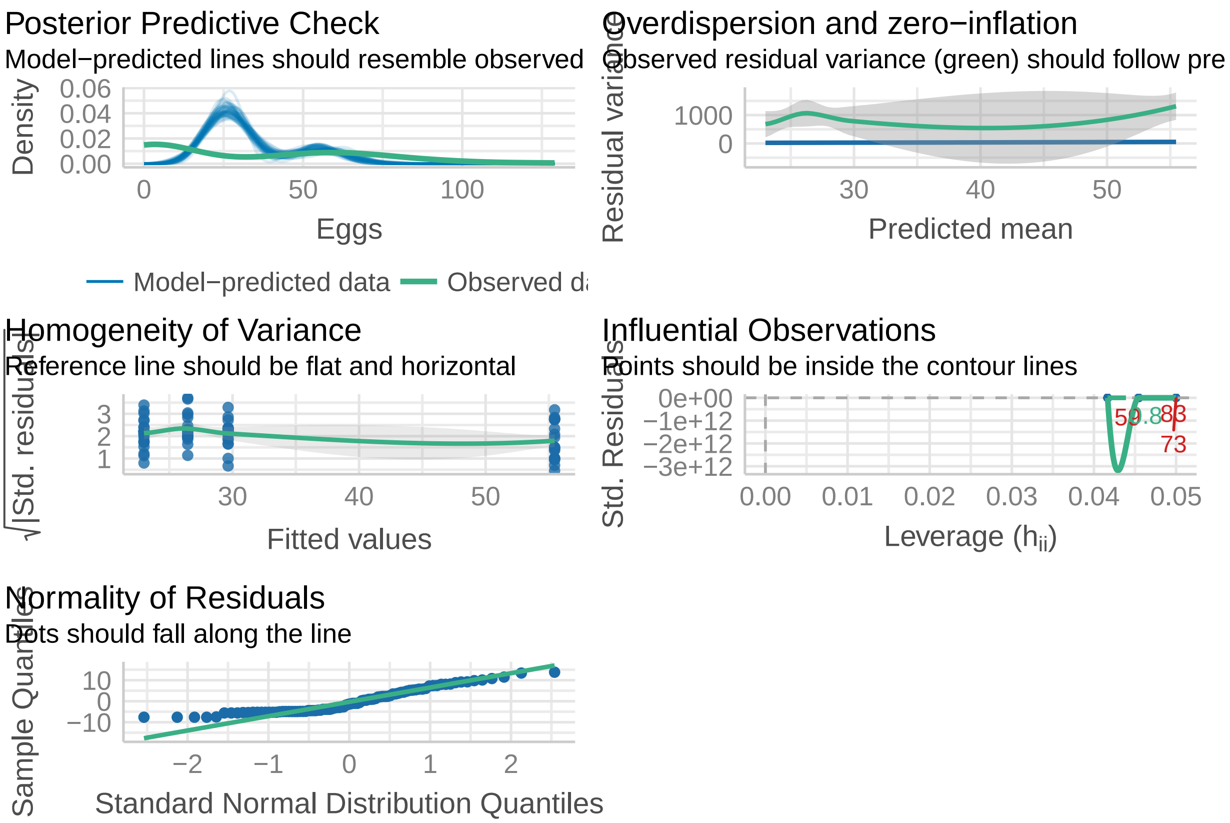

**Plot** .

```
fecund.plot.aeg  = ggplot(data=NULL) +
  geom_point(data = fecund_aeg,
             aes(x = jitter(as.numeric(Line)),
                 y = Eggs, color = Line),
             size = 2, show.legend = FALSE, alpha = 0.6) +
  geom_point(data = pred.fecund.aeg,
             aes(x = as.numeric(Line),
                 y = mean.pro),
             size = 1.5, show.legend = FALSE,
             color = 'black') +
  geom_errorbar(data = pred.fecund.aeg,
                aes(x = as.numeric(Line),
                    ymin = conf.inf,
                    ymax = conf.sup), width = 0.05,
                color = 'black') +
  annotate(geom = 'text', x= fecund.summary.aeg$pos, y = -8,
           label = paste0('N = ', fecund.summary.aeg$value),
           size = 3) +
  annotate(geom = 'line', x= 1:2, y=180) +
```

```
  annotate(geom = 'text', x= 1.5, y=192,
           label = 'n.s.',
           size = 3) +
  annotate(geom = 'line', x= 1:3, y=198) +
  annotate(geom = 'text', x= 2, y=210,
           label = 'n.s.',
           size = 3) +
  annotate(geom = 'line', x= 1:4, y=216) +
  annotate(geom = 'text', x= 2.5, y=228,
           label = '.',
           size = 3) +
  labs(x = 'Line', y = 'Number of eggs per female',
       color = "Line") +
  scale_y_continuous(breaks = seq(0,175,25),limits = c(-10,230)) +
  scale_color_manual(values = line.color.aeg,
                     labels = names(line.color.aeg)) +
  scale_x_continuous(guide = guide_axis(angle = 30), breaks = 1:4,
                     labels = c("Bra (WT)", "Aaeg-M", "Aaeg-m", "Aaeg-CS")) +
  theme_classic() +
  theme(legend.position = 'none')

fecund.plot.aeg
```

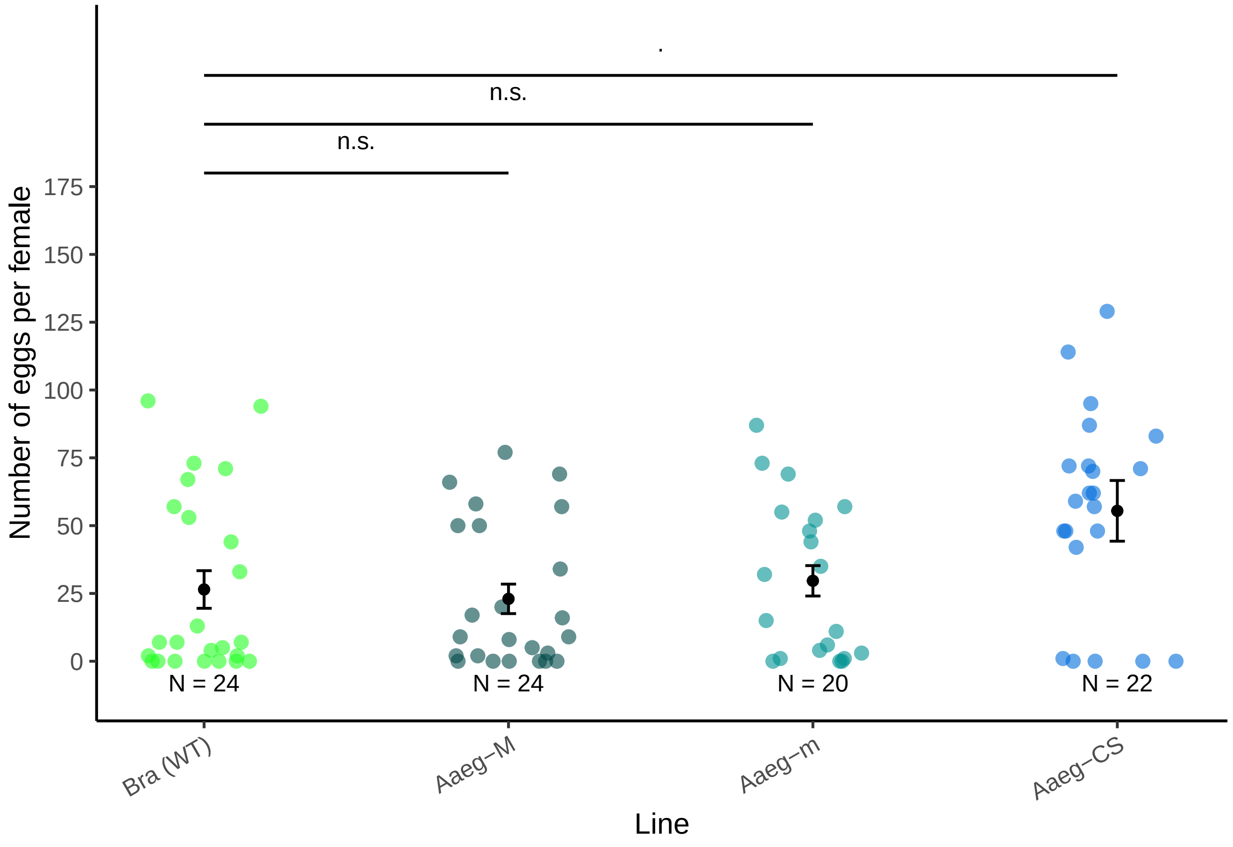

##### Albopictus

###### Data .

```
fecund_albo = read.table("data/Fecundity_albo.csv", header=TRUE, sep=";")

fecund_albo = fecund_albo %>% filter(Eggs != "NA")

fecund_albo$Line = factor(fecund_albo$Line,
                          levels = c('BiA (WT)', 'Aal-M', 'Aal-m', 'Aal-CS'))

fecund_albo$Species = "Ae. albopictus"

head(fecund_albo)
```

```
      Line Well Eggs        Species
1 BiA (WT)   A1    3 Ae. albopictus
2 BiA (WT)   A2  100 Ae. albopictus
3 BiA (WT)   A3    1 Ae. albopictus
4 BiA (WT)   A4    0 Ae. albopictus
5 BiA (WT)   A5   97 Ae. albopictus
6 BiA (WT)   A6   17 Ae. albopictus
```

```
fecund.summary.alb = fecund_albo %>%
  group_by(Line) %>%
  dplyr::summarise(value = n()) %>%
  data.frame() %>%
  mutate(pos = c(1, 2, 3, 4))
```

###### Model .

```
fecund_albo.glm = glm(formula = 'Eggs ~ Line',
                      data = fecund_albo,
                      family = 'poisson')

fec_albo_sum = summary(fecund_albo.glm)
fec_albo_sum
```

```
Call:
glm(formula = "Eggs ~ Line", family = "poisson", data = fecund_albo)

Deviance Residuals:
    Min       1Q   Median       3Q      Max
-11.752   -8.501   -2.258    5.312   12.365

Coefficients:
            Estimate Std. Error z value Pr(>|z|)
(Intercept)  3.64556    0.03369 108.206   <2e-16 ***
LineAal-M    0.58931    0.04356  13.529   <2e-16 ***
LineAal-m    0.04123    0.04668   0.883    0.377
LineAal-CS  -0.05826    0.04892  -1.191    0.234
---
```

```
Signif. codes:  0 '***' 0.001 '**' 0.01 '*' 0.05 '.' 0.1 ' ' 1

(Dispersion parameter for poisson family taken to be 1)

    Null deviance: 5146.1  on 87  degrees of freedom
Residual deviance: 4856.8  on 84  degrees of freedom
AIC: 5201.8

Number of Fisher Scoring iterations: 6
```

```
mod1=pscl::hurdle(Eggs ~ Line,
             data = fecund_albo,
             dist = 'poisson',
             zero.dist = "binomial",
             link="logit")
summary(mod1)
```

```
Call:
pscl::hurdle(formula = Eggs ~ Line, data = fecund_albo, dist = "poisson",
    zero.dist = "binomial", link = "logit")

Pearson residuals:
    Min      1Q  Median      3Q     Max
-2.2251 -1.4223 -0.4688  1.4213  4.3899

Count model coefficients (truncated poisson with log link):
            Estimate Std. Error z value Pr(>|z|)
(Intercept)  3.89069    0.03369 115.482   <2e-16 ***
LineAal-M    0.51603    0.04356  11.847   <2e-16 ***
LineAal-m    0.40921    0.04668   8.767   <2e-16 ***
LineAal-CS   0.07961    0.04892   1.627    0.104
Zero hurdle model coefficients (binomial with logit link):
            Estimate Std. Error z value Pr(>|z|)
(Intercept)   1.2809     0.5055   2.534   0.0113 *
LineAal-M     0.3930     0.8071   0.487   0.6263
LineAal-m    -1.1139     0.6507  -1.712   0.0869 .
LineAal-CS   -0.5188     0.6820  -0.761   0.4468
---
Signif. codes:  0 '***' 0.001 '**' 0.01 '*' 0.05 '.' 0.1 ' ' 1

Number of iterations in BFGS optimization: 10
Log-likelihood: -1314 on 8 Df
```

```
emmeans::emmeans(mod1, specs = 'Line', mode = 'response')
```

```
 Line     emmean   SE df lower.CL upper.CL
 BiA (WT)   38.3 4.40 80     29.5     47.1
 Aal-M      69.1 7.12 80     54.9     83.2
 Aal-m      39.9 7.61 80     24.8     55.1
 Aal-CS     36.1 5.42 80     25.4     46.9

Confidence level used: 0.95
```

```r
check_distribution(mod1)
```

```
# Distribution of Model Family

Predicted Distribution of Residuals

 Distribution Probability
      tweedie         53%
       normal         44%
          chi          3%

Predicted Distribution of Response

               Distribution Probability
 neg. binomial (zero-infl.)        100%
```

```r
check_homogeneity(mod1)
```

```
OK: There is not clear evidence for different variances across groups (Bartlett Test, p = 0.859).
```

```r
check_zeroinflation(mod1)
```

```
# Check for zero-inflation

   Observed zeros: 26
  Predicted zeros: 0
            Ratio: 0.00
```

```r
check_singularity(mod1)
```

```
[1] FALSE
```

```r
performance::compare_performance(mod1, fecund_albo.glm)
```

```
# Comparison of Model Performance Indices

Name            |  Model |      AIC | AIC weights |      BIC | BIC weights |   RMSE |  Sigma | Score_lo
--------------------------------------------------------------------------------------------------------
mod1            | hurdle | 2644.547 |        1.00 | 2664.366 |        1.00 | 45.966 | 48.209 |   -18.20
fecund_albo.glm |    glm | 5201.769 |     < 0.001 | 5211.678 |     < 0.001 | 45.966 |  7.604 |   -29.51
```

```r
# Pairwise comparison
pairewise.f.albo = glht(fecund_albo.glm,
                        mcp(Line = "Tukey"))

f.albo <- summary(pairewise.f.albo)
f.albo
```

```
     Simultaneous Tests for General Linear Hypotheses

Multiple Comparisons of Means: Tukey Contrasts

Fit: glm(formula = "Eggs ~ Line", family = "poisson", data = fecund_albo)

Linear Hypotheses:
                       Estimate Std. Error z value Pr(>|z|)
Aal-M - BiA (WT) == 0   0.58931    0.04356  13.529   <1e-04 ***
Aal-m - BiA (WT) == 0   0.04123    0.04668   0.883    0.813
Aal-CS - BiA (WT) == 0 -0.05826    0.04892  -1.191    0.632
Aal-m - Aal-M == 0     -0.54808    0.04250 -12.897   <1e-04 ***
Aal-CS - Aal-M == 0    -0.64757    0.04495 -14.408   <1e-04 ***
Aal-CS - Aal-m == 0    -0.09949    0.04798  -2.074    0.161
---
Signif. codes:  0 '***' 0.001 '**' 0.01 '*' 0.05 '.' 0.1 ' ' 1
(Adjusted p values reported -- single-step method)
```

```r
pred.fecund.albo =
    emmeans::emmeans(mod1, specs = 'Line', mode = 'response') %>%
  data.frame() %>%
  dplyr::select(
    Line, mean.pro = emmean,
    sd.pro = SE,
    conf.inf = lower.CL,
    conf.sup = upper.CL
  )
dd =emmeans::emmeans(mod1, specs = 'Line', mode = 'count') %>%
  data.frame() %>%
  dplyr::select(
    Line, mean.pro = emmean,
    sd.pro = SE,
    conf.inf = lower.CL,
    conf.sup = upper.CL
  )
  # ggpredict(fecund_albo.glm) %>%
  # data.frame() %>%
  # dplyr::select(
  #   Line = Line.x,
  #   mean.pro = Line.predicted,
  #   sd.pro = Line.std.error,
  #   conf.inf = Line.conf.low,
  #   conf.sup = Line.conf.high)

pred.fecund.albo$Line = factor(pred.fecund.albo$Line,
                               levels = c('BiA (WT)', 'Aal-M', 'Aal-m', 'Aal-CS'))

head(pred.fecund.albo)
```

```
      Line mean.pro   sd.pro conf.inf conf.sup
1 BiA (WT) 38.30436 4.402897 29.54231 47.06640
2    Aal-M 69.05260 7.119660 54.88403 83.22118
```

```
3     Aal-m 39.91666 7.605172 24.78189 55.05144
4    Aal-CS 36.13637 5.416839 25.35652 46.91623
```

Replication

```
       Line  N
1 BiA (WT) 23
2     Aal-M 19
3     Aal-m 24
4    Aal-CS 22
```

Model fit quality

```
check_model(fecund_albo.glm)
```

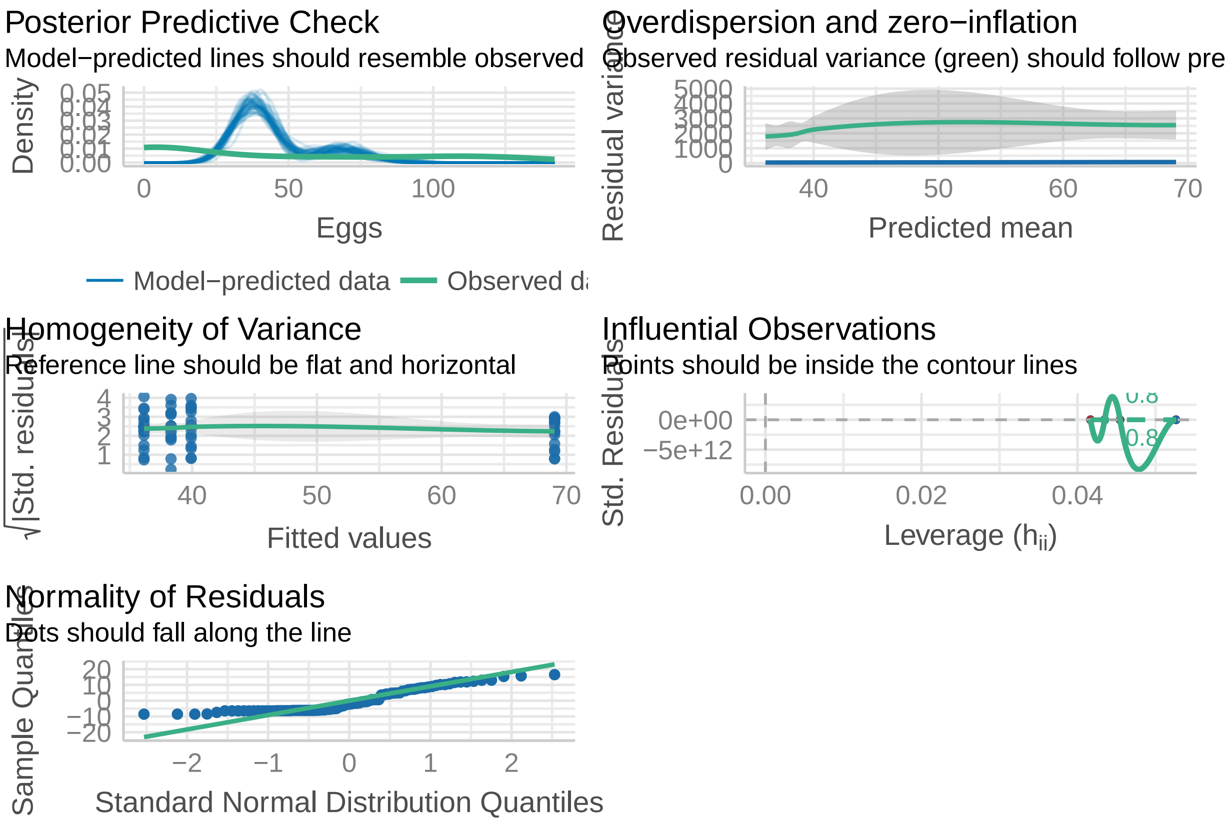

**Plot** .

```
fecund.plot.alb  = ggplot(data=NULL) +
  geom_point(data = fecund_albo,
             aes(x = jitter(as.numeric(Line)),
                 y = Eggs, color = Line),
             size = 2, show.legend = FALSE, alpha = 0.6) +
  geom_point(data = pred.fecund.albo,
             aes(x = as.numeric(Line),
```

```
                 y = mean.pro),
             size = 1.5, show.legend = FALSE,
             color = 'gray90') +
  geom_errorbar(data = pred.fecund.albo,
                aes(x = as.numeric(Line),
                    ymin = conf.inf,
                    ymax = conf.sup), width = 0.05,
                color = 'gray90') +
  geom_point(data = dd,
             aes(x = as.numeric(Line),
                 y = mean.pro),
             size = 1.5, show.legend = FALSE,
             color = 'black') +
  geom_errorbar(data = dd,
                aes(x = as.numeric(Line),
                    ymin = conf.inf,
                    ymax = conf.sup), width = 0.05,
                color = 'black') +
  annotate(geom = 'text', x= fecund.summary.alb$pos, y = -8,
           label = paste0('N = ', fecund.summary.alb$value),
           size = 3) +
  annotate(geom = 'line', x= 1:2, y=180) +
  annotate(geom = 'text', x= 1.5, y=190,
           label = '***',
           size = 3) +
  annotate(geom = 'line', x= 1:3, y=198) +
  annotate(geom = 'text', x= 2, y=210,
           label = 'n.s.',
           size = 3) +
  annotate(geom = 'line', x= 1:4, y=216) +
  annotate(geom = 'text', x= 2.5, y=228,
           label = 'n.s.',
           size = 3) +
  labs(x = 'Line', y = 'Number of eggs per female',
       color = "Line") +
  scale_y_continuous(breaks = seq(0,175,25),limits = c(-10,230)) +
  scale_color_manual(values = line.color.alb,
                     labels = names(line.color.alb)) +
  scale_x_continuous(guide = guide_axis(angle = 30), breaks = 1:4,
                     labels = c("BiA (WT)",
                                "Aal-M",
                                "Aal-m",
                                "Aal-CS")) +
  theme_classic() +
  theme(legend.position = 'none',
        panel.background = element_rect(fill = "grey"))

fecund.plot.alb
```

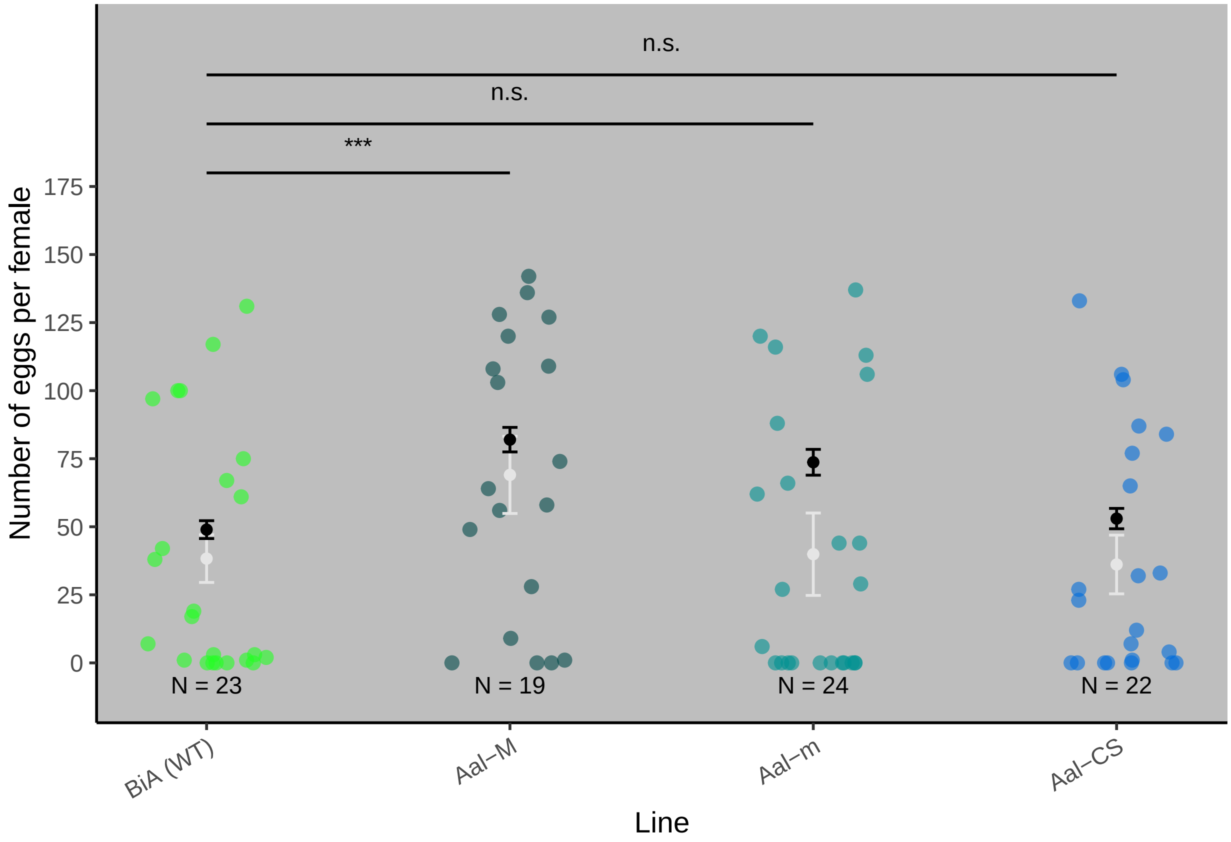

#### Larva survival

##### Aegypti

###### Data .

```
surv.aeg.larv = read.table("data/Survival-larvae_aeg.csv",
                           sep = ";", header = T)
surv.aeg.larv = surv.aeg.larv %>%
  dplyr::select( Replicate, Line, larvae, adults)

surv.aeg.larv$Species = "Ae. aegypti"

surv.aeg.larv$Line = factor(surv.aeg.larv$Line,
                            levels = c('Bra (WT)', 'Aaeg-M', 'Aaeg-m', 'Aaeg-CS'))

surv.aeg.larv$survie = surv.aeg.larv$adults/surv.aeg.larv$larvae * 100

head(surv.aeg.larv)
```

```
  Replicate   Line larvae adults     Species survie
1         1 Aaeg-M    100     87 Ae. aegypti     87
2         2 Aaeg-M    100     82 Ae. aegypti     82
3         3 Aaeg-M    100     93 Ae. aegypti     93
```

```
4         4 Aaeg-M    100     73 Ae. aegypti     73
5         1 Aaeg-m    100     96 Ae. aegypti     96
6         2 Aaeg-m    100    100 Ae. aegypti    100
```

##### Model .

```
mod.tot.sl.aeg = lm(data=surv.aeg.larv, formula = survie ~ Line)
mod.tot.sl.aeg %>% summary()
```

```

Call:
lm(formula = survie ~ Line, data = surv.aeg.larv)

Residuals:
    Min      1Q  Median      3Q     Max 
-16.625  -2.344   0.625   4.125   7.375 

Coefficients:
            Estimate Std. Error t value Pr(>|t|)    
(Intercept)   94.500      3.127  30.225 1.52e-15 ***
LineAaeg-M    -4.875      3.829  -1.273    0.221    
LineAaeg-m     1.000      4.422   0.226    0.824    
LineAaeg-CS    3.750      4.422   0.848    0.409    
---
Signif. codes:  0 '***' 0.001 '**' 0.01 '*' 0.05 '.' 0.1 ' ' 1

Residual standard error: 6.253 on 16 degrees of freedom
Multiple R-squared:  0.2691,	Adjusted R-squared:  0.1321 
F-statistic: 1.964 on 3 and 16 DF,  p-value: 0.1601
```

```
mod.tot.sl.aeg %>% aov %>% TukeyHSD()
```

```

  Tukey multiple comparisons of means
    95% family-wise confidence level

Fit: aov(formula = .)

$Line
                   diff        lwr       upr     p adj
Aaeg-M-Bra (WT)  -4.875 -15.830534  6.080534 0.5920067
Aaeg-m-Bra (WT)   1.000 -11.650361 13.650361 0.9957513
Aaeg-CS-Bra (WT)  3.750  -8.900361 16.400361 0.8308209
Aaeg-m-Aaeg-M     5.875  -5.080534 16.830534 0.4414047
Aaeg-CS-Aaeg-M    8.625  -2.330534 19.580534 0.1514294
Aaeg-CS-Aaeg-m    2.750  -9.900361 15.400361 0.9235384
```

```
df.larv.aeg = ggpredict(mod.tot.sl.aeg) %>%
  data.frame %>%
  dplyr::select(
    Line = Line.x,
    m = Line.predicted,
    sd = Line.std.error,
    conf.inf = Line.conf.low,
```

```
    conf.sup = Line.conf.high
  )
df.larv.aeg$Line = factor(df.larv.aeg$Line,
                          levels = c('Bra (WT)', 'Aaeg-M', 'Aaeg-m', 'Aaeg-CS'))

head(df.larv.aeg)
```

```
      Line      m       sd conf.inf  conf.sup
1 Bra (WT) 94.500 3.126562 88.37205 100.62795
2   Aaeg-M 89.625 2.210813 85.29189  93.95811
3   Aaeg-m 95.500 3.126562 89.37205 101.62795
4  Aaeg-CS 98.250 3.126562 92.12205 104.37795
```

Replication

```
# A tibble: 4 x 3
  Line         N     n
  <fct>    <int> <int>
1 Bra (WT)     4   800
2 Aaeg-M       8   800
3 Aaeg-m       4   400
4 Aaeg-CS      4   400
```

Model quality

.

```
check_model(mod.tot.sl.aeg)
```

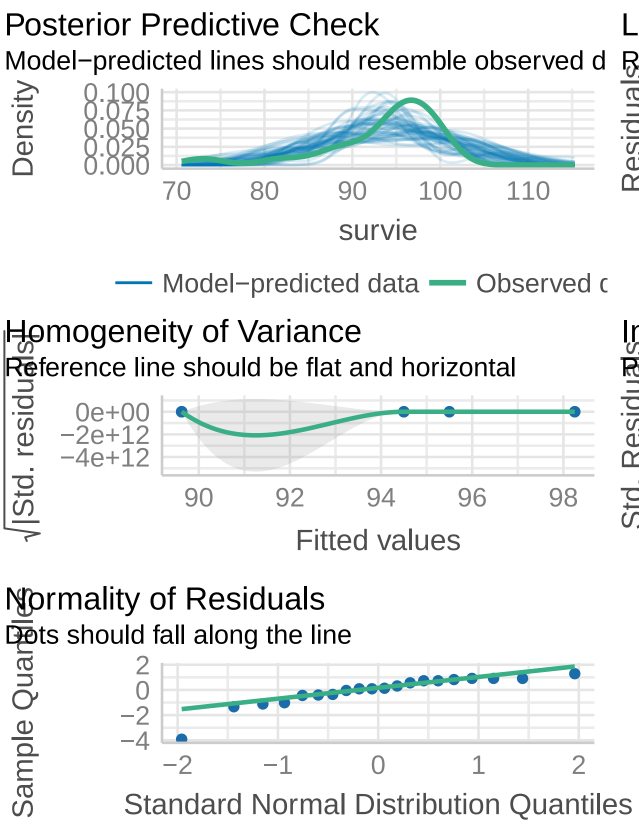

```
larval.surv.plot.aeg = ggplot(data = NULL,
                              aes()) +
  geom_point(data = surv.aeg.larv,
             aes(x = jitter(as.numeric(factor(Line))),
                 y = survie, color = Line),
             size = 2, alpha = 0.6) +
  geom_point(data = df.larv.aeg,
             aes(x=Line, y=m), col = 'black') +
  geom_errorbar(data = df.larv.aeg, aes(x=Line, ymin = conf.inf,
                                         ymax = conf.sup),
                width = 0.05, col = 'black') +
  annotate(geom = 'text', x= 1:4, y=0,
           label = c("N = 4", "N = 8", "N = 4", "N = 4"),
           size = 3) +
  annotate(geom = 'text', x= 1:4, y=-10,
           label = c("n = 800", "n = 400", "n = 400", "n = 400"),
           size = 3) +
  annotate(geom = 'line', x= 1:4, y=118) +
  annotate(geom = 'text', x= 2.5, y=123,
           label = 'n.s.',
           size = 3) +
  annotate(geom = 'line', x= 1:3, y=111) +
  annotate(geom = 'text', x= 2, y=116,
```

```
          label = 'n.s.',
          size = 3) +
  annotate(geom = 'line', x= 1:2, y=103) +
  annotate(geom = 'text', x= 1.5, y=109,
          label = 'n.s.',
          size = 3) +
  scale_x_discrete(limits = levels(df.larv.aeg$Line), labels = c("Bra (WT)",
                                                                  "Aaeg-M",
                                                                  "Aaeg-m",
                                                                  "Aaeg-CS"
  ),
  guide = guide_axis(angle = 30)) +
  scale_y_continuous(breaks = c(0, 25, 50, 75, 100),
                     limits = c(-20, 126), labels = c(0, 25, 50, 75, 100)) +
  scale_color_manual(labels = names(line.color.aeg),
                     values = line.color.aeg) +
  labs(x = 'Line',
       y = 'Larva to adult survival (%)',
       color = 'Line') +
  theme_classic() +
  theme(legend.position = 'none')

larval.surv.plot.aeg
```

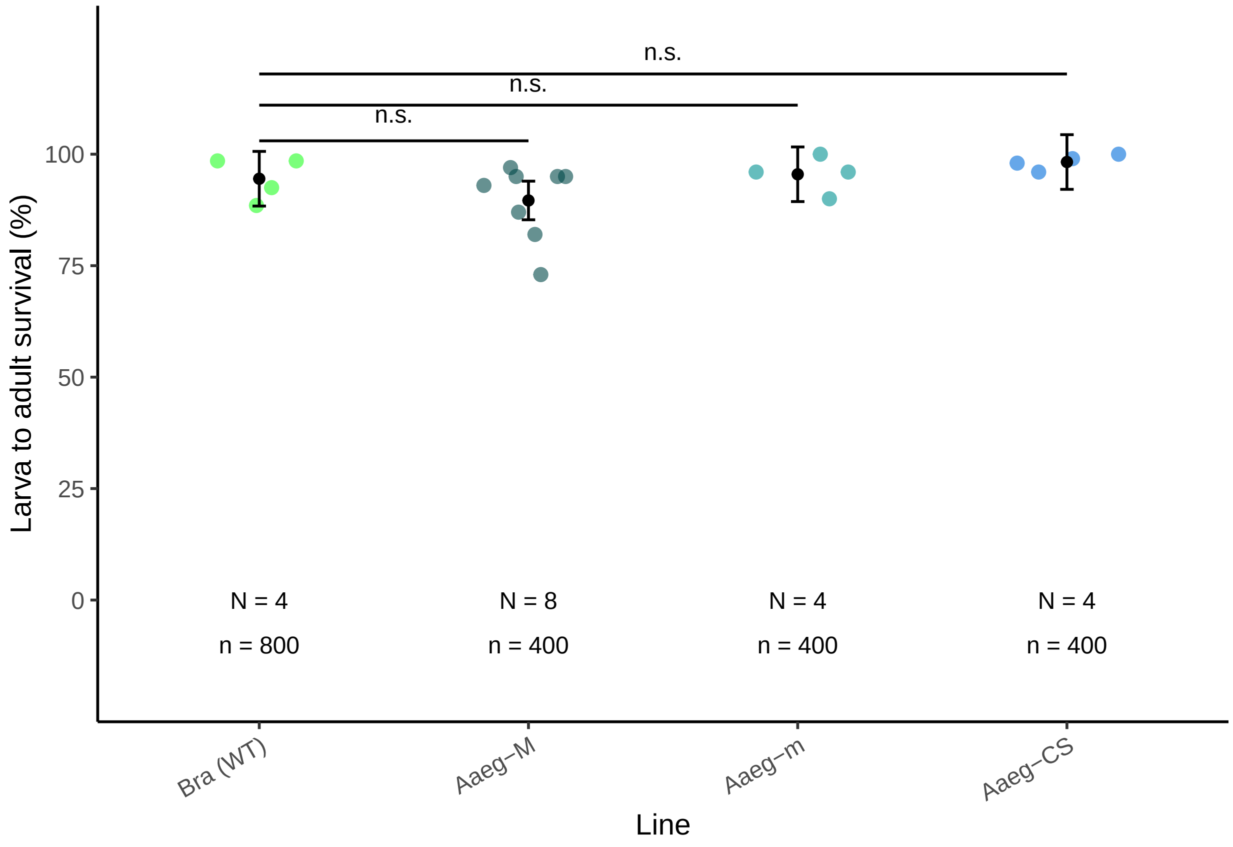

**Plot**

##### Albopictus

###### Data .

```
surv.alb.larv = read.table("data/Survival-larvae_albo.csv", sep = ";", header = T)
surv.alb.larv = surv.alb.larv %>%
  dplyr::select( Replicate, Line, larvae, adults)

surv.alb.larv$Species = "Ae. albopictus"

surv.alb.larv$Line = factor(surv.alb.larv$Line,
                            levels = c('BiA (WT)', 'Aal-M', 'Aal-m', 'Aal-CS'))

surv.alb.larv$survie = surv.alb.larv$adults/surv.alb.larv$larvae * 100

head(surv.alb.larv)
```

```
  Replicate     Line larvae adults        Species survie
1         1 BiA (WT)    200    103 Ae. albopictus   51.5
2         2 BiA (WT)    200    141 Ae. albopictus   70.5
3         3 BiA (WT)    200    129 Ae. albopictus   64.5
4         4 BiA (WT)    200    130 Ae. albopictus   65.0
5         1    Aal-M    100     88 Ae. albopictus   88.0
6         2    Aal-M    100     79 Ae. albopictus   79.0
```

###### Model .

```
mod.tot.sl.alb = lm(data=surv.alb.larv, formula = survie ~ Line)
mod.tot.sl.alb %>% summary()
```

```
Call:
lm(formula = survie ~ Line, data = surv.alb.larv)

Residuals:
    Min      1Q  Median      3Q     Max 
-11.375  -2.750   1.062   2.750   7.625 

Coefficients:
            Estimate Std. Error t value Pr(>|t|)    
(Intercept)   62.875      2.831  22.210 4.09e-11 ***
LineAal-M     18.125      4.004   4.527 0.000693 ***
LineAal-m     16.625      4.004   4.153 0.001341 ** 
LineAal-CS     5.625      4.004   1.405 0.185378    
---
Signif. codes:  0 '***' 0.001 '**' 0.01 '*' 0.05 '.' 0.1 ' ' 1

Residual standard error: 5.662 on 12 degrees of freedom
Multiple R-squared:  0.7043,    Adjusted R-squared:  0.6303 
F-statistic: 9.525 on 3 and 12 DF,  p-value: 0.001695
```

```
mod.tot.sl.alb %>% aov %>% TukeyHSD()
```

```
  Tukey multiple comparisons of means
    95% family-wise confidence level

Fit: aov(formula = .)

$Line
                   diff        lwr        upr     p adj
Aal-M-BiA (WT)   18.125   6.238769 30.0112306 0.0033427
Aal-m-BiA (WT)   16.625   4.738769 28.5112306 0.0063439
Aal-CS-BiA (WT)   5.625  -6.261231 17.5112306 0.5198078
Aal-m-Aal-M      -1.500 -13.386231 10.3862306 0.9812185
Aal-CS-Aal-M    -12.500 -24.386231 -0.6137694 0.0383297
Aal-CS-Aal-m    -11.000 -22.886231  0.8862306 0.0730385
```

```
df.larv.alb = ggpredict(mod.tot.sl.alb) %>%
  data.frame %>%
  dplyr::select(
    Line = Line.x,
    m = Line.predicted,
    sd = Line.std.error,
    conf.inf = Line.conf.low,
    conf.sup = Line.conf.high
  )
df.larv.alb$Line = factor(df.larv.alb$Line,
                          levels = c('BiA (WT)', 'Aal-M', 'Aal-m', 'Aal-CS'))
head(df.larv.alb)
```

```
      Line      m       sd conf.inf conf.sup
1 BiA (WT) 62.875 2.830958 57.32642 68.42358
2    Aal-M 81.000 2.830958 75.45142 86.54858
3    Aal-m 79.500 2.830958 73.95142 85.04858
4   Aal-CS 68.500 2.830958 62.95142 74.04858
```

Replication

```
# A tibble: 4 x 3
  Line         N     n
  <fct>    <int> <int>
1 BiA (WT)     4   800
2 Aal-M        4   400
3 Aal-m        4   400
4 Aal-CS       4   400
```

Model quality

```
check_model(mod.tot.sl.alb)
```

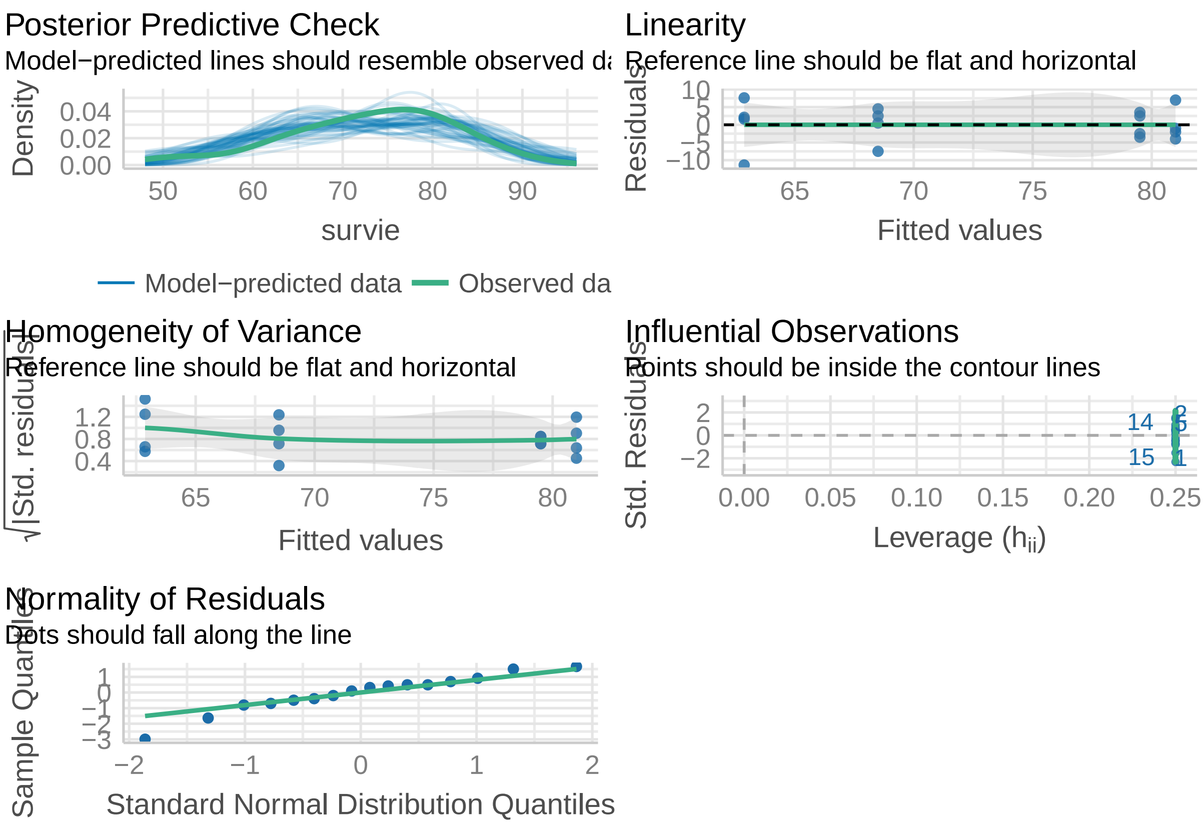

**Plot** .

```
larval.surv.plot.alb = ggplot(data = NULL,
                              aes()) +
  geom_point(data = surv.alb.larv,
             aes(x = jitter(as.numeric(factor(Line))),
                 y = survie, color = Line),
             size = 2, alpha = 0.6) +
  geom_point(data = df.larv.alb,
             aes(x=Line, y=m), col = 'black') +
  geom_errorbar(data = df.larv.alb, aes(x=Line, ymin = conf.inf,
                                        ymax = conf.sup),
                width = 0.05, col = 'black') +
  annotate(geom = 'text', x= 1:4, y=0,
           label = c("N = 4", "N = 4", "N = 4", "N = 4"),
           size = 3) +
  annotate(geom = 'text', x= 1:4, y=-10,
           label = c("n = 800", "n = 400", "n = 400", "n = 400"),
           size = 3) +
  annotate(geom = 'line', x= 1:4, y=118) +
  annotate(geom = 'text', x= 2.5, y=123,
           label = 'n.s.',
           size = 3) +
  annotate(geom = 'line', x= 1:3, y=111) +
  annotate(geom = 'text', x= 2, y=114,
```

```
          label = '**',
          size = 3) +
  annotate(geom = 'line', x= 1:2, y=103) +
  annotate(geom = 'text', x= 1.5, y=106,
          label = '**',
          size = 3) +
  scale_x_discrete(limits = levels(df.larv.alb$Line), labels = c("BiA (WT)",
                                                                  "Aal-M",
                                                                  "Aal-m",
                                                                  "Aal-CS"
  ),
  guide = guide_axis(angle = 30)) +
  scale_y_continuous(breaks = c(0, 25, 50, 75, 100),
                     limits = c(-20, 126), labels = c(0, 25, 50, 75, 100)) +
  scale_color_manual(labels = names(line.color.alb),
                     values = line.color.alb) +
  labs(x = 'Line',
       y = 'Larva to adult survival (%)',
       color = 'Line') +
  theme_classic() +
  theme(legend.position = 'none',
        panel.background = element_rect(fill= "grey"))

larval.surv.plot.alb
```

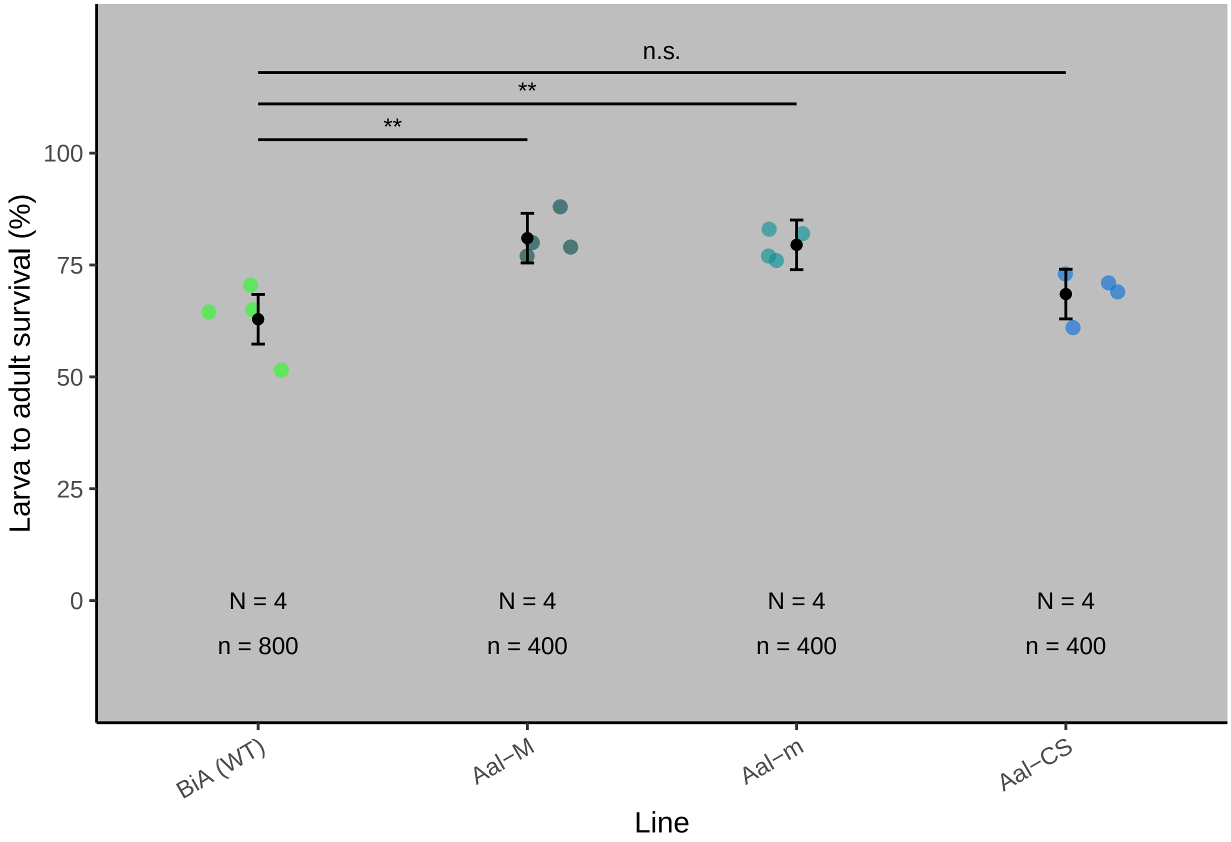

### Figure 4

#### Competitiveness

##### Aegypti

**Data** .

```
compet_aeg<-read.table("data/Competitiveness_aeg.csv", header=TRUE, sep=",")
compet_aeg$total = compet_aeg$tsg + compet_aeg$no_tsg
compet_aeg$comp = compet_aeg$tsg / (compet_aeg$total) * 100
compet_aeg$line =  factor(x = compet_aeg$line,
                          levels = c('Expected','Aaeg-M', 'Aaeg-CS'))
head(compet_aeg)
```

```
      line Replicate  tsg no_tsg total     comp
1 Expected         1 1300   3700  5000 26.00000
2   Aaeg-M         1  632   1758  2390 26.44351
3   Aaeg-M         2  618   1670  2288 27.01049
4   Aaeg-M         3  740   1472  2212 33.45389
5   Aaeg-M         4  493   1775  2268 21.73721
6   Aaeg-M         5  522   1554  2076 25.14451
```

```
compet.aeg =
  rbind(
    data.frame(replicate = compet_aeg[1,2],
               line = compet_aeg[1,1],
               result = rep(1, compet_aeg[1,3])),
    data.frame(replicate = compet_aeg[1,2],
               line = compet_aeg[1,1],
               result = rep(0, compet_aeg[1,4])))

for(i in 2:nrow(compet_aeg)){
  compet.aeg =
    rbind(
      compet.aeg,
      data.frame(replicate = compet_aeg[i,2],
                 line = compet_aeg[i,1],
                 result = rep(1, compet_aeg[i,3])),
      data.frame(replicate = compet_aeg[i,2],
                 line = compet_aeg[i,1],
                 result = rep(0, compet_aeg[i,4]))
    )
}

compet.aeg$line = factor(x = compet.aeg$line,
                          levels = c('Expected','Aaeg-M', 'Aaeg-CS'))
compet.aeg$Species = "Ae. aegypti"

sum.compet.aeg = compet.aeg %>%
  filter(line != 'Expected') %>%
  group_by(line) %>%
  dplyr::summarize(value = n(),
```

```
           N = max(replicate)) %>%
  data.frame() %>%
  mutate(pos = c(1,2))

head(compet.aeg)
```

```
  replicate     line result     Species
1         1 Expected      1 Ae. aegypti
2         1 Expected      1 Ae. aegypti
3         1 Expected      1 Ae. aegypti
4         1 Expected      1 Ae. aegypti
5         1 Expected      1 Ae. aegypti
6         1 Expected      1 Ae. aegypti
```

```
Expected_Aaeg_M = 0.26
Expected_Aaeg_CS = 0.26
```

##### Model .

```
# Generalized linear model with Bernouilli distribution
cpt.aeg = glm(formula = 'result ~ line',
             family = binomial,
             data = compet.aeg)
summary(cpt.aeg)
```

```
Call:
glm(formula = "result ~ line", family = binomial, data = compet.aeg)

Deviance Residuals:
    Min       1Q   Median       3Q      Max
-0.8044  -0.7890  -0.7890   1.6037   1.6414

Coefficients:
            Estimate Std. Error z value Pr(>|z|)
(Intercept) -1.04597    0.03224 -32.442   <2e-16 ***
lineAaeg-M   0.03858    0.03865   0.998   0.3182
lineAaeg-CS  0.08358    0.04388   1.905   0.0568 .
---
Signif. codes:  0 '***' 0.001 '**' 0.01 '*' 0.05 '.' 0.1 ' ' 1

(Dispersion parameter for binomial family taken to be 1)

    Null deviance: 25437  on 21877  degrees of freedom
Residual deviance: 25433  on 21875  degrees of freedom
AIC: 25439

Number of Fisher Scoring iterations: 4
```

```
# Pairwise comparison
cpt.aeg.pred = ggpredict(cpt.aeg) %>% data.frame()
```

```
cpt.aeg.pred %>% dplyr::select(Line = line.x,
                               Mean = line.predicted, SE = line.std.error,
                               CI.low = line.conf.low, CI.high = line.conf.high)
```

```
      Line      Mean         SE    CI.low   CI.high
1 Expected 0.2600000 0.03224127 0.2480275 0.2723411
2   Aaeg-M 0.2674915 0.02131430 0.2593860 0.2757561
3  Aaeg-CS 0.2763997 0.02976378 0.2648858 0.2882179
```

Replication

```
     Line N     n
1  Aaeg-M 5 11234
2 Aaeg-CS 5  5644
```

Model fit quality

```
check_model(cpt.aeg)
```

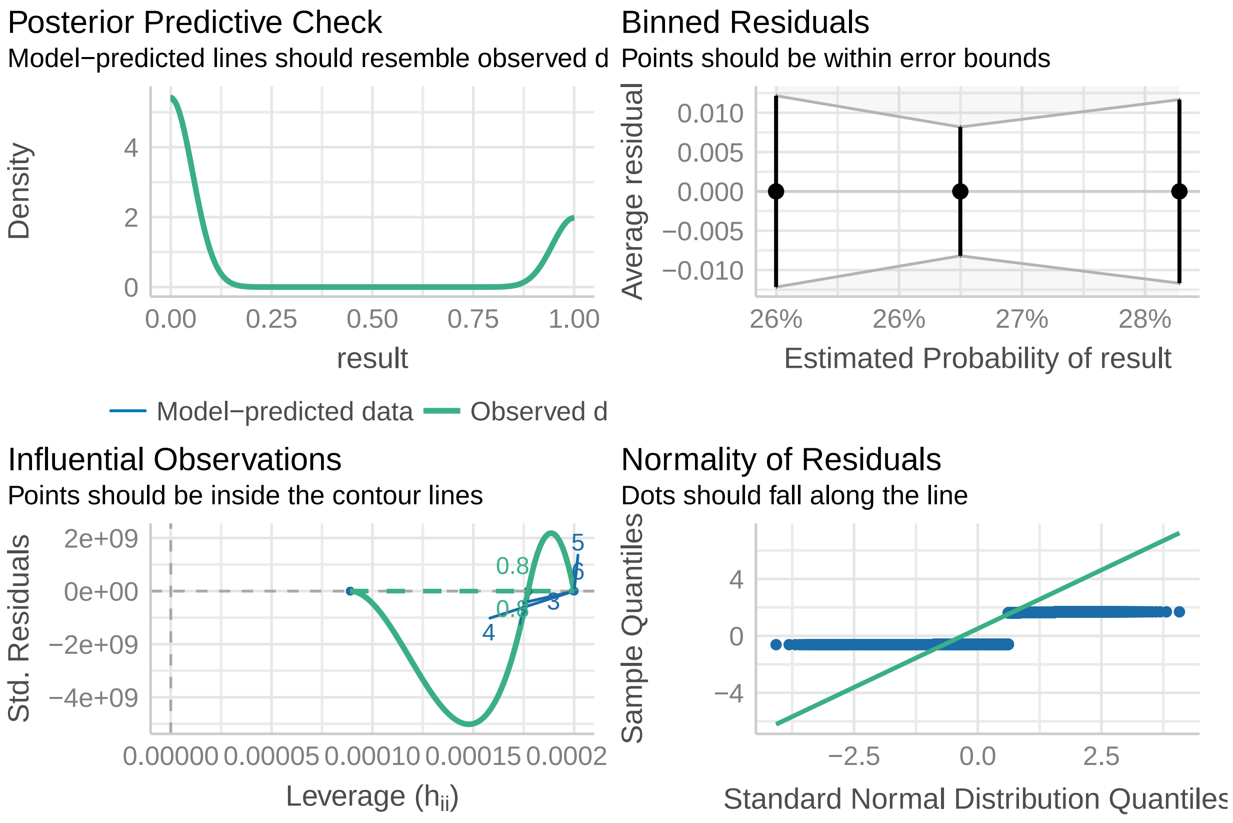

**Plot** .

```
comp.aeg.plot =
  ggplot(data =  NULL) +
  geom_point(data = compet_aeg[compet_aeg$line!='Expected',],
             aes(x = jitter(as.numeric(factor(line))),
                 y = comp,
                 color = line),
             size = 2, alpha = .6)+
  geom_point(data = cpt.aeg.pred[-1,],
             aes(x = (as.numeric(line.x)-1),
                 y = line.predicted * 100),
             size = 1.5, color = 'black') +
  geom_errorbar(data = cpt.aeg.pred[-1,],
                aes(x = (as.numeric(line.x)-1),
                    ymin = line.conf.low* 100,
                    ymax = line.conf.high* 100),
                width = 0.05, color = 'black') +
  geom_point(data = compet.aeg,
             aes(x = jitter((as.numeric(factor(line))-1), factor = 2),
                 y = jitter((result * 95), factor = .25),
                 color = line),
             size = .1) +
  annotate(geom = 'text', x = sum.compet.aeg$pos, y=-20,
           label = paste0('n = ', sum.compet.aeg$value),
           size = 3) +
  annotate(geom = 'text', x= sum.compet.aeg$pos, y=-10,
           label = c('N = 5', 'N = 5'),
           size = 3) +
  annotate(geom = 'line', x = c(.75, 1.25),
           y=rep(Expected_Aaeg_M*100, 2),
           lty = 2, color = 'grey30')  +
  annotate(geom = 'line', x = c(1.75, 2.25),
           y=rep(Expected_Aaeg_CS*100, 2),
           lty = 2, color = 'grey30') +
  annotate(geom = 'text', x= 2, y=110,
           label = 'ns.',
           size = 3) +
  annotate(geom = 'text', x= 1, y=110,
           label = 'ns.',
           size = 3) +
  scale_x_continuous(breaks = 1:2,
                     labels = c("Bra (WT) vs. Aaeg-M",
                                "Aaeg-CS vs. Aaeg-M"),
                     limits = c(.5,2.5)) +
  scale_y_continuous(sec.axis = sec_axis(~., breaks = c(0,95),
                                         labels = c('Non-transgenic', 'Transgenic'),
                                         name = 'Progeny characterization (binomial)'),
                     limits = c(-20,120),
                     breaks = c(0,25,50,75,100)) +
  scale_color_manual(values = line.color.aeg, labels = names(line.color.aeg)) +
  labs(x = 'Line ',
       y = 'Transgenic progeny (%)') +
  theme_classic() +
  theme(legend.position = 'none',
```

```
        axis.title.y.right = element_text(color = 'black'),
        axis.ticks.y.right = element_line(color = 'black'),
        axis.line.y.right = element_line(color = 'black'),
        axis.text.y.right = element_text(angle = -45, hjust = 0, vjust = 0))

comp.aeg.plot
```

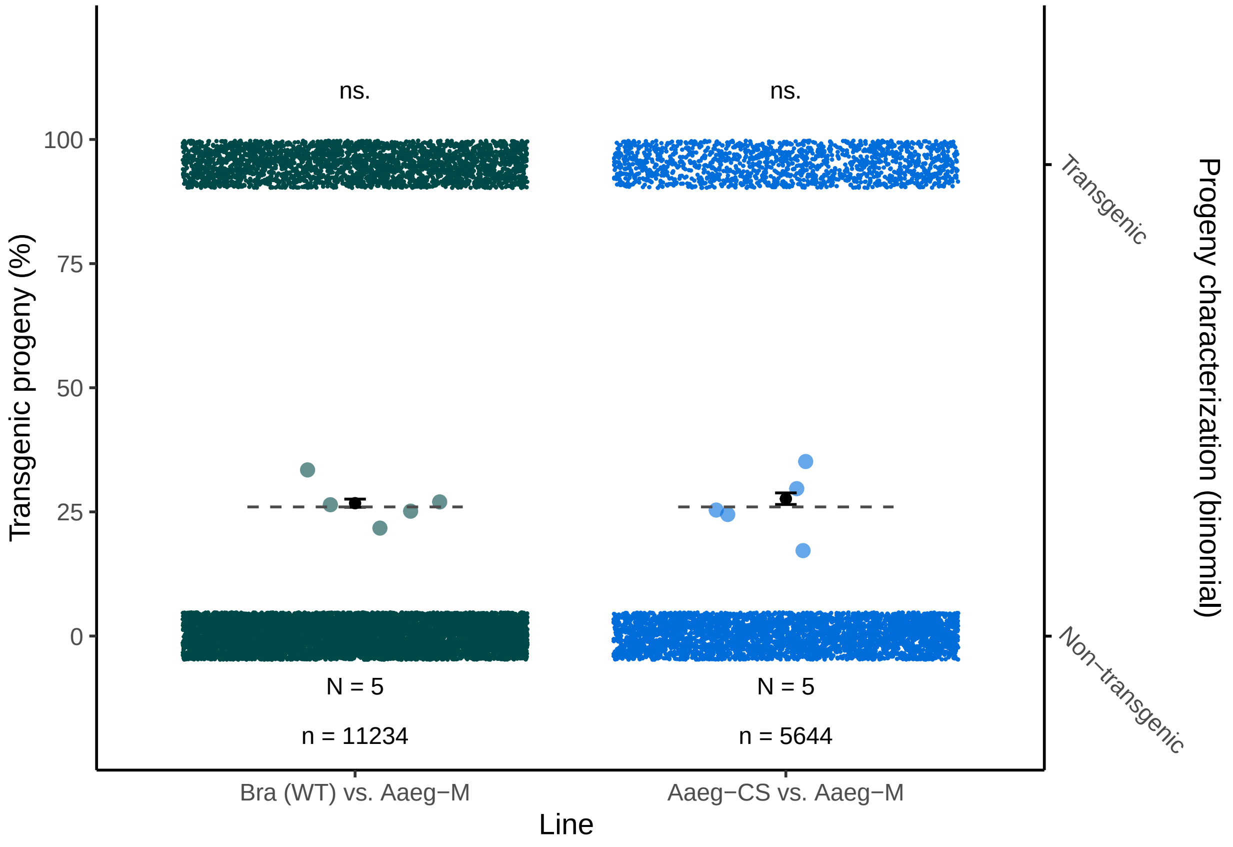

**Albopictus**

**Data** .

```
compet_albo<-read.table("data/Competitiveness_albo.csv", header=TRUE, sep=",")
compet_albo$total = compet_albo$tsg + compet_albo$no_tsg
compet_albo$comp = compet_albo$tsg / (compet_albo$total) * 100
compet_albo$line =  factor(x = compet_albo$line,
                           levels = c('Expected','Aal-M', 'Aal-CS'))
head(compet_albo)
```

```
      line Replicate tsg no_tsg total     comp
1 Expected         1 637   1863  2500 25.48000
2    Aal-M         1  75    185   260 28.84615
3    Aal-M         2  60    247   307 19.54397
4    Aal-M         3 119    335   454 26.21145
5    Aal-M         4 151    887  1038 14.54721
```

```
6    Aal-M         5  67    451   518 12.93436
```

```
compet.albo =
  rbind(
    data.frame(replicate = compet_albo[1,2],
               line = compet_albo[1,1],
               result = rep(1, compet_albo[1,3])),
    data.frame(replicate = compet_albo[1,2],
               line = compet_albo[1,1],
               result = rep(0, compet_albo[1,4])))

for(i in 2:nrow(compet_albo)){
  compet.albo =
    rbind(
      compet.albo,
      data.frame(replicate = compet_albo[i,2],
                 line = compet_albo[i,1],
                 result = rep(1, compet_albo[i,3])),
      data.frame(replicate = compet_albo[i,2],
                 line = compet_albo[i,1],
                 result = rep(0, compet_albo[i,4]))
    )
}

compet.albo$line = factor(x = compet.albo$line,
                            levels = c('Expected','Aal-M', 'Aal-CS'))
compet.albo$Species = "Ae. albopictus"

sum.compet.albo = compet.albo %>%
  filter(line != "Expected") %>%
  group_by(line) %>%
  dplyr::summarize(value = n(),
            N = max(replicate)) %>%
  data.frame() %>%
  mutate(pos = c(1,2))

head(compet.albo)
```

```
  replicate     line result        Species
1         1 Expected      1 Ae. albopictus
2         1 Expected      1 Ae. albopictus
3         1 Expected      1 Ae. albopictus
4         1 Expected      1 Ae. albopictus
5         1 Expected      1 Ae. albopictus
6         1 Expected      1 Ae. albopictus
```

```
Expected_Aal_M = 0.255
Expected_Aal_CS = 0.255
```

##### Model .

```r
# Generalized linear model with Bernouilli distribution
cpt.albo = glm(formula = 'result ~ line',
             family = binomial,
             data = compet.albo)
summary(cpt.albo)
```

```
Call:
glm(formula = "result ~ line", family = binomial, data = compet.albo)

Deviance Residuals:
    Min       1Q   Median       3Q      Max
-0.7835  -0.7835  -0.7670  -0.6361   1.8425

Coefficients:
            Estimate Std. Error z value Pr(>|z|)
(Intercept) -1.07317    0.04590 -23.382  < 2e-16 ***
lineAal-M   -0.42192    0.06856  -6.154 7.55e-10 ***
lineAal-CS   0.04939    0.05174   0.955     0.34
---
Signif. codes:  0 '***' 0.001 '**' 0.01 '*' 0.05 '.' 0.1 ' ' 1

(Dispersion parameter for binomial family taken to be 1)

    Null deviance: 15773  on 14085  degrees of freedom
Residual deviance: 15697  on 14083  degrees of freedom
AIC: 15703

Number of Fisher Scoring iterations: 4
```

```r
# Pairwise comparison
cpt.albo.pred = ggpredict(cpt.albo) %>% data.frame()

cpt.albo.pred %>% dplyr::select(Line = line.x,
                                Mean = line.predicted, SE = line.std.error,
                                CI.low = line.conf.low, CI.high = line.conf.high)
```

```
      Line      Mean         SE    CI.low   CI.high
1 Expected 0.2548000 0.04589797 0.2380986 0.2722543
2    Aal-M 0.1831587 0.05092841 0.1686943 0.1985671
3   Aal-CS 0.2642913 0.02389280 0.2552868 0.2734967
```

Replication

```
    Line N    n
1  Aal-M 5 2577
2 Aal-CS 5 9009
```

Model fit quality

```r
plot(cpt.albo)
```

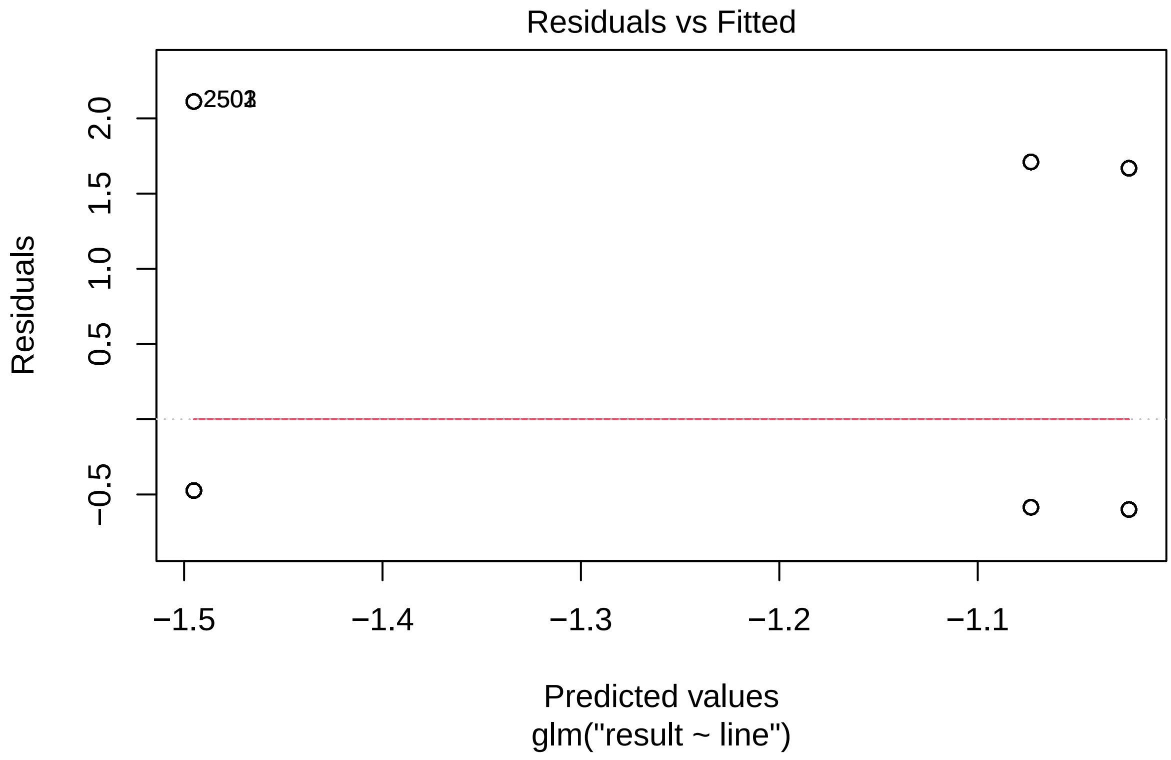
Residuals vs Fitted
2501
Residuals
2.0
1.5
1.0
0.5
−0.5
−1.5
−1.4
−1.3
−1.2
−1.1
Predicted values
glm("result ~ line")

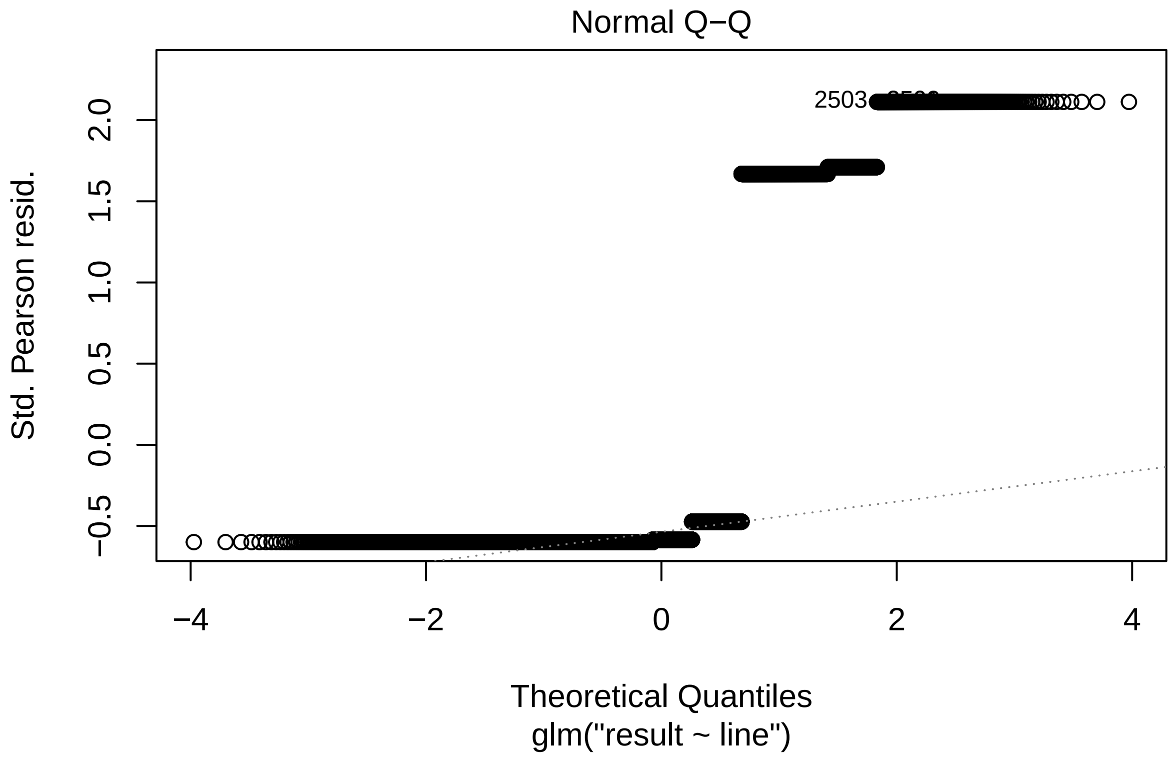
Normal Q−Q
Std. Pearson resid.
2.0
1.5
1.0
0.5
0.0
−0.5
2503
−4
−2
0
2
4
Theoretical Quantiles
glm("result ~ line")

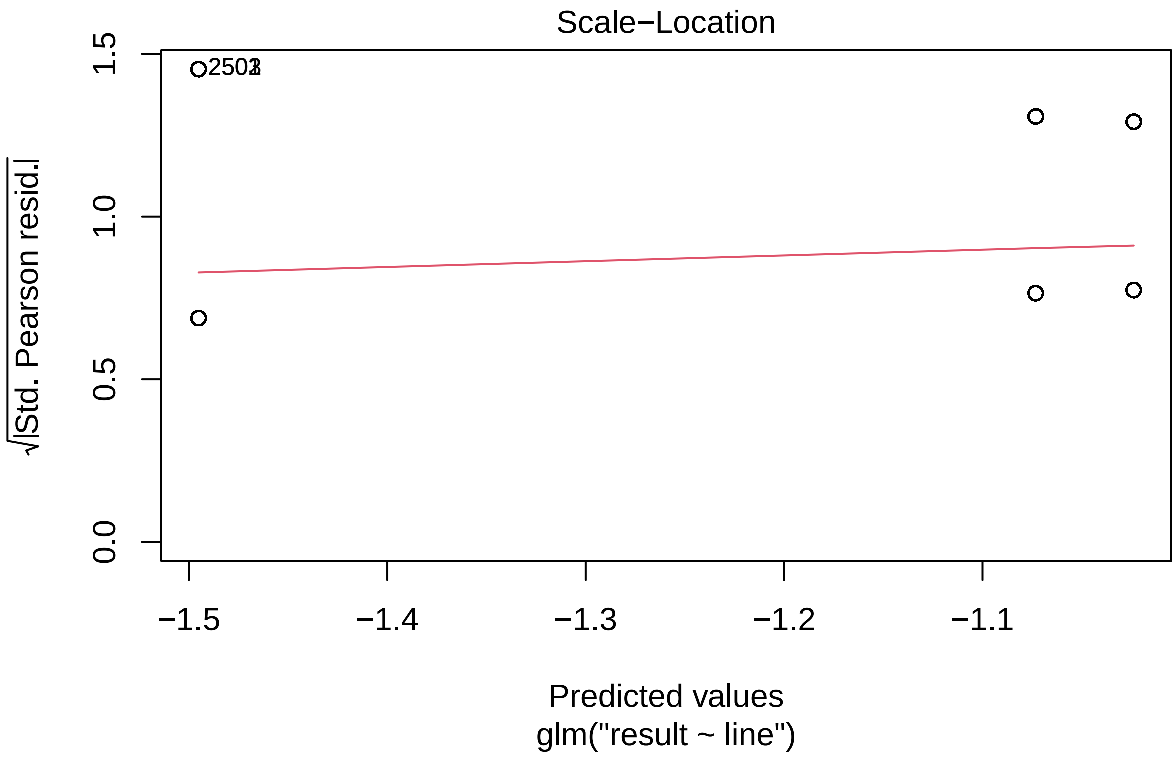
Scale−Location
√|Std. Pearson resid.|
1.5
1.0
0.5
0.0
−1.5
−1.4
−1.3
−1.2
−1.1
Predicted values
glm("result ~ line")

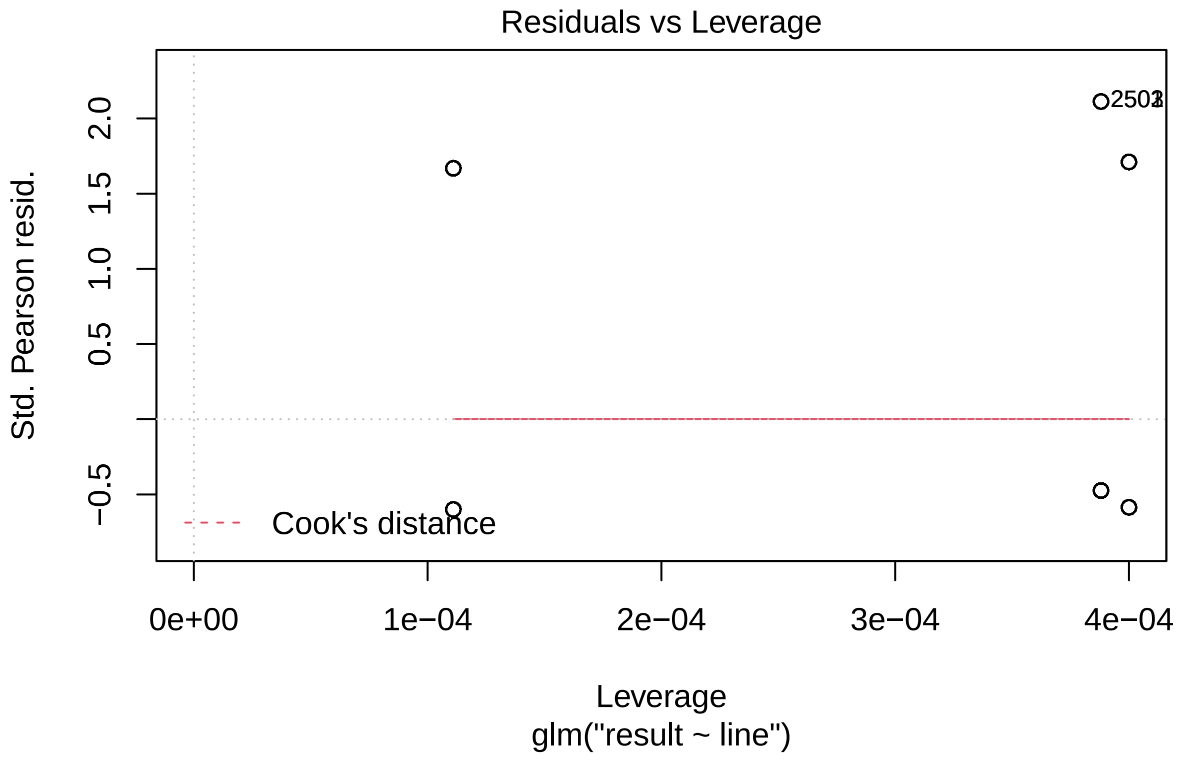

**Plot** .

```
comp.albo.plot =
  ggplot(data =  NULL) +
  geom_point(data = compet_albo[compet_albo$line!='Expected',],
             aes(x = jitter(as.numeric(factor(line))),
                 y = comp,
                 color = line),
             size = 2, alpha = .6)+
  geom_point(data = cpt.albo.pred[-1,],
             aes(x = (as.numeric(line.x)-1),
                 y = line.predicted * 100),
             size = 1.5, color = 'black') +
  geom_errorbar(data = cpt.albo.pred[-1,],
                aes(x = (as.numeric(line.x)-1),
                    ymin = line.conf.low* 100,
                    ymax = line.conf.high* 100),
                width = 0.05, color = 'black') +
  geom_point(data = compet.albo,
             aes(x = jitter((as.numeric(factor(line))-1), factor = 2),
                 y = jitter((result * 95), factor = .25),
                 color = line),
             size = .1) +
  annotate(geom = 'text', x = sum.compet.albo$pos, y=-20,
           label = paste0('n = ', sum.compet.albo$value),
```

```
          size = 3) +
  annotate(geom = 'text', x= sum.compet.albo$pos, y=-10,
           label = c('N = 5', 'N = 5'),
           size = 3) +
  annotate(geom = 'line', x = c(.75, 1.25),
           y=rep(Expected_Aal_M*100, 2),
           lty = 2, color = 'grey30')  +
  annotate(geom = 'line', x = c(1.75, 2.25),
           y=rep(Expected_Aal_CS*100, 2),
           lty = 2, color = 'grey30') +
  annotate(geom = 'text', x= 2, y=110,
           label = 'n.s.',
           size = 3) +
  annotate(geom = 'text', x= 1, y=110,
           label = '***',
           size = 3) +
  scale_x_continuous(breaks = 1:2,
                     labels = c("BiA (WT) vs. Aal-M",
                                "Aal-CS vs. Aal-M"),
                     limits = c(.5,2.5)) +
  scale_y_continuous(sec.axis = sec_axis(~., breaks = c(0,95),
                                         labels = c('Non-transgenic', 'Transgenic'),
                                         name = 'Progeny characterization (binomial)'),
                     limits = c(-20,120),
                     breaks = c(0,25,50,75,100)) +
  scale_color_manual(values = line.color.alb, labels = names(line.color.alb)) +
  labs(x = 'Line ',
       y = 'Transgenic progeny (%)') +
  theme_classic() +
  theme(legend.position = 'none',
        axis.title.y.right = element_text(color = 'black'),
        axis.ticks.y.right = element_line(color = 'black'),
        axis.line.y.right = element_line(color = 'black'),
        axis.text.y.right = element_text(angle = -45, hjust = 0, vjust = 0),
        panel.background = element_rect(fill = 'grey'))

comp.albo.plot
```

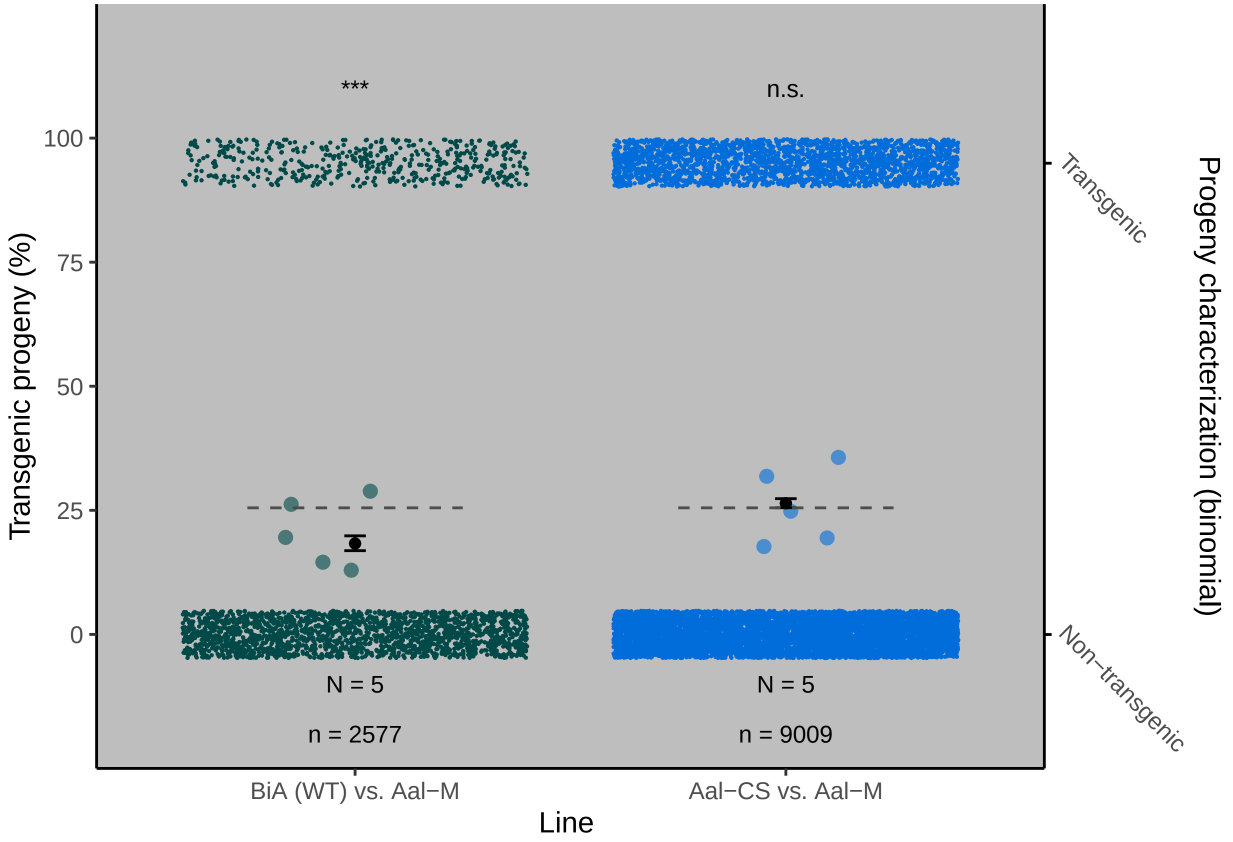

#### Flight test

##### Aegypti

###### Data .

```
flight_aegypti<-read.table("data/Flight_aeg.csv", header=TRUE, sep=";")
flight_aegypti$esc = flight_aegypti$out /
  (flight_aegypti$out + flight_aegypti$in.) * 100
flight_aegypti$Line = factor(flight_aegypti$Line,
                             levels = c("Bra (WT)", "Aaeg-M", "Aaeg-CS"))
head(flight_aegypti)
```

```
  Replicate     Line out in.      esc
1         2   Aaeg-M  65  36 64.35644
2         2  Aaeg-CS  66  36 64.70588
3         2 Bra (WT)  71  22 76.34409
4         3   Aaeg-M  57  31 64.77273
5         3  Aaeg-CS  75  19 79.78723
6         3 Bra (WT)  64  35 64.64646
```

```
flight.aeg =
  rbind(
    data.frame(replicate = flight_aegypti[1,1],
```

```
                treatment = flight_aegypti[1,2],
                result = rep(1, flight_aegypti[1,3])),
    data.frame(replicate = flight_aegypti[1,1],
                treatment = flight_aegypti[1,2],
                result = rep(0, flight_aegypti[1,4])))

for(i in 2:nrow(flight_aegypti)){
  flight.aeg =
    rbind(
      flight.aeg,
      data.frame(replicate = flight_aegypti[i,1],
                  treatment = flight_aegypti[i,2],
                  result = rep(1, flight_aegypti[i,3])),
      data.frame(replicate = flight_aegypti[i,1],
                  treatment = flight_aegypti[i,2],
                  result = rep(0, flight_aegypti[i,4]))
    )
}

flight.aeg$Species = 'Ae. aegypti'
flight.aeg$treatment = factor(flight.aeg$treatment, levels = c("Bra (WT)", "Aaeg-M", "Aaeg-CS"))

head(flight.aeg)
```

```
  replicate treatment result     Species
1         2    Aaeg-M      1 Ae. aegypti
2         2    Aaeg-M      1 Ae. aegypti
3         2    Aaeg-M      1 Ae. aegypti
4         2    Aaeg-M      1 Ae. aegypti
5         2    Aaeg-M      1 Ae. aegypti
6         2    Aaeg-M      1 Ae. aegypti
```

```
flight.aeg.summary = flight.aeg %>% group_by(treatment) %>%
  dplyr::summarise(value = n()) %>% data.frame() %>%
  mutate(pos = 1:3)

head(flight.aeg.summary)
```

```
  treatment value pos
1  Bra (WT)   289   1
2    Aaeg-M   286   2
3   Aaeg-CS   302   3
```

###### Model .

```
flight.aeg.glmer = glmer(formula = 'result ~ treatment + (1|replicate)',
                         data = flight.aeg, family = binomial)

summary(flight.aeg.glmer)
```

```
Generalized linear mixed model fit by maximum likelihood (Laplace
```

```
  Approximation) [glmerMod]
 Family: binomial  ( logit )
Formula: result ~ treatment + (1 | replicate)
   Data: flight.aeg

     AIC      BIC   logLik deviance df.resid
  1045.4   1064.5   -518.7   1037.4      873

Scaled residuals:
    Min      1Q  Median      3Q     Max
-1.8340 -1.4218  0.6088  0.6315  0.7033

Random effects:
 Groups    Name        Variance Std.Dev.
 replicate (Intercept) 0.02668  0.1633
Number of obs: 877, groups:  replicate, 3

Fixed effects:
                 Estimate Std. Error z value Pr(>|z|)
(Intercept)       1.03584    0.16391   6.320 2.62e-10 ***
treatmentAaeg-M  -0.22027    0.18562  -1.187    0.235
treatmentAaeg-CS -0.04688    0.18647  -0.251    0.801
---
Signif. codes:  0 '***' 0.001 '**' 0.01 '*' 0.05 '.' 0.1 ' ' 1

Correlation of Fixed Effects:
            (Intr) trtA-M
tretmntAg-M -0.590
trtmntAg-CS -0.587  0.519
```

```
pairewise.f.aeg = glht(flight.aeg.glmer,
                       mcp(treatment="Tukey"))

summary(pairewise.f.aeg)
```

```
     Simultaneous Tests for General Linear Hypotheses

Multiple Comparisons of Means: Tukey Contrasts

Fit: glmer(formula = result ~ treatment + (1 | replicate), data = flight.aeg,
    family = binomial)

Linear Hypotheses:
                        Estimate Std. Error z value Pr(>|z|)
Aaeg-M - Bra (WT) == 0  -0.22027    0.18562  -1.187    0.461
Aaeg-CS - Bra (WT) == 0 -0.04688    0.18647  -0.251    0.966
Aaeg-CS - Aaeg-M == 0    0.17340    0.18254   0.950    0.609
(Adjusted p values reported -- single-step method)
```

Replication

```
# A tibble: 3 x 3
```

```
  Line          N     n
  <fct>     <int> <int>
1 Bra (WT)      3   289
2 Aaeg-M        3   286
3 Aaeg-CS       3   302
```

Model fit quality

```
check_model(flight.aeg.glmer)
```

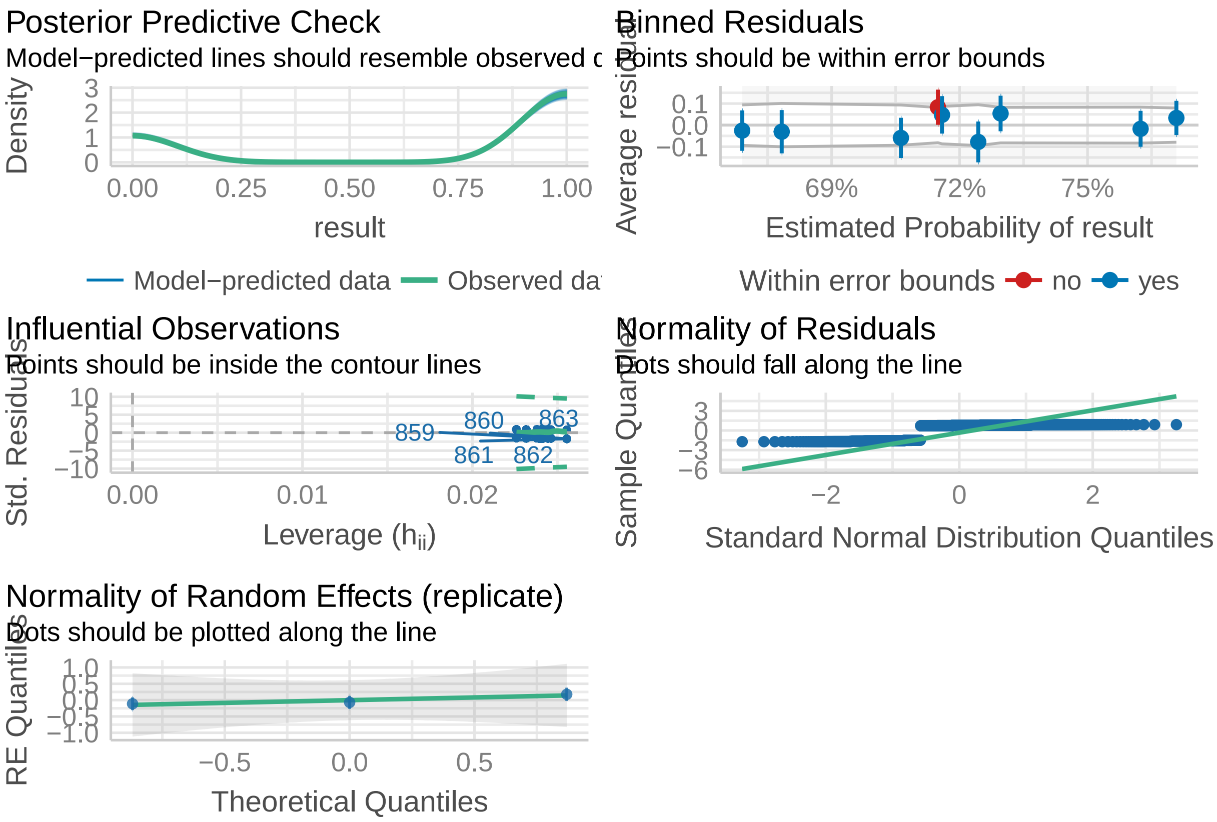

**Plot** .

```
ft.aeg = data.frame(ggpredict(model = flight.aeg.glmer,
                              terms = c("treatment")))
ft.aeg$x = factor(ft.aeg$x, levels = c("Bra (WT)", "Aaeg-M", "Aaeg-CS"))
ft.aeg$Species = 'Ae. aegypti'

ft.plot.aeg =
  ggplot(data =  NULL) +
  geom_point(data = flight_aegypti,
             aes(x = jitter(as.numeric(factor(Line))),
                 y = esc,
                 color = Line),
             size = 2, alpha = .6)+
```

```
  geom_point(data = ft.aeg,
             aes(x = as.numeric(x),
                 y = predicted * 100),
             size = 1.5, color = 'black') +
  geom_errorbar(data = ft.aeg,
                aes(x = as.numeric(x),
                    ymin = conf.low* 100,
                    ymax = conf.high* 100),
                width = 0.05, color = 'black') +
  geom_point(data = flight.aeg,
             aes(x = jitter(as.numeric(factor(treatment)), factor = 2),
                 y = jitter((result * 95), factor = .25),
                 color = treatment),
             #alpha = .25,
             size = .1) +
  annotate(geom = 'text', x= flight.aeg.summary$pos, y=-20,
           label = paste0('n = ', flight.aeg.summary$value),
           size = 3) +
  annotate(geom = 'text', x= flight.aeg.summary$pos, y=-10,
           label = 'N = 3',
           size = 3) +
  annotate(geom = 'line', x= 1:3, y=113) +
  annotate(geom = 'text', x= 2, y=118,
           label = 'n.s.',
           size = 3) +
  annotate(geom = 'line', x= 1:2, y=105) +
  annotate(geom = 'text', x= 1.5, y=110,
           label = 'n.s.',
           size = 3) +
  scale_x_continuous(breaks = 1:3,
                     labels = c("Bra (WT)",
                                "Aaeg-M",
                                "Aaeg-CS"),
                     limits = c(.5,3.5)) +
  scale_y_continuous(sec.axis = sec_axis(~., breaks = c(0,95),
                                         labels = c('In', 'Out'),
                                         name = 'Escape status (binomial)'),
                     limits = c(-20,120),
                     breaks = c(0,25,50,75,100)) +
  scale_color_manual(values = line.color.aeg, labels = names(line.color.aeg)) +
  labs(x = 'Line',
       y = 'Escaped males (%)') +
  theme_classic() +
  theme(legend.position = 'none',
        axis.title.y.right = element_text(color = 'black'),
        axis.ticks.y.right = element_line(color = 'black'),
        axis.line.y.right = element_line(color = 'black'))

ft.plot.aeg
```

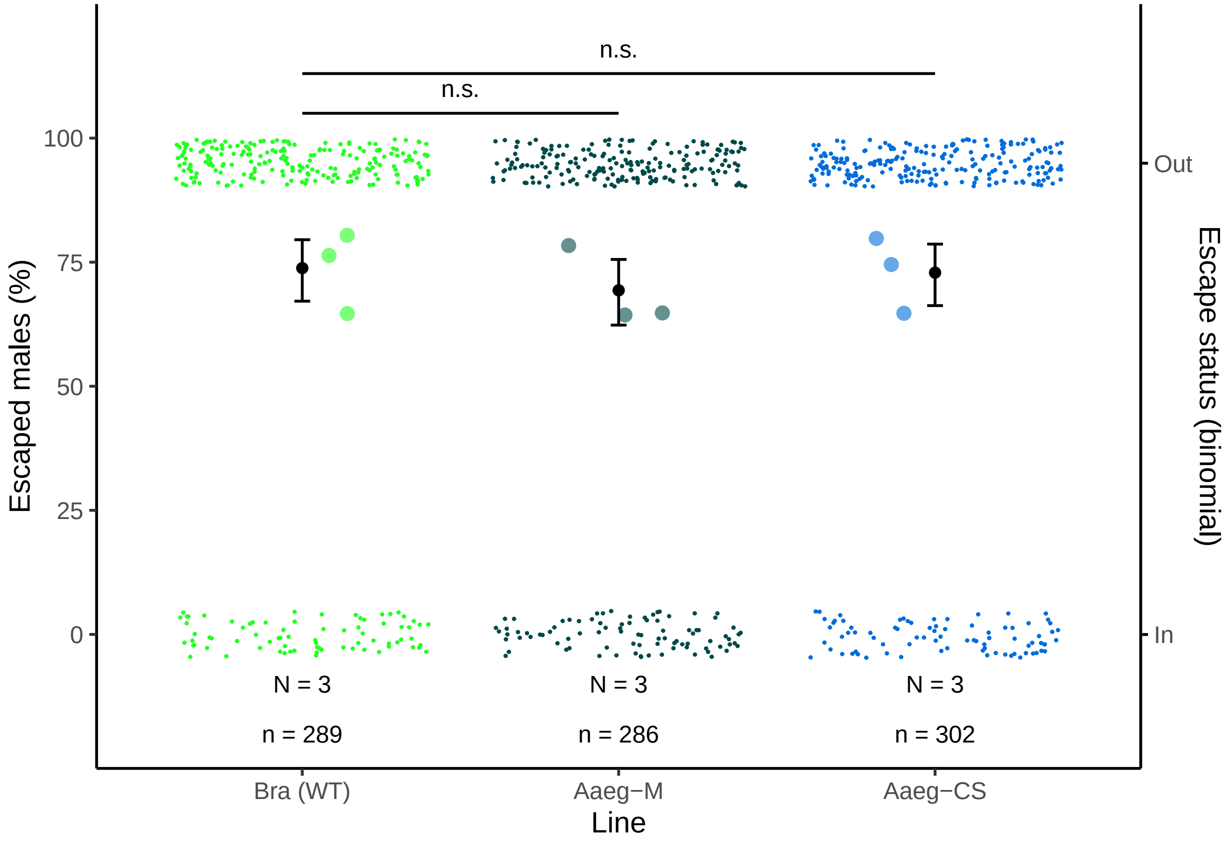

##### Albopictus

###### Data .

```
flight_albo<-read.table("data/Flight_albo.csv", header=TRUE, sep=";")
flight_albo$esc = flight_albo$out / (flight_albo$out + flight_albo$in.) * 100
flight_albo$Line = factor(flight_albo$Line,
                          levels = c("BiA (WT)", "Aal-M", "Aal-CS"))

head(flight_albo)
```

```
  Replicate     Line out in.      esc
1         2    Aal-M  26  63 29.21348
2         2   Aal-CS  45  48 48.38710
3         2 BiA (WT)  38  48 44.18605
4         3    Aal-M  57  31 64.77273
5         3   Aal-CS  62  16 79.48718
6         3 BiA (WT)  58  31 65.16854
```

```
flight.albo =
  rbind(
    data.frame(replicate = flight_albo[1,1],
               treatment = flight_albo[1,2],
               result = rep(1, flight_albo[1,3])),
```

```
    data.frame(replicate = flight_albo[1,1],
               treatment = flight_albo[1,2],
               result = rep(0, flight_albo[1,4])))

for(i in 2:nrow(flight_albo)){
  flight.albo =
    rbind(
      flight.albo,
      data.frame(replicate = flight_albo[i,1],
                 treatment = flight_albo[i,2],
                 result = rep(1, flight_albo[i,3])),
      data.frame(replicate = flight_albo[i,1],
                 treatment = flight_albo[i,2],
                 result = rep(0, flight_albo[i,4]))
    )
}

flight.albo$Species = 'Ae. albopictus'
flight.albo$treatment = factor(flight.albo$treatment,
                               levels = c("BiA (WT)", "Aal-M", "Aal-CS"))

head(flight.albo)
```

```
  replicate treatment result        Species
1         2     Aal-M      1 Ae. albopictus
2         2     Aal-M      1 Ae. albopictus
3         2     Aal-M      1 Ae. albopictus
4         2     Aal-M      1 Ae. albopictus
5         2     Aal-M      1 Ae. albopictus
6         2     Aal-M      1 Ae. albopictus
```

```
flight.alb.summary = flight.albo %>% group_by(treatment) %>%
  dplyr::summarise(value = n()) %>% data.frame() %>%
  mutate(pos = 1:3)

head(flight.alb.summary)
```

```
  treatment value pos
1  BiA (WT)   264   1
2     Aal-M   273   2
3    Aal-CS   244   3
```

##### Model .

```
mod.glmer.albo = glmer(formula = 'result ~ treatment + (1|replicate)',
                       data = flight.albo, family = binomial)

summary(mod.glmer.albo)
```

```
Generalized linear mixed model fit by maximum likelihood (Laplace
  Approximation) [glmerMod]
```

```
 Family: binomial  ( logit )
Formula: result ~ treatment + (1 | replicate)
   Data: flight.albo

     AIC      BIC   logLik deviance df.resid
  1036.3   1054.9   -514.1   1028.3      777

Scaled residuals:
    Min      1Q  Median      3Q     Max
-1.7839 -0.9200  0.5606  0.9063  1.4280

Random effects:
 Groups    Name        Variance Std.Dev.
 replicate (Intercept) 0.2407   0.4906
Number of obs: 781, groups:  replicate, 3

Fixed effects:
                Estimate Std. Error z value Pr(>|z|)
(Intercept)       0.2196     0.3106   0.707   0.4795
treatmentAal-M   -0.3636     0.1784  -2.038   0.0415 *
treatmentAal-CS   0.3490     0.1867   1.869   0.0616 .
---
Signif. codes:  0 '***' 0.001 '**' 0.01 '*' 0.05 '.' 0.1 ' ' 1

Correlation of Fixed Effects:
            (Intr) trtA-M
tretmntAl-M -0.293
trtmntAl-CS -0.279  0.485
```

```
pairwise.f.albo = glht(mod.glmer.albo,
                       mcp(treatment="Tukey"))

summary(pairwise.f.albo)
```

```
     Simultaneous Tests for General Linear Hypotheses

Multiple Comparisons of Means: Tukey Contrasts

Fit: glmer(formula = result ~ treatment + (1 | replicate), data = flight.albo,
    family = binomial)

Linear Hypotheses:
                       Estimate Std. Error z value Pr(>|z|)
Aal-M - BiA (WT) == 0   -0.3636     0.1784  -2.038    0.103
Aal-CS - BiA (WT) == 0   0.3490     0.1867   1.869    0.148
Aal-CS - Aal-M == 0      0.7126     0.1854   3.842   <0.001 ***
---
Signif. codes:  0 '***' 0.001 '**' 0.01 '*' 0.05 '.' 0.1 ' ' 1
(Adjusted p values reported -- single-step method)
```

Replication

```
# A tibble: 3 x 3
```

```
  Line          N     n
  <fct>     <int> <int>
1 BiA (WT)      3   264
2 Aal-M         3   273
3 Aal-CS        3   244
```

Model fit quality

```
check_model(mod.glmer.albo)
```

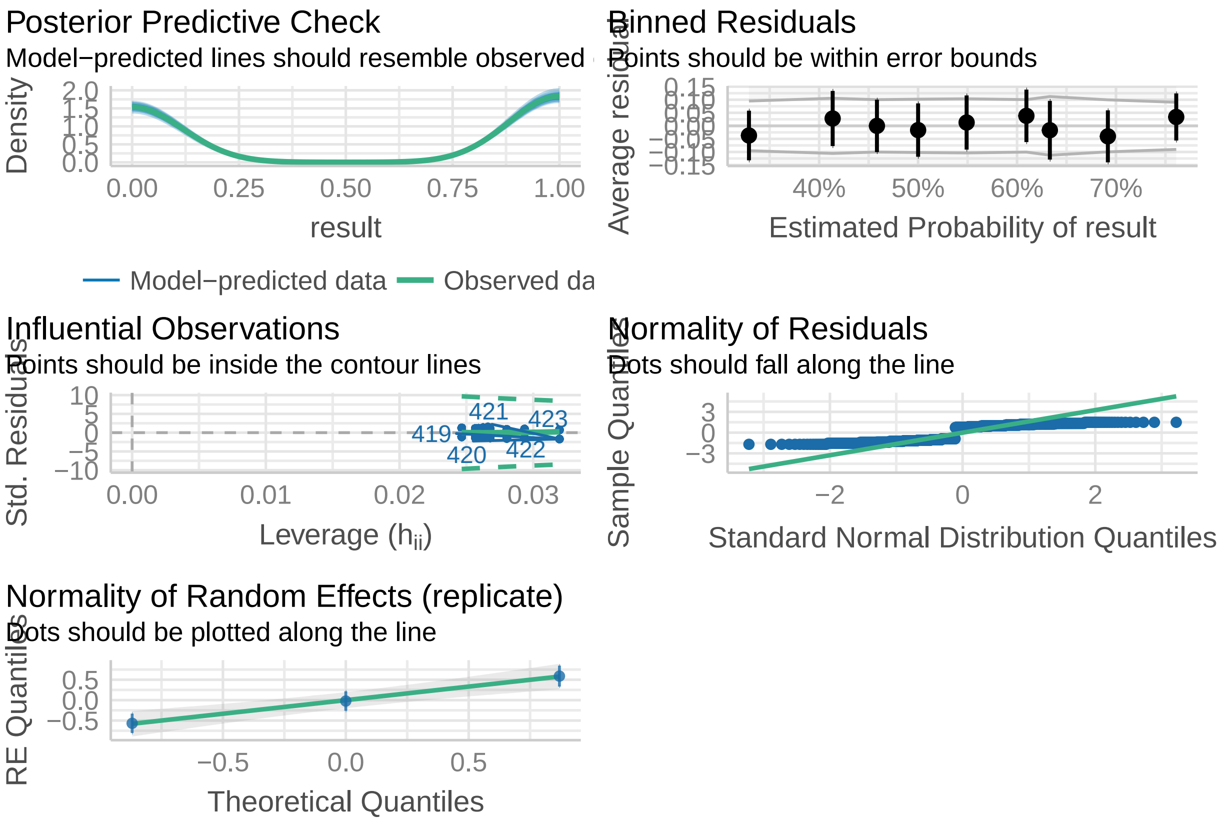

**Plot** .

```
ft.albo = data.frame(ggpredict(model = mod.glmer.albo, terms = c("treatment")))
ft.albo$Specft.albo$Species = 'Ae. albopictus'

ft.plot.alb =
  ggplot(data =  NULL) +
  geom_point(data = flight_albo,
             aes(x = jitter(as.numeric(factor(Line))),
                 y = esc, color = Line),
             size = 2, alpha = .6) +
  geom_point(data = ft.albo,
             aes(x = as.numeric(x),
                 y = predicted * 100),
```

```r
             size = 1.5, color = 'black') +
  geom_errorbar(data = ft.albo,
                aes(x = as.numeric(x),
                    ymin = conf.low* 100,
                    ymax = conf.high* 100),
                width = 0.05, color = 'black') +
  geom_point(data = flight.albo,
             aes(x = jitter(as.numeric(factor(treatment)), factor = 2),
                 y = jitter((result * 95), factor = .25),
                 color = treatment),
             #alpha = .25,
             size = .1) +
  annotate(geom = 'text', x= flight.alb.summary$pos, y=-20,
           label = paste0('n = ', flight.alb.summary$value),
           size = 3) +
  annotate(geom = 'text', x= flight.alb.summary$pos, y=-10,
           label = 'N = 3',
           size = 3) +
  annotate(geom = 'line', x= 1:3, y=113) +
  annotate(geom = 'text', x= 2, y=118,
           label = 'n.s.',
           size = 3) +
  annotate(geom = 'line', x= 1:2, y=105) +
  annotate(geom = 'text', x= 1.5, y=110,
           label = 'n.s.',
           size = 3) +
  scale_x_continuous(breaks = 1:3,
                     labels = c("BiA (WT)",
                                "Aal-M",
                                "Aal-CS"),
                     limits = c(.5,3.5)) +
  scale_y_continuous(sec.axis = sec_axis(~., breaks = c(0,95),
                                         labels = c('In', 'Out'),
                                         name = 'Escape status (binomial)'),
                     limits = c(-20,120),
                     breaks = c(0,25,50,75,100)) +
  scale_color_manual(values = line.color.alb,
                     labels = names(line.color.alb)) +
  labs(x = 'Line',
       y = 'Escaped males (%)') +
  theme_classic() +
  theme(legend.position = 'none',
        axis.title.y.right = element_text(color = 'black'),
        axis.ticks.y.right = element_line(color = 'black'),
        axis.line.y.right = element_line(color = 'black'),
        panel.background = element_rect(fill = 'grey'))

ft.plot.alb
```

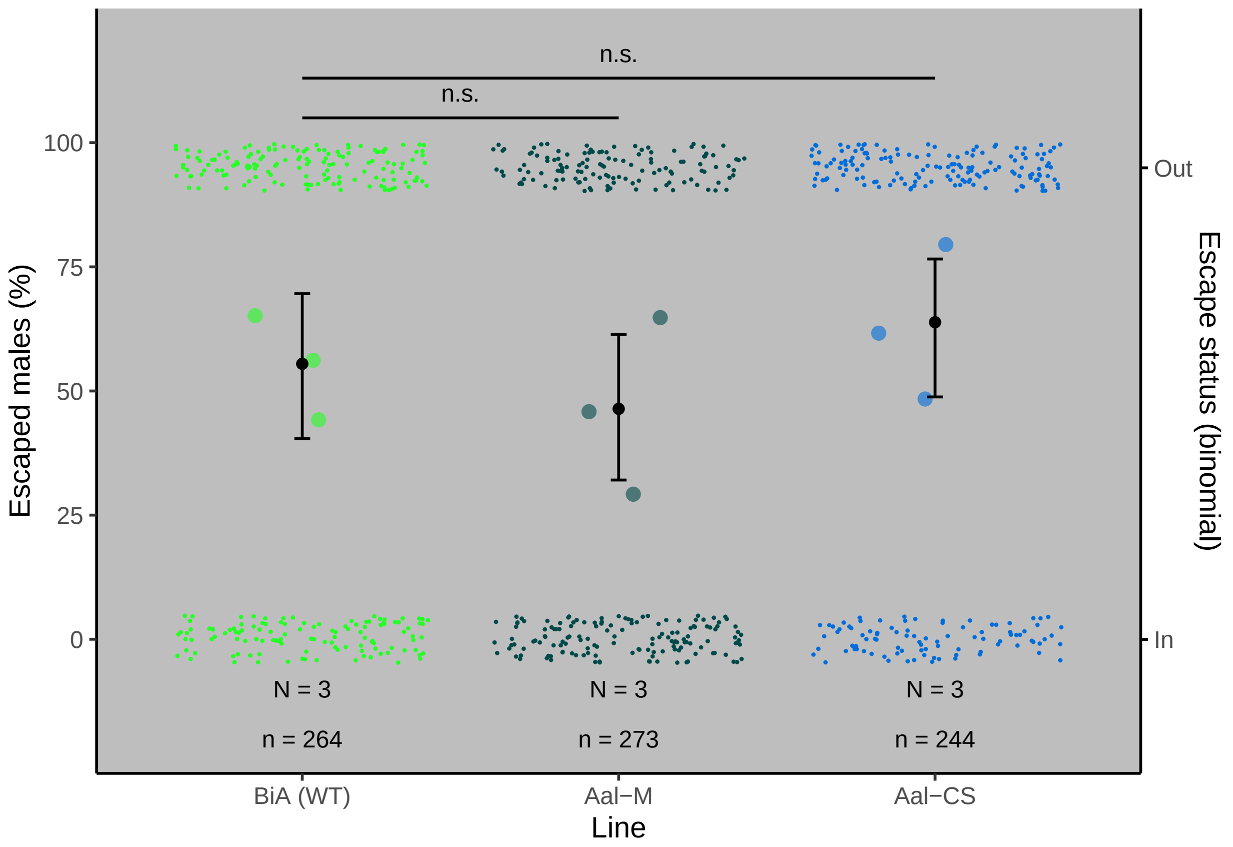

#### Adult survival

##### Aegypti

###### Data .

```
data_aeg <- read.table("data/Survival_aeg.txt", h=T)
data_aeg$Line[data_aeg$Line=='Bra-(WT)'] = 'Bra (WT)'
data_aeg$Line = factor(data_aeg$Line,
                       levels = c('Bra (WT)', 'Aaeg-M','Aaeg-CS'))
head(data_aeg)
```

```
  Replicate     Line larvae adults larval_deaths male_adults D1 D2 D3 D4 D5 D6
1         1   Aaeg-M    100     87            13          87  0  0  0  0  0  0
2         2   Aaeg-M    100     82            18          82  0  0  0  0  0  0
3         3   Aaeg-M    100     93             7          93  1  0  0  0  1  0
4         4   Aaeg-M    100     73            27          73  0  0  0  0  0  0
5         1 Bra (WT)    200    185            15          82  0  0  0  0  0  0
6         2 Bra (WT)    200    197             3          97  0  0  0  0  2  0
  D7 D8 D9 D10 D11 D12 D13 D14
1  0  0  0   0   0   0   0   0
2  0  0  0   0   0   0   0   0
3  0  0  0   0   0   0   0   0
4  0  1  0   0   0   0   0   1
```

```
5  0  0  0   0   0   0   1   0
6  0  0  0   0   0   0   0   0
```

```
# First, melt the data.frame
data1 <- melt(data_aeg , .(Replicate,Line, larvae, adults, larval_deaths, male_adults))
# Transform days in numbers
data1$variable <- as.numeric(str_replace(data1$variable, "D",""))

#Sort by Line, Replicate and variable.
data2 <- arrange(data1, Line, Replicate, variable )
event <- subset(data2, value>0)

dat <- NULL
temporary_event <- NULL
for (i in 1:nrow(event)) {
  N <- event[i, "value"]
  dat <- data.frame(Replicate=rep(event[i,"Replicate"],N) ,Line=rep(event[i,"Line"],N) ,larvae=rep(event

  temporary_event  <- rbind(temporary_event ,dat )
}

dat <- data.frame(Replicate=1, Line="Aaeg-M", larvae=100,  adults=87, larval_deaths=13, male_adults=87,
dat1 <- data.frame(Replicate=2, Line="Aaeg-M", larvae=100,  adults=82, larval_deaths=18, male_adults=82

temporary_event <- rbind(temporary_event, dat, dat1)

# Add the censored values.
censored <- unique(ddply(temporary_event,
                         .(Replicate,Line,larvae,adults,
                           larval_deaths,male_adults),
                         summarise,
                         variable=Day,
                         Event=sum(Event),
                         Censored=(male_adults-Event))[,-7])
censored$variable <- max(data2$variable)

dat <- NULL
temporary <- NULL
for (i in 1:nrow(censored)) {
  N <- censored[i, "Censored"]
  dat <- data.frame(Replicate=rep(censored[i,"Replicate"],N) ,Line=rep(censored[i,"Line"],N) ,larvae=rep
                    Day=rep(censored[i,"variable"],N),
                    Event=rep(0,N) )

  temporary <- rbind(temporary,dat )
}

data_surv <- rbind(temporary_event, temporary)
data_surv <- arrange(data_surv, Line,Replicate,desc(Event),Day)

SurvObj <- with(data_surv, Surv(Day, Event))
```

**Model** .

```r
my.KMest <- survfit(SurvObj ~ Line , conf.int=0.95, data = data_surv)
summary(my.KMest)
```

```
Call: survfit(formula = SurvObj ~ Line, data = data_surv, conf.int = 0.95)

                Line=Bra (WT)
 time n.risk n.event survival std.err lower 95% CI upper 95% CI
    1    791       5    0.994 0.00282        0.988        0.999
    2    786       2    0.991 0.00333        0.985        0.998
    5    784       4    0.986 0.00416        0.978        0.994
    6    780       1    0.985 0.00435        0.976        0.993
    7    779       2    0.982 0.00469        0.973        0.992
   10    777       2    0.980 0.00501        0.970        0.990
   12    775       1    0.979 0.00516        0.968        0.989
   13    774       2    0.976 0.00544        0.965        0.987

                Line=Aaeg-M
 time n.risk n.event survival std.err lower 95% CI upper 95% CI
    1    757       5    0.993 0.00294        0.988        0.999
    2    752       1    0.992 0.00322        0.986        0.998
    5    751       2    0.989 0.00372        0.982        0.997
    7    749       2    0.987 0.00415        0.979        0.995
    8    747       3    0.983 0.00472        0.974        0.992
   11    744       1    0.982 0.00490        0.972        0.991
   14    743       1    0.980 0.00507        0.970        0.990

                Line=Aaeg-CS
 time n.risk n.event survival std.err lower 95% CI upper 95% CI
    2    420       2    0.995 0.00336        0.989        1.000
    5    418       1    0.993 0.00411        0.985        1.000
   11    417       1    0.990 0.00474        0.981        1.000
   14    416       1    0.988 0.00529        0.978        0.999
```

```r
df = data.frame(t = my.KMest$time,
                nb.rk = my.KMest$n.risk,
                line = c(rep(names(my.KMest$strata[1]),
                             my.KMest$strata[1]),
                         rep(names(my.KMest$strata[2]),
                             my.KMest$strata[2]),
                         rep(names(my.KMest$strata[3]),
                             my.KMest$strata[3]))
)

# df$t[df$line == 'Line=MyriaF' & df$t==4] = 7
df$t[df$line == 'Line=Aaeg-CS' & df$t==5] = 7
# df$t[df$line == 'Line=MyriaF' & df$t==3] = 1
df$t[df$line == 'Line=Aaeg-CS' & df$t==2] = 1

# Test differences of survival at 7 and 14 days
data_aeg.sum =
  data_aeg %>%
  group_by(Line, Replicate) %>%
  dplyr::summarise(sur.7 = (male_adults -
```

```
                              (D1 + D2 + D3 + D4 + D5 + D6 + D7))/
                     male_adults,
            sur.14 = (male_adults - (D1 + D2 + D3 + D4 + D5 +
                                        D6 + D7 + D8 + D9 + D10 +
                                        D11 + D14 + D13 + D14))/
              male_adults)

mod.surv.7 = lm(data = data_aeg.sum,
   formula = sur.7 ~ Line)

summary(mod.surv.7)
```

```
Call:
lm(formula = sur.7 ~ Line, data = data_aeg.sum)

Residuals:
      Min        1Q    Median        3Q       Max 
-0.030523 -0.004965  0.002688  0.008733  0.017097 

Coefficients:
            Estimate Std. Error t value Pr(>|t|)    
(Intercept) 0.982904   0.004902 200.529   <2e-16 ***
LineAaeg-M  0.004884   0.006932   0.705    0.491    
LineAaeg-CS 0.009954   0.008490   1.172    0.257    
---
Signif. codes:  0 '***' 0.001 '**' 0.01 '*' 0.05 '.' 0.1 ' ' 1

Residual standard error: 0.01386 on 17 degrees of freedom
Multiple R-squared:  0.07789,	Adjusted R-squared:  -0.0306 
F-statistic: 0.7179 on 2 and 17 DF,  p-value: 0.502
```

```
TukeyHSD(aov(mod.surv.7))
```

```
  Tukey multiple comparisons of means
    95% family-wise confidence level

Fit: aov(formula = mod.surv.7)

$Line
                        diff         lwr        upr     p adj
Aaeg-M-Bra (WT)  0.004884479 -0.01289814 0.02266710 0.7640413
Aaeg-CS-Bra (WT) 0.009953603 -0.01182557 0.03173277 0.4848501
Aaeg-CS-Aaeg-M   0.005069124 -0.01671005 0.02684830 0.8235245
```

```
mod.surv.14 = lm(data = data_aeg.sum,
   formula = sur.14 ~ Line)

summary(mod.surv.14)
```

```
Call:
lm(formula = sur.14 ~ Line, data = data_aeg.sum)
```

```
Residuals:
      Min        1Q    Median        3Q       Max 
-0.025193 -0.009310  0.002312  0.010522  0.020920 

Coefficients:
            Estimate Std. Error t value Pr(>|t|)    
(Intercept) 0.977574   0.005019 194.776   <2e-16 ***
LineAaeg-M  0.001505   0.007098   0.212    0.835    
LineAaeg-CS 0.008140   0.008693   0.936    0.362    
---
Signif. codes:  0 '***' 0.001 '**' 0.01 '*' 0.05 '.' 0.1 ' ' 1

Residual standard error: 0.0142 on 17 degrees of freedom
Multiple R-squared:  0.05089,	Adjusted R-squared:  -0.06077 
F-statistic: 0.4558 on 2 and 17 DF,  p-value: 0.6415
```

```
TukeyHSD(aov(mod.surv.14))
```

```
  Tukey multiple comparisons of means
    95% family-wise confidence level

Fit: aov(formula = mod.surv.14)

$Line
                        diff         lwr        upr     p adj
Aaeg-M-Bra (WT)  0.001505310 -0.01670328 0.01971390 0.9755453
Aaeg-CS-Bra (WT) 0.008139992 -0.01416088 0.03044087 0.6254566
Aaeg-CS-Aaeg-M   0.006634682 -0.01566619 0.02893556 0.7299193
```

Replication

```
# A tibble: 3 x 3
  Line          N     n
  <fct>     <int> <int>
1 Bra (WT)      8   791
2 Aaeg-M        8   755
3 Aaeg-CS       4   420
```

Model quality

```
check_model(mod.surv.7)
```

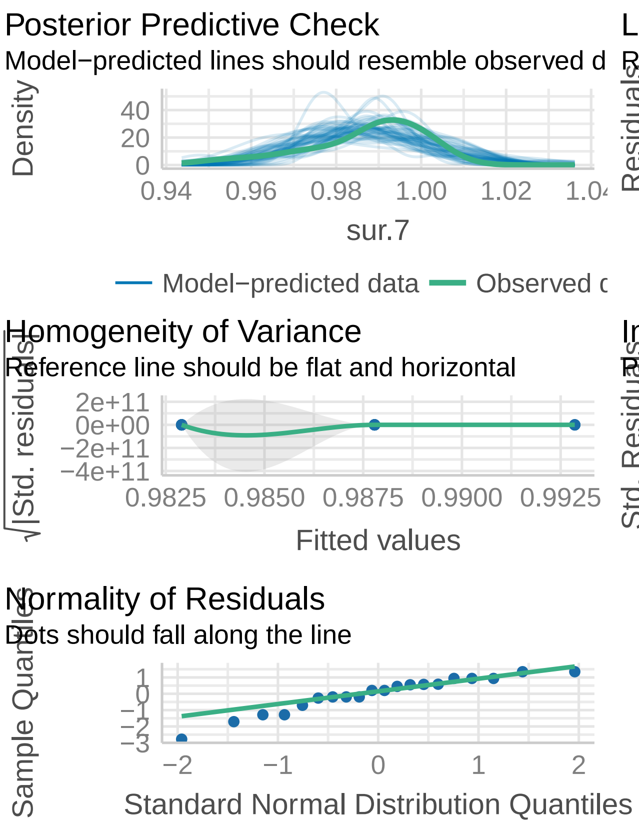

```
check_model(mod.surv.14)
```

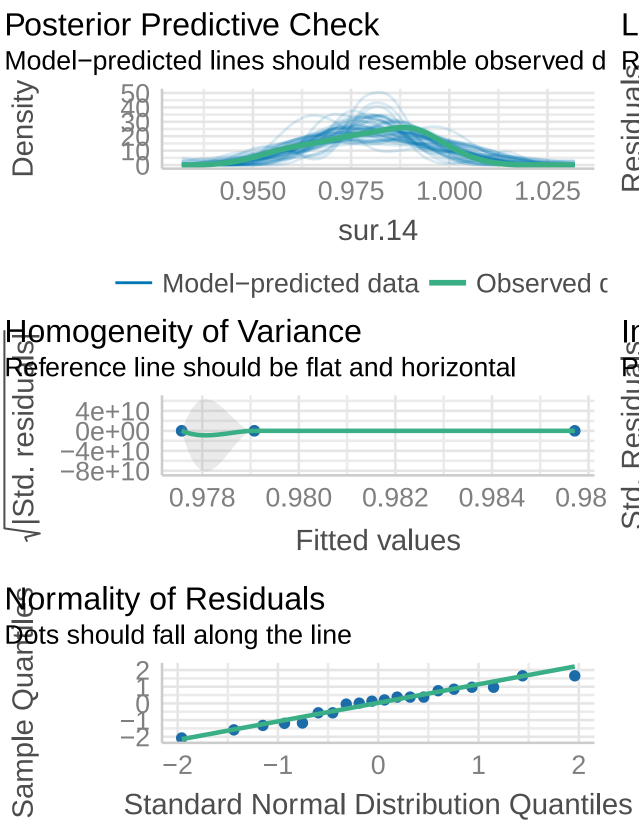

**Plot** .

```
ggsurv <-
  ggsurvplot(my.KMest,
             data = data_surv,
             conf.int = T,
             risk.table = T,
             pval = T,
             risk.table.height = 0.25)

line.color.aeg = c("#24ff24","#004949", "#006ddb")
names(line.color.aeg) = c('Line=Bra (WT)', 'Line=Aaeg-M', 'Line=Aaeg-CS')

p.aeg =
  ggsurv$plot +
  labs(x = 'Time (days)') +
  scale_y_continuous(limits =  c(.84, 1),
                     breaks = c(.84, .90, .95, 1),
                     labels = c(0, .90, .95,1)) +
  scale_x_continuous(breaks = c(0,7,14)) +
  scale_color_manual(breaks = names(line.color.aeg),
                     values = line.color.aeg,
                     guide = 'none') +
  scale_fill_manual(breaks = names(line.color.aeg),
                    values = line.color.aeg,
```

```
                    labels = c("Bra (WT)", "Aaeg-M", "Aaeg-CS"),
                    name = "Line") +
  theme(axis.line.y = element_blank(),
        legend.position = 'right',
        panel.background = element_rect(fill = 'white')) +
  annotate(geom = 'segment', x = -Inf, xend = -Inf,
           y = -Inf, yend = Inf) +
  annotate(geom = 'segment', x = -Inf, xend = -Inf,
           y = .85, yend = .89, linetype = 'dashed', color = 'white') +
  annotate(geom = 'text',
           x = -.5, y = .895,
           label = "Number at risk",
           hjust = 0, size = 3) +
  annotate(geom = 'text',
           x = rep(0, 3), y =  seq(.841, .881, .02),
           label = rev(df$nb.rk[df$t == 1]),
           color = rev(c("#24ff24","#004949", "#006ddb")),
           size = 3) +
  annotate(geom = 'text',
           x = rep(7, 3), y =  seq(.841, .881, .02),
           label = rev(df$nb.rk[df$t == 7]),
           color = rev(c("#24ff24","#004949", "#006ddb")),
           size = 3) +
  annotate(geom = 'text',
           x = rep(14, 3), y =  seq(.841, .881, .02),
           label = rev(df$nb.rk[df$t == 14]),
           color = rev(c("#24ff24","#004949", "#006ddb")),
           size = 3) +
  annotate(geom = 'text', x = 14.8, y = .875, label = 'n.s', size = 3) +
  annotate(geom = "line", x = rep(14.4,2), y = c(.865,.885)) +
  annotate(geom = 'text', x = 15.5, y = .855, label = 'n.s', size = 3) +
  annotate(geom = "line", x = rep(15.15,2), y = c(.845,.885))

p.aeg
```

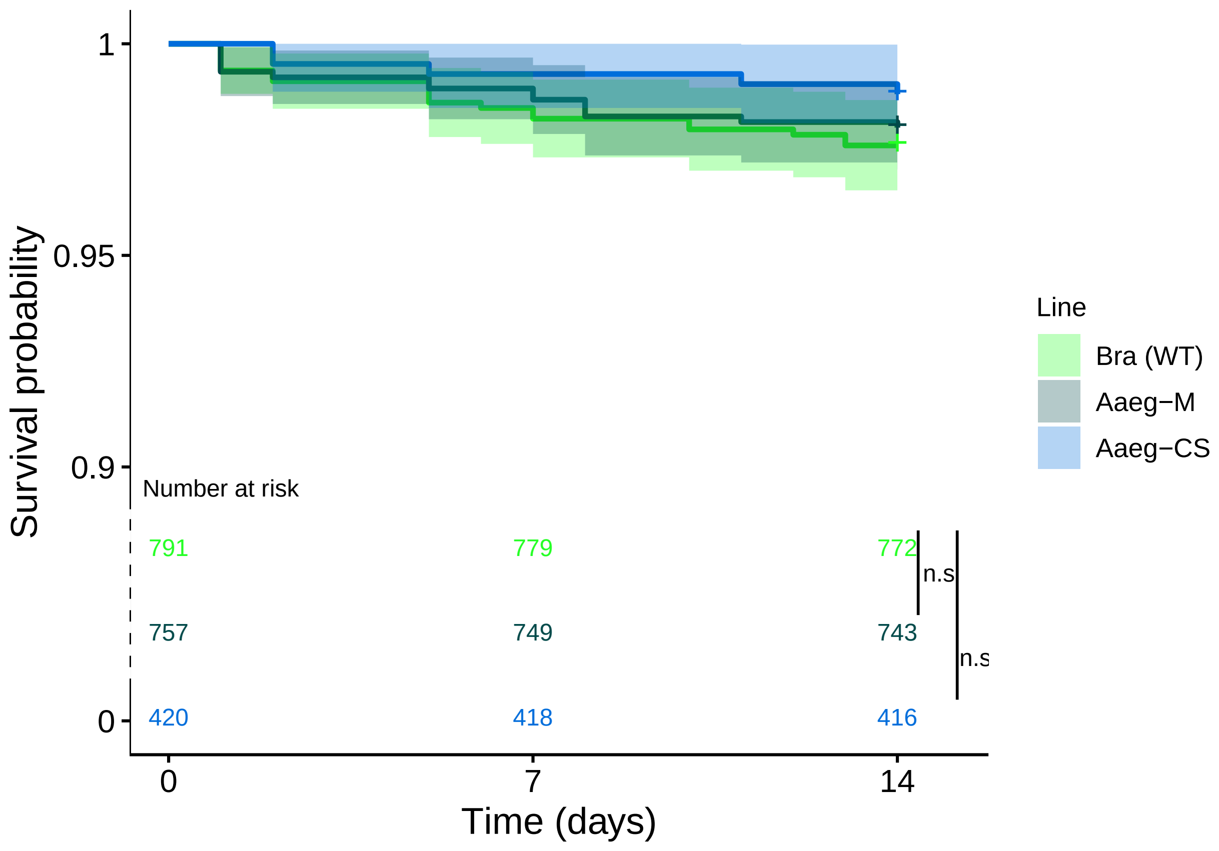

**Albopictus**

**Data** .

```
data_albo <- read.table("data/Survival_albo.txt", h=T)
data_albo$Line[data_albo$Line=='BiA-(WT)'] = 'BiA (WT)'
data1 <- melt(data_albo, .(Replicate,Line, larvae, adults, larval_deaths, male_adults))
#Then transform days in numbers
data1$variable <- as.numeric(str_replace(data1$variable, "D",""))
#Sort by Line, Replicate and variable.
data2 <- arrange(data1, Line, Replicate, variable )
#We now separate the data set in two parts: individuals that has experienced an event(death) and censor

event <- subset(data2, value>0)

#We will now work of the dataset with individuals experiencing an event.

dat <- NULL
temporary_event <- NULL
for (i in 1:nrow(event)) {
  N <- event[i, "value"]
  dat <- data.frame(Replicate=rep(event[i,"Replicate"],N) ,Line=rep(event[i,"Line"],N) ,larvae=rep(even

  temporary_event  <- rbind(temporary_event ,dat )
}
```

```r
temporary_event
```

```
   Replicate     Line larvae adults larval_deaths male_adults Day Event
1          1   Aal-CS    100     71            29          71   5     1
2          1   Aal-CS    100     71            29          71   7     1
3          1   Aal-CS    100     71            29          71  13     1
4          1   Aal-CS    100     71            29          71  13     1
5          2   Aal-CS    100     73            27          73   4     1
6          2   Aal-CS    100     73            27          73   4     1
7          2   Aal-CS    100     73            27          73  11     1
8          2   Aal-CS    100     73            27          73  13     1
9          3   Aal-CS    100     61            39          61  13     1
10         4   Aal-CS    100     69            31          69   8     1
11         1    Aal-M    100     88            12          88   1     1
12         1    Aal-M    100     88            12          88   2     1
13         1    Aal-M    100     88            12          88   4     1
14         1    Aal-M    100     88            12          88   4     1
15         1    Aal-M    100     88            12          88  12     1
16         2    Aal-M    100     79            21          79   1     1
17         2    Aal-M    100     79            21          79   2     1
18         2    Aal-M    100     79            21          79   6     1
19         2    Aal-M    100     79            21          79  10     1
20         3    Aal-M    100     80            20          80   1     1
21         3    Aal-M    100     80            20          80   8     1
22         3    Aal-M    100     80            20          80  12     1
23         4    Aal-M    100     77            23          73   1     1
24         1 BiA (WT)    200    103            97          56   1     1
25         1 BiA (WT)    200    103            97          56   1     1
26         1 BiA (WT)    200    103            97          56   2     1
27         1 BiA (WT)    200    103            97          56   4     1
28         2 BiA (WT)    200    141            59          96   1     1
29         2 BiA (WT)    200    141            59          96   2     1
30         2 BiA (WT)    200    141            59          96  13     1
31         4 BiA (WT)    200    130            70          76   1     1
32         4 BiA (WT)    200    130            70          76   9     1
```

```r
# here is or transformed dataset with one
# row for each time we have an event.

# BIA rep3 has no event over the course of
# the study, we add it manually so that it would appear.
dat <- data.frame(Replicate=3, Line="BiA (WT)", larvae=200,  adults=129, larval_deaths=71, male_adults=

temporary_event <- rbind(temporary_event, dat)

#We now add the censored values.

censored <- unique(ddply(temporary_event, .(Replicate,Line,larvae,adults,larval_deaths,male_adults), su
censored$variable <- max(data2$variable)

dat <- NULL
temporary <- NULL
for (i in 1:nrow(censored)) {
```

```
  N <- censored[i, "Censored"]
  dat <- data.frame(Replicate=rep(censored[i,"Replicate"],N) ,Line=rep(censored[i,"Line"],N) ,larvae=re

  temporary <- rbind(temporary,dat )
}

#Bind the final dataset
data_surv <- rbind(temporary_event, temporary)
data_surv <- arrange(data_surv, Line,Replicate,desc(Event),Day)

SurvObj <- with(data_surv, Surv(Day, Event))
```

##### Model .

```
my.KMest <- survfit(SurvObj ~ Line , conf.int=0.95, data = data_surv)
summary(my.KMest)
```

```
Call: survfit(formula = SurvObj ~ Line, data = data_surv, conf.int = 0.95)

                Line=Aal-CS
 time n.risk n.event survival std.err lower 95% CI upper 95% CI
    4    274       2    0.993 0.00514        0.983        1.000
    5    272       1    0.989 0.00629        0.977        1.000
    7    271       1    0.985 0.00725        0.971        1.000
    8    270       1    0.982 0.00809        0.966        0.998
   11    269       1    0.978 0.00884        0.961        0.996
   13    268       4    0.964 0.01133        0.942        0.986

                Line=Aal-M
 time n.risk n.event survival std.err lower 95% CI upper 95% CI
    1    320       4    0.988 0.00621        0.975        1.000
    2    316       2    0.981 0.00758        0.967        0.996
    4    314       2    0.975 0.00873        0.958        0.992
    6    312       1    0.972 0.00924        0.954        0.990
    8    311       1    0.969 0.00973        0.950        0.988
   10    310       1    0.966 0.01018        0.946        0.986
   12    309       2    0.959 0.01104        0.938        0.981

                Line=BiA (WT)
 time n.risk n.event survival std.err lower 95% CI upper 95% CI
    1    320       4    0.988 0.00621        0.975        1.000
    2    316       2    0.981 0.00758        0.967        0.996
    4    314       1    0.978 0.00818        0.962        0.994
    9    313       1    0.975 0.00873        0.958        0.992
   13    312       1    0.972 0.00924        0.954        0.990
```

```
df = data.frame(t = my.KMest$time,
                nb.rk = my.KMest$n.risk,
                line = c(rep(names(my.KMest$strata[1]),
                             my.KMest$strata[1]),
                         rep(names(my.KMest$strata[2]),
                             my.KMest$strata[2]),
```

```r
                         rep(names(my.KMest$strata[3]),
                             my.KMest$strata[3]))
)

df$t[df$line == 'Line=Aal-M' & df$t==6] = 7
df$t[df$line == 'Line=BiA (WT)' & df$t==4] = 7
df$t[df$line == 'Line=Aal-CS' & df$t==4] = 1

df$line = factor(df$line, levels = c('Line=BiA (WT)','Line=Aal-M','Line=Aal-CS'))
df = df[order(df$line),]

data_albo.sum =
  data_albo %>%
  group_by(Line, Replicate) %>%
  dplyr::summarise(sur.7 = (male_adults -
                              (D1 + D2 + D3 + D4 + D5 + D6 + D7))/
                     male_adults,
            sur.14 = (male_adults - (D1 + D2 + D3 + D4 + D5 +
                                       D6 + D7 + D8 + D9 + D10 +
                                       D11 + D14 + D13 + D14))/
              male_adults)

data_albo.sum$Line = data_albo.sum$Line %>%
  factor(., levels = c('BiA (WT)','Aal-M', 'Aal-CS'))

mod.surv.7 = lm(data = data_albo.sum,
   formula = sur.7 ~ Line)

summary(mod.surv.7)
```

```

Call:
lm(formula = sur.7 ~ Line, data = data_albo.sum)

Residuals:
      Min        1Q    Median        3Q       Max
-0.045074 -0.013699  0.009359  0.013892  0.026355

Coefficients:
             Estimate Std. Error t value Pr(>|t|)
(Intercept)  0.973645   0.011243  86.600 1.85e-14 ***
LineAal-M   -0.001052   0.015900  -0.066    0.949
LineAal-CS   0.012463   0.015900   0.784    0.453
---
Signif. codes:  0 '***' 0.001 '**' 0.01 '*' 0.05 '.' 0.1 ' ' 1

Residual standard error: 0.02249 on 9 degrees of freedom
Multiple R-squared:  0.09038,   Adjusted R-squared:  -0.1118
F-statistic: 0.4471 on 2 and 9 DF,  p-value: 0.6529
```

```r
TukeyHSD(aov(mod.surv.7))
```

```

  Tukey multiple comparisons of means
```

```
    95% family-wise confidence level

Fit: aov(formula = mod.surv.7)

$Line
                         diff         lwr        upr     p adj
Aal-M-BiA (WT)  -0.001052015 -0.04544497 0.04334094 0.9975900
Aal-CS-BiA (WT)  0.012463381 -0.03192957 0.05685634 0.7216963
Aal-CS-Aal-M     0.013515396 -0.03087756 0.05790835 0.6831225
```

```r
mod.surv.14 = lm(data = data_albo.sum,
   formula = sur.14 ~ Line)

summary(mod.surv.14)
```

```
Call:
lm(formula = sur.14 ~ Line, data = data_albo.sum)

Residuals:
      Min        1Q    Median        3Q       Max
-0.039180 -0.017525  0.003466  0.019333  0.032249

Coefficients:
             Estimate Std. Error t value Pr(>|t|)
(Intercept)  0.967751   0.011931  81.111 3.33e-14 ***
LineAal-M   -0.001448   0.016873  -0.086    0.933
LineAal-CS  -0.003256   0.016873  -0.193    0.851
---
Signif. codes:  0 '***' 0.001 '**' 0.01 '*' 0.05 '.' 0.1 ' ' 1

Residual standard error: 0.02386 on 9 degrees of freedom
Multiple R-squared:  0.004137,  Adjusted R-squared:  -0.2172
F-statistic: 0.0187 on 2 and 9 DF,  p-value: 0.9815
```

```r
TukeyHSD(aov(mod.surv.14))
```

```
  Tukey multiple comparisons of means
    95% family-wise confidence level

Fit: aov(formula = mod.surv.14)

$Line
                         diff         lwr        upr     p adj
Aal-M-BiA (WT)  -0.001447932 -0.04855825 0.04566239 0.9959503
Aal-CS-BiA (WT) -0.003256096 -0.05036641 0.04385422 0.9797262
Aal-CS-Aal-M    -0.001808164 -0.04891848 0.04530215 0.9936934
```

Replication

```
# A tibble: 3 x 3
  Line          N     n
  <chr>     <int> <int>
```

```
1 Aal-CS        4   274
2 Aal-M         4   320
3 BiA (WT)      4   319
```

Model quality

```
check_model(mod.surv.7)
```

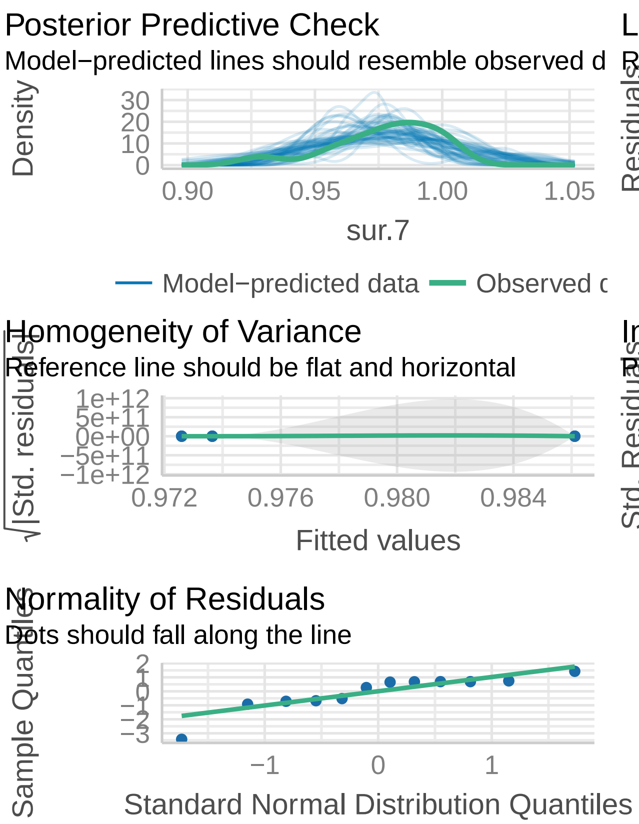

```
check_model(mod.surv.14)
```

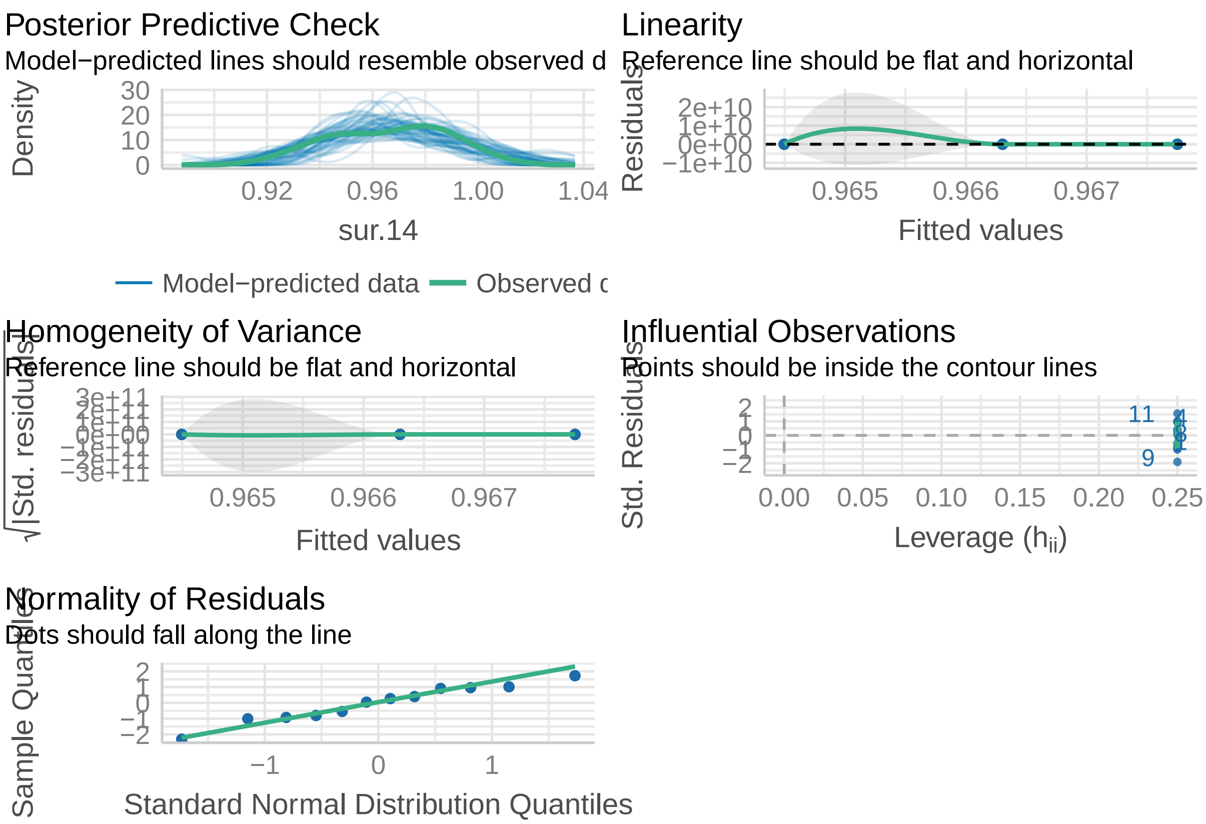

**Plot** .

```
ggsurv <-
  ggsurvplot(my.KMest,
             data = data_surv,
             conf.int = T,
             risk.table = T,
             pval = T,
             risk.table.height = 0.25)

line.color.alb = c("#24ff24","#004949", "#006ddb")
names(line.color.alb) = c('Line=BiA (WT)', 'Line=Aal-M', 'Line=Aal-CS')

p.albo =
  ggsurv$plot +
  labs(x = 'Time (days)') +
  scale_y_continuous(limits =  c(.84, 1),
                     breaks = c(.84, .90, .95, 1),
                     labels = c(0, .90, .95,1)) +
  scale_x_continuous(breaks = c(0,7,14)) +
  scale_color_manual(breaks = names(line.color.alb),
                     values = line.color.alb,
                     guide = 'none') +
  scale_fill_manual(breaks = names(line.color.alb),
                    values = line.color.alb,
```

```
                    labels = c("BiA (WT)", "Aal-M", "Aal-CS"),
                    name = "Line: ") +
  theme(axis.line.y = element_blank(),
        legend.position = 'right',
        panel.background = element_rect(fill = 'grey')) +
  annotate(geom = 'segment', x = -Inf, xend = -Inf,
           y = -Inf, yend = Inf) +
  annotate(geom = 'segment', x = -Inf, xend = -Inf,
           y = .85, yend = .89, linetype = 'dashed', color = 'white') +
  annotate(geom = 'text',
           x = -.5, y = .895,
           label = "Number at risk",
           hjust = 0, size = 3) +
  annotate(geom = 'text',
           x = rep(0, 3), y =  seq(.841, .881, .02),
           label = rev(df$nb.rk[df$t == 1]),
           color = rev(c("#24ff24","#004949", "#006ddb")),
           size = 3) +
  annotate(geom = 'text',
           x = rep(7, 3), y =  seq(.841, .881, .02),
           label = rev(df$nb.rk[df$t == 7]),
           color = rev(c("#24ff24","#004949", "#006ddb")),
           size = 3) +
  annotate(geom = 'text',
           x = rep(14, 3), y =  seq(.841, .881, .02),
           label = rev(df$nb.rk[df$t == 14]),
           color = rev(c("#24ff24","#004949", "#006ddb")),
           size = 3) +
  annotate(geom = 'text', x = 14.8, y = .875, label = 'n.s', size = 3) +
  annotate(geom = "line", x = rep(14.4,2), y = c(.865,.885)) +
  annotate(geom = 'text', x = 15.5, y = .855, label = 'n.s', size = 3) +
  annotate(geom = "line", x = rep(15.15,2), y = c(.845,.885))

p.albo
```

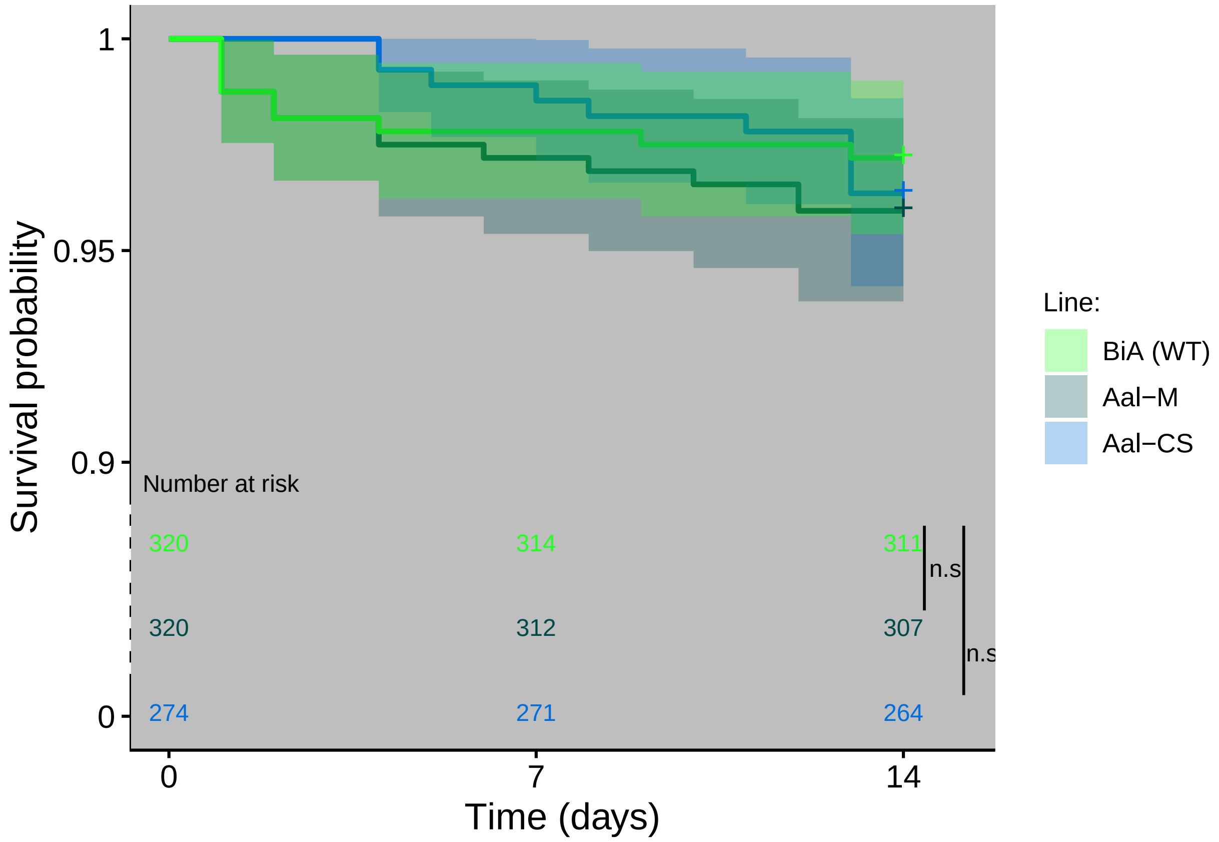

1
0.95
0.9
0
Survival probability
Number at risk
320
314
311
320
312
307
274
271
264
n.s
n.s
0
7
14
Time (days)
Line:
BiA (WT)
Aal−M
Aal−CS

### Figure 5

#### Data

```
speed.aeg = read.csv('data/Speed-test_aeg.csv', sep = ';') %>%
  mutate(recovery = 100*recovered_males/total_males) %>%
  mutate(contamination = 100*contamination/total_males)

speed.albo = read.csv('data/Speed-test_albo.csv', sep = ';') %>%
  mutate(recovery = 100*recovered_males/total_males) %>%
  mutate(contamination = 100*contamination/total_males)

head(speed.aeg)
```

```
  Replicate Volume Larval_concentration Mean_speed total_males recovered_males
1         1    500                   20          6    3924.536            3619
2         2    500                   20          6    3898.820            3578
3         3    500                   20          6    3829.696            3556
4         1    250                   40         12    3672.180            3328
5         2    250                   40         12    3366.765            2927
6         3    250                   40         12    3408.405            3037
  contamination recovery
1             0 92.21472
2             0 91.77136
3             0 92.85332
4             0 90.62737
5             0 86.93805
6             0 89.10326
```

#### Model

##### Recovery

.

```
speed.lm = lm(formula = recovery~Mean_speed,
              data = speed.aeg)

speed.poly2 = lm(formula = recovery~poly(Mean_speed, degree = 2),
              data = speed.aeg)

speed.poly3 = lm(formula = recovery~poly(Mean_speed, degree = 3),
              data = speed.aeg)

speed.poly4 = lm(formula = recovery~poly(Mean_speed, degree = 4),
              data = speed.aeg)

speed.log = lm(formula = recovery~log(Mean_speed),
              data = speed.aeg)

compare_performance(speed.lm, speed.log,
```

```
                    speed.poly2, speed.poly3,
                    speed.poly4)
```

```
# Comparison of Model Performance Indices

Name        | Model |     AIC | AIC weights |     BIC | BIC weights |    R2 | R2 (adj.) |  RMSE | Sigma
----------------------------------------------------------------------------------------------------
speed.lm    |    lm | 107.090 |     < 0.001 | 109.215 |     < 0.001 | 0.917 |     0.911 | 7.034 | 7.556
speed.log   |    lm | 103.154 |     < 0.001 | 105.278 |     < 0.001 | 0.936 |     0.932 | 6.169 | 6.626
speed.poly2 |    lm |  82.932 |       0.409 |  85.764 |       0.518 | 0.986 |     0.983 | 2.941 | 3.288
speed.poly3 |    lm |  82.824 |       0.432 |  86.365 |       0.383 | 0.987 |     0.984 | 2.742 | 3.201
speed.poly4 |    lm |  84.823 |       0.159 |  89.072 |       0.099 | 0.987 |     0.982 | 2.742 | 3.358
```

Best model:

```
summary(speed.poly2)
```

```
Call:
lm(formula = recovery ~ poly(Mean_speed, degree = 2), data = speed.aeg)

Residuals:
    Min      1Q  Median      3Q     Max 
-6.2202 -1.0021  0.2709  1.1871  6.7801 

Coefficients:
                              Estimate Std. Error t value Pr(>|t|)    
(Intercept)                     65.498      0.849  77.144  < 2e-16 ***
poly(Mean_speed, degree = 2)1  -90.774      3.288 -27.605 3.15e-12 ***
poly(Mean_speed, degree = 2)2   24.746      3.288   7.526 6.99e-06 ***
---
Signif. codes:  0 '***' 0.001 '**' 0.01 '*' 0.05 '.' 0.1 ' ' 1

Residual standard error: 3.288 on 12 degrees of freedom
Multiple R-squared:  0.9856,	Adjusted R-squared:  0.9831 
F-statistic: 409.3 on 2 and 12 DF,  p-value: 9.089e-12
```

##### Contamination

.

```
conta.lm = lm(formula = contamination~Mean_speed,
              data = speed.aeg)

conta.poly2 = lm(formula = contamination~poly(Mean_speed, degree = 2),
              data = speed.aeg)

conta.poly3 = lm(formula = contamination~poly(Mean_speed, degree = 3),
              data = speed.aeg)

conta.poly4 = lm(formula = contamination~poly(Mean_speed, degree = 4),
              data = speed.aeg)
```

```r
conta.log = lm(formula = contamination~log(Mean_speed),
              data = speed.aeg)

compare_performance(conta.lm,
                    conta.log,
                    conta.poly2,
                    conta.poly3,
                    conta.poly4)
```

```
# Comparison of Model Performance Indices

Name        | Model |     AIC | AIC weights |     BIC | BIC weights |    R2 | R2 (adj.) |  RMSE | Sigma
------------------------------------------------------------------------------------------------------
conta.lm    |    lm | -17.088 |       0.138 | -14.963 |       0.207 | 0.626 |     0.597 | 0.112 | 0.120
conta.log   |    lm |  -8.249 |       0.002 |  -6.125 |       0.002 | 0.326 |     0.274 | 0.150 | 0.162
conta.poly2 |    lm | -19.887 |       0.561 | -17.055 |       0.588 | 0.729 |     0.683 | 0.096 | 0.107
conta.poly3 |    lm | -18.007 |       0.219 | -14.467 |       0.161 | 0.731 |     0.657 | 0.095 | 0.111
conta.poly4 |    lm | -16.007 |       0.081 | -11.759 |       0.042 | 0.731 |     0.623 | 0.095 | 0.117
```

Best model

```r
summary(conta.lm)
```

```
Call:
lm(formula = contamination ~ Mean_speed, data = speed.aeg)

Residuals:
     Min       1Q   Median       3Q      Max 
-0.25076 -0.06549  0.03905  0.04709  0.24102 

Coefficients:
              Estimate Std. Error t value Pr(>|t|)    
(Intercept) -0.0551361  0.0422440  -1.305 0.214460    
Mean_speed   0.0013403  0.0002872   4.667 0.000441 ***
---
Signif. codes:  0 '***' 0.001 '**' 0.01 '*' 0.05 '.' 0.1 ' ' 1

Residual standard error: 0.1204 on 13 degrees of freedom
Multiple R-squared:  0.6262,    Adjusted R-squared:  0.5975 
F-statistic: 21.78 on 1 and 13 DF,  p-value: 0.0004409
```

##### Plot

.

```r
pred.speed = ggpredict(speed.poly2) %>%
  data.frame()
pred.conta = ggpredict(conta.lm) %>%
  data.frame()

reco.plot =
```

```r
ggplot(data = NULL) +
annotate(geom = 'line', x = c(60, 60), y = c(-Inf,70),
         lty =2, color = 'gray', size = 1 ) +
annotate(geom = 'line', x = c(-Inf, 60), y = c(70,70),
         lty =2, color = 'gray', size = 1 ) +
geom_ribbon(data = pred.speed,
            aes(x = Mean_speed.x,
                ymin = Mean_speed.conf.low,
                ymax = Mean_speed.conf.high),
            fill = "grey", alpha=.3) +
geom_line(data= pred.speed,
          aes(Mean_speed.x,
              Mean_speed.predicted),
          col = "black") +
geom_ribbon(data = pred.conta,
            aes(x = Mean_speed.x,
                ymin = Mean_speed.conf.low,
                ymax = Mean_speed.conf.high),
            fill = "red", alpha=.3) +
geom_line(data= pred.conta,
          aes(x = Mean_speed.x,
              y = Mean_speed.predicted),
          col="red") +
geom_point(data = speed.aeg,
           aes(x = Mean_speed,
               y = recovery,
               col = 'Recovery', pch = "Ae. Aegypti")) +
geom_point(data = speed.aeg,
           aes(x = Mean_speed,
               y = contamination,
               col = 'Contamination', pch = "Ae. Aegypti")) +
geom_point(data = speed.albo,
           aes(x = Mean_speed,
               y = recovery,
               col = 'Recovery', pch = "Ae. Albopictus")) +
geom_point(data = speed.albo,
           aes(x = Mean_speed,
               y = contamination,
               col = 'Contamination', pch = "Ae. Albopictus")) +
annotate(geom='text', x = -27, y =70, label = bquote(italic('70')), color = 'gray') +
annotate(geom='text', x = 275, y =50, label = bquote(R^2 ~ '= 98.3%'), color = 'black') +
annotate(geom='text', x = 275, y =10, label = bquote(R^2 ~ '= 55.6%'), color = 'red') +
labs(x = 'Sorting speed (larvae/sec) \n [larval concentration (larvae/mL)]',
     y = 'Percentage of recovery\n and contamination (%)',
     color = "Measurement" ,
     pch = "Species") +
scale_y_continuous(breaks = seq(0,100,25)) +
scale_x_continuous(breaks = seq(0, 300, 60),
                   labels = c("0 [0]", '60 [200]', '120 [400]',
                              '180 [600]', '240 [800]', '300 [1000]')) +
scale_shape_manual(labels = c(bquote(italic("Ae. aegypti")),
                              bquote(italic("Ae. albopictus"))),
                   values = c(16, 17)) +
```

```
  scale_color_manual(labels = c("Recovery",
                                "Contamination"),
                     values = c(Recovery = "black", Contamination = "red")) +
  coord_cartesian(xlim = c(0,300), ylim = c(-5,100), clip = 'off') +
  theme_classic() +
  theme(text = element_text(size = 15),
        legend.position = 'right')

reco.plot
```

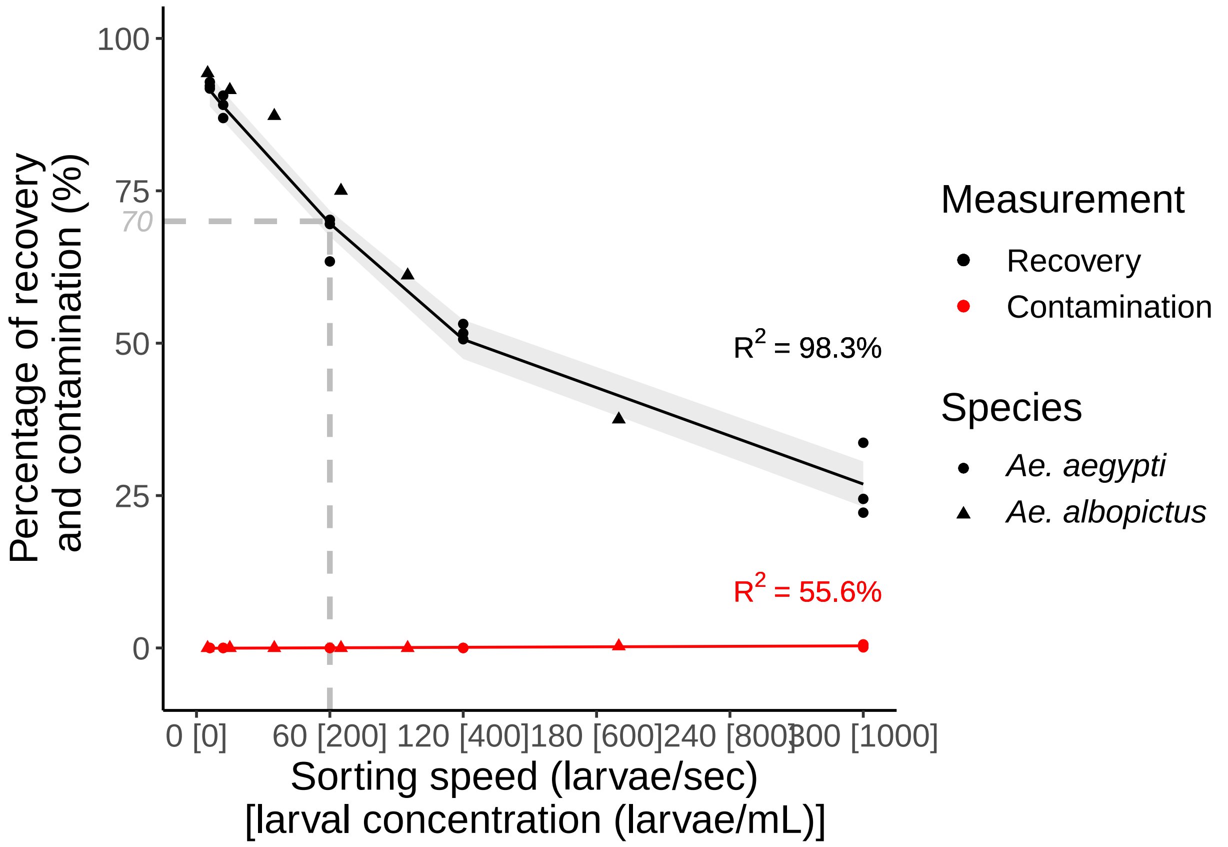

### Supplementary Data 3 - Production costs and initial parameters

Lutrat et al. 2022

**Initial parameters** 9

### Mass rearing processes

#### Size-based sorting

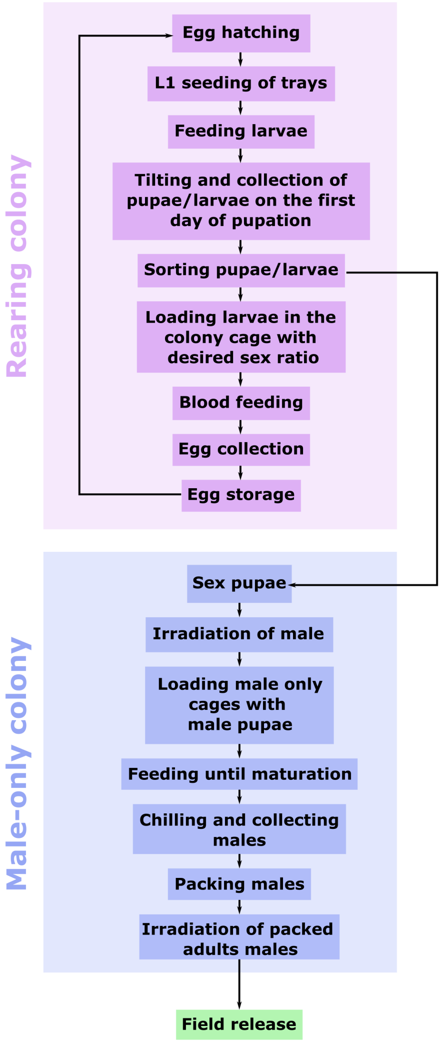

GSS

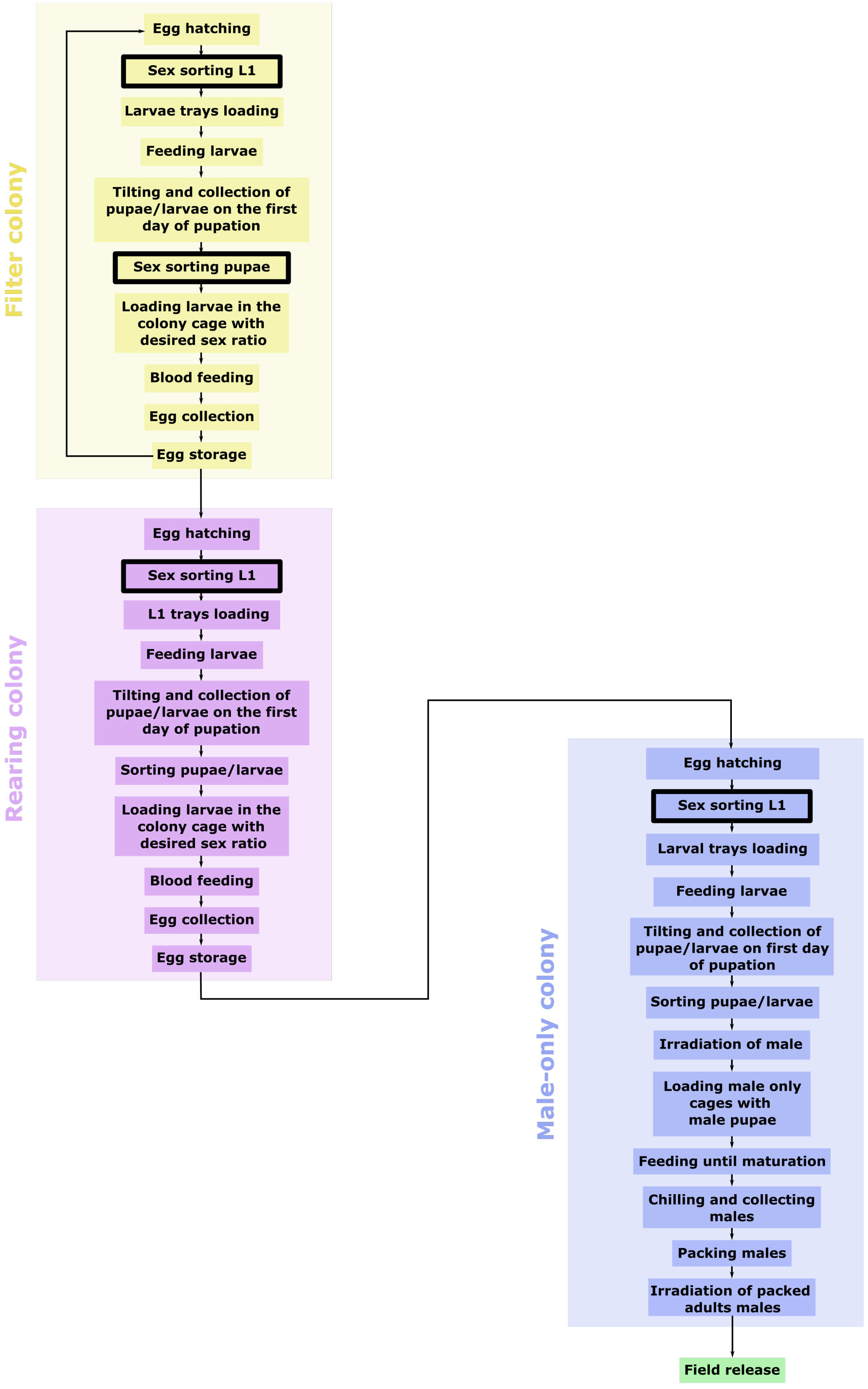

Filter colony
Egg hatching
Sex sorting L1
Larvae trays loading
Feeding larvae
Tilting and collection of pupae/larvae on the first day of pupation
Sex sorting pupae
Loading larvae in the colony cage with desired sex ratio
Blood feeding
Egg collection
Egg storage
Rearing colony
Egg hatching
Sex sorting L1
L1 trays loading
Feeding larvae
Tilting and collection of pupae/larvae on the first day of pupation
Sorting pupae/larvae
Loading larvae in the colony cage with desired sex ratio
Blood feeding
Egg collection
Egg storage
Male-only colony
Egg hatching
Sex sorting L1
Larval trays loading
Feeding larvae
Tilting and collection of pupae/larvae on first day of pupation
Sorting pupae/larvae
Irradiation of male
Loading male only cages with male pupae
Feeding until maturation
Chilling and collecting males
Packing males
Irradiation of packed adults males
Field release

#### GSS-CS

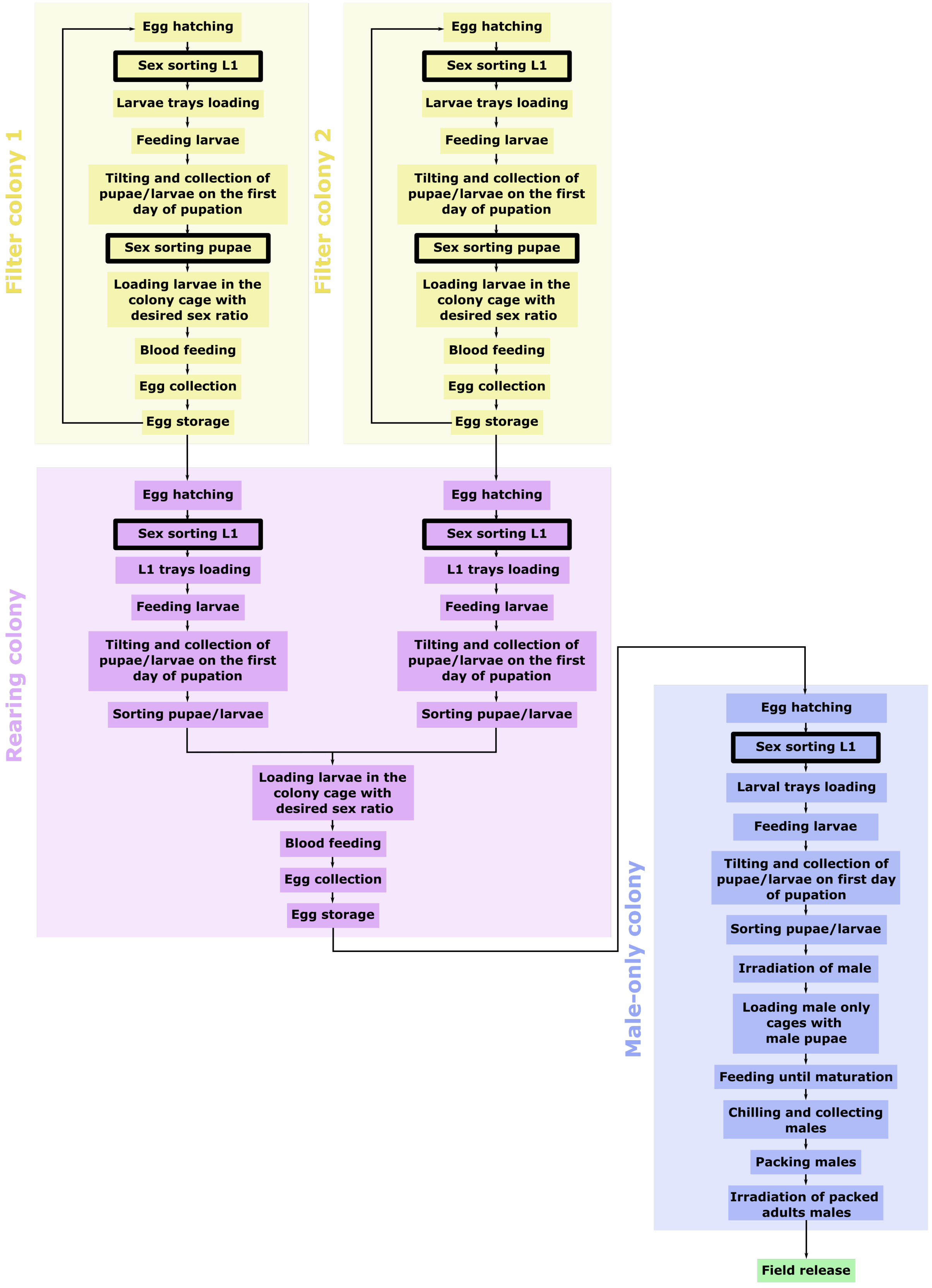

Filter colony 1
Egg hatching
Sex sorting L1
Larvae trays loading
Feeding larvae
Tilting and collection of pupae/larvae on the first day of pupation
Sex sorting pupae
Loading larvae in the colony cage with desired sex ratio
Blood feeding
Egg collection
Egg storage
Filter colony 2
Egg hatching
Sex sorting L1
Larvae trays loading
Feeding larvae
Tilting and collection of pupae/larvae on the first day of pupation
Sex sorting pupae
Loading larvae in the colony cage with desired sex ratio
Blood feeding
Egg collection
Egg storage
Rearing colony
Egg hatching
Sex sorting L1
L1 trays loading
Feeding larvae
Tilting and collection of pupae/larvae on the first day of pupation
Sorting pupae/larvae
Egg hatching
Sex sorting L1
L1 trays loading
Feeding larvae
Tilting and collection of pupae/larvae on the first day of pupation
Sorting pupae/larvae
Loading larvae in the colony cage with desired sex ratio
Blood feeding
Egg collection
Egg storage
Male-only colony
Egg hatching
Sex sorting L1
Larval trays loading
Feeding larvae
Tilting and collection of pupae/larvae on first day of pupation
Sorting pupae/larvae
Irradiation of male
Loading male only cages with male pupae
Feeding until maturation
Chilling and collecting males
Packing males
Irradiation of packed adults males
Field release

### Simulation outputs

#### Insect production

| Number of released males (M) | Size manual | Size auto | GSS | GSS-CS | GSS vs. Size manual | GSS-CS vs. Size manual |
|---|---|---|---|---|---|---|
| **Number of eggs to be hatched (M/day)** | | | | | | |
| 5 | 7.96 | 8.44 | 5.49 | 6.46 | -2.46 (-31%) | -1.49 (-18.8%) |
| 10 | 15.92 | 16.88 | 10.99 | 12.93 | -4.93 (-31%) | -2.99 (-18.8%) |
| 20 | 31.83 | 33.75 | 21.98 | 25.86 | -9.86 (-31%) | -5.98 (-18.8%) |
| 50 | 79.58 | 84.38 | 54.94 | 64.64 | -24.64 (-31%) | -14.94 (-18.8%) |
| 100 | 159.16 | 168.76 | 109.88 | 129.28 | -49.29 (-31%) | -29.89 (-18.8%) |
| **Number of larvae in the mass-rearing facility (M)** | | | | | | |
| 5 | 32.98 | 34.96 | 11.56 | 17.82 | -21.42 (-65%) | -15.16 (-46%) |
| 10 | 65.95 | 69.93 | 23.00 | 35.53 | -42.95 (-65.1%) | -30.42 (-46.1%) |
| 20 | 131.91 | 139.86 | 46.01 | 71.06 | -85.90 (-65.1%) | -60.84 (-46.1%) |
| 50 | 329.77 | 349.65 | 114.59 | 176.90 | -215.18 (-65.3%) | -152.86 (-46.4%) |
| 100 | 659.53 | 699.30 | 228.96 | 353.48 | -430.57 (-65.3%) | -306.05 (-46.4%) |
| **Number of adults in the mass-rearing facility (M)** | | | | | | |
| 5 | 4.84 | 4.84 | 3.63 | 4.44 | -1.21 (-25%) | -0.40 (-8.3%) |
| 10 | 9.28 | 9.68 | 6.45 | 7.66 | -2.82 (-30.4%) | -1.61 (-17.4%) |
| 20 | 18.15 | 18.96 | 12.91 | 15.33 | -5.24 (-28.9%) | -2.82 (-15.6%) |
| 50 | 44.77 | 47.59 | 31.46 | 37.51 | -13.31 (-29.7%) | -7.26 (-16.2%) |
| 100 | 89.14 | 94.78 | 62.92 | 75.02 | -26.22 (-29.4%) | -14.12 (-15.8%) |

#### Facility area

| Number of released males (M) | Size manual | Size auto | GSS | GSS-CS | GSS vs. Size manual | GSS-CS vs. Size manual |
|---|---|---|---|---|---|---|
| **Mass rearing facility area (m2)** | | | | | | |
| 5 | 535 | 539 | 417 | 470 | -118 (-22.1%) | -65 (-12.2%) |
| 10 | 839 | 876 | 565 | 661 | -274 (-32.6%) | -178 (-21.2%) |
| 20 | 1,466 | 1,519 | 893 | 1,098 | -573 (-39.1%) | -368 (-25.1%) |
| 50 | 3,321 | 3,516 | 1,895 | 2,389 | -1,427 (-43%) | -933 (-28.1%) |
| 100 | 6,437 | 6,800 | 3,553 | 4,533 | -2,884 (-44.8%) | -1,903 (-29.6%) |
| **Release facility area (m2)** | | | | | | |
| 5 | 174 | 174 | 174 | 174 | 0 (0%) | 0 (0%) |
| 10 | 228 | 228 | 228 | 228 | 0 (0%) | 0 (0%) |
| 20 | 338 | 338 | 338 | 338 | 0 (0%) | 0 (0%) |
| 50 | 667 | 667 | 667 | 667 | 0 (0%) | 0 (0%) |
| 100 | 1,215 | 1,215 | 1,215 | 1,215 | 0 (0%) | 0 (0%) |

#### Workload (both facilities)

| Number of released males (M) | Size manual | Size auto | GSS | GSS-CS | GSS vs. Size manual | GSS-CS vs. Size manual |
|---|---|---|---|---|---|---|
| **Number of rearing labourers** | | | | | | |
| 5 | 40 | 42 | 28 | 32 | -12 (-30%) | -8 (-20%) |
| 10 | 76 | 79 | 48 | 57 | -28 (-36.8%) | -19 (-25%) |
| 20 | 151 | 156 | 95 | 110 | -56 (-37.1%) | -41 (-27.2%) |
| 50 | 368 | 385 | 230 | 269 | -138 (-37.5%) | -99 (-26.9%) |
| 100 | 733 | 766 | 453 | 531 | -280 (-38.2%) | -202 (-27.6%) |
| **Number of QC technicians** | | | | | | |
| 5 | 4 | 4 | 2 | 3 | -2 (-50%) | -1 (-25%) |
| 10 | 6 | 6 | 3 | 4 | -3 (-50%) | -2 (-33.3%) |
| 20 | 14 | 14 | 8 | 9 | -6 (-42.9%) | -5 (-35.7%) |
| 50 | 36 | 37 | 21 | 25 | -15 (-41.7%) | -11 (-30.6%) |
| 100 | 72 | 75 | 41 | 50 | -31 (-43.1%) | -22 (-30.6%) |
| **Number of maintenance officers** | | | | | | |
| 5 | 2 | 2 | 1 | 1 | -1 (-50%) | -1 (-50%) |
| 10 | 2 | 2 | 1 | 1 | -1 (-50%) | -1 (-50%) |
| 20 | 6 | 6 | 3 | 4 | -3 (-50%) | -2 (-33.3%) |
| 50 | 18 | 19 | 11 | 13 | -7 (-38.9%) | -5 (-27.8%) |
| 100 | 38 | 40 | 23 | 27 | -15 (-39.5%) | -11 (-28.9%) |
| **Total number of staff for both facilities** | | | | | | |
| 5 | 55 | 57 | 39 | 45 | -16 (-29.1%) | -10 (-18.2%) |
| 10 | 102 | 105 | 65 | 78 | -37 (-36.3%) | -24 (-23.5%) |
| 20 | 200 | 206 | 127 | 147 | -73 (-36.5%) | -53 (-26.5%) |
| 50 | 483 | 504 | 303 | 353 | -180 (-37.3%) | -130 (-26.9%) |
| 100 | 957 | 1,000 | 588 | 692 | -369 (-38.6%) | -265 (-27.7%) |

#### Water requirements

| Number of released males (M) | Size manual | Size auto | GSS | GSS-CS | GSS vs. Size manual | GSS-CS vs. Size manual |
|---|---|---|---|---|---|---|
| **Daily water requirements (L)** | | | | | | |
| 5 | 3,704 | 3,900 | 1,580 | 2,202 | -2,124 (-57.3%) | -1,502 (-40.6%) |
| 10 | 7,403 | 7,799 | 3,142 | 4,383 | -4,261 (-57.6%) | -3,021 (-40.8%) |
| 20 | 14,802 | 15,594 | 6,284 | 8,766 | -8,518 (-57.5%) | -6,037 (-40.8%) |
| 50 | 36,999 | 38,986 | 15,658 | 21,831 | -21,341 (-57.7%) | -15,168 (-41%) |
| 100 | 73,993 | 77,968 | 31,296 | 43,630 | -42,698 (-57.7%) | -30,364 (-41%) |

#### Construction budget

| Number of released males (M) | Size manual | Size auto | GSS | GSS-CS | GSS vs. Size manual | GSS-CS vs. Size manual |
|---|---|---|---|---|---|---|
| **Construction cost of the mass rearing facility ($)** | | | | | | |
| 5 | 653,350 | 656,800 | 511,188 | 575,122 | -142,163 (-21.8%) | -78,228 (-12%) |
| 10 | 1,019,988 | 1,063,840 | 690,876 | 805,563 | -329,112 (-32.3%) | -214,424 (-21%) |
| 20 | 1,776,099 | 1,840,034 | 1,086,457 | 1,333,551 | -689,642 (-38.8%) | -442,548 (-24.9%) |
| 50 | 4,015,630 | 4,250,948 | 2,296,329 | 2,893,922 | -1,719,301 (-42.8%) | -1,121,708 (-27.9%) |

*(continued)*

| Number of released males (M) | Size manual | Size auto | GSS | GSS-CS | GSS vs. Size manual | GSS-CS vs. Size manual |
|---|---|---|---|---|---|---|
| 100 | 7,774,879 | 8,213,995 | 4,300,135 | 5,485,650 | -3,474,744 (-44.7%) | -2,289,229 (-29.4%) |
| **Yearly depreciation of the mass rearing facility ($)** | | | | | | |
| 5 | 32,668 | 32,840 | 25,559 | 28,756 | -7,108 (-21.8%) | -3,911 (-12%) |
| 10 | 50,999 | 53,192 | 34,544 | 40,278 | -16,456 (-32.3%) | -10,721 (-21%) |
| 20 | 88,805 | 92,002 | 54,323 | 66,678 | -34,482 (-38.8%) | -22,127 (-24.9%) |
| 50 | 200,782 | 212,547 | 114,816 | 144,696 | -85,965 (-42.8%) | -56,085 (-27.9%) |
| 100 | 388,744 | 410,700 | 215,007 | 274,282 | -173,737 (-44.7%) | -114,461 (-29.4%) |
| **Construction cost of the release facility ($)** | | | | | | |
| 5 | 237,901 | 237,901 | 237,901 | 237,901 | 0 (0%) | 0 (0%) |
| 10 | 311,603 | 311,603 | 311,603 | 311,603 | 0 (0%) | 0 (0%) |
| 20 | 459,006 | 459,006 | 459,006 | 459,006 | 0 (0%) | 0 (0%) |
| 50 | 901,214 | 901,214 | 901,214 | 901,214 | 0 (0%) | 0 (0%) |
| 100 | 1,638,082 | 1,638,082 | 1,638,082 | 1,638,082 | 0 (0%) | 0 (0%) |
| **Yearly depreciation of the release facility ($)** | | | | | | |
| 5 | 11,895 | 11,895 | 11,895 | 11,895 | 0 (0%) | 0 (0%) |
| 10 | 15,580 | 15,580 | 15,580 | 15,580 | 0 (0%) | 0 (0%) |
| 20 | 22,950 | 22,950 | 22,950 | 22,950 | 0 (0%) | 0 (0%) |
| 50 | 45,061 | 45,061 | 45,061 | 45,061 | 0 (0%) | 0 (0%) |
| 100 | 81,904 | 81,904 | 81,904 | 81,904 | 0 (0%) | 0 (0%) |

#### Equipment budget

| Number of released males (M) | Size manual | Size auto | GSS | GSS-CS | GSS vs. Size manual | GSS-CS vs. Size manual |
|---|---|---|---|---|---|---|
| **Mass rearing facility equipment cost ($)** | | | | | | |
| 5 | 1,093,510 | 1,831,030 | 1,247,100 | 1,448,810 | 153,590 (14%) | 355,300 (32.5%) |
| 10 | 1,791,920 | 3,249,980 | 1,913,010 | 2,314,060 | 121,090 (6.8%) | 522,140 (29.1%) |
| 20 | 3,260,960 | 6,055,430 | 3,238,590 | 4,103,040 | -22,370 (-0.7%) | 842,080 (25.8%) |
| 50 | 7,534,210 | 14,563,840 | 7,270,080 | 9,493,760 | -264,130 (-3.5%) | 1,959,550 (26%) |
| 100 | 14,751,890 | 28,653,800 | 13,933,530 | 18,352,250 | -818,360 (-5.5%) | 3,600,360 (24.4%) |
| **Mass rearing facility equipment yearly depreciation ($)** | | | | | | |
| 5 | 132,888 | 207,608 | 136,497 | 159,878 | 3,608 (2.7%) | 26,990 (20.3%) |
| 10 | 222,067 | 369,903 | 210,832 | 257,063 | -11,235 (-5.1%) | 34,997 (15.8%) |
| 20 | 408,607 | 692,415 | 357,415 | 456,713 | -51,192 (-12.5%) | 48,107 (11.8%) |
| 50 | 950,178 | 1,665,433 | 803,147 | 1,059,787 | -147,032 (-15.5%) | 109,608 (11.5%) |
| 100 | 1,865,932 | 3,279,073 | 1,541,245 | 2,050,885 | -324,687 (-17.4%) | 184,953 (9.9%) |
| **Release facility equipment cost** | | | | | | |
| 5 | 88,508 | 88,508 | 88,508 | 88,508 | 0 (0%) | 0 (0%) |
| 10 | 141,016 | 141,016 | 141,016 | 141,016 | 0 (0%) | 0 (0%) |
| 20 | 246,032 | 246,032 | 246,032 | 246,032 | 0 (0%) | 0 (0%) |
| 50 | 561,080 | 561,080 | 561,080 | 561,080 | 0 (0%) | 0 (0%) |
| 100 | 1,086,076 | 1,086,076 | 1,086,076 | 1,086,076 | 0 (0%) | 0 (0%) |
| **Release facility equipment yearly depreciation ($)** | | | | | | |
| 5 | 17,660 | 17,660 | 17,660 | 17,660 | 0 (0%) | 0 (0%) |
| 10 | 29,121 | 29,121 | 29,121 | 29,121 | 0 (0%) | 0 (0%) |
| 20 | 52,041 | 52,041 | 52,041 | 52,041 | 0 (0%) | 0 (0%) |
| 50 | 120,803 | 120,803 | 120,803 | 120,803 | 0 (0%) | 0 (0%) |
| 100 | 235,386 | 235,386 | 235,386 | 235,386 | 0 (0%) | 0 (0%) |
| **Total equipment cost** | | | | | | |
| 5 | 1,182,018 | 1,919,538 | 1,335,608 | 1,537,318 | 153,590 (13%) | 355,300 (30.1%) |

*(continued)*

| Number of released males (M) | Size manual | Size auto | GSS | GSS-CS | GSS vs. Size manual | GSS-CS vs. Size manual |
|---|---|---|---|---|---|---|
| 10 | 1,932,936 | 3,390,996 | 2,054,026 | 2,455,076 | 121,090 (6.3%) | 522,140 (27%) |
| 20 | 3,506,992 | 6,301,462 | 3,484,622 | 4,349,072 | -22,370 (-0.6%) | 842,080 (24%) |
| 50 | 8,095,290 | 15,124,920 | 7,831,160 | 10,054,840 | -264,130 (-3.3%) | 1,959,550 (24.2%) |
| 100 | 15,837,966 | 29,739,876 | 15,019,606 | 19,438,326 | -818,360 (-5.2%) | 3,600,360 (22.7%) |
| **Total equipment yearly depreciation ($)** | | | | | | |
| 5 | 150,549 | 225,269 | 154,157 | 177,539 | 3,608 (2.4%) | 26,990 (17.9%) |
| 10 | 251,187 | 399,024 | 239,952 | 286,184 | -11,235 (-4.5%) | 34,997 (13.9%) |
| 20 | 460,648 | 744,456 | 409,456 | 508,755 | -51,192 (-11.1%) | 48,107 (10.4%) |
| 50 | 1,070,982 | 1,786,237 | 923,950 | 1,180,590 | -147,032 (-13.7%) | 109,608 (10.2%) |
| 100 | 2,101,317 | 3,514,459 | 1,776,631 | 2,286,271 | -324,687 (-15.5%) | 184,953 (8.8%) |

#### Diet and consumables yearly cost

| Number of released males (M) | Size manual | Size auto | GSS | GSS-CS | GSS vs. Size manual | GSS-CS vs. Size manual |
|---|---|---|---|---|---|---|
| **Yearly diet cost** | | | | | | |
| 5 | 30,720 | 32,346 | 13,376 | 18,742 | -17,344 (-56.5%) | -11,978 (-39%) |
| 10 | 61,388 | 64,692 | 26,599 | 37,327 | -34,789 (-56.7%) | -24,061 (-39.2%) |
| 20 | 122,723 | 129,331 | 53,169 | 74,585 | -69,553 (-56.7%) | -48,138 (-39.2%) |
| 50 | 306,728 | 323,340 | 132,452 | 185,717 | -174,276 (-56.8%) | -121,011 (-39.5%) |
| 100 | 613,402 | 646,628 | 264,721 | 371,166 | -348,681 (-56.8%) | -242,236 (-39.5%) |
| **Yearly consumables cost** | | | | | | |
| 5 | 8,110 | 8,435 | 4,641 | 5,714 | -3,469 (-42.8%) | -2,396 (-29.5%) |
| 10 | 15,554 | 16,214 | 8,596 | 10,741 | -6,958 (-44.7%) | -4,812 (-30.9%) |
| 20 | 30,441 | 31,763 | 16,531 | 20,814 | -13,911 (-45.7%) | -9,628 (-31.6%) |
| 50 | 75,105 | 78,427 | 40,250 | 50,903 | -34,855 (-46.4%) | -24,202 (-32.2%) |
| 100 | 149,544 | 156,189 | 79,807 | 101,096 | -69,736 (-46.6%) | -48,447 (-32.4%) |
| **Yearly cost for diet and consumables** | | | | | | |
| 5 | 38,830 | 40,781 | 18,017 | 24,456 | -20,813 (-53.6%) | -14,374 (-37%) |
| 10 | 76,941 | 80,906 | 35,194 | 48,069 | -41,747 (-54.3%) | -28,873 (-37.5%) |
| 20 | 153,164 | 161,094 | 69,700 | 95,399 | -83,464 (-54.5%) | -57,765 (-37.7%) |
| 50 | 381,832 | 401,768 | 172,701 | 236,619 | -209,131 (-54.8%) | -145,213 (-38%) |
| 100 | 762,946 | 802,817 | 344,528 | 472,263 | -418,418 (-54.8%) | -290,683 (-38.1%) |

#### Estimate of the total yearly cost for construction, equipment, diet and consumables

| Number of released males (M) | Size manual | Size auto | GSS | GSS-CS | GSS vs. Size manual | GSS-CS vs. Size manual |
|---|---|---|---|---|---|---|
| 5 | 233,941 | 310,785 | 209,629 | 242,646 | -24,313 (-10.4%) | 8,705 (3.7%) |
| 10 | 394,708 | 548,703 | 325,271 | 390,111 | -69,438 (-17.6%) | -4,597 (-1.2%) |
| 20 | 725,567 | 1,020,502 | 556,430 | 693,781 | -169,138 (-23.3%) | -31,786 (-4.4%) |
| 50 | 1,698,656 | 2,445,613 | 1,256,529 | 1,606,966 | -442,128 (-26%) | -91,690 (-5.4%) |
| 100 | 3,334,911 | 4,809,880 | 2,418,070 | 3,114,720 | -916,842 (-27.5%) | -220,192 (-6.6%) |

### Initial parameters

Weekly flying male production (production) [5, 10, 20, 50, 100] M males

| Parameters | Size manual | Size auto | GSS | GSS-CS |
|---|---|---|---|---|
| **Production goals** | | | | |
| Weekly pupal production (Mpp/week) | production / 0.95 | production / 0.95 | production / 0.95 | production / 0.95 |
| *Rearing efficienty* | | | | |
| Number of weeks with releases per year | 52 | 52 | 52 | 52 |
| **Biological parameters** | | | | |
| Egg hatching rate | 0.85 | 0.85 | 0.85 | 0.85 |
| Sex ratio (M/(F+M)) | 0.5 | 0.5 | 0.5 | 0.5 |
| Male pupae recovery after first sorting | 0.65 | 0.65 | 0.65 | 0.65 |
| Male pupae recovery after second sorting | 0.15 | 0.15 | 0.15 | 0.15 |
| Male pupae recovery after third sorting | 0.04 | 0.04 | 0.04 | 0.04 |
| Female pupae recovery after first sorting | 0.32 | 0.32 | 0.32 | 0.32 |
| Female pupae recovery after second sorting | 0.35 | 0.35 | 0.35 | 0.35 |
| Female pupae recovery after third sorting | 0.36 | 0.36 | 0.36 | 0.36 |
| Number of tilting or sex sorting operations for MO | 1 | 1 | 1 | 1 |
| **Number of tilting or sex sorting operations for colony** | **2** | **2** | **1** | **1** |
| Survival pupae to flying males | 0.95 | 0.95 | 0.95 | 0.95 |
| Survival pupae to flying females | 0.95 | 0.95 | 0.95 | 0.95 |
| Average number of eggs per females for the 1st gonotrophic cycle | 30 | 30 | 30 | 30 |
| Average number of eggs per females for the 2nd gonotrophic cycle | 20 | 20 | 20 | 20 |
| Average number of eggs per females for the 3rd gonotrophic cycle | 10 | 10 | 10 | 10 |
| *Life cycle information* | | | | |
| Pre oviposition period (days) | 8 | 8 | 8 | 8 |
| Duration of gonotrophic cycle (days) | 7 | 7 | 7 | 7 |
| Number of gonotrophic cycles colonies | 3 | 3 | 3 | 3 |
| Duration of males holding before release (days) | 4 | 4 | 4 | 4 |
| Duration of the larval cycle Colony (days) | 8 | 8 | 8 | 8 |
| Egg density (eggs/mL) | 91000 | 91000 | 91000 | 91000 |
| Pupa density (pupae/L) | 333333 | 333333 | 333333 | 333333 |
| **Capacity of main equipment** | | | | |
| *Larval trays* | | | | |
| Density of larvae in colony trays (larvae/mL) | 3 | 3 | 3 | 3 |
| Depth of colony larval tray (cm) | 1 | 1 | 1 | 1 |
| Larval tray length (cm) | 60 | 60 | 60 | 60 |
| Larval tray width (cm) | 100 | 100 | 100 | 100 |
| Height of tray columns (cm) | 170 | 170 | 170 | 170 |
| Free space between trays (cm) | 2.5 | 2.5 | 2.5 | 2.5 |

*(continued)*

| Parameters | Size manual | Size auto | GSS | GSS-CS |
|---|---|---|---|---|
| *MO cages* | | | | |
| Vertical resting space in MO cages (cm2/adult) | 1 | 1 | 1 | 1 |
| MO cage compartment (cm2 per cage) | 12 | 12 | 12 | 12 |
| MO cage height (cm) | 25 | 25 | 25 | 25 |
| MO cage width (cm) | 10 | 10 | 10 | 10 |
| MO cage length (cm) | 6 | 6 | 6 | 6 |
| *Colony cages* | | | | |
| Vertical resting place for colony cages (cm2/adult) | 1.2 | 1.2 | 1.2 | 1.2 |
| Net colony cages height (cm) | 100 | 100 | 100 | 100 |
| Net colony cages width (cm) | 20 | 20 | 20 | 20 |
| Net colony cages length (cm) | 90 | 90 | 90 | 90 |
| **Rearing task schedule** | | | | |
| Frequency of blood feeding (days) | 3 | 3 | 3 | 3 |
| Frequency of egg collection (days) | 3 | 3 | 3 | 3 |
| Number of feeding rounds per blood membrane | 5 | 5 | 5 | 5 |
| Frequency of colony cage washing (holding cycle) | 1 | 1 | 1 | 1 |
| Frequency of MO cage washing (holding cycle) | 1 | 1 | 1 | 1 |
| Increase factor mother colony | 1.08 | 1.08 | 1.08 | 1.08 |
| **Diet formula** | | | | |
| *Larval diet* | | | | |
| Beef liver powder (%) | 0 | 0 | 0 | 0 |
| Tuna meal (%) | 50 | 50 | 50 | 50 |
| Brewer yeast (%) | 15 | 15 | 15 | 15 |
| BSF yeast (%) | 35 | 35 | 35 | 35 |
| Component 5 (%) | 0 | 0 | 0 | 0 |
| Component 6 (%) | 0 | 0 | 0 | 0 |
| Concentration in water (g/mL) | 4 | 4 | 4 | 4 |
| Ration per one larva on day 1 (mg ingredients/larva) | 0.66 | 0.66 | 0.66 | 0.66 |
| Ration per one larva on day 2 (mg ingredients/larva) | 0 | 0 | 0 | 0 |
| Ration per one larva on day 3 (mg ingredients/larva) | 0 | 0 | 0 | 0 |
| Ration per one larva on day 4 (mg ingredients/larva) | 0.66 | 0.66 | 0.66 | 0.66 |
| Ration per one larva on day 5 (mg ingredients/larva) | 0.44 | 0.44 | 0.44 | 0.44 |
| Ration per one larva on day 6 (mg ingredients/larva) | 0.66 | 0.66 | 0.66 | 0.66 |
| Ration per one larva on day 7 after 1st sorting (mg ingredients/larva) | 0 | 0 | 0 | 0 |
| Ration per one larva on day 8 after 2nd sorting (mg ingredients/larva) | 0 | 0 | 0 | 0 |
| *Adult diet* | | | | |
| Sugar percentage in adult diets | 10 | 10 | 10 | 10 |
| Weight of sugar per colony cage (kg/cage) | 0.03 | 0.03 | 0.03 | 0.03 |
| Weight of sugar per MO cage (kg/cage) | 0.1 | 0.1 | 0.1 | 0.1 |
| Replenishment of water in adult cages (days) | 5 | 5 | 5 | 5 |

*(continued)*

| Parameters | Size manual | Size auto | GSS | GSS-CS |
|---|---|---|---|---|
| Average Blood intake per female (mg/female/intake) | 1.5 | 1.5 | 1.5 | 1.5 |
| **Storage of ingredients** | | | | |
| Beef liver powder storage time (days) | 90 | 90 | 90 | 90 |
| Beef liver powder storage bags per m3 | 6 | 6 | 6 | 6 |
| Beef liver powder (kg/unit) | 25 | 25 | 25 | 25 |
| Beef liver powder height (cm) | 1 | 1 | 1 | 1 |
| Tuna meal storage time (days) | 90 | 90 | 90 | 90 |
| Tuna meal storage bags per m3 | 6 | 6 | 6 | 6 |
| Tuna meal (kg/unit) | 25 | 25 | 25 | 25 |
| Tuna meal height (cm) | 1 | 1 | 1 | 1 |
| Brewer yeast storage time (days) | 90 | 90 | 90 | 90 |
| Brewer yeast storage bags per m3 | 6 | 6 | 6 | 6 |
| Brewer yeast (kg/unit) | 25 | 25 | 25 | 25 |
| Brewer yeast height (cm) | 1 | 1 | 1 | 1 |
| BSF yeast storage time (days) | 90 | 90 | 90 | 90 |
| BSF yeast storage bags per m3 | 6 | 6 | 6 | 6 |
| BSF yeast (kg/unit) | 25 | 25 | 25 | 25 |
| BSF yeast height (cm) | 1 | 1 | 1 | 1 |
| Component 5 storage time (days) | 90 | 90 | 90 | 90 |
| Component 5 storage bags per m3 | 6 | 6 | 6 | 6 |
| Component 5 (kg/unit) | 25 | 25 | 25 | 25 |
| Component 5 height (cm) | 1 | 1 | 1 | 1 |
| Component 6 storage time (days) | 90 | 90 | 90 | 90 |
| Component 6 storage bags per m3 | 6 | 6 | 6 | 6 |
| Component 6 (kg/unit) | 25 | 25 | 25 | 25 |
| Component 6 height (cm) | 1 | 1 | 1 | 1 |
| Total sugar for adult diet storage time (days) | 90 | 90 | 90 | 90 |
| Total sugar for adult diet storage bags per m3 | 6 | 6 | 6 | 6 |
| Total sugar for adult diet (kg/unit) | 50 | 50 | 50 | 50 |
| Total sugar for adult diet height (cm) | 1 | 1 | 1 | 1 |
| **Rearing equipment** | | | | |
| *Oversize factors* | | | | |
| Oversize factor larval trays | 1.3 | 1.3 | 1.3 | 1.3 |
| Oversize factor racks | 1.2 | 1.2 | 1.2 | 1.2 |
| Oversize factor cages | 1.2 | 1.2 | 1.2 | 1.2 |
| Oversize factor RM cages | 1.3 | 1.3 | 1.3 | 1.3 |
| *Irradiators* | | | | |
| Number of pupae per irradiation canister | 375000 | 375000 | 375000 | 375000 |
| Duration of one irradiation operation (hour) | 0.17 | 0.17 | 0.17 | 0.17 |
| Irradiator max time (h/day) | 12 | 12 | 12 | 12 |
| Irradiator backup | 0 | 0 | 0 | 0 |
| *L1 sex sorter* | | | | |
| L1 sex sorter male recovery | 0.696 | 0.696 | 0.696 | 0.696 |
| L1 sex sorter female recovery | 0.696 | 0.696 | 0.696 | 0.696 |
| L1 sex sorter throwput (M/hour) | 0.133 | 0.133 | 0.133 | 0.133 |
| L1 sex sorter max time (h/day) | 5 | 5 | 5 | 5 |
| L1 sex sorter backup | 1 | 1 | 1 | 1 |
| *Pupal sorter* | | | | |
| **Pupal sex sorter male recovery** | **0.427** | **0.403** | **0.403** | **0.403** |
| Pupal sex sorter female recovery | 0.3806 | 0.3806 | 0.3806 | 0.3806 |

*(continued)*

| Parameters | Size manual | Size auto | GSS | GSS-CS |
|---|---|---|---|---|
| Pupal sorter throwput (Mpp/h) | 0.05 | 0.05 | 0.05 | 0.05 |
| Pupal sorter max time (h/day) | 5 | 5 | 5 | 5 |
| Pupal sorter backup | 1 | 1 | 1 | 1 |
| Male recovery for GSS filter double check | 0.9 | 0.9 | 0.9 | 0.9 |
| *L1 counter* | | | | |
| L1 counter capacity (tray/h) | 20 | 20 | 20 | 20 |
| L1 counter max time | 4 | 4 | 4 | 4 |
| L1 counter backup | 1 | 1 | 1 | 1 |
| *Larval diet mixer* | | | | |
| Larval diet mixer capacity (trays/h) | 40 | 40 | 40 | 40 |
| Larval diet mixer duration per batch (h) | 0.25 | 0.25 | 0.25 | 0.25 |
| Larval diet mixer max time (h/day) | 3 | 3 | 3 | 3 |
| Larval diet mixer backup | 0 | 0 | 0 | 0 |
| *Adult diet mixer* | | | | |
| Adult diet mixer capacity (L/operation) | 20 | 20 | 20 | 20 |
| Adult diet mixer duration per batch (h) | 0.25 | 0.25 | 0.25 | 0.25 |
| Adult diet mixer max time (h/day) | 3 | 3 | 3 | 3 |
| Adult diet mixer backup | 0 | 0 | 0 | 0 |
| *Larval diet feeder* | | | | |
| Larval feeder capacity (trays/h) | 45 | 45 | 45 | 45 |
| Larval feeder max time (h/day) | 4 | 4 | 4 | 4 |
| Larval feeder backup | 1 | 1 | 1 | 1 |
| *Blood feeder* | | | | |
| Blood feeder capacity (cages/h) | 2 | 2 | 2 | 2 |
| Blood feeder max time (h/day) | 6 | 6 | 6 | 6 |
| Blood feeder backup | 1 | 1 | 1 | 1 |
| *Tray washer* | | | | |
| Tray washer capacity (trays/h) | 200 | 200 | 200 | 200 |
| Tray washer max time (h/day) | 6 | 6 | 6 | 6 |
| Tray washer backup | 0 | 0 | 0 | 0 |
| *Cage washer* | | | | |
| Cage washer capacity (cages/h) | 2.4 | 2.4 | 2.4 | 2.4 |
| Cage washer max time (h/day) | 6 | 6 | 6 | 6 |
| Cage washer backup | 0 | 0 | 0 | 0 |
| *MO cage washer* | | | | |
| MO cage washer capacity (cages/h) | 12 | 12 | 12 | 12 |
| MO cage washer max time (h/day) | 6 | 6 | 6 | 6 |
| MO cage washer backup | 0 | 0 | 0 | 0 |
| **Water requirement** | | | | |
| Water per tray wash (L/tray) | 0.5 | 0.5 | 0.5 | 0.5 |
| Water per cage wash (L/cage) | 2 | 2 | 2 | 2 |
| Water per MO cage wash (L/cage) | 3 | 3 | 3 | 3 |
| Percentage of water for room cleaning (%) | 0.4 | 0.4 | 0.4 | 0.4 |
| **Floor area calculation** | | | | |
| *Mass rearing facility* | | | | |
| Rack oversize factor | 2 | 2 | 2 | 2 |

*(continued)*

| Parameters | Size manual | Size auto | GSS | GSS-CS |
|---|---|---|---|---|
| Colony cage oversize factor | 2 | 2 | 2 | 2 |
| Storage oversize factor | 3 | 3 | 3 | 3 |
| Offices size per staff (m2/staff) | 5 | 5 | 5 | 5 |
| WC size per staff (m2/staff) | 3 | 3 | 3 | 3 |
| Percentage corridors (%) | 0.1 | 0.1 | 0.1 | 0.1 |
| Diet preparation area (m2) | 10 | 10 | 10 | 10 |
| Tray washing area (m2) | 15 | 15 | 15 | 15 |
| Cage washing area (m2) | 15 | 15 | 15 | 15 |
| QC lab area (m2) | 20 | 20 | 20 | 20 |
| Workshop area (m2) | 20 | 20 | 20 | 20 |
| Warehouse area (m2) | 20 | 20 | 20 | 20 |
| Office staff (m2) | 6 | 6 | 6 | 6 |
| WC staff (m2) | 10 | 10 | 10 | 10 |
| *Realese facility* | | | | |
| MO cage oversize factor | 2 | 2 | 2 | 2 |
| MO cages loading area (m2) | 15 | 15 | 15 | 15 |
| MO QC lab area (m2) | 20 | 20 | 20 | 20 |
| MO adult packaging area (m2) | 15 | 15 | 15 | 15 |
| MO diet preparation area (m2) | 15 | 15 | 15 | 15 |
| MO Warehouse area (m2) | 10 | 10 | 10 | 10 |
| MO office staff (m2) | 2 | 2 | 2 | 2 |
| MO wc staff (m2) | 6 | 6 | 6 | 6 |
| **Construction cost** | | | | |
| Cost m2 office labs wc irradiation corridor ($/m2) | 1500 | 1500 | 1500 | 1500 |
| Cost m2 rearing rooms ($/m2) | 1200 | 1200 | 1200 | 1200 |
| Cost m2 storage warehouse ($/m2) | 1000 | 1000 | 1000 | 1000 |
| Expected lifespan mass rearing facility (years) | 20 | 20 | 20 | 20 |
| Cost m2 chilling room ($/m2) | 2000 | 2000 | 2000 | 2000 |
| Expected lifespan release facility (years) | 20 | 20 | 20 | 20 |
| **Workload** | | | | |
| *Mass rearing facility* | | | | |
| Egg hatching work rate (h/day) | 2 | 2 | 2 | 2 |
| Tray hanging work rate (h/rack) | 0.45 | 0.45 | 0.45 | 0.45 |
| L1 dosage work rate (h/rack) | 0.25 | 0.25 | 0.25 | 0.25 |
| L4 loading work rate (h/rack) | 0.05 | 0.05 | 0.05 | 0.05 |
| Larval feeding work rate (h/rack) | 0.25 | 0.25 | 0.25 | 0.25 |
| Tray tilting work rate (h/rack) | 1 | 1 | 1 | 1 |
| Insect packing for irradiation work rate (h/canister) | 0 | 0 | 0 | 0 |
| Colony cage loading work rate (h/cage) | 0.2 | 0.2 | 0.2 | 0.2 |
| Colony cage bloodfeeding work rate (h/cage) | 0.05 | 0.05 | 0.05 | 0.05 |
| Egg collection work rate (h/cage) | 0.05 | 0.05 | 0.05 | 0.05 |
| Egg storage work rate (h/day) | 0 | 0 | 0 | 0 |
| Larval diet prep work rate (hour/batch) | 0.15 | 0.15 | 0.15 | 0.15 |
| Adult diet prep work rate (hour/batch) | 0.1 | 0.1 | 0.1 | 0.1 |
| Blood collection weekly | 1 | 1 | 1 | 1 |
| Blood collection work rate (hour/collection) | 5 | 5 | 5 | 5 |
| Blood doses prep work rate (hour/dose) | 0.1 | 0.1 | 0.1 | 0.1 |
| Net working time per staff per day (h/day) | 7.5 | 7.5 | 7.5 | 7.5 |

*(continued)*

| Parameters | Size manual | Size auto | GSS | GSS-CS |
|---|---|---|---|---|
| *Realese facility* | | | | |
| MO cage loading work rate (h/cage) | 0.1 | 0.1 | 0.1 | 0.1 |
| MO cage chilling work rate (h/cage) | 0.1 | 0.1 | 0.1 | 0.1 |
| MO adult collection work rate (hour/cage) | 0.15 | 0.15 | 0.15 | 0.15 |
| MO adult packing for release work rate (hour/cage) | 0.1 | 0.1 | 0.1 | 0.1 |
| *Majoring factors* | | | | |
| Majoring factor labourer | 1.7 | 1.7 | 1.7 | 1.7 |
| Majoring factor team leader | 1.5 | 1.5 | 1.5 | 1.5 |
| Majoring factor QC manager | 1 | 1 | 1 | 1 |
| Majoring factor QC technician | 1 | 1 | 1 | 1 |
| Majoring factor maintenance manager | 1 | 1 | 1 | 1 |
| Majoring factor maintenance officer | 1 | 1 | 1 | 1 |
| Majoring factor admin assistant | 1 | 1 | 1 | 1 |
| Majoring factor manager | 1 | 1 | 1 | 1 |
| QC managers needed mass rearing | 1 | 1 | 1 | 1 |
| Manager needed mass rearing | 1 | 1 | 1 | 1 |
| **Equipment budget** | | | | |
| *Mass rearing facility* | | | | |
| Larval trays unit price ($) | 80 | 80 | 80 | 80 |
| Larval trays life expectancy | 6 | 6 | 6 | 6 |
| Racks for larval trays unit price ($) | 7000 | 7000 | 7000 | 7000 |
| Racks for larval trays life expectancy (years) | 10 | 10 | 10 | 10 |
| Cages for Colonies unit price ($) | 350 | 350 | 350 | 350 |
| Cages for Colonies life expectancy (years) | 10 | 10 | 10 | 10 |
| Irradiator unit price ($) | 250000 | 250000 | 250000 | 250000 |
| Irradiator life expectancy (years) | 10 | 10 | 10 | 10 |
| L1 Sex sorter unit price ($) | 50000 | 50000 | 50000 | 50000 |
| L1 Sex sorter life expectancy (years) | 10 | 10 | 10 | 10 |
| **Pupal sex sorter unit price ($)** | **40000** | **40000** | **40000** | **40000** |
| Pupal sex sorter life expectancy (years) | 10 | 10 | 10 | 10 |
| L1 counter unit price ($) | 2000 | 2000 | 2000 | 2000 |
| L1 counter life expectancy (years) | 5 | 5 | 5 | 5 |
| Larval diet mixer unit price ($) | 3000 | 3000 | 3000 | 3000 |
| Larval diet mixer life expectancy (years) | 5 | 5 | 5 | 5 |
| Adult diet mixer unit price ($) | 2000 | 2000 | 2000 | 2000 |
| Adult diet mixer life expectancy (years) | 6 | 6 | 6 | 6 |
| Larval diet feeder unit price ($) | 3000 | 3000 | 3000 | 3000 |
| Larval diet feeder life expectancy (years) | 5 | 5 | 5 | 5 |
| Blood feeders unit price ($) | 2000 | 2000 | 2000 | 2000 |
| Blood feeders life expectancy (years) | 5 | 5 | 5 | 5 |
| Tray washing machine in mass rearing facility unit price ($) | 20000 | 20000 | 20000 | 20000 |

*(continued)*

| Parameters | Size manual | Size auto | GSS | GSS-CS |
|---|---|---|---|---|
| Tray washing machine in mass rearing facility life expectancy (years) | 6 | 6 | 6 | 6 |
| Cage washing machine in mass rearing facility unit price ($) | 20000 | 20000 | 20000 | 20000 |
| Cage washing machine in mass rearing facility life expectancy (years) | 6 | 6 | 6 | 6 |
| Equipment for basic QC lab number | 1 | 1 | 1 | 1 |
| Equipment for basic QC lab unit price ($) | 12000 | 12000 | 12000 | 12000 |
| Equipment for basic QC lab life expectancy (years) | 5 | 5 | 5 | 5 |
| Workshop equipment number | 1 | 1 | 1 | 1 |
| Workshop equipment unit price ($) | 25000 | 25000 | 25000 | 25000 |
| Workshop equipment life expectancy (years) | 10 | 10 | 10 | 10 |
| Blood storage number | 1 | 1 | 1 | 1 |
| Blood storage unit price ($) | 5000 | 5000 | 5000 | 5000 |
| Blood storage life expectancy (years) | 6 | 6 | 6 | 6 |
| *Realese facility* | | | | |
| Cages for Release Males unit price ($) | 84 | 84 | 84 | 84 |
| Cages for Release Males life expectancy (years) | 4 | 4 | 4 | 4 |
| Cage washing machine in release facility unit price ($) | 20000 | 20000 | 20000 | 20000 |
| Cage washing machine in release facility life expectancy (years) | 6 | 6 | 6 | 6 |
| Equipment for basic QC lab MO number | 1 | 1 | 1 | 1 |
| Equipment for basic QC lab MO unit price ($) | 6000 | 6000 | 6000 | 6000 |
| Equipment for basic QC lab MO life expectancy (years) | 5 | 5 | 5 | 5 |
| Workshop equipment MO number | 1 | 1 | 1 | 1 |
| Workshop equipment MO unit price ($) | 10000 | 10000 | 10000 | 10000 |
| Workshop equipment MO life expectancy (years) | 6 | 6 | 6 | 6 |
| Beef liver powder unit cost ($/kg) | 80 | 80 | 80 | 80 |
| **Diet and consumable costs** | | | | |
| Tuna meal unit cost ($/kg) | 0.7 | 0.7 | 0.7 | 0.7 |
| Brewer yeast unit cost ($/kg) | 8.5 | 8.5 | 8.5 | 8.5 |
| BSF powder unit cost ($/kg) | 8.5 | 8.5 | 8.5 | 8.5 |
| Component 5 unit cost ($/kg) | 0 | 0 | 0 | 0 |
| Component 6 unit cost ($/kg) | 0 | 0 | 0 | 0 |
| Water including the initial load of trays unit cost ($/L) | 3 | 3 | 3 | 3 |
| Sugar for adult colony unit cost ($/kg) | 1 | 1 | 1 | 1 |
| Blood for adult diet unit cost ($/L) | 2 | 2 | 2 | 2 |
| Radiation dosimeters unit cost ($) | 1.5 | 1.5 | 1.5 | 1.5 |
| Consumables without inventoring unit cost ($) | 0.2 | 0.2 | 0.2 | 0.2 |

**Supplementary Data 4: Plasmid and genomic integration nucleotide sequences**

1. pX4 plasmid used for CRISPR/Cas9 knock-in of a Pub-GFP cassette into exon 6 of the *Aedes aegypti* M-linked gene AAEL019619 (Addgene #183903)

2. Nucleotide sequence of three gRNA expression cassettes, each under the control of a different *Aedes aegypti* U6 promoter, used to knock-in the pX4 GFP cassette in the *Ae. aegypti* M locus (Addgene #183912, #183913, #183914)

3. pX3 plasmid used for CRISPR/Cas9 knock-in into the *Aedes aegypti* m-linked gene AAEL019619 (Addgene #183904)

4. ppBAalbNixE1E3E4 PUb-YFP plasmid that produced M-linked *piggyBac* insertion Aalb-M on *Aedes albopictus* chromosome 1 (Addgene #173666)

5. *piggyBac* plasmid containing *Cas9* under control of the *Exuperentia* promoter, used to obtain the m-linked m-albR9 *Ae. albopictus* line (Addgene #183905)

6. pENTRPUb-OptpBTransposase helper plasmid expressing a hyper-active, codon-optimized and NLS-modified piggyBac transposase under the control of the Ae. aegypti Polyubiquitin promoter

7. pattBPUb-GFP, attB site-containg plasmid expressing *GFP* under control of the *Ae. aegypti* PUb promoter, inserted in the m-linked attP site of *Ae. albopictus* strain mX1 derived from m-albR9 (Addgene #183911)

8. Helper plasmid (*piggyBac* construct) used to express PhiC31 integrase under control of the PUb promoter (Addgene #183966)

9. Genomic sequence flanking the Aalb-M insertion on the 3' piggyBac side.

10. Genomic sequence flanking the Aalb-m insertion on the 3' piggyBac side.

**1. pX4 plasmid used for CRISPR/Cas9 knock-in of a Pub-GFP cassette into exon 6 of the Aedes aegypti M-linked gene AAEL019619**

```
LOCUS       pX4                6785 bp    DNA     circular       19-MAY-2021
FEATURES             Location/Qualifiers
     misc_feature    complement(603..727)
                     /note="attR4"
     misc_feature    complement(5090..5108)
                     /note="pDONR-RP"
     misc_feature    537..552
                     /note="M13F"
     misc_feature    complement(5046..5064)
                     /note="M13R"
     exon            1666..1738
                     /note="exon_id=AAEL019619-RC-E5.1"
     exon            1666..1738
                     /note="exon_id=AAEL019619-RA-E6.1"
     misc_feature    complement(1737..1738)
                     /note="gRNA1 X4 "
     CDS             1666..1687
                     /gene="AAEL019619.1"
                     /protein_id="AAEL019619-PC"
                     /note="transcript_id=AAEL019619-RC"
                     /db_xref="RefSeq_peptide:XP_021708495.1"
                     /db_xref="RefSeq_peptide:XP_021708495.1"
                     /db_xref="RefSeq_dna:XM_021852803.1"
                     /db_xref="Uniprot/SPTREMBL:A0A1S4FXT0"
                     /db_xref="Uniprot/SPTREMBL:Q16KX0"
                     /db_xref="GO:0005856"
                     /db_xref="GO:0030036"
                     /db_xref="UniParc:UPI000B789663"
                     /translation="MPFVQRVVTPKYVARSTKPSHSRGTAALPVQDYELEAITNLTLS
                     NALRQLASLVLISNQIFTELNKELASVSERSLGIKQRIDNLSKRVEEFDPKQVAVPES
                     DLVAFSQIKNHYSTKYHIDTCLFTAETRTETLQELYDAAAKTPVSAIAEMDRIAGYNE
                     SDHLGSDAFLCTPVLGQTRRKLRAKVDMDIETRLPSAVEDLRKWTSIEAIGDTTVPPD
                     CTVRLTGNGSVILGNVSQNASPSYQAGTSASFDETDDIIMVDSSSRGLGREQDTPLDH
                     RLPSPEEQCQMIALKFPAETIKVDTSGKRFDRMCATRKSLLHFVSTSEQAETNAAGGQ
                     QADGSDGDTIRRRSRPRRSRGKRRNTIAGTDQKEIAEVVNNGLSTFVP"
                     /dnas_title="AAEL019619.1"
     misc_feature    778..1738
                     /note="X4 5%82%C4%F4flk region"
     misc_feature    1922..2637
                     /note="GFP"
     misc_feature    1739..1740
                     /note="perfect intron splice acceptor site!"
     misc_feature    complement(1753..1921)
                     /note="Dm beta-Tub56D terminator"
     misc_feature    complement(1808..1813)
                     /note="polyA site"
     misc_feature    1739..1752
                     /note="PRESERVED"
```

misc_feature 1739..1920
/note="gBlock"
misc_feature complement(2643..3998)
/note="PUb promoter"
misc_feature 4024..4053
/note="intron"
misc_feature 4063..4081
/note="gRNA2 X4"
variation 4464..4464
/replace="A/C"
/db_xref="VBP0000121:supercont1.93:2165054"
exon 4054..4137
/note="exon_id=AAEL019619-RC-E5.1"
misc_feature complement(4054..4070)
/note="gRNA1 X4 "
mRNA 4054..4137
/gene="AAEL019619.1"
/standard_name="AAEL019619-RC"
/dnas_title="AAEL019619.1"
variation 4720..4720
/replace="G/A"
/db_xref="VBP0000177:AX-93227275"
misc_feature 4054..5018
/note="X4 5%82%C4%F4 flk"
source 1..6785
/dnas_title="knockin plasmid X4"

ORIGIN
1 CTTTCCTGCG TTATCCCCTG ATTCTGTGGA TAACCGTATT ACCGCCTTTG AGTGAGCTGA
61 TACCGCTCGC CGCAGCCGAA CGACCGAGCG CAGCGAGTCA GTGAGCGAGG AAGCGGAAGA
121 GCGCCCAATA CGCAAACCGC CTCTCCCCGC GCGTTGGCCG ATTCATTAAT GCAGCTGGCA
181 CGACAGGTTT CCCGACTGGA AAGCGGGCAG TGAGCGCAAC GCAATTAATA CGCGTACCGC
241 TAGCCAGGAA GAGTTTGTAG AAACGCAAAA AGGCCATCCG TCAGGATGGC CTTCTGCTTA
301 GTTTGATGCC TGGCAGTTTA TGGCGGGCGT CCTGCCCGCC ACCCTCCGGG CCGTTGCTTC
361 ACAACGTTCA AATCCGCTCC CGGCGGATTT GTCCTACTCA GGAGAGCGTT CACCGACAAA
421 CAACAGATAA AACGAAAGGC CCAGTCTTCC GACTGAGCCT TTCGTTTTAT TTGATGCCTG
481 GCAGTTCCCT ACTCTCGCGT TAACGCTAGC ATGGATGTTT TCCCAGTCAC GACGTTGTAA
541 AACGACGGCC AGTCTTAAGC TCGGGCCCCT ACAGGTCACT AATACCATCT AAGTAGTTGA
601 TTCATAGTGA CTGGATATGT TGTGTTTTAC AGTATTATGT AGTCTGTTTT TTATGCAAAA
661 TCTAATTTAA TATATTGATA TTTATATCAT TTTACGTTTC TCGTTCAACT TTTCTATACA
721 AAGTTGGTAC CGGGCCCCCC GCTAGCGTCG ACGGTATCGA TAAGCTTGAT cggatccGCT
781 CCGTTGATTG ATGAATCATT GATTCAGTAA CTTAGTTGAT CAATAACTCT AACACGAAAA
841 TTACGGATAT AATTTCGTAA TTTCGAATTT TGAGGGTTTT GTTAGATGAA CTTTGATTTA
901 AGATCTTGTT TCGTAGGAAG ATAAATGTCA CAATCATTGT TTTGCAATAC CCTATTATTG
961 ATATAAAAAA TCTGAAATTT TTTTAAATTT AGTTTATAGG CACTCGTTGT TTACGAAATA
1021 TGATGCAAAA TTCAGTTTAT TATAAAAAAG TTTTTTaTTT TTCGAATTAG GGTAGCATCC
1081 CAAATTATGT CACGCTAAAT TTCAACTTTT TCGACCCCCT TCCCTCCTCT ACTTTGTCAC
1141 ATTTTTTGTA TGAGTCCTCC GAAAATTTTG TAAGGCTTGA CCCCCTCCAC CCCTCTTAAA
1201 GCGTGACGTA ATTTGTGCAT GACCCCTTAG ATAGAAATGT TGAAAACCTA TCAACATAAA
1261 TCTGGGCAAA ACTTCAATCA AAATAAATGC AATGTACACT GAACAGGTAA GTGTAGGAGG
1321 TAAGAAAAGA TAGCTAAAAA AAACTCGACT CGACTTGATG ATGGTTTTGA AAAAACTCAA
1381 AAACGACATG TAACAATAAA GGTAAGTATT GAATGGATAC TTGTATACAA AGTACTTTGA

1441 AGTTCAATTC TTGTTGAAAT TGAACTGTCT TCCATAACAT TTATTCTTAA ACACTAAATT
1501 TCTAAGTTTT AACGGCATTT CGTGGTGCGC ACTGAACTTG TTCTCTTTTC TCCTGATACA
1561 TAACTCACAT GTAGCTCTGA TGTTGAAACA CTTTCATATC TATCATTAAG TCAGCAATCT
1621 CAGGACAATT TTCATCATAT TTTTTCTCTT GTATTTTTTC AACAGGTTAT CTACATTCGT
1681 CCCCTGATGA GTACGTTCAA TATTATGGAC CACAAACTCA CACATACACG CTTCCCAGGT
1741 AaGTTCCTAT ACgaaacccc aacaaaaacc ataattgttt AGACTTGTGA ACAAAATTGG
1801 ATCCGACTTT ATTGATTACG TTGTTAAGAG AACAAATCTT TTACAACTGA ATTCATTTGT
1861 TCTCGTTTCA TTTTTTTTCG CAAAACATTG ATCGAGAATT CGATTGATTT CCGATTCGAA
1921 TTTACTTGTA CAGCTCGTCC ATGCCGAGAG TGATCCCGGC GGCGGTCACG AACTCCAGCA
1981 GGACCATGTG ATCGCGCTTC TCGTTGGGGT CTTTGCTCAG GGCGGACTGG GTGCTCAGGT
2041 AGTGGTTGTC GGGCAGCAGC ACGGGGCCGT CGCCGATGGG GGTGTTCTGC TGGTAGTGGT
2101 CGGCGAGCTG CACGCTGCCG TCCTCGATGT TGTGGCGGAT CTTGAAGTTC ACCTTGATGC
2161 CGTTCTTCTG CTTGTCGGCC ATGATATAGA CGTTGTGGCT GTTGTAGTTG TACTCCAGCT
2221 TGTGCCCCAG GATGTTGCCG TCCTCCTTGA AGTCGATGCC CTTCAGCTCG ATGCGGTTCA
2281 CCAGGGTGTC GCCCTCGAAC TTCACCTCGG CGCGGGTCTT GTAGTTGCCG TCGTCCTTGA
2341 AGAAGATGGT GCGCTCCTGG ACGTAGCCTT CGGGCATGGC GGACTTGAAG AAGTCGTGCT
2401 GCTTCATGTG GTCGGGGTAG CGGCTGAAGC ACTGCACGCC GTAGGTCAGG GTGGTCACGA
2461 GGGTGGGCCA GGGCACGGGC AGCTTGCCGG TGGTGCAGAT GAACTTCAGG GTCAGCTTGC
2521 CGTAGGTGGC ATCGCCCTCG CCCTCGCCGG ACACGCTGAA CTTGTGGCCG TTTACGTCGC
2581 CGTCCAGCTC GACCAGGATG GGCACCACCC CGGTGAACAG CTCCTCGCCC TTGCTCAcca
2641 tgGTTGAAAT CTCTGTTGAG CAGAAAAAGA AACGAGGAAA CGCTTCAGTA ATTGGTTGTG
2701 AAATGCAAAC TCTCATTTGA TATTGATTCA TTGCCTTTGG CTTCGAGCAC GACACGACAG
2761 GTTTTAAACT TGTTTTGCTT TGTCTGCGTT TGCAGTCGCA GGCCAAGTGA AAAATATACA
2821 CTTGAAGGTG ATGACGTCAC AACAACGCCC TACTTTTTAG TGAAAAATTA ACTTGTTTTC
2881 GACTTTGAAC TACGTAGTTT GGAAATTGCG TATCTTCAAG TTTTCACGAT TTCCTCAAGG
2941 TTTTTCTCGA TATGTGTTAA TATTACCTTA ATGGGTAATT ATCACCACAA TATTTAATTT
3001 TAGAAATATG GCGAATTCGC GCTTGCTCTA GAAATTTAAT TCAATTAGCA GCGATGATTC
3061 AACGAAATAT GATTATCGCC GTGAATCACA ATTGGGTTTT ATCAATGATG ATGAAACTGC
3121 GTTGCAAATT TTCACTAATC ACTCAAAGCT CAATAGTCGC CATCTTGGAA AATAGTTTGC
3181 TCATTCAAAG ACAAAGGAAT CATTCACCAA ACTAGTTTTC GCTCATAGCT ATAATTTCAT
3241 CATTTAATTT ACCTACCTTC ACTAGAAGAT TCCCTTCACC GAAATGCACT TTCACAAATT
3301 TAATAGTAAT TGTCCTTTGA ACAAAGCTTT GTTCACTCTG AAATTTTCTC CTCTGGCTAA
3361 TTGGATCACT CTTTTTCACT AGAGACTTCA CTTCACTTGC ACTGGCACTG CTTACTTGGA
3421 CCGCGTAATG TTCACTCCAC TAGGAAACGT ATTCGATTGA GCTGGTTTTG CCTTTGCAGG
3481 GGCGTTTTAT AGACACTGCC GTAGTGGTGG TTGTACTTCT AGAAAATTTC TGCAATCCAT
3541 TCATACATAT ACCTCTGTTC GGTTGGATGG CTCTAAATCG ATCGACTATG GGTACCATCC
3601 CTGTGTCTGA ATGGACAGCA AAACGTGCTT GTGTCTGTTA GTCGTTCATT CTGTACCTTG
3661 AACGATGCAG TTCAACTTCT GGCAAAGACG TCAATGTACC TACCATTCGT GTATATGGCA
3721 TACAGAGAAA GATGTGCGAA TGCCTTTTTC GATAGAGAAA GGATTTCCTG ATTTGATCGA
3781 CAATTTCGGT AGCTATTCTT AGCTTCGAAT GTAATCTGAT AACCAAATCC AGAGACAAAG
3841 CAGTAATTTG GGAATTTCAC TTGAATCATT CTAATTGTAT CATTTGCTGT ATTAGTGCAG
3901 TCAGCAAAGT GACGTCAACC CTTCTAAATC GATATACTTC TGGGAAGCTT TCTTTCTTGT
3961 CTGGCTCAGC TGGTGCCAAG GCAAATTATA ATTGGATTCA ATGCACAAGC TACATGTAAA
4021 GATCTTTTCT ACGCCTACCC CCTCGGACAA CAGGCGtCCC ATGATTCATC AATCAACGGA
4081 GCGaCCTCAT AGAAATATTA CGCCAACTGG AATACCAATG GCATCCAGAC ATATCTATGG
4141 GTATGACGCC CAAGTTGACA ATAATGCAAG CACATTCCAC CGGAACAACG ATATGTACTG
4201 GACCTTGCAA ACTCGTCGTC GACCAATTCC AATGGACCAG CCGAATATGC GTGAAGCTCT
4261 TAATTTATCC CAGCAGAGCG TTAGACCAAT AGAAGAACTC TATTCAACAC CAAACAAGAT
4321 CAAATCAAAT GCACTGAAAA CTACACCAAC TGGGTTTGCT ACGTACTTTT CTCCTATCGC
4381 TAAAAGTAAC ACGGGTGGTG AATTAAACGA CGATGTGGAT AGCAGTTTTA AGATTTCTCC
4441 GATCGACCCT CGGAGTATTG CTCATTTCAA AACATCTACT CCGTCAAAAC CAAACGAGAC

4501 ATCACTTTCT CCTACAAGAT CGCTTACCGA AGAGCTTCGA TATAGATTAA GACTGCAACA
4561 GGTCGGAGTT CGATCACATG GGAATTCTCC GGTAAGCTCC GGCCGTTCAA CACCAAAAAA
4621 CGTACTTGAA GCACAAGCTC CACGTGGCAG ACACAGTTGG GCTTCCAACA GTACCGACAT
4681 TCCACAGACA TGCTCTGATC GACTTGGGAC ACCAAAAACG AGCCTTATGG ATTTCAAAAA
4741 GTTGCTTCTT GCGCATGGAT CGAAATCTAA TATTTCGTCT GGAAGTAAAA TTTCTGCCGT
4801 TGAACTACTT AAGAAATCAA AAAGTAACGC TCCAGTTAAC CCAGTTTCTC CAGTTACAAA
4861 AACTTCTGCA AATAGTAGTT TGAATATTTT GGACCTTTCG GGATCTCCAA AAACATTCGC
4921 TACAAGACGC ATGATTCGAC AAGGAAACTT TGGAAATGGT TCACCATCAA AGCTCGGTAA
4981 TGTGTCTAAA CATTCTTCAA GGGGTGGTTG GCGGTACAgc ttATCCCCTA TAGTGAGTCG
5041 TATTACATGG TCATAGCTGT TTCCTGGCAG CTCTGGCCCG TGTCTCAAAA TCTCTGATGT
5101 TACATTGCAC AAGATAAAAA TATATCATCA TGAACAATAA AACTGTCTGC TTACATAAAC
5161 AGTAATACAA GGGGTGTTAT GAGCCATATT CAACGGGAAA CGTCGAGGCC GCGATTAAAT
5221 TCCAACATGG ATGCTGATTT ATATGGGTAT AAATGGGCTC GCGATAATGT CGGGCAATCA
5281 GGTGCGACAA TCTATCGCTT GTATGGGAAG CCCGATGCGC CAGAGTTGTT TCTGAAACAT
5341 GGCAAAGGTA GCGTTGCCAA TGATGTTACA GATGAGATGG TCAGACTAAA CTGGCTGACG
5401 GAATTTATGC CTCTTCCGAC CATCAAGCAT TTTATCCGTA CTCCTGATGA TGCATGGTTA
5461 CTCACCACTG CGATCCCCGG AAAAACAGCA TTCCAGGTAT TAGAAGAATA TCCTGATTCA
5521 GGTGAAAATA TTGTTGATGC GCTGGCAGTG TTCCTGCGCC GGTTGCATTC GATTCCTGTT
5581 TGTAATTGTC CTTTTAACAG CGATCGCGTA TTTCGTCTCG CTCAGGCGCA ATCACGAATG
5641 AATAACGGTT TGGTTGATGC GAGTGATTTT GATGACGAGC GTAATGGCTG GCCTGTTGAA
5701 CAAGTCTGGA AAGAAATGCA TAAACTTTTG CCATTCTCAC CGGATTCAGT CGTCACTCAT
5761 GGTGATTTCT CACTTGATAA CCTTATTTTT GACGAGGGGA AATTAATAGG TTGTATTGAT
5821 GTTGGACGAG TCGGAATCGC AGACCGATAC CAGGATCTTG CCATCCTATG GAACTGCCTC
5881 GGTGAGTTTT CTCCTTCATT ACAGAAACGG CTTTTTCAAA AATATGGTAT TGATAATCCT
5941 GATATGAATA AATTGCAGTT TCATTTGATG CTCGATGAGT TTTTCTAATC AGAATTGGTT
6001 AATTGGTTGT AACACTGGCA GAGCATTACG CTGACTTGAC GGGACGGCGC AAGCTCATGA
6061 CCAAAATCCC TTAACGTGAG TTACGCGTCG TTCCACTGAG CGTCAGACCC CGTAGAAAAG
6121 ATCAAAGGAT CTTCTTGAGA TCCTTTTTTT CTGCGCGTAA TCTGCTGCTT GCAAACAAAA
6181 AAACCACCGC TACCAGCGGT GGTTTGTTTG CCGGATCAAG AGCTACCAAC TCTTTTTCCG
6241 AAGGTAACTG GCTTCAGCAG AGCGCAGATA CCAAATACTG TTCTTCTAGT GTAGCCGTAG
6301 TTAGGCCACC ACTTCAAGAA CTCTGTAGCA CCGCCTACAT ACCTCGCTCT GCTAATCCTG
6361 TTACCAGTGG CTGCTGCCAG TGGCGATAAG TCGTGTCTTA CCGGGTTGGA CTCAAGACGA
6421 TAGTTACCGG ATAAGGCGCA GCGGTCGGGC TGAACGGGGG GTTCGTGCAC ACAGCCCAGC
6481 TTGGAGCGAA CGACCTACAC CGAACTGAGA TACCTACAGC GTGAGCTATG AGAAAGCGCC
6541 ACGCTTCCCG AAGGGAGAAA GGCGGACAGG TATCCGGTAA GCGGCAGGGT CGGAACAGGA
6601 GAGCGCACGA GGGAGCTTCC AGGGGGAAAC GCCTGGTATC TTTATAGTCC TGTCGGGTTT
6661 CGCCACCTCT GACTTGAGCG TCGATTTTTG TGATGCTCGT CAGGGGGGCG GAGCCTATGG
6721 AAAAACGCCA GCAACGCGGC CTTTTTACGG TTCCTGGCCT TTTGCTGGCC TTTTGCTCAC
6781 ATGTT
//

**2. Nucleotide sequence of three gRNA expression cassettes, each under the control of a different *Aedes aegypti* U6 promoter, used to knock-in the pX4 GFP cassette in the *Ae. aegypti* M locus**

LOCUS Plasmid1243 632 bp ds-DNA linear SYN 13-OCT-2021
DEFINITION synthetic DNA fragment
ORGANISM synthetic DNA construct
REFERENCE 1 (bases 1 to 632)
FEATURES Location/Qualifiers
source 1..632
/organism="synthetic DNA construct"
/mol_type="other DNA"
misc_feature 8..481
/label=promoter of Chr1 U6 AAEL017702
/note="promoter of Chr1 U6 AAEL017702"
misc_feature 421..433
/label=Conserved atccatcgctaga
misc_feature 455..460
/label=TATA
misc_feature 482..504

misc_feature 505..590
/label=pX330 chimeric guide RNA scaffold
/note="pX330 chimeric guide RNA scaffold"
misc_feature 505..515
/label=CRISPR end
/note="CRISPR end"
misc_feature 510..540
/label=mutations to optimize gRNA according to
/note="mutations to optimize gRNA according to Dang et al."
misc_feature 525..590
/label=tracer
/note="tracer"
misc_feature complement(574..590)
/label=EM334
/note="EM334"
misc_feature 591..595
/label=U6 Aedes - AAEL017702
ORIGIN
1 ggtctcattc catagtaaat atactacaaa taatattaat gttccatgaa aaaaggagta
61 agagtctggt aaccctagtg cacgcaaata tctcgcgggc atatttggtt gctgaggtat
121 atttatattt gaacgccatg agaaaaagcg gaagaaattg gctcatggcc gattttaagg
181 atatttaaaa attgtacaat gtacatataa ttaaacatcc gttccttcaa tgtgttcttt
241 tttttaagcg tgtgttaaaa gtttgctctg gtggtgaatt cacgctctac ccgttcaggc
301 agcattcatc gaaaagccct atctgctcgc acacatttac aaaatgctga ttgcgttgtg
361 tgctgaatgg gtcactcgtc cgtcactgct tgctgtgtac actgtacagt tacgcagtct
421 gtgcatcgct agaatcatat ttacggaaaa gtattatata tacccaatgc gttgctcatc
481 ggttgtggca tagcgccgtg tggagtttca gagctatgct ggaaacagca tagcaagttg
541 aaataaggct agtccgttat caacttgaaa aagtggcacc gagtcggtgc ttttttgtg
601 gttttattat tcgataattg tggatagaga cc
//

LOCUS       Plasmid1244                520 bp ds-DNA     linear SYN 13-OCT-2021
DEFINITION  synthetic DNA fragment
SOURCE      synthetic DNA construct
  ORGANISM  synthetic DNA construct
REFERENCE   1  (bases 1 to 520)
FEATURES             Location/Qualifiers
     source          1..520
                     /organism="synthetic DNA construct"
                     /mol_type="other DNA"
     misc_feature    8..328
                     /label=pU6 promoter Chr3 AAEL017905 Konet2007
                     /note="pU6 promoter Chr3 AAEL017905 Konet2007"
     misc_feature    288..301
                     /label=conserved
     misc_feature    322..327
                     /label=TATA
     misc_feature    329..350
                     /label=this section from AAEL017774 U6
                     /note="this section from AAEL017774 U6"
     misc_feature    351..372
                     /label=Linker
     misc_feature    373..458
                     /label=pX330 chimeric guide RNA scaffold
                     /note="pX330 chimeric guide RNA scaffold"
     misc_feature    373..383
                     /label=CRISPR  end
                     /note="CRISPR  end"
     misc_feature    378..408
                     /label=mutations to optimize gRNA according to
                     /note="mutations to optimize gRNA according to Dang et al."
     misc_feature    393..458
                     /label=tracer
                     /note="tracer"
     misc_RNA        459..463
                     /label=AAEL017905-RA
                     /note="snRNA"
ORIGIN
        1 ggtctcactc taattggagc tgccagacaa atttgatttg tccgtgcggt atgcatatgt
       61 acctacttac gtctgtcgtt ttgtctccgt ttaccgggag ggaaagtctg gaaacatgga
      121 aactctatag ttgccaggta gaccatctgc ctccgtcggc tggctggatt ccaatttgaa
      181 tattggctaa ttggaagaga tggaagtttt tgaatggatg attgaataat tgaagcgact
      241 ccgggtacct gtttgtaagc tctgcaacag tgccatagat tcgtgtcagt ccatcactag
      301 aatcaaatca acttgtactt gatatataag agcagaggca agagtagtga aatgtcttaa
      361 atgaaagagg cggtttcaga gctatgctgg aaacagcata gcaagttgaa ataaggctag
      421 tccgttatca acttgaaaaa gtggcaccga gtcggtgctt tttttacaaa tctattatgc
      481 atgaggtact atgttggcag agtaatgaat tccagagacc
//

LOCUS       Plasmid1245                530 bp ds-DNA     linear SYN 13-OCT-2021
DEFINITION  synthetic DNA fragment

```
SOURCE      synthetic DNA construct
ORGANISM  synthetic DNA construct
REFERENCE   1  (bases 1 to 530)
FEATURES             Location/Qualifiers
     source          1..530
                     /organism="synthetic DNA construct"
                     /mol_type="other DNA"
     misc_feature    8..364
                     /label=U6 prom from Chr3 AAEL017763
                     /note="U6 prom from Chr3 AAEL017763"
     misc_feature    304..317
                     /label=Conserved
     misc_feature    337..342
                     /label=TATA
     misc_feature    365..386
                     /label=Linker
     misc_feature    387..472
                     /label=pX330 chimeric guide RNA scaffold
                     /note="pX330 chimeric guide RNA scaffold"
     misc_feature    387..397
                     /label=CRISPR  end
                     /note="CRISPR  end"
     misc_feature    392..422
                     /label=mutations to optimize gRNA according to
                     /note="mutations to optimize gRNA according to Dang et al."
     misc_feature    407..472
                     /label=tracer
                     /note="tracer"
     misc_RNA        473..477
                     /label=U6 AAEL017763-RA
                     /note="snRNA"
ORIGIN
        1 ggtctcatgt tcgttcttca acacctctcc atggtgataa cggatacggt ttcattgtca
       61 gcatccatcc tccgaaaaat acattacgcc ttgaaatatg caatcgcaaa cacggatctg
      121 tttggaacat ttattttact atgaagagat gcgataggta atatttattt gagcgtttaa
      181 gatactcatt gttctctcaa agaatgtcat tgaaagccaa cgaggtcaaa tcaaatatta
      241 taataaaaag gtcaaagagg actaacttaa agctctcttt atggatagga aaaaatattt
      301 tcgcccatcg ctagaacttt taccgtttcc attgagtata taactaagat gaatgaggct
      361 aattgatgta tcatgcgtat tgcgaggttt cagagctatg ctggaaacag catagcaagt
      421 tgaaataagg ctagtccgtt atcaacttga aaaagtggca ccgagtcggt gctttttttaa
      481 gatgcttcgg caattaataa taaggaatgg taaagcattc tctagagacc
//
```

**3. pX3 plasmid used for CRISPR/Cas9 knock-in into the Aedes aegypti m-linked gene AAEL019619**

LOCUS       pX3                    6628 bp ds-DNA     circular SYN 20-NOV-2019
DEFINITION  synthetic circular DNA
FEATURES             Location/Qualifiers
    terminator      268..295
                    /label=rrnB T2 terminator
                    /note="transcription terminator T2 from the E. coli rrnB
                    gene"
    terminator      387..473
                    /gene="Escherichia coli rrnB"
                    /label=rrnB T1 terminator
                    /note="transcription terminator T1 from the E. coli rrnB
                    gene"
    primer_bind     537..553
                    /label=M13 fwd
                    /note="common sequencing primer, one of multiple similar
                    variants"
    misc_feature    774..1707
                    /label=X3 5' flk region
    misc_feature    complement(1721..1889)
                    /label=DmTub56Dterm
    CDS             complement(1890..2609)
                    /codon_start=1
                    /product="the original enhanced GFP (Yang et al., 1996)"
                    /label=EGFP
                    /note="mammalian codon-optimized"
                    /translation="MVSKGEELFTGVVPILVELDGDVNGHKFSVSGEGEGDATYGKLTL
                    KFICTTGKLPVPWPTLVTTLTYGVQCFSRYPDHMKQHDFFKSAMPEGYVQERTIFFKDD
                    GNYKTRAEVKFEGDTLVNRIELKGIDFKEDGNILGHKLEYNYNSHNVYIMADKQKNGIK
                    VNFKIRHNIEDGSVQLADHYQQNTPIGDGPVLLPDNHYLSTQSALSKDPNEKRDHMVLL
                    EFVTAAGITLGMDELYK"
    misc_feature    complement(2610..3986)
                    /label=AePUb promoter
    misc_feature    3987..4861
                    /label=X3 3' flk region
    promoter        complement(4870..4888)
                    /label=T7 promoter
                    /note="promoter for bacteriophage T7 RNA polymerase"
    primer_bind     complement(4893..4909)
                    /label=M13 rev
                    /note="common sequencing primer, one of multiple similar
                    variants"
    CDS             5022..5831
                    /codon_start=1
                    /gene="aph(3')-Ia"
                    /product="aminoglycoside phosphotransferase"
                    /label=KanR
                    /note="confers resistance to kanamycin in bacteria or G418
                    (Geneticin(R)) in eukaryotes"
                    /translation="MSHIQRETSRPRLNSNMDADLYGYKWARDNVGQSGATIYRLYGKP

DAPELFLKHGKGSVANDVTDEMVRLNWLTEFMPLPTIKHFIRTPDDAWLLTTAIPGKTA
FQVLEEYPDSGENIVDALAVFLRRLHSIPVCNCPFNSDRVFRLAQAQSRMNNGLVDASD
FDDERNGWPVEQVWKEMHKLLPFSPDSVVTHGDFSLDNLIFDEGKLIGCIDVGRVGIAD
RYQDLAILWNCLGEFSPSLQKRLFQKYGIDNPDMNKLQFHLMLDEFF"

rep_origin 5978..6566
/direction=RIGHT
/label=ori
/note="high-copy-number ColE1/pMB1/pBR322/pUC origin of replication"

ORIGIN
1 ctttcctgcg ttatcccctg attctgtgga taaccgtatt accgcctttg agtgagctga
61 taccgctcgc cgcagccgaa cgaccgagcg cagcgagtca gtgagcgagg aagcggaaga
121 gcgcccaata cgcaaaccgc ctctccccgc gcgttggccg attcattaat gcagctggca
181 cgacaggttt cccgactgga aagcgggcag tgagcgcaac gcaattaata cgcgtaccgc
241 tagccaggaa gagtttgtag aaacgcaaaa aggccatccg tcaggatggc cttctgctta
301 gtttgatgcc tggcagttta tggcgggcgt cctgcccgcc accctccggg ccgttgcttc
361 acaacgttca aatccgctcc cggcggattt gtcctactca ggagagcgtt caccgacaaa
421 caacagataa aacgaaaggc ccagtcttcc gactgagcct ttcgttttat ttgatgcctg
481 gcagttccct actctcgcgt taacgctagc atggatgttt tcccagtcac gacgttgtaa
541 aacgacggcc agtcttaagc tcgggcccct acaggtcact aataccatct aagtagttga
601 ttcatagtga ctggatatgt tgtgttttac agtattatgt agtctgtttt ttatgcaaaa
661 tctaatttaa tatattgata tttatatcat tttacgtttc tcgttcaact tttctataca
721 aagttggtac cgggcccccc gctagcgtcg acggtatcga taagcttgat cggatccgga
781 gagtacgttc gcattgccga agtaaaaact accaagaaaa ttcaacacag cggtggcgtt
841 tactattcaa acgatgtggt aaaacgtaat ttagccacag tcctctcggg aaatctcaac
901 gaagaaaccg aatatgtatc cttgaatgaa ttgccgtgca atatgcgatg cgagagcaat
961 gttctctcgc aaaacgacac ccctcctggc ggaagcggaa aactgaatac ttcgggtacg
1021 ggaggagaag caccctcggc aggagaaaac aacgaacatg ctataaaaag ggtgcacgc
1081 gtgaaactgg atgcgcacgg taaggttatc tacagttcgg atagtctgaa gaggagaaag
1141 ggtgcgcaca cgacctttgc gccgggtcca tttgtgaaag acgtaaatac ggcacagcca
1201 ctgacgacca acactcctac tacaaacact acaactcggt cagcatcacc agcagcatcc
1261 acagaaaatg gcaaatgtgc tgttgttgca tctccattac taagtggtgg tctagctaac
1321 aggaaaccaa tagcagttaa accagtgata tcgcagtccg cattgaaaac gaaagcaatc
1381 gtcaacacca atagtaatat aagtaatcat agcgacggag tggcagccca tcgtatggcc
1441 ccaattgcca gtattcttcc caacaacagt acggtgcaag aagtggaaaa aatagctgca
1501 gcatcacaca tcatgtcctc cggagttcta aaaggtgcat acgttaatgt acaggaatct
1561 aaaccaccgt cttcaaaacc tcaatcgatt cactatatgg aagcaacaaa cttacagcat
1621 ccccaaacag gtaagcaagc taaatcgcaa aagaataaca aaatagttgg tgcttcgcca
1681 aaggctagga acgtaatatt tccctccggg gttcctatac gaaaccccaa caaaaaccat
1741 aattgtttag acttgtgaac aaaattggat ccgactttat tgattacgtt gttaagagaa
1801 caaatctttt acaactgaat tcatttgttc tcgtttcatt ttttttcgca aaacattgat
1861 cgagaattcg attgatttcc gattcgaatt tacttgtaca gctcgtccat gccgagagtg
1921 atcccggcgg cggtcacgaa ctccagcagg accatgtgat cgcgcttctc gttggggtct
1981 ttgctcaggg cggactgggt gctcaggtag tggttgtcgg gcagcagcac ggggccgtcg
2041 ccgatggggg tgttctgctg gtagtggtcg gcgagctgca cgctgccgtc ctcgatgttg
2101 tggcggatct tgaagttcac cttgatgccg ttcttctgct tgtcggccat gatatagacg
2161 ttgtggctgt tgtagttgta ctccagcttg tgccccagga tgttgccgtc ctccttgaag
2221 tcgatgccct tcagctcgat gcggttcacc agggtgtcgc cctcgaactt cacctcggcg
2281 cgggtcttgt agttgccgtc gtccttgaag aagatggtgc gctcctggac gtagccttcg
2341 ggcatggcgg acttgaagaa gtcgtgctgc ttcatgtggt cggggtagcg gctgaagcac
2401 tgcacgccgt aggtcagggt ggtcacgagg gtgggccagg gcacgggcag cttgccggtg

2461 gtgcagatga acttcagggt cagcttgccg taggtggcat cgccctcgcc ctcgccggac
2521 acgctgaact tgtggccgtt tacgtcgccg tccagctcga ccaggatggg caccaccccg
2581 gtgaacagct cctcgccctt gctcaccatg gttgaaatct ctgttgagca gaaaaagaaa
2641 cgaggaaacg cttcagtaat tggttgtgaa atgcaaactc tcatttgata ttgattcatt
2701 gcctttggct tcgagcacga cacgacaggt tttaaacttg ttttgctttg tctgcgtttg
2761 cagtcgcagg ccaagtgaaa aatatacact tgaaggtgat gacgtcacaa caacgcccta
2821 ctttttagtg aaaaattaac ttgttttcga ctttgaacta cgtagtttgg aaattgcgta
2881 tcttcaagtt ttcacgattt cctcaaggtt tttctcgata tgtgttaata ttaccttaat
2941 gggtaattat caccacaata tttaatttta gaaatatggc gaattcgcgc ttgctctaga
3001 aatttaattc aattagcagc gatgattcaa cgaaatatga ttatcgccgt gaatcacaat
3061 tgggttttat caatgatgat gaaactgcgt tgcaaatttt cactaatcac tcaaagctca
3121 atagtcgcca tcttggaaaa tagtttgctc attcaaagac aaaggaatca ttcaccaaac
3181 tagttttcgc tcatagctat aatttcatca tttaatttac ctaccttcac tagaagattc
3241 ccttcaccga aatgcacttt cacaaattta atagtaattg tcctttgaac aaagctttgt
3301 tcactctgaa attttctcct ctggctaatt ggatcactct ttttcactag agacttcact
3361 tcacttgcac tggcactgct tacttggacc gcgtaatgtt cactccacta ggaaacgtat
3421 tcgattgagc tggttttgcc tttgcagggg cgttttatag acactgccgt agtggtggtt
3481 gtacttctag aaaatttctg caatccattc atacatatac ctctgttcgg ttggatggct
3541 ctaaatcgat cgactatggg taccatccct gtgtctgaat ggacagcaaa acgtgcttgt
3601 gtctgttagt cgttcattct gtaccttgaa cgatgcagtt caacttctgg caaagacgtc
3661 aatgtaccta ccattcgtgt atatggcata cagagaaaga tgtgcgaatg cctttttcga
3721 tagagaaagg atttcctgat ttgatcgaca atttcggtag ctattcttag cttcgaatgt
3781 aatctgataa ccaaatccag agacaaagca gtaatttggg aatttcactt gaatcattct
3841 aattgtatca tttgctgtat tagtgcagtc agcaaagtga cgtcaaccct tctaaatcga
3901 tatacttctg ggaagctttc tttcttgtct ggctcagctg gtgccaaggc aaattataat
3961 tggattcaat gcacaagcta catgtaaaga cgcatgatac tcatcacatt agacgtaata
4021 aggacattga cccagatagc aattctacta tcaagcgcag gaatagctat agaaatgcga
4081 actcgttcaa aatcgatact gatgctgcga gttttaccga tttcttggaa gaacatgtaa
4141 ataaacaaga ggaacggcta cagtcctccg cggtgaagaa tgatagcaat tcatataaca
4201 agtataccca tttgcatgat caaaatagcg ataaggacga tggctacgaa gaaatcagcg
4261 atattgactt ggacgaggat agcaaatcta ttgctcgttc gtttgaaaga ttactcaatc
4321 tatcagatga ggtgtttggg cttcagaaga tggatagcac tagacccagt atcctggaga
4381 agtgcaattt cgatgaatta gataccagta cagtggagaa taatgacgac gagttggaga
4441 ttgtattacg tgacctgata ccttccacac ctaccggttg tagaagaagt ttctctattc
4501 ctcttacacc cagtaagttg actatacctt cagatggggg tcagcatcgt gctagagttc
4561 ttagcttcag cgctaccagc aacgacaact ctacagatat ctggtgagtc ggacgtttta
4621 atataatttt caatggtagt gtttaacact catttctccc aataaatcac aaaagtacaa
4681 attattgtca tctagacaca tttttaaata atcgccttta tcttggctct attctattct
4741 aattattttt gttaaataag acaaatacac catgctttat aatgaaggcc attaatcaaa
4801 aagttacaaa atgacaacca ttcaagtgca tgtccgactg tggcatagcg ccgtgtggac
4861 ggcttatccc ctatagtgag tcgtattaca tggtcatagc tgtttcctgg cagctctggc
4921 ccgtgtctca aaatctctga tgttacattg cacaagataa aaatatatca tcatgaacaa
4981 taaaactgtc tgcttacata aacagtaata caaggggtgt tatgagccat attcaacggg
5041 aaacgtcgag gccgcgatta aattccaaca tggatgctga tttatatggg tataaatggg
5101 ctcgcgataa tgtcgggcaa tcaggtgcga caatctatcg cttgtatggg aagcccgatg
5161 cgccagagtt gtttctgaaa catggcaaag gtagcgttgc caatgatgtt acagatgaga
5221 tggtcagact aaactggctg acggaattta tgcctcttcc gaccatcaag cattttatcc
5281 gtactcctga tgatgcatgg ttactcacca ctgcgatccc cggaaaaaca gcattccagg
5341 tattagaaga atatcctgat tcaggtgaaa atattgttga tgcgctggca gtgttcctgc
5401 gccggttgca ttcgattcct gtttgtaatt gtccttttaa cagcgatcgc gtatttcgtc
5461 tcgctcaggc gcaatcacga atgaataacg gtttggttga tgcgagtgat tttgatgacg

5521 agcgtaatgg ctggcctgtt gaacaagtct ggaaagaaat gcataaactt ttgccattct
5581 caccggattc agtcgtcact catggtgatt tctcacttga taaccttatt tttgacgagg
5641 ggaaattaat aggttgtatt gatgttggac gagtcggaat cgcagaccga taccaggatc
5701 ttgccatcct atggaactgc ctcggtgagt tttctccttc attacagaaa cggctttttc
5761 aaaaatatgg tattgataat cctgatatga ataaattgca gtttcatttg atgctcgatg
5821 agtttttcta atcagaattg gttaattggt tgtaacactg gcagagcatt acgctgactt
5881 gacgggacgg cgcaagctca tgaccaaaat cccttaacgt gagttacgcg tcgttccact
5941 gagcgtcaga ccccgtagaa aagatcaaag gatcttcttg agatcctttt tttctgcgcg
6001 taatctgctg cttgcaaaca aaaaaaccac cgctaccagc ggtggtttgt ttgccggatc
6061 aagagctacc aactcttttt ccgaaggtaa ctggcttcag cagagcgcag ataccaaata
6121 ctgttcttct agtgtagccg tagttaggcc accacttcaa gaactctgta gcaccgccta
6181 catacctcgc tctgctaatc ctgttaccag tggctgctgc cagtggcgat aagtcgtgtc
6241 ttaccgggtt ggactcaaga cgatagttac cggataaggc gcagcggtcg ggctgaacgg
6301 ggggttcgtg cacacagccc agcttggagc gaacgaccta caccgaactg agatacctac
6361 agcgtgagct atgagaaagc gccacgcttc ccgaagggag aaaggcggac aggtatccgg
6421 taagcggcag ggtcggaaca ggagagcgca cgagggagct tccaggggga aacgcctggt
6481 atctttatag tcctgtcggg tttcgccacc tctgacttga gcgtcgattt ttgtgatgct
6541 cgtcaggggg gcggagccta tggaaaaacg ccagcaacgc ggccttttta cggttcctgg
6601 ccttttgctg gccttttgct cacatgtt
//

**4. ppBAalbNixE1E3E4 PUb-YFP plasmid that produced M-linked piggyBac insertion Aalb-M on Aedes albopictus chromosome 1 (Addgene #173666)**

```
LOCUS       AlbNixE1E2E3_PubYFP      8956 bp ds-DNA     circular SYN 22-JUN-2021
SOURCE      synthetic DNA construct
  ORGANISM  synthetic DNA construct
REFERENCE   1  (bases 1 to 8956)
  AUTHORS   .
  TITLE     Direct Submission
  JOURNAL   Exported Friday, Jul 2, 2021 from SnapGene 5.2.5
            https://www.snapgene.com
FEATURES             Location/Qualifiers
     source          1..8956
                     /dnas_title="Exported"
                     /organism="synthetic DNA construct"
                     /mol_type="other DNA"
     primer_bind     4..20
                     /label=M13 rev
     misc_feature    complement(61..196)
                     /label=PiggyBac 5'TR
     misc_feature    77..95
                     /label=TR2
     misc_feature    complement(371..591)
                     /label=attP'
     protein_bind    427..526
                     /label=phage phi-C31 attP
     protein_bind    602..635
                     /label=loxP
     misc_feature    638..640
                     /label=lac promoter
     promoter        645..2559
                     /label=Nix promoter
     exon            2560..3305
                     /label=Exon 1
     misc_feature    2560..2631
                     /label=5' UTR
     misc_feature    2632..2634
                     /label=Start
     exon            3306..3370
                     /label=Exon 3
     exon            3371..3712
                     /label=Exon 4
     misc_feature    3471..3473
                     /label=stop?
     misc_feature    3472..3712
                     /label=3' UTR
     misc_feature    3713..3714
                     /label=added to match isof 1
     misc_feature    3719..3941
```

/label=SV40 term
polyA_signal 3822..3904
/label=SV40 poly(A) signal
promoter 3951..5336
/label=PUb
misc_feature 5337..6053
/label=YFP
misc_feature 6058..6311
/label=SV40
primer_bind 6074..6096
/label=EM820
misc_feature 6175..6296
/label=SV40 polyA signal
protein_bind 6352..6385
/label=loxP
misc_feature complement(6478..6635)
/label=original PiggyBac 3'region
misc_feature complement(6573..6591)
/label=TR2
misc_feature 6587..6607
/label=direct repeat
misc_feature complement(6623..6635)
/label=TR1
primer_bind complement(6765..6781)
/label=M13 fwd
misc_feature join(6813..8956,1..33)
/label=pDONR backbone
terminator 6946..6973
/label=rrnB T2 terminator
rep_origin complement(7303..7891)
/direction=LEFT
/label=ori
CDS complement(8038..8847)
/label=KanR
misc_feature 8918..8936
/label=pDONR-RP
ORIGIN
1 tgccaggaaa cagctatgac catgtaatac gacgatatga tcctgatgca gctagattaa
61 ccctagaaag atagtctgcg taaaattgac gcatgcattc ttgaaatatt gctctctctt
121 tctaaatagc gcgaatccgt cgctgtgcat ttaggacatc tcagtcgccg cttggagctc
181 ccgtgaggcg tgcttgtcaa tgcggtaagt gtcactgatt ttgaactata acgaccgcgt
241 gagtcaaaat gacgcatgat tatcttttac gtgactttta agatttaact catacgataa
301 ttatattgtt atttcatgtt ctacttacgt gataacttat tatatatata ttttcttgtt
361 atagattaga tcgcgctcgc gcgactgacg gtcgtaagca cccgcgtacg tgtccacccc
421 ggtcacaacc ccttgtgtca tgtcggcgac cctacgcccc caactgagag aactcaaagg
481 ttaccccagt tggggcacta ctcccgaaaa ccgcttctga cctgggaaaa cgtgaagccc
541 cggggcatcc gctgagggtt gccgccgggg cttcggtgtg tccgtcagta ctgcaggtac
601 cataacttcg tatagcatac attatacgaa gttataccgg atcctcgcat tttatgagta
661 aaggcccatt actcatatat gggtaaagtg ctttttgtaa aaaatgagta aatcgattta
721 ctcataaatt ataatttcac cgtgtttact tttcgtcttt tgttatattt ttgaactgtt
781 aactattttg tatttatcat ttaattcata caaaattttg cgtttatttt gtagtttatt

841 gcaatctacg aaaattatta atgaattttt acatgtttca aatactctta atcattttcc
901 tcctggaaaa cagaccttcc gagacgtagc tctttgaagt tttcataatg atccatgtat
961 gaaattttcc cctccggctg ttgttcatcg cgcagttgtt cctgttcttc cgttcctcgt
1021 tcttccggcg cttgaagcct tggtgcagtc ggaactgtgg aaaagactga tatattttta
1081 tttcacggtc gttaatgaaa atgatgctgt cactatggaa cttacctgca aattccatgc
1141 ttttgtagcc tttcttgtag ctccacatga tgatttgttg ctcctgtatt atcttcgagt
1201 accaaacctt gtgaaacgga aataatccat cgtaaatgaa aattgaacgg tatcaattta
1261 tccaggaagt acttaccgat taaaatatca atgtatttgg gaacctcgac tctaatccgc
1321 gtgggaatcc gatggacgcg gtatcggtca ttgaacccga tgctgaaggg cctcgactgg
1381 ctatagcggt gccgtgaaaa tcaaaatatc taaaatttcc cgatagtgtt atgatcaggg
1441 gatccgcatg cggcaaggta ggcctactgc tcatcttttg cgagtgtgtt agtgtgactg
1501 gcaacgcgac cagaagtcta gtcgaatcct gcttggaaat cggtttgttt tgtcgatggg
1561 ggtggcccgc gaaatcgcat tgtgttgttt gggaaggccc gctcgatgca tgttaggtag
1621 gcgtagtttg ttcgagtttg acgtgcctgc accctccctg gttatgatgg cgcactaatg
1681 cattccgaga aaagttgcta acttgaaaat tttcatccaa atatgggtaa gttcatttac
1741 ccatgtttgg gtatagcgat tttacctata atatgggcaa atttgaccgt ttttcaaagg
1801 tatattttac ccatattatg ggtaatgcat atgaaaccca aatatgggta aatcaactac
1861 tgatctatgg gtaaactgat cttagcgtgt atgtttcaca cgtgacattc actggtttta
1921 caattggctt ctttagcagt gttgggataa ttccatatta tgaatctgca tgccaaactg
1981 ggccgaaatc caaattttca tcaattttgg tgcacgggaa cctatttaaa tatcaatttg
2041 aagtttgtat gggagcgatt tgtcgaatca cccctcgttg cattttgtac tgggcggagc
2101 tgtcaaacag ttgcctagct gtcaaaaggt gatttcgaat aatctctttg aaattgattt
2161 taggtatcaa aataaagttc taaaaatctg aaaaaaatca tagtggctca gaaaaaggtg
2221 ctctttcgta taaaatcaaa aaatgaacac ttttttcaaa atttaaaaac ccaattttcg
2281 caaatctata gggctctgca cgaagttctc tccctctctt tcgctctcat tgagattttg
2341 taaacaacaa ggccaggaaa tgtcaaaatc ccatacaaaa tcaaaacagt gcagtgccct
2401 atatgtaaaa cacatcactc cgacgtgtaa atttttttga gtgttgattt aatcaaagtg
2461 aataaaaata ttagttttat gacatacttg ttttctgagt gtagcaaaaa tatgaaaaca
2521 catttttgta ctttgaatgt taagcgtgta tgctttttgt gtcaatatgt caattgttaa
2581 acccatgtta atagttttaa ttttttttta atcaaattct ttttttaagt aatgtacagt
2641 aaaagtgaac ttaatctcat taacaatcaa tttgaataca ttaaaaaaata ttgcatatac
2701 attggaaaca ttcccgccga agtatcgaaa acagatttaa ttgcaaaatt ttccgtattt
2761 ggtgaaatat ctaacttata catgaagtca ttcattcagt tttgtgatgt gaaaccggca
2821 gttgttcgtt acagactgat gaaaagtgta aaggaatctt caagtttaca caatagtcga
2881 tatattcaat cggttttaat agttctgcca ctagattctt cctacaataa ttactttctt
2941 ccttacaaca cttgtgttgt ggtatacact tataacaaat ttggcatggt agatttttat
3001 caaaaattca gtaaattagg agatatacat gcgatgaaga aagctacaaa tgtcatggtt
3061 tacattagct ttgtatcaga aagagctgca aggaccattc tggatactaa gcctacagat
3121 atacatataa atgtacaaac aattaatcat gttacacgaa atattaacgt atgcttaata
3181 gattttgaaa aggaatgtac atcaaatacg gcgataaaat taacactttt atataatcgc
3241 tcaattggaa tattcggact accatctaat ttcacagaag caaaactgca cgatgaattt
3301 tcaagctgcg gaccgttcaa gaaattttgg cagcggaaac atttcttgac cttttccaca
3361 atacttatca gctttggaaa atccccagag gtagatcaat cctactccgc caaaaacttc
3421 cctggcgatg gaagctcttc cgaaccgatc ttcccgagtc ggtgtcgagc tagattctgt
3481 tttttccggt tggagactgg tctccggttc actggcgccc cagcgaccgt gctttccatg
3541 acgttatcga tctagcctac tctcccggcg gtggacctag cctactctga tggaaaaaat
3601 ccccagcgaa agaagttctc ccgaaaccga gaattcaatc tcgaccatcc aggtgccgag
3661 gtcgacccag cgatgccggt tggaggttga cttccagttc actatcactc cgattaagca
3721 taatcagcca taccacattt gtagaggttt tacttgcttt aaaaaacctc ccacacctcc
3781 ccctgaacct gaaacataaa atgaatgcaa ttgttgttgt taacttgttt attgcagctt
3841 ataatggtta caaataaagc aatagcatca caaatttcac aaataaagca tttttcttca

3901 ctgcattcta gttgtggttt gtccaaactc atcaatgtat ctaagtatcg aatccatctt
3961 tacatgtagc ttgtgcattg aatccaatta taatttgcct tggcaccagc tgagccagac
4021 aagaaagaaa gcttcccaga agtatatcga tttagaaggg ttgacgtcac tttgctgact
4081 gcactaatac agcaaatgat acaattagaa tgattcaagt gaaattccca aattactgct
4141 ttgtctctgg atttggttat cagattacat tcgaagctaa gaatagctac cgaaattgtc
4201 gatcaaatca ggaaatcctt tctctatcga aaaaggcatt cgcacatctt tctctgtatg
4261 ccatatacac gaatggtagg tacattgacg tctttgccag aagttgaact gcatcgttca
4321 aggtacagaa tgaacgacta acagacacaa gcacgttttg ctgtccattc agacacaggg
4381 atggtaccca tagtcgatcg atttagagcc atccaaccga acagaggtat atgtatgaat
4441 ggattgcaga aattttctag aagtacaacc accactacgg cagtgtctat aaaacgcccc
4501 tgcaaaggca aaaccagctc aatcgaatac gtttcctagt ggagtgaaca ttacgcggtc
4561 caagtaagca gtgccagtgc aagtgaagtg aagtctctag tgaaaaagag tgatccaatt
4621 agccagagga gaaaatttca gagtgaacaa agctttgttc aaaggacaat tactattaaa
4681 tttgtgaaag tgcatttcgg tgaagggaat cttctagtga aggtaggtaa attaaatgat
4741 gaaattatag ctatgagcga aaactagttt ggtgaatgat tcctttgtct ttgaatgagc
4801 aaactatttt ccaagatggc gactattgag ctttgagtga ttagtgaaaa tttgcaacgc
4861 agtttcatca tcattgataa aacccaattg tgattcacgg cgataatcat atttcgttga
4921 atcatcgctg ctaattgaat taaatttcta gagcaagcgc gaattcgcca tatttctaaa
4981 attaaatatt gtggtgataa ttacccatta aggtaatatt aacacatatc gagaaaaacc
5041 ttgaggaaat cgtgaaaact tgaagatacg caatttccaa actacgtagt tcaaagtcga
5101 aaacaagtta atttttcact aaaaagtagg gcgttgttgt gacgtcatca ccttcaagtg
5161 tatatttttc acttggcctg cgactgcaaa cgcagacaaa gcaaaacaag tttaaaacct
5221 gtcgtgtcgt gctcgaagcc aaaggcaatg aatcaatatc aaatgagagt ttgcatttca
5281 caaccaatta ctgaagcgtt tcctcgtttc tttttctgct caacagagat ttcaacatgg
5341 tgagcaaggg cgaggagctg ttcaccgggg tggtgcccat cctggtcgag ctggacggcg
5401 acgtaaacgg ccacaagttc agcgtgtccg gcgagggcga gggcgatgcc acctacggca
5461 agctgaccct gaagttcatc tgcaccaccg gcaagctgcc cgtgccctgg cccaccctcg
5521 tgaccacctt cggctacggc ctgcagtgct tcgcccgcta ccccgaccac atgaagcagc
5581 acgacttctt caagtccgcc atgcccgaag gctacgtcca ggagcgcacc atcttcttca
5641 aggacgacgg caactacaag acccgcgccg aggtgaagtt cgagggcgac accctggtga
5701 accgcatcga gctgaagggc atcgacttca aggaggacgg caacatcctg gggcacaagc
5761 tggagtacaa ctacaacagc cacaacgtct atatcatggc cgacaagcag aagaacggca
5821 tcaaggtgaa cttcaagatc cgccacaaca tcgaggacgg cagcgtgcag ctcgccgacc
5881 actaccagca gaacaccccc atcggcgacg gccccgtgct gctgcccgac aaccactacc
5941 tgagctacca gtccgccctg agcaaagacc ccaacgagaa gcgcgatcac atggtcctgc
6001 tggagttcgt gaccgccgcc gggatcactc tcggcatgga cgagctgtac aagaagcggc
6061 cgcgactcta gatcataatc agccatacca catttgtaga ggttacttgc tttaaaaaac
6121 ctcccacacc tcccccctgaa cctgaaacat aaaatgaatg caattgttgt tgttaacttg
6181 tttattgcag cttataatgg ttacaaataa agcaatagca tcacaaattt cacaaataaa
6241 gcattttttt cactgcattc tagttgtggt ttgtccaaac tcatcaatgt atcttagctt
6301 gtaattcgtc gacgcggata tcgtttaaac cgttgcttgt aattcgtcga cataacttcg
6361 tatagcatac attatacgaa gttatgagct caattcgata aaagttttgt tactttatag
6421 aagaaatttt gagtttttgt ttttttaat aaataaataa acataaataa attgtttgtt
6481 gaatttatta ttagtatgta agtgtaaata taataaaact taatatctat tcaaattaat
6541 aaataaacct cgatatacag accgataaaa cacatgcgtc aattttacgc atgattatct
6601 ttaacgtacg tcacaatatg attatctttc tagggttaat ctagctgcgt gttctgcagc
6661 gtgtcgagca tcttcatctg ctccatcacg ctgtaaaaca catttgcacc gcgagtctgc
6721 ccgtcctcca cgggttcaaa aacgtgaatg aacgaggcgc gctcactggc cgtcgtttta
6781 caggggatgt cttcatatat atgaagactc ccatctgttg tttgtcggtg aacgctctcc
6841 tgagtaggac aaatccgccg ggagcggatt tgaacgttgt gaagcaacgg cccggagggt
6901 ggcgggcagg acgcccgcca taaactgcca ggcatcaaac taagcagaag gccatcctga

6961 cggatggcct ttttgcgttt ctacaaactc ttcctggcta gcggtacgcg tattaattgc
7021 gttgcgctca ctgcccgctt tccagtcggg aaacctgtcg tgccagctgc attaatgaat
7081 cggccaacgc gcggggagag gcggtttgcg tattgggcgc tcttccgctt cctcgctcac
7141 tgactcgctg cgctcggtcg ttcggctgcg gcgagcggta tcagctcact caaaggcggt
7201 aatacggtta tccacagaat caggggataa cgcaggaaag aacatgtgag caaaaggcca
7261 gcaaaaggcc aggaaccgta aaaaggccgc gttgctggcg tttttccata ggctccgccc
7321 ccctgacgag catcacaaaa atcgacgctc aagtcagagg tggcgaaacc cgacaggact
7381 ataaagatac caggcgtttc cccctggaag ctccctcgtg cgctctcctg ttccgaccct
7441 gccgcttacc ggatacctgt ccgcctttct cccttcggga agcgtggcgc tttctcatag
7501 ctcacgctgt aggtatctca gttcggtgta ggtcgttcgc tccaagctgg gctgtgtgca
7561 cgaacccccc gttcagcccg accgctgcgc cttatccggt aactatcgtc ttgagtccaa
7621 cccggtaaga cacgacttat cgccactggc agcagccact ggtaacagga ttagcagagc
7681 gaggtatgta ggcggtgcta cagagttctt gaagtggtgg cctaactacg gctacactag
7741 aagaacagta tttggtatct gcgctctgct gaagccagtt accttcggaa aaagagttgg
7801 tagctcttga tccggcaaac aaaccaccgc tggtagcggt ggtttttttg tttgcaagca
7861 gcagattacg cgcagaaaaa aaggatctca agaagatcct ttgatctttt ctacggggtc
7921 tgacgctcag tggaacgacg cgtaactcac gttaagggat tttggtcatg agcttgcgcc
7981 gtcccgtcaa gtcagcgtaa tgctctgcca gtgttacaac caattaacca attctgatta
8041 gaaaaactca tcgagcatca aatgaaactg caatttattc atatcaggat tatcaatacc
8101 atatttttga aaaagccgtt tctgtaatga aggagaaaac tcaccgaggc agttccatag
8161 gatggcaaga tcctggtatc ggtctgcgat tccgactcgt ccaacatcaa tacaacctat
8221 taatttcccc tcgtcaaaaa taaggttatc aagtgagaaa tcaccatgag tgacgactga
8281 atccggtgag aatggcaaaa gtttatgcat ttctttccag acttgttcaa caggccagcc
8341 attacgctcg tcatcaaaat cactcgcatc aaccaaaccg ttattcattc gtgattgcgc
8401 ctgagcgaga cgaaatacgc gatcgctgtt aaaaggacaa ttacaaacag gaatcgaatg
8461 caaccggcgc aggaacactg ccagcgcatc aacaatattt tcacctgaat caggatattc
8521 ttctaatacc tggaatgctg ttttccgggg gatcgcagtg gtgagtaacc atgcatcatc
8581 aggagtacgg ataaaatgct tgatggtcgg aagaggcata aattccgtca gccagtttag
8641 tctgaccatc tcatctgtaa catcattggc aacgctacct ttgccatgtt tcagaaacaa
8701 ctctggcgca tcgggcttcc catacaagcg atagattgtc gcacctgatt gcccgacatt
8761 atcgcgagcc catttatacc catataaatc agcatccatg ttggaattta atcgcggcct
8821 cgacgtttcc cgttgaatat ggctcataac accccttgta ttactgttta tgtaagcaga
8881 cagttttatt gttcatgatg atatattttt atcttgtgca atgtaacatc agagattttg
8941 agacacgggc cagagc
//

**5. piggyBac plasmid containing *Cas9* under control of the *Exuperentia* promoter, used to obtain the m-linked m-albR9 *Ae. albopictus* line**

```
LOCUS       pBExuCas9        13831 bp ds-DNA     circular SYN 23-OCT-2017
DEFINITION  synthetic circular DNA
SOURCE      synthetic DNA construct
FEATURES             Location/Qualifiers
     source          1..13831
                     /organism="recombinant plasmid"
                     /mol_type="other DNA"
     rep_origin      complement(63..651)
                     /direction=LEFT
                     /label=ori
                     /note="high-copy-number ColE1/pMB1/pBR322/pUC origin of
                     replication"
     CDS             complement(798..1607)
                     /codon_start=1
                     /gene="aph(3')-Ia"
                     /product="aminoglycoside phosphotransferase"
                     /label=KanR
                     /note="confers resistance to kanamycin in bacteria or G418
                     (Geneticin(R)) in eukaryotes"
                     /translation="MSHIQRETSRPRLNSNMDADLYGYKWARDNVGQSGATIYRLYGKP
                     DAPELFLKHGKGSVANDVTDEMVRLNWLTEFMPLPTIKHFIRTPDDAWLLTTAIPGKTA
                     FQVLEEYPDSGENIVDALAVFLRRLHSIPVCNCPFNSDRVFRLAQAQSRMNNGLVDASD
                     FDDERNGWPVEQVWKEMHKLLPFSPDSVVTHGDFSLDNLIFDEGKLIGCIDVGRVGIAD
                     RYQDLAILWNCLGEFSPSLQKRLFQKYGIDNPDMNKLQFHLMLDEFF"
     primer_bind     1720..1736
                     /label=M13 rev
                     /note="common sequencing primer, one of multiple similar
                     variants"
     misc_feature    1722..1740
                     /label=M13R
                     /note="M13R"
     misc_feature    complement(1777..1912)
                     /label=PiggyBac 5'TR
                     /note="PiggyBac 5'TR"
     misc_feature    1793..1811
                     /label=TR2
                     /note="TR2"
                     /label=Rv2 - PiggyBac NO attP NO lox
     misc_feature    complement(2087..2307)
                     /label=attP'
                     /note="attP'"
     protein_bind    2143..2242
                     /label=phage phi-C31 attP
                     /bound_moiety="phage phi-C31 integrase"
                     /note="attachment site of phage phi-C31"
     protein_bind    2318..2351
                     /label=loxP
                     /bound_moiety="Cre recombinase"
```

/note="Cre-mediated recombination occurs in the 8-bp core
sequence (GCATACAT)."
misc_feature 2354..2356
/label=lac promoter
/note="lac promoter"
misc_feature 2357..3135
/label=ExupPromoter
CDS 3137..3205
/codon_start=1
/label=3xFLAG
/note="3xFLAG"
/translation="MDYKDHDGDYKDHDIDYKDDDDK"
CDS 3206..3256
/codon_start=1
/label=NLS
/note="NLS"
/translation="MAPKKKRKVGIHGVPAA"
CDS 3257..7357
/codon_start=2
/product="Cas9 endonuclease from the Streptococcus pyogenes
Type II CRISPR/Cas system, mutated to improve targeting
specificity (Slaymaker et al., 2016)"
/label=eSpCas9(1.1)
/note="carries the mutations K848A, K1003A, and R1060A"
misc_feature complement(3486..3519)
/label=EM686
/note="EM686"
CDS 7358..7405
/codon_start=1
/product="bipartite nuclear localization signal from
nucleoplasmin"
/label=nucleoplasmin NLS
/translation="KRPAATKKAGQAKKKK"
misc_feature 7397..7637
/label=MCS
/note="pBluescript multiple cloning site"
misc_feature 7397..7405
/label=NLS
/note="NLS"
misc_feature 7415..7637
/label=SV40 term
/note="SV40 term"
3'UTR join(8245..8423,9028..9037)
intron 8424..9027
/label=3'UTRintron
CDS 9042..9722
/codon_start=1
/product="wild-type DsRed"
/label=DsRed1
/note="mammalian codon-optimized"
/translation="MVRSSKNVIKEFMRFKVRMEGTVNGHEFEIEGEGEGRPYEGHNTV

KLKVTKGGPLPFAWDILSPQFQYGSKVYVKHPADIPDYKKLSFPEGFKWERVMNFEDGG
VVTVTQDSSLQDGCFIYKVKFIGVNFPSDGPVMQKKTMGWEASTERLYPRDGVLKGEIH
KALKLKDGGHYLVEFKSIYMAKKPVQLPGYYYVDSKLDITSHNEDYTIVEQYERTEGRH
HLFL"
misc_feature 9724..9960
/label=SV40 Term
/note="SV40 Term"
primer_bind complement(9765..9784)
/label=GATC-pEGFP_C2-RP
polyA_signal 9843..9961
/label=SV40 poly(A) signal
/note="SV40 polyadenylation signal"
protein_bind 9979..10012
/label=loxP
/bound_moiety="Cre recombinase"
/note="Cre-mediated recombination occurs in the 8-bp core
sequence (GCATACAT)."
primer_bind 10012..10041
/label=Fw2 - PiggyBac NO attP NO lox EM834
misc_feature complement(10105..10262)
/label=original PiggyBac 3'region
misc_feature complement(10200..10218)
/label=TR2
/note="TR2"
misc_feature 10214..10234
/label=direct repeat
misc_feature complement(10250..10262)
/label=TR1
/note="TR1"
primer_bind complement(10392..10408)
/label=M13 fwd
/note="common sequencing primer, one of multiple similar
variants"
misc_feature 10681..10878
/label=hs promoter
/note="hs promoter"
misc_feature 10879..11267
/label=hs promoter
/note="hs promoter"
/label=rrnB terminator, EM33
/note="rrnB terminator, EM33"
terminator 11268..11313
/label=rrnB T1 terminator
/note="transcription terminator T1 from the E. coli rrnB
gene"
CDS 11397..13181
/codon_start=1
/label=transposase
/note="transposase"
/translation="MGCSLDDEHILSALLQSDDELVGEDSDSEISDHVSEDDVQSDTEE
AFIDEVHEVQPTSSGSEILDEQNVIEQPGSSLASNRILTLPQRTIRGKNKHCWSTSKST

RRSRVSALNIVRSQRGPTRMCRNIYDPLLCFKLFFTDEIISEIVKWTNAEISLKRRESM
TGATFRDTNEDEIYAFFGILVMTAVRKDNHMSTDDLFDRSLSMVYVSVMSRDRFDFLIR
CLRMDDKSIRPTLRENDVFTPVRKIWDLFIHQCIQNYTPGAHLTIDEQLLGFRGRCPFR
MYIPNKPSKYGIKILMMCDSGTKYMINGMPYLGRGTQTNGVPLGEYYVKELSKPVHGSC
RNITCDNWFTSIPLAKNLLQEPYKLTIVGTVRSNKREIPEVLKNSRSRPVGTSMFCFDG
PLTLVSYKPKPAKMVYLLSSCDEDASINESTGKPQMVMYYNQTKGGVDTLDQMCSVMTC
SRKTNRWPMALLYGMINIACINSFIIYSHNVSSKGEKVQSRKKFMRNLYMSLTSSFMRK
RLEAPTLKRYLRDNISNILPNEVPGTSDDSTEEPVMKKRTYCTYCPSKIRRKANASCKK
CKKVICREHNIDMCQSCF"

terminator 13537..13564
/label=rrnB T2 terminator
/note="transcription terminator T2 from the E. coli rrnB
gene"

ORIGIN
1 aacatgtgag caaaaggcca gcaaaaggcc aggaaccgta aaaaggccgc gttgctggcg
61 tttttccata ggctccgccc ccctgacgag catcacaaaa atcgacgctc aagtcagagg
121 tggcgaaacc cgacaggact ataaagatac caggcgtttc cccctggaag ctccctcgtg
181 cgctctcctg ttccgaccct gccgcttacc ggatacctgt ccgcctttct cccttcggga
241 agcgtggcgc tttctcatag ctcacgctgt aggtatctca gttcggtgta ggtcgttcgc
301 tccaagctgg gctgtgtgca cgaacccccc gttcagcccg accgctgcgc cttatccggt
361 aactatcgtc ttgagtccaa cccggtaaga cacgacttat cgccactggc agcagccact
421 ggtaacagga ttagcagagc gaggtatgta ggcggtgcta cagagttctt gaagtggtgg
481 cctaactacg gctacactag aagaacagta tttggtatct gcgctctgct gaagccagtt
541 accttcggaa aaagagttgg tagctcttga tccggcaaac aaaccaccgc tggtagcggt
601 ggtttttttg tttgcaagca gcagattacg cgcagaaaaa aaggatctca agaagatcct
661 ttgatctttt ctacggggtc tgacgctcag tggaacgacg cgtaactcac gttaagggat
721 tttggtcatg agcttgcgcc gtcccgtcaa gtcagcgtaa tgctctgcca gtgttacaac
781 caattaacca attctgatta gaaaaactca tcgagcatca aatgaaactg caatttattc
841 atatcaggat tatcaatacc atatttttga aaaagccgtt tctgtaatga aggagaaaac
901 tcaccgaggc agttccatag gatggcaaga tcctggtatc ggtctgcgat tccgactcgt
961 ccaacatcaa tacaacctat taatttcccc tcgtcaaaaa taaggttatc aagtgagaaa
1021 tcaccatgag tgacgactga atccggtgag aatggcaaaa gtttatgcat ttctttccag
1081 acttgttcaa caggccagcc attacgctcg tcatcaaaat cactcgcatc aaccaaaccg
1141 ttattcattc gtgattgcgc ctgagcgaga cgaaatacgc gatcgctgtt aaaaggacaa
1201 ttacaaacag gaatcgaatg caaccggcgc aggaacactg ccagcgcatc aacaatattt
1261 tcacctgaat caggatattc ttctaatacc tggaatgctg tttttccggg gatcgcagtg
1321 gtgagtaacc atgcatcatc aggagtacgg ataaaatgct tgatggtcgg aagaggcata
1381 aattccgtca gccagtttag tctgaccatc tcatctgtaa catcattggc aacgctacct
1441 ttgccatgtt tcagaaacaa ctctggcgca tcgggcttcc catacaagcg atagattgtc
1501 gcacctgatt gcccgacatt atcgcgagcc catttatacc catataaatc agcatccatg
1561 ttggaattta atcgcggcct cgacgtttcc cgttgaatat ggctcataac accccttgta
1621 ttactgttta tgtaagcaga cagttttatt gttcatgatg atatattttt atcttgtgca
1681 atgtaacatc agagattttg agacacgggc cagagctgcc aggaaacagc tatgaccatg
1741 taatacgacg atatgatcct gatgcagcta gattaaccct agaaagatag tctgcgtaaa
1801 attgacgcat gcattcttga aatattgctc tctctttcta aatagcgcga atccgtcgct
1861 gtgcatttag gacatctcag tcgccgcttg gagctcccgt gaggcgtgct tgtcaatgcg
1921 gtaagtgtca ctgattttga actataacga ccgcgtgagt caaaatgacg catgattatc
1981 ttttacgtga cttttaagat ttaactcata cgataattat attgttattt catgttctac
2041 ttacgtgata acttattata tatatatttt cttgttatag attagatcgc gctcgcgcga
2101 ctgacggtcg taagcacccg cgtacgtgtc caccccggtc acaacccctt gtgtcatgtc
2161 ggcgacccta cgcccccaac tgagagaact caaaggttac cccagttggg gcactactcc

2221 cgaaaaccgc ttctgacctg ggaaaacgtg aagccccggg gcatccgctg agggttgccg
2281 ccggggcttc ggtgtgtccg tcagtactgc aggtaccata acttcgtata gcatacatta
2341 tacgaagtta taccggatcc aatatagata ttttgatgct tgaggcgaaa aaaaaggtcc
2401 gcagtgttat ttttgaaaaa tttaaggtgg aaaacattgg atgacacttt tccgtgcatt
2461 gaaatcaata cataaaaata aattacctgg catccctgtc cggtagtggt gagtgcacaa
2521 tgtgatggca ttggtgacat tggtgagtgc atgcatgatt aaaaaatata aacggtcgac
2581 atttttcgca tttcttgcca caaattccat tggcatctca aatattacaa taattccgca
2641 tcgagtataa attttcactt ccatcaatgg tcacatttaa actaatattc gaatttttcc
2701 gccggatatt ccacaaaacc cggcacgtaa tcaactataa ctcattctag taccgcaccg
2761 acactaacga aattttgctc ccgtttcgtc cgacttattt ttgctctaga ccagatcgac
2821 cttgccaacc ttcacttcaa gttcatcaag ccaaggacga gcatcgccgc gttggaagaa
2881 aaaaaaggaa agtttcagaa aagtgttctg tggaatattt atcctcaaaa gtgaaaatct
2941 aaaacacaaa cgggtcgcta aaagcgtgcg tgaaaacatt gataaaaacc tgtgtgaaaa
3001 ttgcaaattt cttcgaaaga gtattggtaa gtgtcgttct tttttccacc atgtgaccat
3061 tttgtagaat tttccttttc ggaggaaaag ctggttgctg ggataaaaga ggttatcttt
3121 tgtctacaga gtgaaaatgg actataagga ccacgacgga gactacaagg atcatgatat
3181 tgattacaaa gacgatgacg ataagatggc cccaaagaag aagcggaagg tcggtatcca
3241 cggagtccca gcagccgaca agaagtacag catcggcctg gacatcggca ccaactctgt
3301 gggctgggcc gtgatcaccg acgagtacaa ggtgcccagc aagaaattca aggtgctggg
3361 caacaccgac cggcacagca tcaagaagaa cctgatcgga gccctgctgt tcgacagcgg
3421 cgaaacagcc gaggccaccc ggctgaagag aaccgccaga agaagataca ccagacggaa
3481 gaaccggatc tgctatctgc aagagatctt cagcaacgag atggccaagg tggacgacag
3541 cttcttccac agactggaag agtccttcct ggtggaagag gataagaagc acgagcggca
3601 ccccatcttc ggcaacatcg tggacgaggt ggcctaccac gagaagtacc ccaccatcta
3661 ccacctgaga aagaaactgg tggacagcac cgacaaggcc gacctgcggc tgatctatct
3721 ggccctggcc cacatgatca agttccgggg ccacttcctg atcgagggcg acctgaaccc
3781 cgacaacagc gacgtggaca agctgttcat ccagctggtg cagacctaca accagctgtt
3841 cgaggaaaac cccatcaacg ccagcggcgt ggacgccaag gccatcctgt ctgccagact
3901 gagcaagagc agacggctgg aaaatctgat cgcccagctg cccggcgaga agaagaatgg
3961 cctgttcgga aacctgattg ccctgagcct gggcctgacc cccaacttca agagcaactt
4021 cgacctggcc gaggatgcca aactgcagct gagcaaggac acctacgacg acgacctgga
4081 caacctgctg gcccagatcg gcgaccagta cgccgacctg tttctggccg ccaagaacct
4141 gtccgacgcc atcctgctga gcgacatcct gagagtgaac accgagatca ccaaggcccc
4201 cctgagcgcc tctatgatca agagatacga cgagcaccac caggacctga ccctgctgaa
4261 agctctcgtg cggcagcagc tgcctgagaa gtacaaagag attttcttcg accagagcaa
4321 gaacggctac gccggctaca ttgacggcgg agccagccag gaagagttct acaagttcat
4381 caagcccatc ctggaaaaga tggacggcac cgaggaactg ctcgtgaagc tgaacagaga
4441 ggacctgctg cggaagcagc ggaccttcga caacggcagc atcccccacc agatccacct
4501 gggagagctg cacgccattc tgcggcggca ggaagatttt tacccattcc tgaaggacaa
4561 ccgggaaaag atcgagaaga tcctgacctt ccgcatcccc tactacgtgg gccctctggc
4621 caggggaaac agcagattcg cctggatgac cagaaagagc gaggaaacca tcaccccctg
4681 gaacttcgag gaagtggtgg acaagggcgc ttccgcccag agcttcatcg agcggatgac
4741 caacttcgat aagaacctgc ccaacgagaa ggtgctgccc aagcacagcc tgctgtacga
4801 gtacttcacc gtgtataacg agctgaccaa agtgaaatac gtgaccgagg gaatgagaaa
4861 gcccgccttc ctgagcggcg agcagaaaaa ggccatcgtg gacctgctgt tcaagaccaa
4921 ccggaaagtg accgtgaagc agctgaaaga ggactacttc aagaaaatcg agtgcttcga
4981 ctccgtggaa atctccggcg tggaagatcg gttcaacgcc tccctgggca cataccacga
5041 tctgctgaaa attatcaagg acaaggactt cctggacaat gaggaaaacg aggacattct
5101 ggaagatatc gtgctgaccc tgacactgtt tgaggacaga gagatgatcg aggaacggct
5161 gaaaacctat gcccacctgt tcgacgacaa agtgatgaag cagctgaagc ggcggagata
5221 caccggctgg ggcaggctga gccggaagct gatcaacggc atccgggaca agcagtccgg

5281 caagacaatc ctggatttcc tgaagtccga cggcttcgcc aacagaaact tcatgcagct
5341 gatccacgac gacagcctga cctttaaaga ggacatccag aaagcccagg tgtccggcca
5401 gggcgatagc ctgcacgagc acattgccaa tctggccggc agccccgcca ttaagaaggg
5461 catcctgcag acagtgaagg tggtggacga gctcgtgaaa gtgatgggcc ggcacaagcc
5521 cgagaacatc gtgatcgaaa tggccagaga gaaccagacc acccagaagg gacagaagaa
5581 cagccgcgag agaatgaagc ggatcgaaga gggcatcaaa gagctgggca gccagatcct
5641 gaaagaacac cccgtggaaa acacccagct gcagaacgag aagctgtacc tgtactacct
5701 gcagaatggg cgggatatgt acgtggacca ggaactggac atcaaccggc tgtccgacta
5761 cgatgtggac catatcgtgc ctcagagctt tctggccgac gactccatcg acaacaaggt
5821 gctgaccaga agcgacaaga accggggcaa gagcgacaac gtgccctccg aagaggtcgt
5881 gaagaagatg aagaactact ggcggcagct gctgaacgcc aagctgatta cccagagaaa
5941 gttcgacaat ctgaccaagg ccgagagagg cggcctgagc gaactggata aggccggctt
6001 catcaagaga cagctggtgg aaacccggca gatcacaaag cacgtggcac agatcctgga
6061 ctcccggatg aacactaagt acgacgagaa tgacaagctg atccgggaag tgaaagtgat
6121 caccctgaag tccaagctgg tgtccgattt ccggaaggat ttccagtttt acaaagtgcg
6181 cgagatcaac aactaccacc acgcccacga cgcctacctg aacgccgtcg tgggaaccgc
6241 cctgatcaaa aagtaccctg cgctggaaag cgagttcgtg tacggcgact acaaggtgta
6301 cgacgtgcgg aagatgatcg ccaagagcga gcaggaaatc ggcaaggcta ccgccaagta
6361 cttcttctac agcaacatca tgaacttttt caagaccgag attaccctgg ccaacggcga
6421 gatccggaag gcgcctctga tcgagacaaa cggcgaaacc ggggagatcg tgtgggataa
6481 gggccgggat tttgccaccg tgcggaaagt gctgagcatg ccccaagtga atatcgtgaa
6541 aaagaccgag gtgcagacag gcggcttcag caaagagtct atcctgccca agaggaacag
6601 cgataagctg atcgccagaa agaaggactg ggaccctaag aagtacggcg gcttcgacag
6661 ccccaccgtg gcctattctg tgctggtggt ggccaaagtg gaaaagggca agtccaagaa
6721 actgaagagt gtgaaagagc tgctggggat caccatcatg gaaagaagca gcttcgagaa
6781 gaatcccatc gactttctgg aagccaaggg ctacaaagaa gtgaaaaagg acctgatcat
6841 caagctgcct aagtactccc tgttcgagct ggaaaacggc cggaagagaa tgctggcctc
6901 tgccggcgaa ctgcagaagg gaaacgaact ggccctgccc tccaaatatg tgaacttcct
6961 gtacctggcc agccactatg agaagctgaa gggctccccc gaggataatg agcagaaaca
7021 gctgtttgtg gaacagcaca agcactacct ggacgagatc atcgagcaga tcagcgagtt
7081 ctccaagaga gtgatcctgg ccgacgctaa tctggacaaa gtgctgtccg cctacaacaa
7141 gcaccgggat aagcccatca gagagcaggc cgagaatatc atccacctgt ttaccctgac
7201 caatctggga gcccctgccg ccttcaagta ctttgacacc accatcgacc ggaagaggta
7261 caccagcacc aaagaggtgc tggacgccac cctgatccac cagagcatca ccggcctgta
7321 cgagacacgg atcgacctgt ctcagctggg aggcgacaaa aggccggcgg ccacgaaaaa
7381 ggccggccag gcaaaaaaga aaaagtaaga attctagaca taatcagcca taccacattt
7441 gtagaggttt tacttgcttt aaaaaacctc ccacacctcc ccctgaacct gaaacataaa
7501 atgaatgcaa ttgttgttgt taacttgttt attgcagctt ataatggtta caaataaagc
7561 aatagcatca caaatttcac aaataaagca tttttttcca ctgcattcta gttgtggttt
7621 gtccaaactc atcaatgtat ctaggatgtg cgatccatct ttacatgtag cttgtgcatt
7681 gaatccaatt ataatttgcc ttggcaccag ctgagccaga caagaaagaa agcttcccag
7741 aagtatatcg atttagaagg gttgacgtca ctttgctgac tgcactaata cagcaaatga
7801 tacaattaga atgattcaag tgaaattccc aaattactgc tttgtctctg gatttggtta
7861 tcagattaca ttcgaagcta agaatagcta ccgaaattgt cgatcaaatc aggaaatcct
7921 ttctctatcg aaaaaggcat tcgcacatct ttctctgtat gccatataca cgaatggtag
7981 gtacattgac gtctttgcca gaagttgaac tgcatcgttc aaggtacaga atgaacgact
8041 aacagacaca agcacgtttt gctgtccatt cagacacagg gatggtaccc atagtcgatc
8101 gatttagagc catccaaccg aacagaggta tatgtatgaa tggattgcag aaattttcta
8161 gaagtacaac caccactacg gcagtgtcta taaaacgccc ctgcaaaggc aaaaccagct
8221 caatcgaata cgtttcctag tggagtgaac attacgcggt ccaagtaagc agtgccagtg
8281 caagtgaagt gaagtctcta gtgaaaaaga gtgatccaat tagccagagg agaaaatttc

8341 agagtgaaca aagctttgtt caaaggacaa ttactattaa atttgtgaaa gtgcatttcg
8401 gtgaagggaa tcttctagtg aaggtaggta aattaaatga tgaaattata gctatgagcg
8461 aaaactagtt tggtgaatga ttcctttgtc tttgaatgag caaactattt tccaagatgg
8521 cgactattga gctttgagtg attagtgaaa atttgcaacg cagtttcatc atcattgata
8581 aaacccaatt gtgattcacg gcgataatca tatttcgttg aatcatcgct gctaattgaa
8641 ttaaatttct agagcaagcg cgaattcgcc atatttctaa aattaaatat tgtggtgata
8701 attacccatt aaggtaatat taacacatat cgagaaaaac cttgaggaaa tcgtgaaaac
8761 ttgaagatac gcaatttcca aactacgtag ttcaaagtcg aaaacaagtt aatttttcac
8821 taaaaagtag ggcgttgttg tgacgtcatc accttcaagt gtatattttt cacttggcct
8881 gcgactgcaa acgcagacaa agcaaaacaa gtttaaaacc tgtcgtgtcg tgctcgaagc
8941 caaaggcaat gaatcaatat caaatgagag tttgcatttc acaaccaatt actgaagcgt
9001 ttcctcgttt ctttttctgc tcaacagaga tttcaacaat gatggtgcgc tcctccaaga
9061 acgtcatcaa ggagttcatg cgcttcaagg tgcgcatgga gggcaccgtg aacggccacg
9121 agttcgagat cgagggcgag ggcgagggcc gcccctacga gggccacaac accgtgaagc
9181 tgaaggtgac caagggcggc cccctgccct tcgcctggga catcctgtcc ccccagttcc
9241 agtacggctc caaggtgtac gtgaagcacc ccgccgacat ccccgactac aagaagctgt
9301 ccttccccga gggcttcaag tgggagcgcg tgatgaactt cgaggacggc ggcgtggtga
9361 ccgtgaccca ggactcctcc ctgcaggacg gctgcttcat ctacaaggtg aagttcatcg
9421 gcgtgaactt cccctccgac ggccccgtaa tgcagaagaa gaccatgggc tgggaggcct
9481 ccaccgagcg cctgtacccc cgcgacggcg tgctgaaggg cgagatccac aaggccctga
9541 agctgaagga cggcggccac tacctggtgg agttcaagtc catctacatg gccaagaagc
9601 ccgtgcagct gcccggctac tactacgtgg actccaagct ggacatcacc tcccacaacg
9661 aggactacac catcgtggag cagtacgagc gcaccgaggg ccgccaccac ctgttcctgt
9721 agcggccgcg actctagatc ataatcagcc ataccacatt tgtagaggtt ttacttgctt
9781 taaaaaacct cccacacctc cccctgaacc tgaaacataa aatgaatgca attgttgttg
9841 ttaacttgtt tattgcagct tataatggtt acaaataaag caatagcatc acaaatttca
9901 caaataaagc atttttttca ctgcattcta gttgtggttt gtccaaactc atcaatgtat
9961 cgcttgtaat tcgtcgacat aacttcgtat agcatacatt atacgaagtt atgagctcaa
10021 ttcgataaaa gttttgttac tttatagaag aaattttgag tttttgtttt ttttaataaa
10081 taaataaaca taaataaatt gtttgttgaa tttattatta gtatgtaagt gtaaatataa
10141 taaaacttaa tatctattca aattaataaa taaacctcga tatacagacc gataaaacac
10201 atgcgtcaat tttacgcatg attatcttta acgtacgtca caatatgatt atctttctag
10261 ggttaatcta gctgcgtgtt ctgcagcgtg tcgagcatct tcatctgctc catcacgctg
10321 taaaacacat ttgcaccgcg agtctgcccg tcctccacgg gttcaaaaac gtgaatgaac
10381 gaggcgcgct cactggccgt cgttttacaa cgtcgtgact gggaaaaccc tggcgttacc
10441 caacttaatc gccttgcagc acatccccct ttcgccagct ggcgtaatag cgaagaggcc
10501 cgcaccgatc gcccttccca acagttgcgc agcctgaatg gcgaatggga cgcgccctgt
10561 agcggcgcat taagcgcggc gggtgtggtg gttacgcgca gcgtgaccgc tacacttgcc
10621 agcgccctag cgcccgctcc tttcgctttc ttcccttcct ttctcgccac gttcgccgat
10681 cagataagtt caatgatatc cagtgcagta aaaaaaaaaa tgttttttt atctactttc
10741 cgcaaaaatg ggttttatta acttacatac atactagaat tgatccccga tccccctaga
10801 atcccaaaac aaactggtta ttgtggtagg tcatttgttt ggcagaaaga aaactcgaga
10861 aatttctctg gccgttattc gttattctct cttttctttt tgagtctctc cctctctgca
10921 ctaatgctct ctcactctgt cacacagtaa acggcatact gctctcgttg gttcgagaga
10981 gcgcgcctcg aatgttcgcg aaaagagcgc cggagtataa atagaggcgc ttcgtctacg
11041 gagcgacaat tcaattcaaa caagcaaagt gaacacgtcg ctaagcgaaa gctaagcaaa
11101 taaacaagcg cagctgaaca agctaaacaa tctgcagtaa agtgcaagtt aaagtgaatc
11161 aattaaaagt aaccagcaac caagtaaatc aactgcaact actgaaatct gccaagaagt
11221 aattattgaa tacaagaaga gaactctgaa tagggaattg ggaatttcaa ataaaacgaa
11281 aggctcagtc gaaagactgg gcctttcgtt ttatctgtct agaaggccta attccagctg
11341 agcgccggtc gctaccatta ccagttggtc tggtgtcggg gatcctatat aataaaatgg

11401 gatgttcttt agacgatgag catatcctct ctgctcttct gcaaagcgat gacgagcttg
11461 ttggtgagga ttctgacagt gaaatatcag atcacgtaag tgaagatgac gtccagagcg
11521 atacagaaga agcgtttata gatgaggtac atgaagtgca gccaacgtca agcggtagtg
11581 aaatattaga cgaacaaaat gttattgaac aaccaggttc ttcattggct tctaacagaa
11641 tcttgacctt gccacagagg actattagag gtaagaataa acattgttgg tcaacttcaa
11701 agtccacgag gcgtagccga gtctctgcac tgaacattgt cagatctcaa agaggtccga
11761 cgcgtatgtg ccgcaatata tatgacccac ttttatgctt caaactattt tttactgatg
11821 agataatttc ggaaattgta aaatggacaa atgctgagat atcattgaaa cgtcgggaat
11881 ctatgacagg tgctacattt cgtgacacga atgaagatga aatctatgct ttctttggta
11941 ttctggtaat gacagcagtg agaaaagata accacatgtc cacagatgac ctctttgatc
12001 gatctttgtc aatggtgtac gtctctgtaa tgagtcgtga tcgttttgat tttttgatac
12061 gatgtcttag aatggatgac aaaagtatac ggcccacact tcgagaaaac gatgtattta
12121 ctcctgttag aaaaaatatgg gatctcttta tccatcagtg catacaaaat tacactccag
12181 gggctcattt gaccatagat gaacagttac ttggttttag aggacggtgt ccgtttagga
12241 tgtatatccc aaacaagcca agtaagtatg gaataaaaat cctcatgatg tgtgacagtg
12301 gtacgaagta tatgataaat ggaatgcctt atttgggaag aggaacacag accaacggag
12361 taccactcgg tgaatactac gtgaaggagt tatcaaagcc tgtgcacggt agttgtcgta
12421 atattacgtg tgacaattgg ttcacctcaa tccctttagc aaaaaactta ctacaagaac
12481 cgtataagtt aaccattgtg ggaaccgtgc gatcaaacaa acgcgagata ccggaagtac
12541 tgaaaaacag tcgctccagg ccagtgggaa catcgatgtt ttgttttgac ggaccccctta
12601 ctctcgtctc atataaaccg aagccagcta agatggtata cttattatca tcttgtgatg
12661 aggatgcttc tatcaacgaa agtaccggta aaccgcaaat ggttatgtat tataatcaaa
12721 ctaaaggcgg agtggacacg ctagaccaaa tgtgttctgt gatgacctgc agtaggaaga
12781 cgaataggtg gcctatggca ttattgtacg gaatgataaa cattgcctgc ataaattctt
12841 ttattatata cagccataat gtcagtagca agggagaaaa ggttcaaagt cgcaaaaaat
12901 ttatgagaaa cctttacatg agcctgacgt catcgtttat gcgtaagcgt ttagaagctc
12961 ctactttgaa gagatatttg cgcgataata tctctaatat tttgccaaat gaagtgcctg
13021 gtacatcaga tgacagtact gaagagccag taatgaaaaa acgtacttac tgtacttact
13081 gcccctctaa aataaggcga aaggcaaatg catcgtgcaa aaaatgcaaa aaagttattt
13141 gtcgagagca taatattgat atgtgccaaa gttgtttctg actgactaat aagtataatt
13201 tgtttctatt atgtataagt taaactaatt acttatttta taatacaaca tgactgtttt
13261 taaagtacaa aataagttta ttttgtaaa ggagagaatg tttaaaagtt ttgttacttt
13321 atagaagaaa ttttgagttt ttgttttttt ttaataaata aataaacata aattgtttgt
13381 tgaatttgga tccgtcgagg ctttcctgtt gtttgtcggt gaacgctctc ctgagtagga
13441 caaatccgcc gggagcggat ttgaacgttg tgaagcaacg gcccggaggg tggcgggcag
13501 gacgcccgcc ataaactgcc aggcatcaaa ctaagcagaa ggccatcctg acggatggcc
13561 tttttgcgtt tctacaaact cttcctggct agcggtacgc gtattaattg cgttgcgctc
13621 actgcccgct ttccagtcgg gaaacctgtc gtgccagctg cattaatgaa tcggccaacg
13681 cgcggggaga ggcggtttgc gtattgggcg ctcttccgct tcctcgctca ctgactcgct
13741 gcgctcggtc gttcggctgc ggcgagcggt atcagctcac tcaaaggcgg taatacggtt
13801 atccacagaa tcaggggata acgcaggaaa g
//

**6. pENTRPUb-OptpBTransposase helper plasmid expressing a hyper-active, codon-optimized and NLS-modified piggyBac transposase under the control of the *Ae. aegypti* Polyubiquitin promoter**

```
LOCUS       Exported              6013 bp ds-DNA     circular SYN 28-MAY-2019
DEFINITION  synthetic circular DNA
ACCESSION   .
VERSION     .
KEYWORDS    .
SOURCE      synthetic DNA construct
  ORGANISM  synthetic DNA construct
REFERENCE   1  (bases 1 to 6013)
  AUTHORS   .
  TITLE     Direct Submission
  JOURNAL   Exported Wednesday, Feb 3, 2021 from SnapGene 5.2.4
            https://www.snapgene.com
FEATURES             Location/Qualifiers
     promoter        38..1418
                     /label=Aedes aegypti PUb promoter
     3'UTR           join(626..804,1409..1418)
     intron          805..1408
                     /label=3'UTRintron
     CDS             1420..3252
                     /codon_start=1
                     /label=codon optimized
                     /note="codon optimized"
                     /translation="MAPKKKRKVGIHGVPAAGSSLDDEHILSALLQSDDELVGEDSDS
                     EISDHVSEDDVQSDTEEAFIDEVHEVQPTSSGSEILDEQNVIEQPGSSLASNRILTLP
                     QRTIRGKNKHCWSTSKSTRRSRVSALNIVRSQRGPTRMCRNIYDPLLCFKLFFTDEII
                     SEIVKWTNAEISLKRRESMTGATFRDTNEDEIYAFFGILVMTAVRKDNHMSTDDLFDR
                     SLSMVYVSVMSRDRFDFLIRCLRMDDKSIRPTLRENDVFTPVRKIWDLFIHQCIQNYT
                     PGAHLTIDEQLLGFRGRCPFRMYIPNKPSKYGIKILMMCDSGTKYMINGMPYLGRGTQ
                     TNGVPLGEYYVKELSKPVHGSCRNITCDNWFTSIPLAKNLLQEPYKLTIVGTVRSNKR
                     EIPEVLKNSRSRPVGTSMFCFDGPLTLVSYKPKPAKMVYLLSSCDEDASINESTGKPQ
                     MVMYYNQTKGGVDTLDQMCSVMTCSRKTNRWPMALLYGMINIACINSFIIYSHNVSSK
                     GEKVQSRKKFMRNLYMSLTSSFMRKRLEAPTLKRYLRDNISNILPNEVPGTSDDSTEE
                     PVMKKRTYCTYCPSKIRRKANASCKKCKKVICREHNIDMCQSCF"
     misc_feature    1420..1470
                     /label=NLS
                     /note="NLS"
     misc_feature    1471..3252
                     /label=piggyBac transposase ORF
                     /note=""
     misc_feature    3253..3485
                     /label=SV40 terminator
                     /note="SV40"
     misc_feature    complement(3268..3291)
                     /label=EM226
                     /note="EM226"
     misc_feature    3269..3290
                     /label=AG105
                     /note="AG105"
```

misc_feature    complement(3458..3485)
/label=EM503
/note="EM503"
misc_feature    3463..3485
/label=EM172
/note="EM172"
misc_feature    complement(3507..3510)
/label=EM157
/note="EM157"
promoter    complement(3515..3533)
/label=T7 promoter
/note="promoter for bacteriophage T7 RNA polymerase"
primer_bind    complement(3538..3554)
/label=M13 rev
/note="common sequencing primer, one of multiple similar
variants"
misc_feature    complement(3578..3596)
/label=pDONR-RP
/note="pDONR-RP"
CDS    3667..4476
/codon_start=1
/gene="aph(3')-Ia"
/product="aminoglycoside phosphotransferase"
/label=KanR
/note="confers resistance to kanamycin in bacteria or
G418 (Geneticin(R)) in eukaryotes"
/translation="MSHIQRETSRPRLNSNMDADLYGYKWARDNVGQSGATIYRLYGK
PDAPELFLKHGKGSVANDVTDEMVRLNWLTEFMPLPTIKHFIRTPDDAWLLTTAIPGK
TAFQVLEEYPDSGENIVDALAVFLRRLHSIPVCNCPFNSDRVFRLAQAQSRMNNGLVD
ASDFDDERNGWPVEQVWKEMHKLLPFSPDSVVTHGDFSLDNLIFDEGKLIGCIDVGRV
GIADRYQDLAILWNCLGEFSPSLQKRLFQKYGIDNPDMNKLQFHLMLDEFF"
rep_origin    4623..5211
/direction=RIGHT
/label=ori
/note="high-copy-number ColE1/pMB1/pBR322/pUC origin of
replication"
terminator    5541..5568
/label=rrnB T2 terminator
/note="transcription terminator T2 from the E. coli rrnB
gene"
terminator    5660..5746
/gene="Escherichia coli rrnB"
/label=rrnB T1 terminator
/note="transcription terminator T1 from the E. coli rrnB
gene"
primer_bind    complement(5680..5701)
/label=EM841
primer_bind    5810..5826
/label=M13 fwd
/note="common sequencing primer, one of multiple similar
variants"

misc_feature 5810..5825
/label=M13F
/note="M13F"
protein_bind complement(5876..6000)
/gene="mutant version of attR"
/label=attR4
/bound_moiety="LR Clonase(TM)"
/note="recombination site for the Gateway(R) LR reaction"
source 1..6013
/dnas_title="Exported"
ORIGIN
1 gctagcgtcg acggtatcga taagcttgat cggatccatc tttacatgta gcttgtgcat
61 tgaatccaat tataatttgc cttggcacca gctgagccag acaagaaaga aagcttccca
121 gaagtatatc gatttagaag ggttgacgtc actttgctga ctgcactaat acagcaaatg
181 atacaattag aatgattcaa gtgaaattcc caaattactg ctttgtctct ggatttggtt
241 atcagattac attcgaagct aagaatagct accgaaattg tcgatcaaat caggaaatcc
301 tttctctatc gaaaaaggca ttcgcacatc tttctctgta tgccatatac acgaatggta
361 ggtacattga cgtctttgcc agaagttgaa ctgcatcgtt caaggtacag aatgaacgac
421 taacagacac aagcacgttt tgctgtccat tcagacacag ggatggtacc catagtcgat
481 cgatttagag ccatccaacc gaacagaggt atatgtatga atggattgca gaaattttct
541 agaagtacaa ccaccactac ggcagtgtct ataaaacgcc cctgcaaagg caaaaccagc
601 tcaatcgaat acgtttccta gtggagtgaa cattacgcgg tccaagtaag cagtgccagt
661 gcaagtgaag tgaagtctct agtgaaaaag agtgatccaa ttagccagag gagaaaattt
721 cagagtgaac aaagctttgt tcaaaggaca attactatta aatttgtgaa agtgcatttc
781 ggtgaaggga atcttctagt gaaggtaggt aaattaaatg atgaaattat agctatgagc
841 gaaaactagt ttggtgaatg attcctttgt ctttgaatga gcaaactatt ttccaagatg
901 gcgactattg agctttgagt gattagtgaa aatttgcaac gcagtttcat catcattgat
961 aaaacccaat tgtgattcac ggcgataatc atatttcgtt gaatcatcgc tgctaattga
1021 attaaatttc tagagcaagc gcgaattcgc catatttcta aaattaaata ttgtggtgat
1081 aattacccat taaggtaata ttaacacata tcgagaaaaa ccttgaggaa atcgtgaaaa
1141 cttgaagata cgcaatttcc aaactacgta gttcaaagtc gaaaacaagt taatttttca
1201 ctaaaaagta gggcgttgtt gtgacgtcat caccttcaag tgtatatttt tcacttggcc
1261 tgcgactgca aacgcagaca aagcaaaaca agtttaaaac ctgtcgtgtc gtgctcgaag
1321 ccaaaggcaa tgaatcaata tcaaatgaga gtttgcattt cacaaccaat tactgaagcg
1381 tttcctcgtt tctttttctg ctcaacagag atttcaacaa tggcacccaa gaagaaacgt
1441 aaagtgggaa tacacggtgt cccggctgcg ggatcctcgc tcgacgacga gcacatactt
1501 tcggcactgc ttcaatcgga cgacgagtta gtaggcgagg attcggacag cgagatcagc
1561 gatcacgtct ccgaggacga tgttcaaagc gacacagagg aagccttcat tgacgaagta
1621 cacgaggtgc aaccgacaag ctcgggatcg gaaatcctcg atgagcagaa tgtcatcgag
1681 cagccaggaa gtagcctggc gtcgaacaga atcttaaccc taccgcagcg aaccattagg
1741 ggtaaaaaca aacactgttg gtccaccagt aagagcaccc gccgcagccg tgtctccgcg
1801 ttgaacattg tgcgctcgca gcgtggccct acccgcatgt gtaggaatat ttacgatccg
1861 ctcttgtgtt ttaagctctt tttcactgac gagatcattt ccgaaattgt gaagtggacg
1921 aacgcagaga tctcgctgaa gcgtcgcgag tcgatgacag gcgctacctt tcgtgacacg
1981 aacgaggacg agatctacgc gttcttcggc atcctcgtga tgactgcagt ccggaaggac
2041 aatcatatgt cgaccgatga cctcttcgac cgttcactca gcatggtgta cgtgtcggtg
2101 atgtcgcgcg accgtttcga tttcctgatt cggtgtctgc ggatggacga caagtcgatc
2161 agacccacgt tgcgtgagaa tgatgtcttt acgccagtgc gcaagatctg ggatctcttc
2221 atccaccagt gtattcagaa ctacacaccc ggcgcccacc tgaccatcga cgagcagctg
2281 ctgggttttc gcggtagatg cccattccgc atgtacatcc ccaacaagcc aagcaagtat
2341 ggcatcaaga tcctgatgat gtgcgattct ggtaccaagt acatgatcaa cgggatgcca

2401 tacctgggcc gcggcacgca gacgaacgga gttccgctgg gtgagtacta cgtcaaagaa
2461 ctgtctaagc ccgtccacgg ctcctgtcgg aacattacat gcgataactg gttcaccagc
2521 atccctctgg caaagaatct cctgcaggag ccctacaagc tgaccatcgt cggcacggtt
2581 cgctcgaaca agcgggagat cccggaggtg ctgaagaact cccgtagtcg tccggtcggc
2641 acctccatgt tctgtttcga cggcccgctg accttagtgt cgtataagcc gaagccggcc
2701 aagatggtgt acttactgtc cagctgtgac gaggacgcca gcatcaacga gtcgaccggc
2761 aagccgcaga tggtgatgta ctataaccag accaagggtg gtgtcgatac gctggaccag
2821 atgtgctctg ttatgacatg cagtcgcaag acgaatcgtt ggccgatggc actgctgtac
2881 ggaatgatca acatcgcctg cattaacagc ttcattatct actcgcataa cgtgtctagc
2941 aagggagaga aggtgcagtc gagaaagaag ttcatgcgca atctgtacat gagtctgacg
3001 agctccttca tgcgcaagcg cctggaagct ccgacgctga agcgctacct acgcgataac
3061 atcagcaaca tcctgccgaa cgaggttccg ggaacgtcgg atgactcgac agaggagccg
3121 gttatgaaga agcgcacgta ctgcacgtac tgcccgtcga agatccgccg caaggccaac
3181 gcgtcgtgca agaagtgcaa gaaggtcatc tgccgcgagc ataacatcga catgtgccag
3241 agttgcttct aagctagcta gacataatca gccataccac atttgtagag gttttacttg
3301 ctttaaaaaa cctcccacac ctccccctga acctgaaaca taaaatgaat gcaattgttg
3361 ttgttaactt gtttattgca gcttataatg gttacaaata aagcaatagc atcacaaatt
3421 tcacaaataa agcatttttc ttcactgcat tctagttgtg gtttgtccaa actcatcaat
3481 gtatcgctag cccattagga tgtgcggctt atcccctata gtgagtcgta ttacatggtc
3541 atagctgttt cctggcagct ctggcccgtg tctcaaaatc tctgatgtta cattgcacaa
3601 gataaaaata tatcatcatg aacaataaaa ctgtctgctt acataaacag taatacaagg
3661 ggtgttatga gccatattca acgggaaacg tcgaggccgc gattaaattc caacatggat
3721 gctgatttat atgggtataa atgggctcgc gataatgtcg ggcaatcagg tgcgacaatc
3781 tatcgcttgt atgggaagcc cgatgcgcca gagttgtttc tgaaacatgg caaaggtagc
3841 gttgccaatg atgttacaga tgagatggtc agactaaact ggctgacgga atttatgcct
3901 cttccgacca tcaagcattt tatccgtact cctgatgatg catggttact caccactgcg
3961 atccccggaa aaacagcatt ccaggtatta gaagaatatc ctgattcagg tgaaaatatt
4021 gttgatgcgc tggcagtgtt cctgcgccgg ttgcattcga ttcctgtttg taattgtcct
4081 tttaacagcg atcgcgtatt tcgtctcgct caggcgcaat cacgaatgaa taacggtttg
4141 gttgatgcga gtgattttga tgacgagcgt aatggctggc ctgttgaaca agtctggaaa
4201 gaaatgcata aacttttgcc attctcaccg gattcagtcg tcactcatgg tgatttctca
4261 cttgataacc ttatttttga cgaggggaaa ttaataggtt gtattgatgt tggacgagtc
4321 ggaatcgcag accgatacca ggatcttgcc atcctatgga actgcctcgg tgagttttct
4381 ccttcattac agaaacggct ttttcaaaaa tatggtattg ataatcctga tatgaataaa
4441 ttgcagtttc atttgatgct cgatgagttt ttctaatcag aattggttaa ttggttgtaa
4501 cactggcaga gcattacgct gacttgacgg gacggcgcaa gctcatgacc aaaatccctt
4561 aacgtgagtt acgcgtcgtt ccactgagcg tcagaccccg tagaaaagat caaaggatct
4621 tcttgagatc cttttttct gcgcgtaatc tgctgcttgc aaacaaaaaa accaccgcta
4681 ccagcggtgg tttgtttgcc ggatcaagag ctaccaactc tttttccgaa ggtaactggc
4741 ttcagcagag cgcagatacc aaatactgtt cttctagtgt agccgtagtt aggccaccac
4801 ttcaagaact ctgtagcacc gcctacatac ctcgctctgc taatcctgtt accagtggct
4861 gctgccagtg gcgataagtc gtgtcttacc gggttggact caagacgata gttaccggat
4921 aaggcgcagc ggtcgggctg aacggggggt tcgtgcacac agcccagctt ggagcgaacg
4981 acctacaccg aactgagata cctacagcgt gagctatgag aaagcgccac gcttcccgaa
5041 gggagaaagg cggacaggta tccggtaagc ggcagggtcg gaacaggaga gcgcacgagg
5101 gagcttccag ggggaaacgc ctggtatctt tatagtcctg tcgggtttcg ccacctctga
5161 cttgagcgtc gatttttgtg atgctcgtca ggggggcgga gcctatggaa aaacgccagc
5221 aacgcggcct ttttacggtt cctggccttt tgctggcctt ttgctcacat gttctttcct
5281 gcgttatccc ctgattctgt ggataaccgt attaccgcct ttgagtgagc tgataccgct
5341 cgccgcagcc gaacgaccga gcgcagcgag tcagtgagcg aggaagcgga agagcgccca
5401 atacgcaaac cgcctctccc cgcgcgttgg ccgattcatt aatgcagctg gcacgacagg

5461 tttcccgact ggaaagcggg cagtgagcgc aacgcaatta atacgcgtac cgctagccag
5521 gaagagtttg tagaaacgca aaaaggccat ccgtcaggat ggccttctgc ttagtttgat
5581 gcctggcagt ttatggcggg cgtcctgccc gccaccctcc gggccgttgc ttcacaacgt
5641 tcaaatccgc tcccggcgga tttgtcctac tcaggagagc gttcaccgac aaacaacaga
5701 taaaacgaaa ggcccagtct tccgactgag cctttcgttt tatttgatgc ctggcagttc
5761 cctactctcg cgttaacgct agcatggatg ttttcccagt cacgacgttg taaaacgacg
5821 gccagtctta agctcgggcc cctacaggtc actaatacca tctaagtagt tgattcatag
5881 tgactggata tgttgtgttt tacagtatta tgtagtctgt tttttatgca aaatctaatt
5941 taatatattg atatttatat cattttacgt ttctcgttca acttttctat acaaagttgg
6001 taccgggccc ccc
//

**7. pattBPUb-GFP, attB site-containg plasmid expressing *GFP* under control of the *Ae. aegypti* Pub promoter, inserted in the m-linked attP site of *Ae. albopictus* strain mX1 derived from m-albR9**

```
LOCUS       attBPUbGFP               4729 bp ds-DNA     circular SYN 19-OCT-2020
DEFINITION  synthetic circular DNA
SOURCE      synthetic DNA construct
  ORGANISM  recombinant plasmid
FEATURES             Location/Qualifiers
     source          1..4729
                     /organism="recombinant plasmid"
                     /mol_type="other DNA"
     misc_feature    537..552
                     /label=M13F
                     /note="M13F"
     promoter        564..1944
                     /label=AePUb promoter
     3'UTR           join(1152..1330,1935..1944)
     intron          1331..1934
                     /label=3'UTRintron
     CDS             1946..2662
                     /codon_start=1
                     /product="enhanced GFP"
                     /label=EGFP
                     /note="mammalian codon-optimized"
                     /translation="MVSKGEELFTGVVPILVELDGDVNGHKFSVSGEGEGDATYGKLTL
                     KFICTTGKLPVPWPTLVTTLTYGVQCFSRYPDHMKQHDFFKSAMPEGYVQERTIFFKDD
                     GNYKTRAEVKFEGDTLVNRIELKGIDFKEDGNILGHKLEYNYNSHNVYIMADKQKNGIK
                     VNFKIRHNIEDGSVQLADHYQQNTPIGDGPVLLPDNHYLSTQSALSKDPNEKRDHMVLL
                     EFVTAAGITLGMDELYK"
     polyA_signal    2787..2869
                     /label=SV40 poly(A) signal
                     /note="SV40 polyadenylation signal"
     misc_feature    2911..2966
                     /label=attB 2
                     /note="attB 2"
     misc_feature    2914..2951
                     /label=minimal attB
                     /note="minimal attB"
     misc_feature    complement(2990..3008)
                     /label=M13R
                     /note="M13R"
ORIGIN
        1 ctttcctgcg ttatcccctg attctgtgga taaccgtatt accgcctttg agtgagctga
       61 taccgctcgc cgcagccgaa cgaccgagcg cagcgagtca gtgagcgagg aagcggaaga
      121 gcgcccaata cgcaaaccgc ctctccccgc gcgttggccg attcattaat gcagctggca
      181 cgacaggttt cccgactgga aagcgggcag tgagcgcaac gcaattaata cgcgtaccgc
      241 tagccaggaa gagtttgtag aaacgcaaaa aggccatccg tcaggatggc cttctgctta
      301 gtttgatgcc tggcagttta tggcgggcgt cctgcccgcc accctccggg ccgttgcttc
      361 acaacgttca aatccgctcc cggcggattt gtcctactca ggagagcgtt caccgacaaa
```

421 caacagataa aacgaaaggc ccagtcttcc gactgagcct ttcgttttat ttgatgcctg
481 gcagttccct actctcgcgt taacgctagc atggatgttt tcccagtcac gacgttgtaa
541 aacgacggcc agtcttaaga tccatcttta catgtagctt gtgcattgaa tccaattata
601 atttgccttg gcaccagctg agccagacaa gaaagaaagc ttcccagaag tatatcgatt
661 tagaagggtt gacgtcactt tgctgactgc actaatacag caaatgatac aattagaatg
721 attcaagtga aattcccaaa ttactgcttt gtctctggat ttggttatca gattacattc
781 gaagctaaga atagctaccg aaattgtcga tcaaatcagg aaatcctttc tctatcgaaa
841 aaggcattcg cacatctttc tctgtatgcc atatacacga atggtaggta cattgacgtc
901 tttgccagaa gttgaactgc atcgttcaag gtacagaatg aacgactaac agacacaagc
961 acgttttgct gtccattcag acacaggat ggtacccata gtcgatcgat ttagagccat
1021 ccaaccgaac agaggtatat gtatgaatgg attgcagaaa ttttctagaa gtacaaccac
1081 cactacggca gtgtctataa aacgcccctg caaaggcaaa accagctcaa tcgaatacgt
1141 ttcctagtgg agtgaacatt acgcggtcca agtaagcagt gccagtgcaa gtgaagtgaa
1201 gtctctagtg aaaaagagtg atccaattag ccagaggaga aaatttcaga gtgaacaaag
1261 ctttgttcaa aggacaatta ctattaaatt tgtgaaagtg catttcggtg aagggaatct
1321 tctagtgaag gtaggtaaat taaatgatga aattatagct atgagcgaaa actagtttgg
1381 tgaatgattc ctttgtcttt gaatgagcaa actattttcc aagatggcga ctattgagct
1441 ttgagtgatt agtgaaaatt tgcaacgcag tttcatcatc attgataaaa cccaattgtg
1501 attcacggcg ataatcatat ttcgttgaat catcgctgct aattgaatta aatttctaga
1561 gcaagcgcga attcgccata tttctaaaat taaatattgt ggtgataatt acccattaag
1621 gtaatattaa cacatatcga gaaaaacctt gaggaaatcg tgaaaacttg aagatacgca
1681 atttccaaac tacgtagttc aaagtcgaaa acaagttaat ttttcactaa aaagtagggc
1741 gttgttgtga cgtcatcacc ttcaagtgta tatttttcac ttggcctgcg actgcaaacg
1801 cagacaaagc aaaacaagtt taaaacctgt cgtgtcgtgc tcgaagccaa aggcaatgaa
1861 tcaatatcaa atgagagttt gcatttcaca accaattact gaagcgtttc ctcgtttctt
1921 tttctgctca acagagattt caacaatggt gagcaagggc gaggagctgt tcaccggggt
1981 ggtgcccatc ctggtcgagc tggacggcga cgtaaacggc cacaagttca gcgtgtccgg
2041 cgagggcgag ggcgatgcca cctacggcaa gctgaccctg aagttcatct gcaccaccgg
2101 caagctgccc gtgccctggc ccaccctcgt gaccaccctg acctacggcg tgcagtgctt
2161 cagccgctac cccgaccaca tgaagcagca cgacttcttc aagtccgcca tgcccgaagg
2221 ctacgtccag gagcgcacca tcttcttcaa ggacgacggc aactacaaga cccgcgccga
2281 ggtgaagttc gagggcgaca ccctggtgaa ccgcatcgag ctgaagggca tcgacttcaa
2341 ggaggacggc aacatcctgg ggcacaagct ggagtacaac tacaacagcc acaacgtcta
2401 tatcatggcc gacaagcaga agaacggcat caaggtgaac ttcaagatcc gccacaacat
2461 cgaggacggc agcgtgcagc tcgccgacca ctaccagcag aacaccccca tcggcgacgg
2521 ccccgtgctg ctgcccgaca accactacct gagcacccag tccgccctga gcaaagaccc
2581 caacgagaag cgcgatcaca tggtcctgct ggagttcgtg accgccgccg ggatcactct
2641 cggcatggac gagctgtaca agtaaagcgg ccgcgactct agatcaaatc agccatacca
2701 catttgtaga ggttttactt gctttaaaaa acctcccaca cctccccctg aacctgaaac
2761 ataaaatgaa tgcaattgtt gttgttaact tgtttattgc agcttataat ggttacaaat
2821 aaagcaatag catcacaaat ttcacaaata aagcattttt cttcactgca ttctagttgt
2881 ggtttgtcca aactcatcaa tgtatcgctt tgcgggtgcc agggcgtgcc cttgggctcc
2941 ccgggcgcgt actccacctc acccatatcc cctatagtga gtcgtattac atggtcatag
3001 ctgtttcctg gcagctctgg cccgtgtctc aaaatctctg atgttacatt gcacaagata
3061 aaaatatatc atcatgaaca ataaaactgt ctgcttacat aaacagtaat acaaggggtg
3121 ttatgagcca tattcaacgg gaaacgtcga ggccgcgatt aaattccaac atggatgctg
3181 atttatatgg gtataaatgg gctcgcgata atgtcgggca atcaggtgcg acaatctatc
3241 gcttgtatgg gaagcccgat gcgccagagt tgtttctgaa acatggcaaa ggtagcgttg
3301 ccaatgatgt tacagatgag atggtcagac taaactggct gacggaattt atgcctcttc
3361 cgaccatcaa gcattttatc cgtactcctg atgatgcatg gttactcacc actgcgatcc
3421 ccggaaaaac agcattccag gtattagaag aatatcctga ttcaggtgaa aatattgttg

3481 atgcgctggc agtgttcctg cgccggttgc attcgattcc tgtttgtaat tgtcctttta
3541 acagcgatcg cgtatttcgt ctcgctcagg cgcaatcacg aatgaataac ggtttggttg
3601 atgcgagtga ttttgatgac gagcgtaatg gctggcctgt tgaacaagtc tggaaagaaa
3661 tgcataaact tttgccattc tcaccggatt cagtcgtcac tcatggtgat ttctcacttg
3721 ataaccttat ttttgacgag gggaaattaa taggttgtat tgatgttgga cgagtcggaa
3781 tcgcagaccg ataccaggat cttgccatcc tatggaactg cctcggtgag ttttctcctt
3841 cattacagaa acggcttttt caaaaatatg gtattgataa tcctgatatg aataaattgc
3901 agtttcattt gatgctcgat gagtttttct aatcagaatt ggttaattgg ttgtaacact
3961 ggcagagcat tacgctgact tgacgggacg gcgcaagctc atgaccaaaa tcccttaacg
4021 tgagttacgc gtcgttccac tgagcgtcag accccgtaga aaagatcaaa ggatcttctt
4081 gagatccttt ttttctgcgc gtaatctgct gcttgcaaac aaaaaaacca ccgctaccag
4141 cggtggtttg tttgccggat caagagctac caactctttt tccgaaggta actggcttca
4201 gcagagcgca gataccaaat actgttcttc tagtgtagcc gtagttaggc caccacttca
4261 agaactctgt agcaccgcct acatacctcg ctctgctaat cctgttacca gtggctgctg
4321 ccagtggcga taagtcgtgt cttaccgggt tggactcaag acgatagtta ccggataagg
4381 cgcagcggtc gggctgaacg gggggttcgt gcacacagcc cagcttggag cgaacgacct
4441 acaccgaact gagataccta cagcgtgagc tatgagaaag cgccacgctt cccgaaggga
4501 gaaaggcgga caggtatccg gtaagcggca gggtcggaac aggagagcgc acgagggagc
4561 ttccaggggg aaacgcctgg tatctttata gtcctgtcgg gtttcgccac ctctgacttg
4621 agcgtcgatt tttgtgatgc tcgtcagggg ggcggagcct atggaaaaac gccagcaacg
4681 cggccttttt acggttcctg gccttttgct ggccttttgc tcacatgtt
//

**8. Helper plasmid (piggyBac construct) used to express PhiC31 integrase under control of the PUb promoter**

```
LOCUS       pB PUb-phiC31integrase              7924 bp ds-DNA     circular SYN 07-OCT-2021
DEFINITION  synthetic circular DNA.
ACCESSION   .
VERSION     .
KEYWORDS    .
SOURCE      synthetic DNA construct
  ORGANISM  synthetic DNA construct
REFERENCE   1  (bases 1 to 7924)
  AUTHORS   Li
  TITLE     Direct Submission
  JOURNAL   Exported Thursday, Oct 7, 2021 from SnapGene 5.2.5
            https://www.snapgene.com
FEATURES             Location/Qualifiers
     source          1..7924
                     /organism="synthetic DNA construct"
                     /mol_type="other DNA"
     primer_bind     4..20
                     /label=M13 rev
                     /note="common sequencing primer, one of multiple similar
                     variants"
     misc_feature    complement(61..196)
                     /label=PiggyBac 5'TR
                     /note="PiggyBac 5'TR"
     misc_feature    77..95
                     /label=TR2
                     /note="TR2"
     misc_feature    complement(371..379)
                     /label=attP'
                     /note="attP'"
     promoter        387..1767
                     /label=AePUb promoter
     primer_bind     387..416
                     /label=PUbFw-taag
     3'UTR           join(975..1153,1758..1767)
     intron          1154..1757
                     /label=3'UTRintron
     CDS             1781..1801
                     /codon_start=1
                     /product="nuclear localization signal of SV40 large
                     Tantigen"
                     /label=SV40 NLS
                     /translation="PKKKRKV"
     CDS             1817..3631
                     /codon_start=1
                     /label=integrase
                     /note="integrase"
                     /translation="DTYAGAYDRQSRERENSSAASPATQRSANEDKAADLQREVERDGG
                     RFRFVGHFSEAPGTSAFGTAERPEFERILNECRAGRLNMIIVYDVSRFSRLKVMDAIPI
                     VSELLALGVTIVSTQEGVFRQGNVMDLIHLIMRLDASHKESSLKSAKILDTKNLQRELG
```

GYVGGKAPYGFELVSETKEITRNGRMVNVVINKLAHSTTPLTGPFEFEPDVIRWWWREI
KTHKHLPFKPGSQAAIHPGSITGLCKRMDADAVPTRGETIGKKTASSAWDPATVMRILR
DPRIAGFAAEVIYKKKPDGTPTTKIEGYRIQRDPITLRPVELDCGPIIEPAEWYELQAW
LDGRGRGKGLSRGQAILSAMDKLYCECGAVMTSKRGEESIKDSYRCRRRKVVDPSAPGQ
HEGTCNVSMAALDKFVAERIFNKIRHAEGDEETLALLWEAARRFGKLTEAPEKSGERAN
LVAERADALNALEELYEDRAAGAYDGPVGRKHFRKQQAALTLRQQGAEERLAELEAAEA
PKLPLDQWFPEDADADPTGPKSWWGRASVDDKRVFVGLFVDKIVVTKSTTGRGQGTPIE
KRASITWAKPPTDDDEDDAQDGTEDVAA"
misc_feature 3636..3858
/label=SV40 term
/note="SV40 term"
polyA_signal 3739..3821
/label=SV40 poly(A) signal
/note="SV40 polyadenylation signal"
misc_feature 3864..4164
/label=18xP3
/note="3xP3"
promoter 4165..4374
/label=Hsp70 minimal promoter
misc_feature 4204..4209
/label=TATA
/note="TATA "
CDS 4375..5094
/codon_start=1
/product="enhanced GFP"
/label=EGFP
/note="mammalian codon-optimized"
/translation="MVSKGEELFTGVVPILVELDGDVNGHKFSVSGEGEGDATYGKLTL
KFICTTGKLPVPWPTLVTTLTYGVQCFSRYPDHMKQHDFFKSAMPEGYVQERTIFFKDD
GNYKTRAEVKFEGDTLVNRIELKGIDFKEDGNILGHKLEYNYNSHNVYIMADKQKNGIK
VNFKIRHNIEDGSVQLADHYQQNTPIGDGPVLLPDNHYLSTQSALSKDPNEKRDHMVLL
EFVTAAGITLGMDELYK"
polyA_signal 5216..5297
/label=SV40 poly(A) signal
/note="SV40 polyadenylation signal"
misc_feature complement(5446..5603)
/label=original PiggyBac 3'region
/note="original PiggyBac 3'region"
misc_feature 5555..5575
/label=direct repeat
/note="direct repeat"
misc_feature complement(5591..5603)
/label=TR1
/note="TR1"
primer_bind complement(5733..5749)
/label=M13 fwd
/note="common sequencing primer, one of multiple similar
variants"
terminator 5914..5941
/label=rrnB T2 terminator
/note="transcription terminator T2 from the E. coli

rrnBgene"
rep_origin complement(6271..6859)
/direction=LEFT
/label=ori
/note="high-copy-number ColE1/pMB1/pBR322/pUC origin of replication"
CDS complement(7006..7815)
/codon_start=1
/gene="aph(3')-Ia"
/product="aminoglycoside phosphotransferase"
/label=KanR
/note="confers resistance to kanamycin in bacteria or G418 (Geneticin(R)) in eukaryotes"
/translation="MSHIQRETSRPRLNSNMDADLYGYKWARDNVGQSGATIYRLYGKP
DAPELFLKHGKGSVANDVTDEMVRLNWLTEFMPLPTIKHFIRTPDDAWLLTTAIPGKTA
FQVLEEYPDSGENIVDALAVFLRRLHSIPVCNCPFNSDRVFRLAQAQSRMNNGLVDASD
FDDERNGWPVEQVWKEMHKLLPFSPDSVVTHGDFSLDNLIFDEGKLIGCIDVGRVGIAD
RYQDLAILWNCLGEFSPSLQKRLFQKYGIDNPDMNKLQFHLMLDEFF"
misc_feature 7886..7904
/label=pDONR-RP
/note="pDONR-RP"
ORIGIN
1 tgccaggaaa cagctatgac catgtaatac gacgatatga tcctgatgca gctagattaa
61 ccctagaaag atagtctgcg taaaattgac gcatgcattc ttgaaatatt gctctctctt
121 tctaaatagc gcgaatccgt cgctgtgcat ttaggacatc tcagtcgccg cttggagctc
181 ccgtgaggcg tgcttgtcaa tgcggtaagt gtcactgatt ttgaactata acgaccgcgt
241 gagtcaaaat gacgcatgat tatcttttac gtgactttta agatttaact catacgataa
301 ttatattgtt atttcatgtt ctacttacgt gataacttat tatatatata ttttcttgtt
361 atagattaga tcgcgctcgc ggatccatct ttacatgtag cttgtgcatt gaatccaatt
421 ataatttgcc ttggcaccag ctgagccaga caagaaagaa agcttcccag aagtatatcg
481 atttagaagg gttgacgtca ctttgctgac tgcactaata cagcaaatga tacaattaga
541 atgattcaag tgaaattccc aaattactgc tttgtctctg gatttggtta tcagattaca
601 ttcgaagcta agaatagcta ccgaaattgt cgatcaaatc aggaaatcct ttctctatcg
661 aaaaaggcat tcgcacatct ttctctgtat gccatataca cgaatggtag gtacattgac
721 gtctttgcca gaagttgaac tgcatcgttc aaggtacaga atgaacgact aacagacaca
781 agcacgtttt gctgtccatt cagacacagg gatggtaccc atagtcgatc gatttagagc
841 catccaaccg aacagaggta tatgtatgaa tggattgcag aaattttcta gaagtacaac
901 caccactacg gcagtgtcta taaaacgccc ctgcaaaggc aaaaccagct caatcgaata
961 cgtttcctag tggagtgaac attacgcggt ccaagtaagc agtgccagtg caagtgaagt
1021 gaagtctcta gtgaaaaaga gtgatccaat tagccagagg agaaaatttc agagtgaaca
1081 aagctttgtt caaaggacaa ttactattaa atttgtgaaa gtgcatttcg gtgaagggaa
1141 tcttctagtg aaggtaggta aattaaatga tgaaattata gctatgagcg aaaactagtt
1201 tggtgaatga ttcctttgtc tttgaatgag caaactattt tccaagatgg cgactattga
1261 gctttgagtg attagtgaaa atttgcaacg cagtttcatc atcattgata aaacccaatt
1321 gtgattcacg gcgataatca tatttcgttg aatcatcgct gctaattgaa ttaaatttct
1381 agagcaagcg cgaattcgcc atatttctaa aattaaatat tgtggtgata attacccatt
1441 aaggtaatat taacacatat cgagaaaaac cttgaggaaa tcgtgaaaac ttgaagatac
1501 gcaatttcca aactacgtag ttcaaagtcg aaaacaagtt aatttttcac taaaaagtag
1561 ggcgttgttg tgacgtcatc accttcaagt gtatattttt cacttggcct gcgactgcaa
1621 acgcagacaa agcaaaacaa gtttaaaacc tgtcgtgtcg tgctcgaagc caaaggcaat
1681 gaatcaatat caaatgagag tttgcatttc acaaccaatt actgaagcgt ttcctcgttt

1741 ctttttctgc tcaacagaga tttcaacaat gagcggccct ccaaaaaaga agagaaaggt
1801 agaagacccg ggcggcgaca cgtacgcggg tgcttacgac cgtcagtcgc gcgagcgcga
1861 aaattcgagc gcagcaagcc cagcgacaca gcgtagcgcc aacgaagaca aggcggccga
1921 ccttcagcgc gaagtcgagc gcgacggggg ccggttcagg ttcgtcgggc atttcagcga
1981 agcgccgggc acgtcggcgt tcgggacggc ggagcgcccg gagttcgaac gcatcctgaa
2041 cgaatgccgc gccgggcggc tcaacatgat cattgtctat gacgtgtcgc gcttctcgcg
2101 cctgaaggtc atggacgcga ttccgattgt ctcggaattg ctcgccctgg gcgtgacgat
2161 tgtttccact caggaaggcg tcttccggca gggaaacgtc atggacctga ttcacctgat
2221 tatgcggctc gacgcgtcgc acaaagaatc ttcgctgaag tcggcgaaga ttctcgacac
2281 gaagaacctt cagcgcgaat tgggcgggta cgtcggcggg aaggcgcctt acggcttcga
2341 gcttgtttcg gagacgaagg agatcacgcg caacggccga atggtcaatg tcgtcatcaa
2401 caagcttgcg cactcgacca ctccccttac cggacccttc gagttcgagc ccgacgtaat
2461 ccggtggtgg tggcgtgaga tcaagacgca caaacacctt cccttcaagc cgggcagtca
2521 agccgccatt cacccgggca gcatcacggg gctttgtaag cgcatggacg ctgacgccgt
2581 gccgacccgg ggcgagacga ttgggaagaa gaccgcttca agcgcctggg acccggcaac
2641 cgttatgcga atccttcggg acccgcgtat tgcgggcttc gccgctgagg tgatctacaa
2701 gaagaagccg gacggcacgc cgaccacgaa gattgagggt taccgcattc agcgcgaccc
2761 gatcacgctc cggccggtcg agcttgattg cggaccgatc atcgagcccg ctgagtggta
2821 tgagcttcag gcgtggttgg acggcagggg gcgcggcaag gggctttccc gggggcaagc
2881 cattctgtcc gccatggaca agctgtactg cgagtgtggc gccgtcatga cttcgaagcg
2941 cggggaagaa tcgatcaagg actcttaccg ctgccgtcgc cggaaggtgg tcgacccgtc
3001 cgcacctggg cagcacgaag gcacgtgcaa cgtcagcatg gcggcactcg acaagttcgt
3061 tgcggaacgc atcttcaaca agatcaggca cgccgaaggc gacgaagaga cgttggcgct
3121 tctgtgggaa gccgcccgac gcttcggcaa gctcactgag gcgcctgaga agagcggcga
3181 acgggcgaac cttgttgcgg agcgcgccga cgccctgaac gcccttgaag agctgtacga
3241 agaccgcgcg gcaggcgcgt acgacggacc cgttggcagg aagcacttcc ggaagcaaca
3301 ggcagcgctg acgctccggc agcaaggggc ggaagagcgg cttgccgaac ttgaagccgc
3361 cgaagccccg aagcttcccc ttgaccaatg gttccccgaa gacgccgacg ctgacccgac
3421 cggccctaag tcgtggtggg ggcgcgcgtc agtagacgac aagcgcgtgt tcgtcgggct
3481 cttcgtagac aagatcgttg tcacgaagtc gactacgggc agggggcagg gaacgcccat
3541 cgagaagcgc gcttcgatca cgtgggcgaa gccgccgacc gacgacgacg aagacgacgc
3601 ccaggacggc acggaagacg tagcggcgta gtaagcataa tcagccatac cacatttgta
3661 gaggttttac ttgctttaaa aaacctccca cacctccccc tgaacctgaa acataaaatg
3721 aatgcaattg ttgttgttaa cttgtttatt gcagcttata atggttacaa ataaagcaat
3781 agcatcacaa atttcacaaa taaagcattt ttcttcactg cattctagtt gtggtttgtc
3841 caaactcatc aatgtatcta agtatctaat tcaattagag actaattcaa ttagagtcta
3901 attcaattag agttatctaa ttcaattaga gactaattca attagagtct aattcaatta
3961 gagttatcta attcaattag agactaattc aattagagtc taattcaatt agagttatct
4021 aattcaatta gagactaatt caattagagt ctaattcaat tagagttatc taattcaatt
4081 agagactaat tcaattagag tctaattcaa ttagagttat ctaattcaat tagagactaa
4141 ttcaattaga gctaattcaa ttaggatcca agcttatcga tttcgaaccc tcgaccgccg
4201 gagtataaat agaggcgctt cgtctacgga gcgacaattc aattcaaaca agcaaagtga
4261 acacgtcgct aagcgaaagc taagcaaata aacaagcgca gctgaacaag ctaaacaatc
4321 ggggtaccgc tagagtcgac ggtaccgcgg gcccgggatc caccggtcgc caccatggtg
4381 agcaagggcg aggagctgtt caccggggtg gtgcccatcc tggtcgagct ggacggcgac
4441 gtaaacggcc acaagttcag cgtgtccggc gagggcgagg gcgatgccac ctacggcaag
4501 ctgaccctga agttcatctg caccaccggc aagctgcccg tgccctggcc caccctcgtg
4561 accaccctga cctacggcgt gcagtgcttc agccgctacc ccgaccacat gaagcagcac
4621 gacttcttca agtccgccat gcccgaaggc tacgtccagg agcgcaccat cttcttcaag
4681 gacgacggca actacaagac ccgcgccgag gtgaagttcg agggcgacac cctggtgaac
4741 cgcatcgagc tgaagggcat cgacttcaag gaggacggca acatcctggg gcacaagctg

4801 gagtacaact acaacagcca caacgtctat atcatggccg acaagcagaa gaacggcatc
4861 aaggtgaact tcaagatccg ccacaacatc gaggacggca gcgtgcagct cgccgaccac
4921 taccagcaga acacccccat cggcgacggc cccgtgctgc tgcccgacaa ccactacctg
4981 agcacccagt ccgccctgag caaagacccc aacgagaagc gcgatcacat ggtcctgctg
5041 gagttcgtga ccgccgccgg gatcactctc ggcatggacg agctgtacaa gtaaagcggc
5101 cgcgactcta gatcaaatca gccataccac atttgtagag gttttacttg ctttaaaaaa
5161 cctcccacac ctccccctga acctgaaaca taaaatgaat gcaattgttg ttgttaactt
5221 gtttattgca gcttataatg gttacaaata aagcaatagc atcacaaatt tcacaaataa
5281 agcatttttt tcactgcatt ctagttgtgg tttgtccaaa ctcatcaatg tatcgcttgt
5341 aattcgtcga catgagctca attcgataaa agttttgtta ctttatagaa gaaattttga
5401 gtttttgttt tttttaataa ataaataaac ataaataaat tgtttgttga atttattatt
5461 agtatgtaag tgtaaatata ataaaactta atatctattc aaattaataa ataaacctcg
5521 atatacagac cgataaaaca catgcgtcaa ttttacgcat gattatcttt aacgtacgtc
5581 acaatatgat tatctttcta gggttaatct agctgcgtgt tctgcagcgt gtcgagcatc
5641 ttcatctgct ccatcacgct gtaaaacaca tttgcaccgc gagtctgccc gtcctccacg
5701 ggttcaaaaa cgtgaatgaa cgaggcgcgc tcactggccg tcgttttaca ggggatgtct
5761 tcatatatat gaagactccc atctgttgtt tgtcggtgaa cgctctcctg agtaggacaa
5821 atccgccggg agcggatttg aacgttgtga agcaacggcc cggagggtgg cgggcaggac
5881 gcccgccata aactgccagg catcaaacta agcagaaggc catcctgacg gatggccttt
5941 ttgcgtttct acaaactctt cctggctagc ggtacgcgta ttaattgcgt tgcgctcact
6001 gcccgctttc cagtcgggaa acctgtcgtg ccagctgcat taatgaatcg gccaacgcgc
6061 ggggagaggc ggtttgcgta ttgggcgctc ttccgcttcc tcgctcactg actcgctgcg
6121 ctcggtcgtt cggctgcggc gagcggtatc agctcactca aaggcggtaa tacggttatc
6181 cacagaatca ggggataacg caggaaagaa catgtgagca aaaggccagc aaaaggccag
6241 gaaccgtaaa aaggccgcgt tgctggcgtt tttccatagg ctccgccccc ctgacgagca
6301 tcacaaaaat cgacgctcaa gtcagaggtg gcgaaacccg acaggactat aaagatacca
6361 ggcgtttccc cctggaagct ccctcgtgcg ctctcctgtt ccgaccctgc cgcttaccgg
6421 atacctgtcc gcctttctcc cttcgggaag cgtggcgctt tctcatagct cacgctgtag
6481 gtatctcagt tcggtgtagg tcgttcgctc caagctgggc tgtgtgcacg aaccccccgt
6541 tcagcccgac cgctgcgcct tatccggtaa ctatcgtctt gagtccaacc cggtaagaca
6601 cgacttatcg ccactggcag cagccactgg taacaggatt agcagagcga ggtatgtagg
6661 cggtgctaca gagttcttga agtggtggcc taactacggc tacactagaa gaacagtatt
6721 tggtatctgc gctctgctga agccagttac cttcggaaaa agagttggta gctcttgatc
6781 cggcaaacaa accaccgctg gtagcggtgg tttttttgtt tgcaagcagc agattacgcg
6841 cagaaaaaaa ggatctcaag aagatccttt gatcttttct acggggtctg acgctcagtg
6901 gaacgacgcg taactcacgt taagggattt tggtcatgag cttgcgccgt cccgtcaagt
6961 cagcgtaatg ctctgccagt gttacaacca attaaccaat tctgattaga aaaactcatc
7021 gagcatcaaa tgaaactgca atttattcat atcaggatta tcaataccat atttttgaaa
7081 aagccgtttc tgtaatgaag gagaaaactc accgaggcag ttccatagga tggcaagatc
7141 ctggtatcgg tctgcgattc cgactcgtcc aacatcaata caacctatta atttcccctc
7201 gtcaaaaata aggttatcaa gtgagaaatc accatgagtg acgactgaat ccggtgagaa
7261 tggcaaaagt ttatgcattt ctttccagac ttgttcaaca ggccagccat tacgctcgtc
7321 atcaaaatca ctcgcatcaa ccaaaccgtt attcattcgt gattgcgcct gagcgagacg
7381 aaatacgcga tcgctgttaa aaggacaatt acaaacagga atcgaatgca accggcgcag
7441 gaacactgcc agcgcatcaa caatattttc acctgaatca ggatattctt ctaatacctg
7501 gaatgctgtt tttccgggga tcgcagtggt gagtaaccat gcatcatcag gagtacggat
7561 aaaatgcttg atggtcggaa gaggcataaa ttccgtcagc cagtttagtc tgaccatctc
7621 atctgtaaca tcattggcaa cgctaccttt gccatgtttc agaaacaact ctggcgcatc
7681 gggcttccca tacaagcgat agattgtcgc acctgattgc ccgacattat cgcgagccca
7741 tttataccca tataaatcag catccatgtt ggaatttaat cgcggcctcg acgtttcccg
7801 ttgaatatgg ctcataacac cccttgtatt actgtttatg taagcagaca gttttattgt

7861 tcatgatgat atattttat cttgtgcaat gtaacatcag agattttgag acacgggcca
7921 gagc
//

**9. Genomic sequence flanking the Aal-M insertion on the 3' piggyBac side.** One Nanopore read was obtained, including 962bp in the piggyBac transposon and ~1400bp of flanking sequence. We corrected the Nanopore read by PCR, placing the forward primer in the piggyBac transposon and the reverse primer in the flanking sequence. Corrected sequence is displayed in capital letters while the remaining low-quality sequence is in lower case. The NCBI BLAST tool in "discontinued megablast" mode detected a fragmented hit on scaffold 16 with 100% query cover and 90% identity. The hit on scaffold 16 is located between the genes LOC109423234 (uncharacterized, 5-9kb on 5' side) and LOC115263786 (uncharacterized protein K02A2.6-like, 61-65kb on 3' side). According to the latest genomic assembly, scaffold 16 is located on chromosome 1, long arm at position 12 (1q12).

TTAAccacGAATTCTTTTAGGGATTCCTCCAGAGTCCTACCAGAAATTCATTTAGGGATATCTCCAGGAAA
TCCTTGAGAAATTCGTCCAGGGATCCAGGAATTCCTCCAGAGTATCCTTCAGGCATTCCGGTCAGAAATC
TTTAAGGGGTTCATCCTCGGATTTGTTCAGCAATCCATCCATGAATTCATCAAGAATACGCTCTGCTGCTT
CTGCAAATGAATTCTTTCAGAATTTCGTCCTGTGATTCTTTCAGGGATATTTATTATTGAGATTATTTTTC
CGAGAAATCCTTCAAGAAGTTATCCATTGATTTTTTTCAAGAACTCCTCCACAAGTTTTTTCAGGGATGTT
TAGAGAGTGTCCTTCAATGATTCTTCCAAAAAATAGTCAAGTCACTCGTTTAGAAATTCCCTCGGCAATA
CCTCAAGGATTCCCTTCAGCGATTGTTTTAGGACTTATCCTAGGGGTTTGTTCAAATATTACATAACGCTA
AGGGGGAGAGAGGGGGTCTAGCTCTGTGTTACGATTCATACAACATTCTTAAAGTTTTCATATAAAATTT
GTTACGTGGGGGAGGGAGGGGGTCTTAAATTGTGAAATTTTGCGTTACGTAATATTTGAAAGAACCTTA
GGATTTCTTCAGGGATTCCACCAGGGACCTCCTTTCGAGTTTTTTTTTGTAAAGATTCTTTCAGAAATTT
CTCCAAGAAATTGCCTAAAAATTTTCCAGGAATCTCTTAAGCAGTTCCTTCAGGGATTGTATCAGGAATT
CCTATACCTAATAAATCTTTCAGGGATTCTTCAAAGCAACACGCATGTTACAGAAGTGTCACGGCAGTGC
GCAAGGTTTTGTTTTCGAAAAAGTCACTGTGACCTACTTTATACAAATAAGTACCTACATGCTTGAATCG
CTTACAAACANTTTTTCCAGGGAATCTATCAGGAATTCACCCAGAGAAAACTTTTTTGGAATTTTCCCAG
GGTTTGTCCGACAATTACCATAAGAAAAAGTATCAGGAGTTCCCCTAGGAAWTCCTTCTCAAATTCCTTC
ATTTTGAATATATTCTAGCGATTCGTCTAGGAATCCCTTCAGGTGAATTTCAGGATTTTTTCAAGAACTTC
ATTATTTGCGAGTTCTCCGGAAATACCTCAAGAATTTCTCCAGGGATTTCTTACAGGAATACCTCGAGGG
TTTCCTCTAGTGGTAGTTACAAGGTTTAACGAGAAGCCTTCAgaattcttccaaagatccttccggaatacgtcccttgacattcctata
ggattatgctcagggatgtttttatttgtatatttgcacatacaactgtcttgcagataagaaagtatagcaaacatggaaagagagtgttcggagaaactgaataaacgggaaa
aaaacctggaacatattttgata

**10. Genomic sequence flanking the Aal-m insertion on the 3' piggyBac side.** In black: sequence confirmed by sequencing PCR products. In Blue: low-quality sequence (2x coverage) obtained from two Nanopore reads

TTAAGCTAGTATTTACATTCTGGTTTCCGATTTAGTTTAATAAGAATCAATCCGCAAACAGAGATAAGAT
TTTGTATCCTGGCCCGAGGTAGTGGCTCACAATGGCTCAACGCAGACCCAAGCAACACAGAAAACATCT
TTGGACCATCAGTATTTCCAAAGATGTTTTTAAGTGAATTATAAGCGATTCAGAAAACATCTTGTGTTA
ATATCCAGCTCTAATTATGCTGATCGAAAACAAACAACATCAAACATGTTATTGTCTATATCACGTATGA
AATATGTTTATATAACTATCAGTGAATCAAAATTCAACCATCGCGCAACAATAATGACAATGTCGCAAC
CTGTATTTGTTGCGACAAAACAACCCATGCGACATATGTTTGTCCCAACTTGAAAATAATGGGATTAACA
TCGTACACCACGAGTTTAGAGATTTTAAGCCTTGCATTGCGATTTCGAACGGTTACTTCGAGTCGGTAA
ACACTGCAACTCTGGTGAAGCTCTATCATCAAACAGTTTCGACTTTGGATTAGTAATAATTGATTATTCA
CGAGTTGAAAAGTACGCAACAAATTCAGCTTCCGTTTTGTCTGCAACAAAAAATTTCGAGATCATGTCAT
TATCTGTCACCAGTAGGGATAAAGACTATCCATCACATCTTCTTTGTTGCTTGTTTTGTGCCTCTTTTGTCT
GACAATTCTCTTCACTGATAACTATTATATAACAGAAACATCAAGTTGCTAATTCAATCATATGTAAAAG
CAAACGCGACAGAACAACAGAAAGAAGATAAAGATAAAAATAAACCAAAATGCAAACATTGTACCGTC
TACCCCCGATGGTTTGAACGACACCTCATGCAAACCAACGGGGTTCTTTTTTTAATTTGAACTTTTAGTA
ACCCTGTGGACATCAACAAACGTACTTTTGGTAACCTTATTTGTTGTTTTGTTTTGATTCTGCGTTCCGTTT
CACCGCGTTCCATGGGCAGAATGACGTTTGAACCATTTTTAGTTTGAACGATGTGCAGATTGGCGGGGGT
CAAATTAAAAAGTGTTCAGATTAGATGCTGTCAAACCAATGGGGGTAGACGGTATGCATTGATTTTAAC
CATTCCTAGGAGGCTCTGGTCGATGGGGGAGAGTGAGTGGTTCAAGGGACCACGTCCCCCACGTCCGAC
GAACGTCCCCGCGCTAATTAGGATCACCATACACAAATTTGGCCATCATCCTCAAGGCCAAAATGTACA
GGCTGCTGATCTACCAACTGGTCTGTCGAATACTATCCAGAAACACCGCGCAACTTGTATTCCGGCCTAT
GTGAAAGAATCAGCATCCAACAGTCGGTGCTATTCCAACTCCAGCGTGAATTCCGGCACTCAAACTTCA
CCTCTCTGCCGTGTGCAGCATAGTTGTCCCATGTTCCAAAACAGCAAACTGAGAAAAACGCGTTTTAAGT
TTCAGCATTGTTTCCATCTCTACGGACAAGTGTAATAAAAACACAAAATGAATCCACCATCATAGCGTTA
GAATTGGTATTTATCATTGAAGTTTGAACAAAAAGTTTCATAAACATACAATTTACTTTGATTACACGGT
CCGACAAAAAATCAACTTTTGTTCTGCCCAGCTCWTAAATTTCACTTTTTCTGATTACTCCCGACTAGAA
ACTCATAAATCCGAAGTTTTGACCACTCCCCCTCCTGGGAGAGGRRGGGCATTGATTTTKRAAATTTTGA
AAAATCGAAAAATTTCGAGGTTTTCGACCGCCATTTCGGCTAAATGCACGATGTCAAAAAAGGAAAAGG
TTGACCRCRGAGCTTTRGGGTTGTGCAACTATCAASWAAATTTCACGAAGATCCATCAAGCCATTCAAW
SGAAACGGTGTTTTTACGAAATGATGTTTTARAGATTTCTTATAKTGCACGCAAAGAATCCAACTACTAT
ATGGCACAATTGTTGATTTAATCATGTCGTTTAGCATCATRGTGTCTTCGAGGAAGTTTTTCACTACAATA
ACGTGCTTCTTTTAACCTGTTGGTAAAAACGATCAATCCCCCTAAAAGTGAGATAGAAAATTTCTTTTTT
AATTTTTCAAGATRCAAGGTTGATGTCTTCACAAGAGTTGTAGAAAATGCTGTTCTGAATAACTTTGTCG
AAGACGCCAAATTCTARTAACTCCGTTAGTTTCAAGATACGCTACGTTTTTCATTACCGGCGGTCTCTTG
AAAGTGTTTTTTCACCTCAATAACTTTTTTCACTTTTGATTCCACAATTCTATTCTCATAGGTCAATCATTT
TGACTTTCTAAAGGAGACAAKTACTTGTACTTTCGCATTTTGCACTTACATTAAAAGTTTTATTACTAGAG
CTACAGAAAGATCTCGATAAARTAGYAGTGCTATTTTTGAAGGTTACATGATAGGGAATCGTAAAGCAA
TGAAAAGCGAAATTATTGTCTTTCCGCTCAATAGTTATTTTGTGGAGACTGTTAGGTGCGAACTTGYYSC
GCSCACTACTGYTATAAGCAAATGTAAGCAAATTACCTCATACAAATGCGAAAGTACAARTAATTGTCTC
CTTTMGAAAGTCAAAATGAGTAAGMCTATTGGAGACWGYGAGAATTAAATTGTGGAATCCAAAAGATG
AAAAAGAAGTTATTGAGAAAAAACACTTTCAAGAGACCGCCSGCAMTGAAMYRTAACGTAATCTTGAA
AACTAACGGAGTTMGCTAAATTTCATGGATATTCACCATCCATTGACACTCCAAGTGATTTAGATGATGC
CGTTGATACTACAACGACCTTCATCATGGAAGCTTTTAAAGA:A:GCATGCCCTCTACGGTCTGTGAAGAT
CACAAGAGGAACCCCTTGG:TGGAACTCTGATCTGGCGAA::ACTCAGGAAACAATGTAGAAAAACCTTTG
CACAAATGTTTCCAGCTTG:AGTGAAGTCAGTCGGTTAAACAATTTCCTTGCGAAATCTAAGGATTTCCG
GGTGAACGAACTTCGTTTGCCAGARTGGCGATCTGACTTCCTCTGATGAGGAAGTTTTGGAATCTTATT
CAGCACACAYTYCCTGGATGTGTGGATATTACATCTTCCGGATGATCCTGATGTCTTTTCTTGTAGTTATG
ATTCTTTRGCTTTTTAACTRTAGAATCGATTGAGTGGGCTCTTAATAGCTTTGCTCCTTTCAAATCTCCTG

GGGCRGATGGGATTTRTCCTATTTTGCTTCCAGAAGGGATTTGATTATTTCAAACATGTTTTGAAAAAAC
TACTTGTTYRCAGTTTTGCTACARGGGTATATTCCCAAATCCTGGCGGGATATTACTGTAAAAGTTTATTC
CAAAAAGTGAGTCGTTRMATMKTRTGAAGAARCAAAGAGTTTCAGACCTATCAGTTTGACCACTCTTTT
CTTCWSAAATGCTWARAACGCAKTGTGGATCATCACATCMGTGATGTTCATCTGGCCAACGATGCCTCT
TCATGTGAACCAACATGCCTACCAATCTGGTAAGTCCACTGTGACTCTTTTACACAAGGTTGTTTACGAT
ATCGAAGGCATTCGCTCCAAAAGCAATCTTGTTTGGGGTGTTTTTCTTAGATATCGAGGGTGCCTTTGAC
AACGTGCCTTTCGATGCCATAWTGGAAGCCACTTTTGAGTCATGGTATATCTCCGATGATTTCCAATTRG
ATTCACCAAATGCTCAAAAACCGATATCTCTTCTCGACAKTGCGTCTAGCRGRGWTWGGAAATTRGTGT
TTGTGGATGCCCCCAAAGGGAATCTTATCACCGCTTTTGTGRRATCTCGTAGCRRATRCGCTATTGAGGT
AACTCAATAATGSYAGCGKTTTYCCTACTTAGYRGTTTTGCCGATGACTACCTARCATTGTTAGTTGGTAT
GTGCATCAGCACCCTTTTCARACCTGATGCAAAGCGCCCTTCAGGTAGTTGAGGGTTRRGTGTCGCCAAT
ATGGCCTTTCRGTTAATCCGAARTRAAACATCTATTGTTCTTTTCACSGRAAGGCGAAACCGTAATGGCC
GTTCGACCTTTTACGTCTCTCTTTGATTCTGAAATCGATGTGACTGAACGGGTAAAGTACGTTGGAGTGC
ATTCTTGATTCCAAGCTTTCCTGGACACCTCRCATTGAGGTCAGAATCAAAAGCTTAATATAGCCTTCRG
GCAATGCCGGCGTACTTTTGARTRCAGCTWGGGTCTAAAACCCAAGTATATCAAATGGATTTACACAAC
TGTRGTTCGGCYAATAKTGGCTTATGGATGTCTTAKTAGTGGTAGCAAAAGGGTGAAGTAGRGAACGGT
CCAATCAAAATTAGGCCATCTCCAAAGGATGWKMTTAATAGCGATGTCCGGAGCGTTCTCTTCAACACC
YRCGGCAGCGCTCGARGATTCTCTTTGACGTTGCCYYACTACWCGATCTCATMWCAAACAGAAGCACTT
TCTTGCACTTACCGGTTACGGGTACTCGGTCTAGMKGAAACTCCTRTAATGAAACGCACATCAACACAC
CTCGTTGTTTTCRCTTYTGGTGAATTGGGACAAAATTGTTCTTGCTCCAAGTGATCTTACAATGGCTTGTA
ATTTTCCATATAGGACATTTTYCACGRAATTCCCCTTCCMGGGAAGAGTGGACATCTGGTTATCTGGAAA
GAAGTRTTTCAGACGGCATCGTATGTTACACTGATGRCTCCCTTCTCGAAGGTCGAGCAGGTGCTGGTGT
TTATTCTCGTGAGCTAARRCTGTATCAGTTTTACTCRCTYGGTGARWCACTRCACCGTTTTTCAGGCCGA
AATCTTTTGCTCTTATGTGCGGAGTGCAATCAGCACTTYTCAAGGGCGCYTGTTGTGAACCCATTGTGGT
ACGCTTTGTGCACAKKTAAKTRCCCTCTTCCAGGGAGTATTTTTTCTATYTCCTACCTGTCCCTATCCCCA
TCCAAATCCTTTTCCTTCCTTTTCYCTCAGGTAGATGATGAAATAGGCTGTTATTTTTAGCGATGGCACAA
ATGCTCCCAAATGGAGGATAACGTGCCTCTGGRSCCGGCCTTCTGATACCTGATACCTATCATTASTAAC
CTTCAAAATARYASYACATTTTATTGAGATCTTTCTGTAGCTCTAGTAATAAAACTTTTAATGCAAGTAC
AAAATGCGAAAGTACAAGTAATTGTCTCCYTWAGAAAGTCAAAATGATTGACCTATTGGAGACTGTGAG
GAATAGAATTGTGGAATCAAAGTGAAAAAAGAGTTATTGAGGAAAAAACACTTTCRAGAMSKMTGGCY
AAWSAAAACGTAACGTATCTTGAAAACTAACGGAGTTMYGAATTTGGCGTCTTCGACAAAGTTRTWCA
GAACAGCATTTTCTACAACTCTTGTGAAGACATCAACCTTGTATCTTGAAAAATTAAARAAGTAAATTTT
CTATCTCACTTTTAGGGGGGATTGATCGTTTTTACCWRYAGGTTAAAARRAGCACGTTATTGTAGTGAAA
AACTTCCTCGRRRACACTGAWRWTGCTAARCGACATGGTTAAATCAACAATTGTGCCATATAGTAGTTG
GATTCTTTGCGTGCARTATAAAAAGAAATCTTTAAAACATCATTTCGTAAAAATYACCGTTTTCTCGAGA
ATGGTTTGATGGATCTTCGTGAAATTTTGTTGATAGTTGCACARCCTAAACRCTCCGTGGTCAACCTTTTT
CCTTTTTGACATCGTGCATTTAGCCGAARTGRMGGTCSRAAAACCTCGAAATTTTTCGATTTTTCAAAATT
TCAAATCAATGCCCCTCYCCCTCTCCCAGGAGGGRGAGGTCAAAACTTCCCGGATTTATGAGTTTTCTAG
TCAGGGMATAGTMAGAAAAGGTGAAATTTAGAGCTGRRCAAAAAAAYWTKTMRGACMRWGTTAYYTA
RWGAARRACTCTAGTGAKACTGTTATMYGATAAATCTAGCAATTTTCGGGATTTTCTCGGCATAAARCT
GTCCCATGTCATRTGAGATGTTCAGCATAAGCTGTCCCATGTTGTAATTTTCRTCCCWTAAGCTGTCCCA
TGTCGAGTTTTGCRGCATAAGCTGTCCCCATAKTTGAATTTTACAAATGATATCAGGWTTTCCGGAAAAT
GTTCAGGATTCCCTCAGAGATGCTCGGACAGCTTTTTATGCGTATAAACTGAACATGATATARCTTTKWG
ARAAAATAATRAACATGGGACAGCTTATGCTRCAAAACTGGAACATRGGACAACTYATGATGACAAAAT
TGAACATGGGACAAATTGACTTGATGTTGTTTTCATGGAGTTTTCTRRACAAAGTATACTAATCGAGGAA
GTAGATTCTGAAAGAATATGCGAAAATAGAMCYTTTGATGATAAAATGTCAAWMTCGCTYAAAAAGTA
ACATGGGACAACTATGCTGCACACGGCAGTCTAGCACACATGAATTGCACCGGAASCTTTTCTCAAKTGC
TCCAATGCTGCGTCRKCAWTCATTCACTGCTACCGCGRRGATTATRCGGAACACTGAATGGTCGGATAR
ATCRTCGACCACCTATAATTCCACGAAGGMKYACAAGAAGCTCCGTCGGACYATCAGCAGCASCAATTA
KTTACTATTGACAATTAGTTTGCCATTTTGACAGGTATAGCCCCGCTAAAAGTATAGCTATAAATATGTT
ATTTTGGCGTAGTCTAAATGTTGAAAACAGAAGAGTGAGGGTATGTTGAAATGATGTTGAYTKAGACGT

ATTATAAATACTTTTCCGTTATTGATTGCAGARRTTGCTCTCAGRGGTGCAAATTTAACGTATTTTGAACT
TGTTTCAAAAAGCTTGACGTGTGAAGTTTGGTTATTTAAGAAACGTCTTGTAGATTACAAKTGCCGAACA
GATATACAAATTAAAAAAAAATGGAACAAAACCCACGCTTTCATTGCTTGCTCGAAATTATTAARGAAT
CCTRCTTCCAYCAGTTTAAAAACACTCCCGAACTTTRCTTGAGTAATAATCTGTAGTTAGCCAGAGGTTC
CAGTTATTCAAAAAAGTAAATATTACGATTGCAACATTTTTGAACGGCACTGTAAGCCAAKTGGAAARG
CTGCCATACTGCTCACACCGTAATTTGGAAAAAAATAGAAAATTTAGAAAAAAAATGCACCATTTGATT
GATTCAATCTGAATATTTCAACGCAATTCAGGGGCGGTTGAATTTAAGATAGGTAAAATCATATGTTTCA
CCATAAATCTTAAATCGCAATCAAAAAACACTACCCTTCCTGAAAATAAACCATTTRMRTAAATTGTTAT
TAATTTGTTGTAGAAAATATCCCAAGCAACACGACTTGTTTCAAAAGCCAGTACCAGCAACATCAGTGC
AATTTATTCACTGTTCAAATTAAGGACAACAGAACACAATTGCTTCTTCTWYGAAGTACGCAAAAAGCG
YRRAATTKYAAATCGTTATGCAACTWRAKMKGGGAATGATAAAAAAATGAAAAAATATATTCAGACTC
GAACCATGGACACCAGGAGTATTAACCATTCACCTTACATACTACAYAGAAGCTCTGTGAAAGATTTGC
TGCTCAATRCCATAAARMYRCTGAAGTTACGCGTTRCTTCCRTTGTATGMMRTCTTTRCCRCAAGTTAGA
RAGCTGTCAAAATGAAAWGRCRCGCAAGTTTTCCGCAATACATTWAAAMCCTAATCTRRACAAACAAA
CCAGAGTAAGATGTTTCATGTGTGTTACATAGCACATTTAATGCAARCTGACTGTATTGCCACTTGTGAG
GAATATGTAAGCTCCGCCTCTATAAATCGATGTTAAATACGATGTTATTTGTAATTCCTATGAARYAAAG
CTGCARCTTAACRWTATTTGGAGTTTAAATGCAACKCAACTGACAAGATGGAAGTTGCGTGCGCTTTTAT
TRCARCTCTTTTAGACCAARGCATAGAAAACCACATTTTGGATGATGTTGCAGGTGCTRCTTGGGATGRW
YTTTGTCTTCATCACCAGAAGAGAACAATTTTGTCRCGTTRTTRGAGTCATAAAATTATATGTAATATATT
TTTYCTGTCAGTCGCTTTACGTAGTCTATATCTTCATTATGAGATGATATTTATATTATTATATGTAGTTC
GTTATAAACAAGGCTCTCGCAAGTTATCATCCRCTATCAATTTTTCCAATTTGAACCCATGCATGTGGTAT
GTCATATAATAGTATTTTACAAATCGTATTACTAAAAGCAASMARCTTAGTTTCAACCAATAAATATATT
TRTWMTATTGAGAGTGTTTTCCAAGTATCCATTGACGAACAGTAACTTTGTTAATGCATGAAAGCTAAA
TGGAAAGCAATTATTARTACTATAAGAGTATTARGGATACATCTTTATATAACCTTTAAAATGTTTTATA
AGCTMKMCATTTTTTGCGCTGTTTAGKKMTATAAGTATCSCATACGACGTTAAAAATGCAAGGAATAAA
ACAATCGAGTTAGTTCGAAACYKGAATTTTTAGCTWWCTTACTTATATAACAAGCTGTGTTACTTGGGG
AGCAYGTTCGATACGATGGGCTGCTGCGGTGAACAACTGGTTCGATGACRGTRCRAGAAGATTTTGGRW
AYAAAAGAACGCAGCGAGRGGCGGTCRTGCTGCAGCTWYRGGACTCCGTAGTACGTGGAACGATATAA
AAGAACGCGGGAAACAGCGAATCCACGTCTTCCGGGAGAAAAAGCACCGCTTGGAAGTGTCGAAATGT
GCATCGTCTACCACTACACACATCCCAAGCGATAGTGCGCACCTACGTGTGCACCTACCTTGCGCCCCAC
CGCAAGTGGAGAGAGAASACAAAGGCGAGKTGTTAATTGAACCTATCGTTTTTGTTTGATGAAGGATTT
GCACCGGTGGTGTCATGCTACACGATAAGTTTTYRTCTGCAACAACCAGGCAGCTCATCAAAGCAGGCA
TAAATTAGTTTGGTGMTTARMRATGCAGATGTAAACAACAGAAAATCGAAGAAAATTTGGAGTGGAGA
AAGCCCSGGAAACAGTGCTGGARGATTCAAGCAAGGAAATGGTGGACTTCGGTGAAGAGTTCTCCCGAC
GGATTTATAGARGWTYCCTTRCRGGATCTATCCAYSAAATTTTCCCCGAAAAGACTGCWGYAGAACATT
TCCAAGCGGAAATTTRTCCTGATATTTTCTCCAAAATGTCTCACCGAGGATTTTCCAGTGAATTCCTCTAC
CGARAATTCCTCCAGAATTTCCCCRTAAATCTCCYCCTGGGATTTCACCGGAGATTCCACCRGGAGCTCC
TCCCGGGGKTTTCTCCACGGAATTCCCTCCATAATTCCGTCGRGGAATCCTCCTGAYWKTTTTCRGGAAA
TTCASCAGGAATTTCTTCCRGGATCCACTCAGGAATTAATCCGGAAATTCTCYCAGCATTTTTGTTTTCGG
AATTCTCTAAAGAATACCTATGTGGAAAKTCAGGAATTCCACCCAGGGATTTCTCAAGGAATTCTACCW
GGGGTTCCTCCKYSAATTCCAACCGAAGATATTTTTCAGGGATTCATCCAGRAGTTTCCTCTAGAAATTT
CTTCACGAGATTTCYCAGGAATTTCACYGRGGACTCTTCCAGGAATTCCCATGGGGATTCCTCAAGGAGT
TCCTTTGGATATTTTCARCATTTTTTTGAGGTGATTCCTCTGAGTGAAAACTGAATTTCTCCAGGTGCTMC
TMRAGGAATGTTTTCTGGGATTCCTCTCGGAATATCACCRGGAACYYCAACTTTGAMAATTCTCAACGG
GACACCTCCAAGAAGTCCCACTAAAGGTKTTCCTCCRGGAATTTCGGAGATGAGGAAKTCCACRGGTCA
TGTCRRTCGATCCTGAGCGACCATTTCAAAAATATCTGAAACTTTRCWYWGKTTTCCAATTTCATCTAAA
TCGTCATTTTTCGATATCAAACCTTCATATTCACTCAMAGYTAACTTTTCAGAAGGGTGTATRCTAAAAC
AGCTCAAAAATATTCWAAAAGCTGTACRGCAAGAAAACGGATTTTTCGATTGTTRTATAGATTTTCAAK
MAAAGTTAGATAACTAAATGATGATTCCTTAGAAAATATGCACACTGTAAAAAATTTATTTTTTTTACTT
AAAATATGATTTTTGTCAYAAAAACTCAAATATCTCAAAACCCTATCTTTTTTACCAACGTTAATTTTTKG
GAAATAGCGCYTTTATATCAGCTATCTACACCATAAGAATTTTGATGATGGTAAACTGATAAACAAAAA

AGTTATGACATTTCAAACATTTCACAATTTTTACATTTAGTAACAAAAAAAATTTTTTTCTGTGTAAATTA
TTTTGAGAATTATTTTTTGATGCTGATTTYATTGTAAARKYTACCGCCTRAGATTAAACAAGTTGTTTTTC
ATGAAGTACTTTTGTTTGAATTATTCWKMATTTCTATAGTATCTATTTGAAACTTRGAGCGCGATCGAGT
GTTTTGATCAAAAATATTGARAGTGTCATACTTTWTATWRTACGTGACTGGTGAAAAAATCCCTTGATA
ATGGTGTTGAAAAGCTGTTGATTATAAGAAAAATGGAATTAAACTTATTTTTAAGACAAAAGCTGTATA
ATAAAGTTTTATATACGCGTATWCGTCTAGTTTATCTTCAAAAAAATACCTCTTAGAGAACAGACTTTTA
TGATCGGAAGAATGCGAGTAACTTCAACAACAACAAATGCATGAGAGTTACCTACCACGTGAGAAATCC
ATCGCCCAGTTCACACTACCCCCACACCTGGTGGAAATGTTTCAYGAAATACTGCGATGTGCAATATCGT
CATCCAGAAGGCGCTAAKTARTGAAACGTCAAACGCAAAGAAAAATGATGGGCACTCTTCTAGKTATGA
WAACYACAACCAAGATTCAAAATGATCGTTAAARKSRTGGTCARTRRAAAATTTCGTCAGTGTTACRTTT
GTTTGTCTGTATACATACCATGACAATGCTCACGAAAAAGCTAAAAAAAAAAAAAATACTACCGCTATGG
ATAKTAAATAGTAAATTTTTCTTCAGCCAAGCARSCAACACCAAACTAGTCAGTTAAAACTAATAGAGTG
ATTTTGTTTTATTATAATCTAACGAACAAGCATGAGMCAGTCGCTTTCTCAAAAAGGTCTTCCAACTCAA
CTCTCTCTTGTAGACCGTTTGATGAAAAAAAAAATGATCAAAGGAGTTACGAAATCTCACCGAAAGCASC
ATAGATAATCTAGAAAGAGACTGGTTATGCTSCAAATTGAATCCAAATTTACGCATTTGCACGTCCTGCC
GTCTAGACTTTGTRSAARCRRGCCATCTGTATATCATCAGTAAAAGCAGGGGCGTAGGAAGTTCGAAAA
CGACAACAACTGAGAAATTGCCAGAAGTGCTAAGTGCAGAGAGTCATGTCACCGTACCRTCAGCAGYGA
CATCAGTTTAAACAATTAATCACGCCCCTTTTCAAAAATGCTTTGWKATATACTTCAACATCTATRCCCA
TTTYMATATTTTTAACGTTGYWRARCACGCTTTCAACATTCTRTTTGAAGAAGGAAAACACTTGWYMTC
AAATGTAACAGCAAAATAATGGAATAMWYGACTAATGCGCAGGAAGCTGTGGCTKTKMWAAAATCGG
AACATTACTGTACATATAATATACATAATACATAATAAAAACRGCTTGTTTTRCTCACGCAAAGGCCATT
AACAATAAAATCAGCATCAAGCTGCGATTTCMGSCGAAATAACACTAAAAAAAAAAAAATTATTACTAAA
TGTAAAAATTGTGAAATGTTTGAATTGTCATAACTTTTTGTTTATTAGTTTATCATCACCAAATTTTTATR
GTAGATTGCTAATACAATGGACCGTTTTCCCTAAAAAAATTGCGTTGGTAAAAAGATAGGGTTTTGAAAT
ATTTGAGTTTTTTGTGACAAAAATGATATTTTTTAATGTTAAAAAAGATTTTTTTTCACAGTGTATATTTT
CTTARRRATCACCATTTAGTTATCTAACTTTGCTGAACAATAAAATCAGCATCAAACTTGCGATTTCTGA
CCGAGAATAATTTACACAGAAAAAAATATTTACCAAATGTGAAAWTKGYRAAATGTTTGAAATGTTATA
ACTTTTTTGTTTATTAGTTTACCATCACYRAATTTTTTATGGTAGATTGCTAATACAATGGACCGTCTTCC
TMWAAAAATTGCGTTGGTAAAAGATAGGGTTTTKRGATATTTGAGTTTTTTGTGACAAAAATGATATTTT
ATTATTAAAAAAAGATGTTTTTTTTTCACAGTGTATATTTTCATAGGAATCACCATTTAGTTATCTAACTT
TGCTGAAAAATTCATAGCAATCRRWYGAACCATTTTGAAARCWKTACCAGATTTTTGAATATTATTGAA
CCATTTTCACATACACCCTTTCGAAAAGTTGAGTCGTGRTTCAAAATAAAAATTTGATATCGAAAAATGG
CGATTTAGATGGAACTGAAAAACTGYWRARAKTTCAGTTCAATAAGAAARTCATGAATTAAAAATTTCC
TTAATTTTGATGCTGTTRCTTKGAATCGCTCAGAYYWCTCAGGACATCCTTCAGGGATTCTTTCAGGAAG
CCCTCSGGGATTCTTCCTTGTTTTCCACCAAGAAATCCTCMAAAAATTCCCCGGAGATTCCTATAGAGAT
ACCTTCGGGGAAATCTACTGGAATTCACTCCAGGATTTTTTTCCAGGAATGCCTCTGTGGATAACTCCAG
GAATTTCTCTACAGATATTCACTCMGRYATTCCTCGAGTAATTTGTTCGAGGATTCTCCCAAGAATTTCT
CTAGGGATAACTCCARGAATTCCTCACGGGGACTCCTCCTAGGGTTCCTCTGAAAATTTCTCAAAAAATT
CCCAYSAGGAATTCCTCCCACGAATTCCTTCRGGTAGTTCTCCAGCATTATATACTGAGATTCTTCCRGGA
ATTCTTCCTGTTATTTCCCCAGATATTACTCCGGGAATCCTCCATAAATTCCTCCGAGGATATCTCCAGGG
AATCCACCGGGGATTCTTTCAAGGGACTCCACCTGGGATTCCTCCAAAAACTCTTCCTCCCGARAATCCG
CCAGGGTTTCCTCTGAGAATTTTYCGMGAGGATTTCTCCAGGRGTTCCACTGGAAAGTCTTCTGGAGATA
ACTCCAAGATTTCCTCCGAGGATTTCACCGGGGATTTCTTAAGAGCTCCTCGGAGAATCACTCCASAAGT
TTTCTCCGAGGATTCAGGGATTTCATTAGAGATTTTCTTCAGGATCGAAACGTTGGCAATGGTATAAATT
TATCAACGCTTTTTTTCGTGACTGGMAAGCCGAAAAACTCATAACTCTCTTCAGGTATTCCTCAACTTGA
AGCAATCCTCATTGGAAKTTCTGAAGTAATCCCTGGAGAAATTCAAGGAATAATTCCCGGAGGAACTTC
TWMAGAAATCCCTAGAGGAATTCTTGGAGTTATCTCCAGAGGAATTGCTGGAACGATCCCTGGAAATGG
CTGAGATRCCCRGAGGAACTCCTTGGAGGAATTTCTGGAGAAATCCGCGGTGTAATTTCTRGAGGAGTC
GCTGGAGGACTTCYGRCGGAATCCCCAGTGAACACTMSAGGAGTCCCCAAAGGGAATTCCTGGRRATGT
CTCRGAGAAATTCGTGGAGGAATTCCCCGAGGAAATTCCAGATATTCTTAAGAAATTATTCCAGGAATTC
ATTTAGAGGTTCCTGCAGAGAATCCTTCTAAACATGACTTCAGAACTTCTTMRGGGATTCCTCCAGGAAT

TCCTTCAGAGATTCCTTCGAAATTCTTTTGAAGATTCCTCCCAGGAAATTCCTCCGGAGATATCTGCAGG
ATTTTATTCAAAGATTTCTCCAAGAAKTACTCAGARTYCCGGATAAGGAGAGTTCGACTGTTGCGATCTA
TAGAGCTATTAWTTCTGTRTTRSAGTCTTGGGGAACTAAATCGATAGTTTAAGATTTTTCCCTTTTTTTTC
CTAAAACTATCAKTTTAATTCCCCATGCCGAAGTGCAGTAGCTCTCCAGAATTCCTCCAGAAATTCCTCA
AGAACTTTCTTCCGGRGATTCCTTCAAGGGATTCCTCCAGGAATTCTTTCATGGGTTTTCTCCATAAATTC
RTCCGRGGATTCCTCCAAGARATCCTGCKGGAATTCGTTCAGATATTTCTTCAGGGAATCCTCCAGAAAT
ACCTWSCAGWGATTCCATCTGGARTCCCTTTAGAAATTCASMTCARRRATTCTCTCGAGGRAGTTCCTCC
AGRRATTCAGCWYKAGGATCCTATCAAAGAATTCCTTCMGRGATTCCTTCTGGAATTCMTTCARGRAKT
YCTCCAGAATTTTTTCAGAAATCCTTCCGGAATTCTTTTGRRRATTCYYCAGGAATTTCCTCCGRGGATAC
CTCCACGAATTTATTCAGAGATTCCTCCAGGAACTCCTTCAGGGAATTACTCAATTCCTCCCAGGGWTYC
CTTCARGGATTCCTGCAGGAATTT
